# Lymphatic activation of ACKR3 signaling regulates lymphatic response after ischemic heart injury

**DOI:** 10.1101/2024.12.04.626683

**Authors:** Laszlo Balint, Shubhangi Patel, D. Stephen Serafin, Hua Zhang, Kelsey E. Quinn, Amir Aghajanian, Bryan M. Kistner, Kathleen M. Caron

## Abstract

2.

**Background:** Ischemic heart disease is a prevalent cause of death and disability worldwide. Recent studies reported a rapid expansion of the cardiac lymphatic network upon ischemic heart injury and proposed that cardiac lymphatics may attenuate tissue edema and inflammatory mechanisms after ischemic heart injury. Nevertheless, the mechanisms through which hypoxic conditions affect cardiac lymphangiogenesis and function remain unclear. Here, we aimed to characterize the role of the adrenomedullin decoy receptor atypical chemokine receptor 3 (ACKR3) in the lymphatic response following ischemic heart injury.

**Methods:** Spatial assessment of ACKR3 signaling in the heart after ischemic heart injury was conducted using ACKR3-TangoGFP reporter mice. Roles of ACKR3 after ischemic heart injury were characterized in *Ackr3^ΔLyve^*^1^ mice and in cultured human lymphatic endothelial cells (LECs) exposed to hypoxia.

**Results:** Using the novel ACKR3-Tango-GFP reporter mice, we detected activation of ACKR3 signaling in cardiac lymphatics adjacent to the site of ischemic injury of left anterior descending artery (LAD) ligation. *Ackr3^ΔLyve^*^1^ mice exhibited better survival and were protected from the formation of acute tissue edema after ischemic cardiac injury. *Ackr3^ΔLyve^*^1^ mice exhibited a denser cardiac lymphatic network after LAD ligation, especially in the injured tissues. Transcriptomic analysis revealed changes in cardiac lymphatic gene expression patterns that have been associated with extracellular matrix remodeling and immune activation. We also found that ACKR3 plays a critical role in the regulating continuous cell-cell junction dynamics in LECs under hypoxic conditions.

**Conclusions:** Lymphatic expression of ACKR3 governs numerous processes following ischemic heart injury, including the lymphangiogenic response, edema protection and overall survival. These results expand our understanding of how the heart failure biomarker adrenomedullin, regulated by lymphatic ACKR3, may exert its cardioprotective roles after ischemic cardiac injury.

## 3. Introduction

Myocardial edema, tissue inflammation and resultant fibrosis are frequent consequences of ischemic heart injury. Lymphatic vessels drain excess interstitial fluid and modulate immune responses by trafficking leukocytes. Recent studies reported an expansion of the cardiac lymphatic vasculature after ischemic cardiac injury^1,2^. Furthermore, multiple studies suggest that this lymphangiogenic response promotes cardiac edema resolution, and immune cell trafficking, thereby bolstering cardiac health post injury^3–6^. Although stimulation of cardiac lymphatic responses has recently emerged as a potential therapeutic approach to treat myocardial infarction^2,4^, the underlying hypoxia-induced mechanisms promoting the lymphangiogenic response after ischemic cardiac injury are not fully understood.

Levels of adrenomedullin (AM, Gene: *Adm*), a circulating peptide hormone required for lymphatic vessel development and function in mice and humans, are significantly elevated in patients with acute myocardial infarction^7^. Importantly, AM signaling promotes lymphangiogenesis in the setting of cardiac injury, leading to resolution of myocardial edema and improved cardiac function^5^. Atypical chemokine receptor-3 (ACKR3) temporally and spatially regulates levels of AM signaling by sequestering the peptide and promoting its degradation^8^, through subcellular endosomal recycling that is dependent on arrestin recruitment and receptor activity modifying protein 3 (RAMP3)^9,10^. Similar scavenging effects of ACKR3 have been established for C-X-C ligand 12 (CXCL12), CXCL11, PAMP and opioid peptides, among others^11,12^. Both AM and ACKR3 expression are induced by hypoxia^13,14^, and their shared roles in regulating heart growth and placental development underscore the functional relevance of AM-ACKR3 signaling axis in development^15,16^. Characterization of numerous genetic mouse models reveals an expanding role for ACKR3 in cardiovascular disease, inflammation and ischemic diseases^17,18^.

With this information in mind, we hypothesized that expression and signaling of ACKR3 in cardiac lymphatics may contribute to the temporal and spatial regulation of AM and thereby modulate lymphangiogenesis and immune cell clearance in ischemic cardiac injury. Thus, we utilized cultured cells, state-of-the-art imaging, reporter- and conditional mice and genome-wide expression analyses to characterize the role of ACKR3 in regulating the lymphatic response after ischemic heart injury.

## 4. Methods

Full methods and all supporting data are available within the Supplementary Material.

### 4.1 Mice

Ackr3-TangoGFP transgenic mice were generated by the Animal Models Core Facility at the University of North Carolina at Chapel Hill (UNC-CH). To attenuate undesired H2B transcription, mice were kept on doxycycline-supplemented chow (Envigo, TD.01306). To turn on H2B transcription, mice were kept on normal chow for 7 days before LAD ligation. *Ackr3^fl/fl^* and *Ackr3^fl/fl^* ; *Lyve1^Cre^*(*Ackr3^ΔLyve^*^1^) mice were maintained on a C57BL/6 background. Prospero homeobox protein 1 - GFP (*Prox1^GFP^*) mice^19^ (generously provided by Young-Kwon Hong) were maintained on a 129S/SvEv background. For all experiments, 4-to 8-month-old animals were used. Ischemic cardiac injury experiments were conducted on male mice. All animal studies were approved by the Institutional Animal Care and Use Committee at the University of North Carolina Chapel Hill.

### 4.2 Surgical induction of myocardial infarction

A left lateral thoracotomy was performed between the 3^rd^ and 4^th^ rib on anesthetized and ventilated mice followed by the ligation of the left anterior descending artery (LAD). Chest wall, muscle and skin were closed, followed by systemic antiseptic and analgesic treatment. Animals that did not survive permanent LAD ligation surgery or died before recovery from anesthesia were not included in survival data and were excluded from further analysis. Ackr3-Tango-GFP mice were euthanized 2 days post LAD ligation, while *Ackr3^fl/fl^* and *Ackr3^ΔLyve^*^1^ were euthanized 3, 6, or 28 days post ligation.

### 4.3 Histology and immunofluorescence

Mice were euthanized with CO_2_ asphyxiation and trans-cardiac perfusion was performed with 10 mL PBS followed by 10 mL 4% PFA. Dissected organs were fixed in 4% PFA for 24h at 4°C, and then embedded in paraffin for sectioning and staining following standard protocols by the UNC-CH Histology Research Core Facility. Immunofluorescent staining was performed using the following primary and secondary antibodies: rabbit anti-LYVE1 (1:200, Fitzgerald, 70R-LR005), goat anti-CD45 (1:50, Novus Biologicals, NB100-77417), goat anti-F4/80 (1:50, BD Pharminogen, 565409), donkey anti-rabbit Cy3 (1:200, Jackson ImmunoResearch, 711-165-152), donkey anti-goat Cy5 (1:200, Jackson ImmunoResearch, 705-175-147).

### 4.4 Whole mount immunostaining

Mice were euthanized with CO_2_ asphyxiation and trans-cardiac perfusion was performed with 10 mL PBS followed by 10 mL 4% PFA. Dissected organs were fixed for 24h in 4% PFA at 4°C, followed by routine whole mount immunostaining with goat anti-LYVE1 (1:200, R&D Systems, AF2125-SP) primary, and Alexa Fluor 488-conjugated donkey anti-goat (1:200, Jackson ImmunoResearch, 705-545-003) secondary antibodies.

### 4.5 Tissue clearing and whole mount immunostaining

iDisco protocol was performed as described previously^20^. Samples were incubated with rabbit anti-LYVE1 (1:500, Fitzgerald, 70R-LR005) and chicken anti-GFP (1:500, Aves, GFP-1020) primary antibodies for 14 days and incubated with Cy5-conjugated donkey anti-rabbit (1:200, Jackson ImmunoResearch, 711-175-152) and Alexa Fluor 790-conjugated donkey anti-chicken (1:200, Jackson ImmunoResearch, 703-655-155) secondary antibodies for 10 days. For imaging, a LaVision Ultramicroscope II (LaVision) light-sheet system was used. Images were analyzed with the Imaris 9 (Oxford Instruments) software program. Relative lymphatic GFP signal was calculated using the following formula: Relative lymphatic GFP signal = [N(GFP)_LYVE1+_/ N(GFP)_LYVE1-_]_injury site_ / [N(GFP)_LYVE1+_/ N(GFP)_LYVE1+_]_uninjured tissue_.

### 4.6 Microarray analysis

For transcriptomic analysis, LECs and macrophages were collected from *Ackr3^fl/fl^* and *Ackr3^ΔLyve^*^1^ mice at baseline or 7 days post LAD ligation. Mice were euthanized with CO_2_ asphyxiation and trans-cardiac perfusion was performed with 10 mL PBS. The ventricles were then minced and incubated with 2mg/mL collagenase type 2 (Worthington Biochemical, LS004176) in Hank’s balanced salt solution (Gibco, 14025092) at 37°C to make a cell suspension. CD68+, CD68-/LYVE1+ and CD68-/LYVE1-cells were separated by a series of pull-downs using Dynabead Protein G (Thermo Fisher, 10003D) magnetic beads pre-treated with 1µg/µL anti-CD68 (BioRad, MCA1957) and 1µg/µL anti-LYVE1 (Fritzgerald, 70R-LR005) primary antibodies. Cells were incubated with each antibody for 1h at 4°C. For RNA extraction from the captured cells, Trizol (Invitrogen, 15596026)-based protocol was used, following the manufacturer’s recommendations. Microarray analysis was performed in the UNC-CH Functional Genomics Core using the mouse Clariom S Pico Assay kit (Thermo Fisher, 902932) after QC/QA testing. Microarray data was evaluated using the Partek Genomics Suite (Partek) software program. For statistical analysis of the data, one-way ANOVA was performed, and false discovery rate-adjusted p values ≤0.01 were considered as significant. Gene Ontology analysis was performed using the Shiny GO 0.8 tool^21^.

### 4.7 Echocardiography

Conscious echocardiography was performed using a Vevo 2100 ultra high frequency ultrasound (VisualSonics) on male *Ackr3^ΔLyve^*^1^ and *Ackr3^fl/fl^* mice before, and 3,6,14,21 and 28 days post LAD ligation. M-mode parasternal long axis echocardiographs were analyzed by measuring both diastolic and systolic dimensions of three cardiac cycles from three recordings using Vevo LAB software (VisualSonics). The nine measurements were averaged for each mouse on each experimental day. Stroke volume was calculated using the SV= LV Vol;d – LV Vol;s formula. Cardiac output was calculated using the CO=SV/HR formula.

### 4.8 Cell culture, gene silencing, and hypoxia induction

Primary human lymphatic endothelial cells (hLECs), isolated from human foreskin (PromoCell, C-12216) were cultured in Endothelial Cell Growth Medium (PromoCell, C-22111) 37°C under 5% CO2 and used within 5 passages. Gene silencing was induced using *CALCRL-* (Santa Cruz Biotechnology, sc-43705), *ACKR3*- (Santa Cruz Biotechnology, sc-94573) targeting or scramble control (Santa Cruz Biotechnology, sc-37007) siRNA using Lipofectamine RNAiMAX transfection reagent (Thermo Scientific, #13778030) for 24h following the manufacturers’ recommendations. RAMP3 expression was silenced using lentiviral particles containing shRNA targeting RAMP3 (TRCN0000060981) or negative control human beta globin (HBG) (TRCN0000029094) for 24 hours. To study hypoxia-induced changes in LECs, cells were incubated at 1% or 21% O2 in a Tri-gas incubator for the indicated times.

### 4.9 Immunocytochemistry and lymphatic junctional assay

To assess the effects of hypoxia on LEC junctional arrangement, confluent monolayers of LECs were fixed in 4% paraformaldehyde for 20 minutes at RT, permeabilized and immunostained using rabbit anti-VE-cadherin (1:200; Abcam, ab33168) primary, and Cy5-conjugated donkey anti-rabbit (1:200; Jackson Immunoresearch, 711-175-152) secondary antibodies. Average percent ratio of total continuous junctions length among total junctions length (continuous + discontinuous junctions combined) of a multiple field of views from the same biological replicate was used for quantitative assessment of junction dynamics.

### 4.10 Adrenomedullin response assay and Western blot

Intracellular AM signaling was assessed following shRNA-mediated gene silencing of RAMP3 in hLECs by Western blot using the following primary and secondary antibodies: p-p44/42 MAPK (T202/Y204) (CST, 4370) (1:5,000), p44/42 MAPK (ERK1/2) (CST, 9102) (1:5,000), p-AKT (S473) (CST, 4060), AKT (pan) (CST, 4691) (1:5,000), p-CREB (Ser133) (CST, 9198) (1:5,000), CREB (CST, 9197) (1:5000), GAPDH (Novus Biologicals, NB300-221) (1:10,000), IRDye 800CW goat anti-rabbit (Licor, 926-32211) (1:10,000), and IRDye 680LT goat anti-mouse (Licor, 926-68020) (1:10,000).

### 4.11 Quantitative real time polymerase chain reaction

To characterize gene expression levels in cultured hLECs or mouse cells, RNA was extracted using Trizol protocol and cDNA was transcribed using M-MLV reverse transcriptase (Thermo Fisher, 28-025-013) following the manufacturer’s protocol. Real time polymerase chain reaction was performed using the Fast Advanced Master Mix (Thermo Fisher, 4444557), according to the manufacturer’s protocol utilizing a Thermo Fisher StepOnePlus real time PCR system. For human LECs, *ACTB*-(Thermo Fisher, Hs01060665_g1), *ACKR3*-(Thermo Fisher, Hs00664172_s1), *ADM*-(Thermo Fisher, Hs00969450_g1), *CALCRL*-(Thermo Fisher, Hs00907738_m1), and *CDH5*-(Thermo Fisher, Hs00901465_m1) specific probes were used. For mouse cells, *Gapdh-* (Thermo Fisher, Mm99999915_g1), *Cd68-* (Thermo Fisher, Mm03047343_m1), *Lyve1-* (Thermo Fisher, Mm00475056_m1) specific probes were used.

### 4.12 Statistical analyses

Statistical analyses were performed with the GraphPad Prism 10 software program. Specific statistical tests and groups sizes are indicated for each analysis. P values ≤0.05 were considered significant and are shown in each graph. For microarray data, false discovery rate-adjusted p values ≤0.01 were considered significant. Data represented as mean ± standard deviation.

## 5. Results

### 5.1 Detailed spatial characterization of the cardiac lymphatic network

We performed a detailed characterization of the cardiac lymphatic network by leveraging the power of 3-dimensional light-sheet imaging of tissue-cleared whole mount hearts of *Prox1^GFP^* lymphatic reporter mice combined with anti-GFP and anti-lymphatic vessel endothelial hyaluronan receptor 1 (LYVE1) immunostaining. This approach enabled a full spatial reconstruction of the cardiac lymphatic network (Supplementary video 1). As previously described, the heart is surrounded by a network of readily apparent subepicardial lymphatic vessels consisting of LYVE1-negative collecting lymphatics with PROX1^high^ valves, and LYVE1-positive initial lymphatic capillaries (Figure 1A, Supplementary Figure 1). Further, we noted the presence of distinct LYVE1-positive lymphatic vessels penetrating into the deeper myocardial layers up to 300 µm deep (Figures 1B,C, yellow arrowheads). Notably, prevalent lymphatic structures within the cardiac septum of adult animals were detected on the side of septum facing the right ventricle, but not the left ventricle (Figure 1B,D, red arrowheads). Thus, the use of high resolution 3-D light-sheet imaging revealed deeper-reaching and asymmetrically organized cardiac lymphatics that have not been previously appreciated.

**Figure 1.**
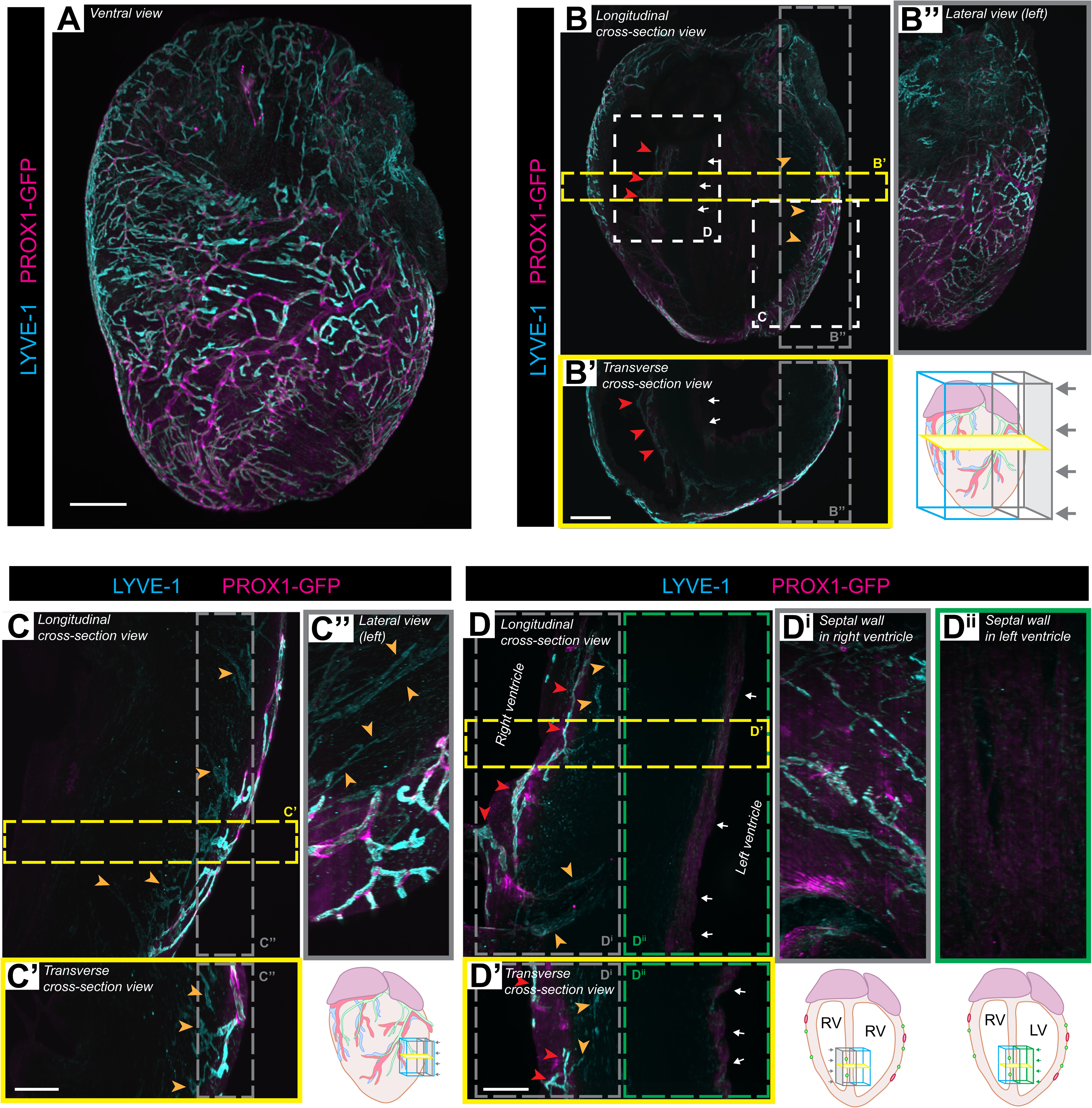
Cardiac lymphatics penetrating into the deeper cardiac tissue layers and asymmetrical lymphatic network in the cardiac septum is revealed by 3-dimensional fluorescent microscopy. **A)** 3-D volumetric view of the cardiac lymphatic network visualized by light-sheet imaging of *Prox1^GFP^* mouse hearts immunostained with anti-LYVE1 and anti-GFP antibodies using the iDisco tissue clearing and staining protocol. Scalebar, 1000 µm. **B-D**) Expanded cross-section views of the cardiac lymphatic vessels of *Prox1^GFP^*mouse hearts. Low-magnification images (**B**, scalebar, 1000 µm), and insets from the left ventricular free wall (**C**) and the cardiac septum (D) (bars, 300 µm) are shown. Yellow arrowheads point at cardiac lymphatic vessels penetrating into the deeper myocardial layers. Red arrowheads point at lymphatic vessels in the cardiac septum facing the right ventricle. White arrows point at the cardiac septal wall facing the left ventricle. Representative images from 3 animals are shown.

### 5.2 Ischemic cardiac injury enhances epicardial lymphatic ACKR3 signaling adjacent to the injury site in a RAMP3-dependent manner

Prior studies reported that both ACKR3 and its ligand, AM are upregulated upon hypoxia in various tissues^13,14^. Similarly, we also found a gradual and statistically significant increase in both *ACKR3* and *ADM* expression levels in human cultured lymphatic endothelial cells (hLECs) exposed to hypoxia (Supplementary Figures 2A, B). To determine whether there is spatial activation of ACKR3 signaling in the mouse heart lymphatics upon ischemic cardiac injury, we generated and characterized a novel ACKR3-TangoGFP reporter mouse model. Briefly, in this model, ligand activation and subsequent β-arrestin recruitment to ACKR3 initiates intracellular H2B-GFP transgene expression, thereby allowing for visualization of spatial localization and activation of ACKR3 in native tissues (Supplementary Figure 3A, adapted from ^22^). To visualize and quantify ACKR3 activity in cardiac lymphatics, we leveraged the power of 3-dimensional light-sheet imaging of tissue-cleared whole mount hearts. Assessment of the H2B-GFP reporter signal in baseline adult male ACKR3-Tango-GFP mice revealed high ACKR3 signaling activity in the endocardium of the left ventricle (Supplementary Figure 3B). Inducing ischemic heart injury by LAD caused an increase of GFP-positive, ACKR3 signaling cells within the epicardial and subepicardial layers of the left ventricle. We found spatial co-localization of TANGO-receptor activity with immunostained LYVE1-positive cardiac lymphatic vessels, while sham operated mice exhibited no GFP signal in LYVE1-positive structures (Figures 2A,B, Supplementary Figure 3C).

**Figure 2.**
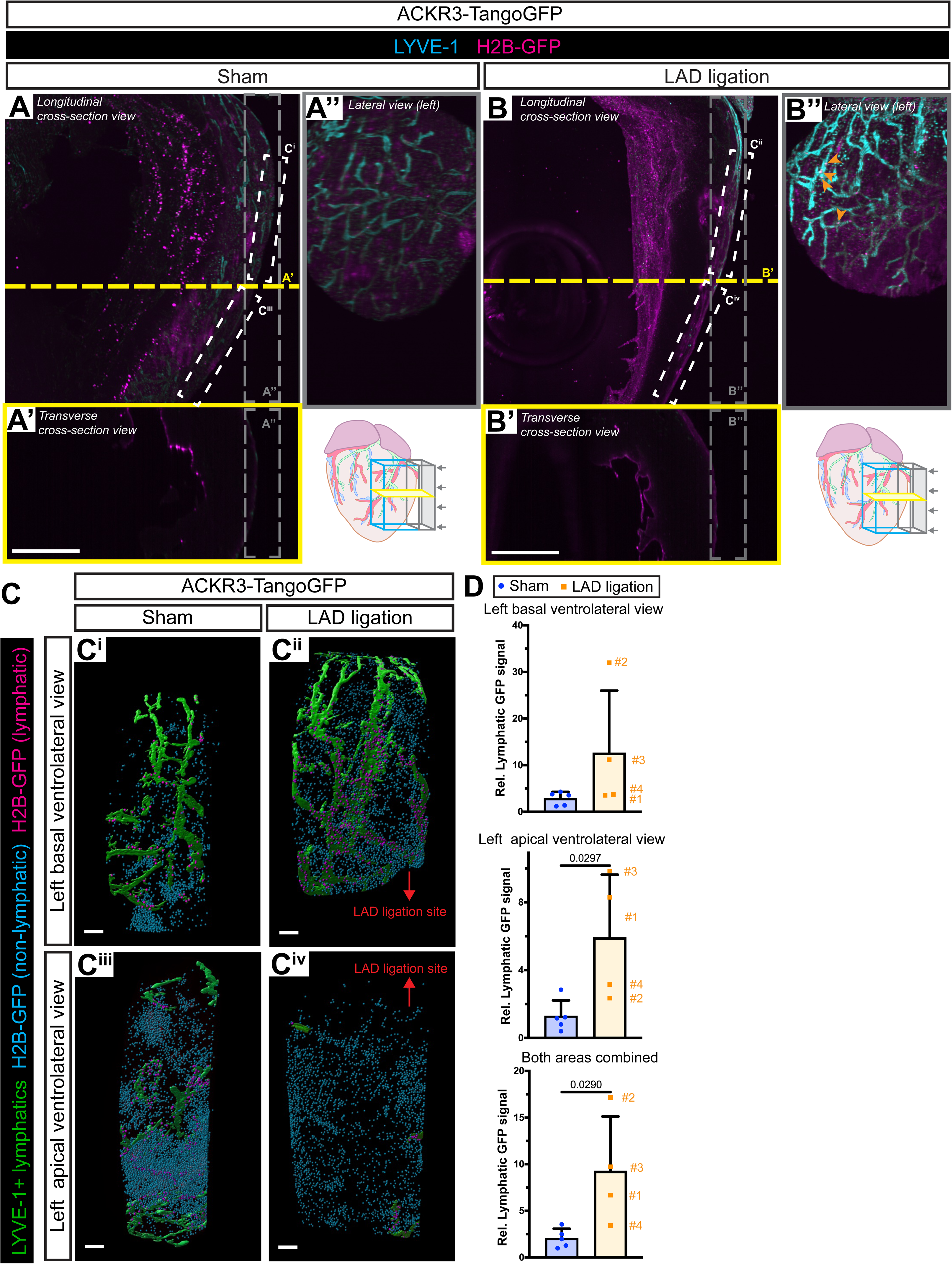
Spatial activation of ACKR3 signaling in cardiac lymphatic vessels upon ischemic heart injury. **A,B)** LYVE-1-positive cardiac lymphatic vessels and ACKR3 activation detected by H2B-GFP expression in ACKR3-TangoGFP mice 2 days post sham surgery **(A)** or LAD ligation **(B)**. Expanded cross-section views are shown. Yellow arrowheads point at H2B-GFP signal overlapping with LYVE1 immunostaining in epicardial LYVE1-positive lymphatic structures in the left basal ventrolateral region of the heart of LAD-ligated mice, adjacent to the ligation site. Bars, 1000 µm. **C)** Rendered pseudocolored images showing LYVE1-positive cardiac lymphatic vessels (green), ACKR3 activation in LYVE1-negative (blue) and LYVE1-positive (magenta) cells in the basal and apical ventrolateral segments of the left ventricle free wall of ACKR3-TangoGFP mice 2 days post sham surgery or LAD ligation. Bars, 200 µm. Representative images from 4-5 animals per group are shown. **D)** Relative lymphatic GFP signal (calculated as [N(GFP)_LYVE1+_ / N(GFP)_LYVE1-_]_injury site_ / [N(GFP)_LYVE1+_ /N(GFP)_LYVE1+_]_uninjured tissue_) in the basal and apical ventrolateral segments of the left ventricle free wall compared to the uninjured apical septal wall facing the right ventricle in ACKR3-TangoGFP mice 2-days after LAD ligation or sham surgery and the average of the values from these two areas. Each symbol represents quantification from one mouse. Mice in the LAD ligated group are identified by numbering. Unpaired t-test, N=4-5 per group.

To precisely map and quantify the levels of ACKR3 signaling in cardiac lymphatics, GFP-positive cells and LYVE1+ lymphatic vessels were defined in 3-D and found to spatially co-localize. The resultant 3-D signaling maps display the activation of ACKR3 in non-lymphatic cells as blue dots and in green-labeled lymphatic vessels as magenta dots (Figure 2C, Supplementary Figure 3D). Induction of ischemic heart injury by LAD ligation resulted in a substantial number of GFP/LYVE-1 double positive cells among all GFP-positive cells in areas adjacent to the infarct zone in LAD ligated mice but not in the corresponding anatomic locations in sham operated mice (Figure 2C). Quantitative analysis of the ratios of lymphatic GFP signal to total GFP signal adjacent to the LAD ligation compared to uninjured areas of the heart revealed that relative lymphatic GFP-positive signal was significantly increased in the infarcted and peri-infarct areas in LAD-ligated mice, compared to the same anatomical locations in sham operated hearts (Figure 2D). This indicates an increased activation of ACKR3 signaling in cardiac lymphatics *in vivo* after ischemic cardiac injury.

We have previously shown that the GPCR allosteric modulator, receptor activity modifying protein 3 (RAMP3), is required for the recycling and scavenging activities of ACKR3 at the plasma membrane^10^. To investigate whether RAMP3 is required for the lymphatic activation of ACKR3 upon ischemic cardiac injury, we crossed ACKR3-TangoGFP mice to the *Ramp3^-/-^*mice and challenged these animals with LAD ligation. Although a weak H2B-GFP signal was detected at the site of LAD ligation, loss of RAMP3 completely abolished the activation of ACKR3 signaling in cardiac lymphatics post-LAD ligation (Figure 3A-C). In addition to confirming that the H2B-GFP reporter is specific to ACKR3 signaling, this data also substantiates the necessity of RAMP3 for lymphatic activation of ACKR3 upon ischemic cardiac injury.

**Figure 3.**
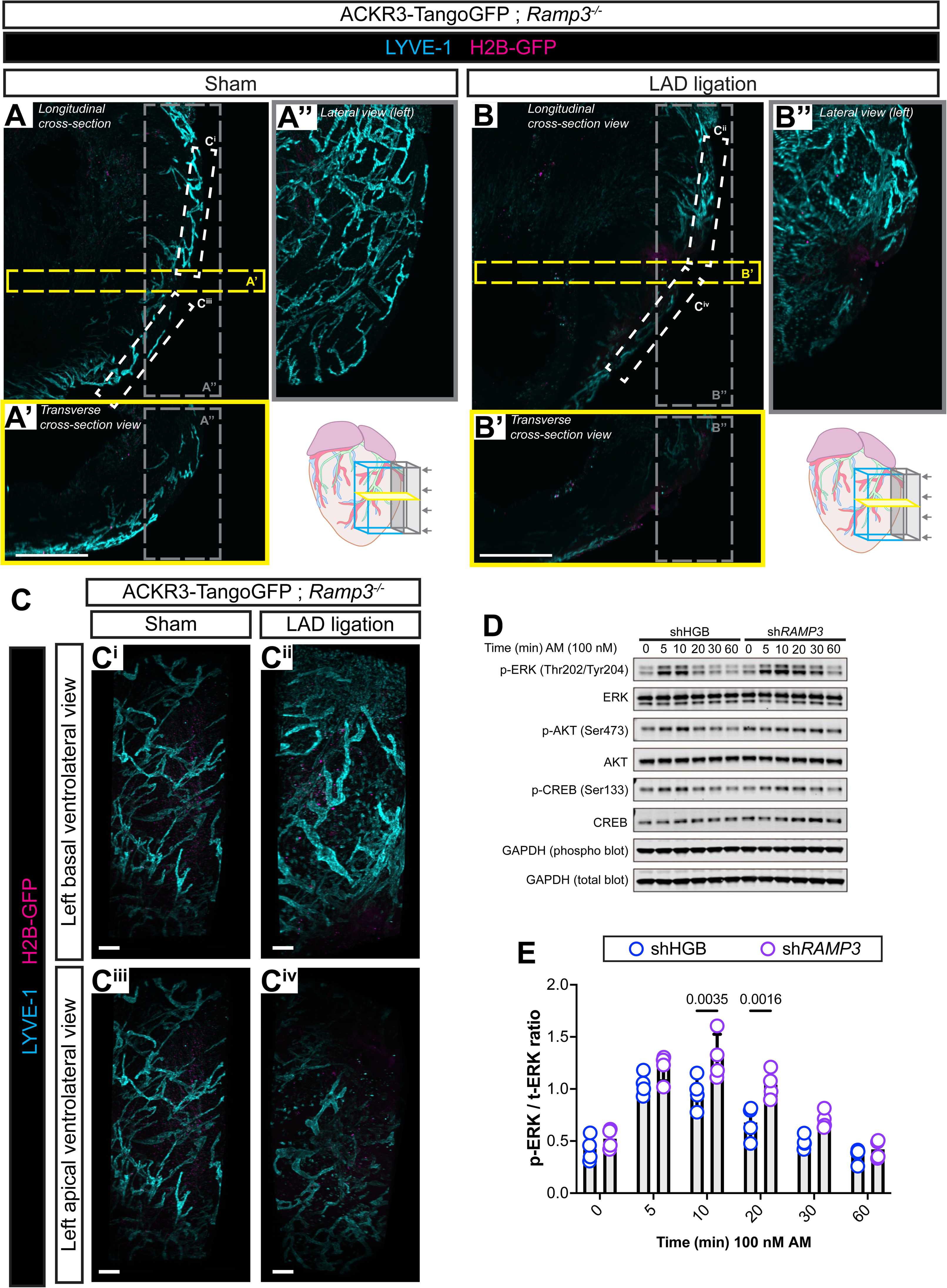
Loss of RAMP3 prevents ACKR3 activation *in vivo* and enhances adrenomedullin-induced ERK phosphorylation in cultured hLECs. **A,B)** LYVE-1-positive cardiac lymphatics and ACKR3 activation detected by H2B-GFP expression in ACKR3-Tango-GFP ; *Ramp3^-/-^*mice 2-days post sham surgery **(A)** or LAD ligation **(B)**. Representative expanded cross-section views from 2 animals per group are shown. Bars, 1000 µm. **C)** 3-D high magnification views showing LYVE1-positive lymphatics and ACKR3 activation detected by H2B-GFP signal in the basal and apical ventrolateral segments of the left ventricle free wall of ACKR3-Tango-GFP ; *Ramp3^-/-^* mice 2-days post sham surgery or LAD ligation. Bars, 200 µm. **D, E)** Representative Western Blot image showing phosphorylation of ERK, AKT, and CREB **(D)** and quantification of p-ERK/t-ERK ratios **(E)** in hLECs treated with adrenomedullin for 5, 10, 20, 30, 60 minutes after treatment with control human beta globin (shHGB) or shRAMP3 constructs. N = 4 per group. Two way ANOVA; Šidak’s multiple comparisons test compared to shHGB.

Among its various physiological roles, ACKR3 acts as a scavenger receptor for the pro-lymphangiogenic peptide AM and regulates its physiological functions. Therefore, we sought to test whether loss of RAMP3 affects the lymphatic response to AM treatment. We found that knockdown of *RAMP3* in cultured hLECs led to a prolonged maintenance of AM-induced ERK phosphorylation (Figures 3D,E), supporting previous findings that loss of RAMP3-mediated ACKR3 recycling can enhance intracellular AM signaling in LECs.

### 5.3 Mice with lymphatic loss of ACKR3 have better survival and are protected against cardiac edema formation after ischemic heart injury

To assess the roles of ACKR3 in the cardiac lymphatic vasculature, we generated and characterized *Ackr3^fl/fl^* ; *Lyve1^Cre^* (*Ackr3^ΔLyve^*^1^) mice, which constitutively lack ACKR3 expression in LYVE1-positive lymphatics. Global deletion of ACKR3 is associated with developmental cardiomegaly, which can be reversed by concurrent genetic reduction of the *Adm* gene^8^. Somewhat unexpectedly, we found that lymphatic-specific deletion of ACKR3 in lymphatics alone leads to cardiomegaly in *Ackr3^ΔLyve^*^1^ mice, evidenced by a significantly greater heart-to-body weight ratio in both sexes (Figure 4A), but no differences in spleen-to-body weight or kidney-to-body weight ratios between *Ackr3^ΔLyve^*^1^ and *Ackr3^fl/fl^* mice (Figures 4B,C). Whole mount anti-LYVE1 immunostaining revealed no obvious differences in the surface cardiac lymphatic density between adult *Ackr3^ΔLyve^*^1^ and sibling control *Ackr3^fl/fl^*mice (Supplementary Figure 4). Thus, expression of ACKR3 in LYVE-1+ cells contribute to the determination of heart size in animals without affecting cardiac lymphatic coverage under physiological conditions.

**Figure 4.**
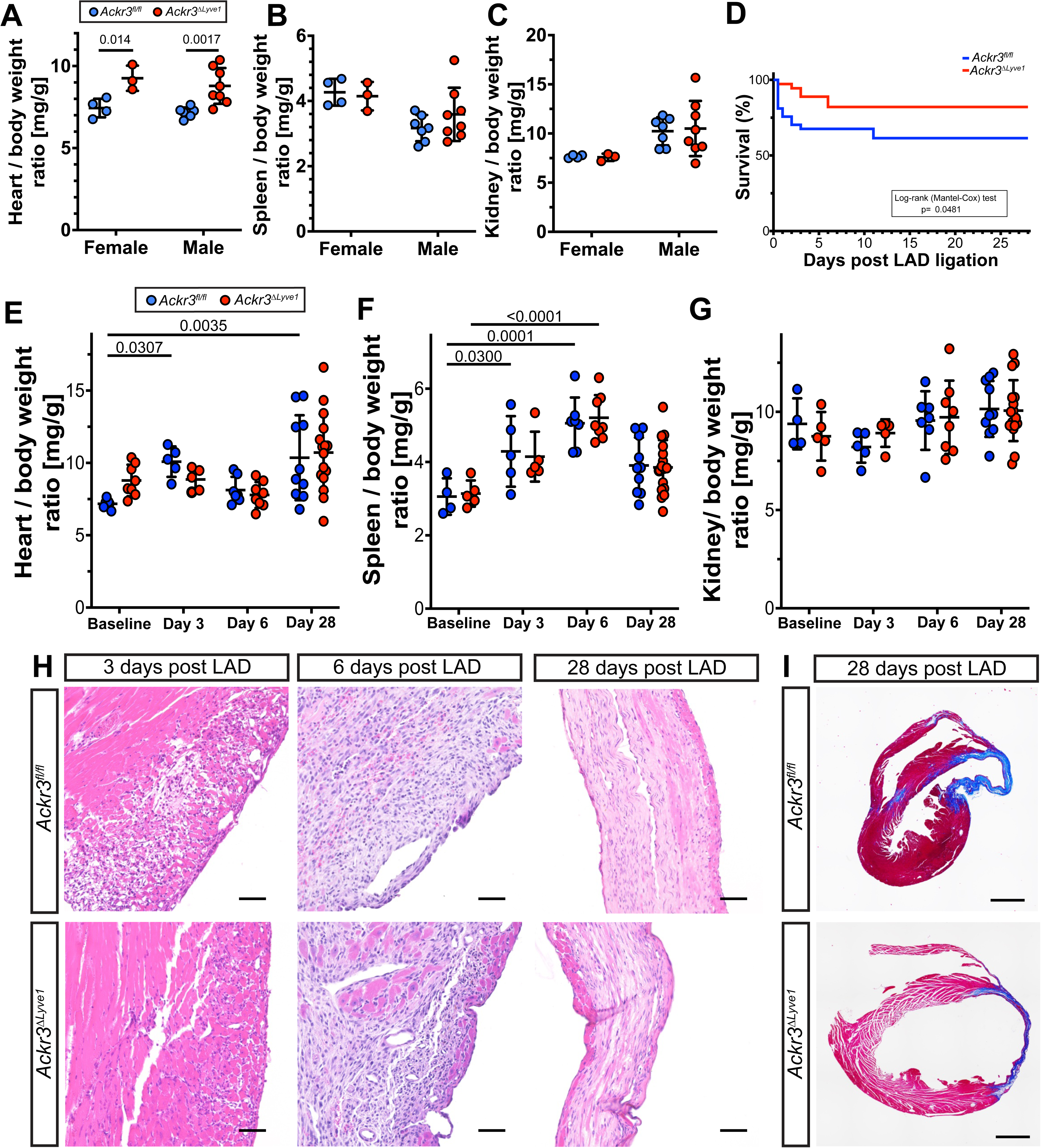
Lymphatic loss of ACKR3 promotes cardioprotection and improves survival after ischemic cardiac injury. **A-C)** Comparison of heart weight / body weight **(A)**, spleen weight / body weight **(B)**, kidney weight / body weight **(C)** ratios of female and male *Ackr3^fl/fl^* and *Ackr3^ΔLyve^*^1^ mice. Two-way ANOVA, Tukey’s post-hoc test. N=3-8 per group. **D)** Survival (%) of *Ackr3^fl/fl^* and *Ackr3^ΔLyve^*^1^ male mice after LAD ligation. N=27-36 per group. Mantel-Cox test, p=0.0481. **E-G)** Assessment of heart weight / body weight **(E)**, spleen weight / body weight **(F)**, kidney weight / body weight **(G)** ratios of male *Ackr3^fl/fl^*and *Ackr3^ΔLyve^*^1^ mice after LAD ligation. Two-way ANOVA, Dunnett’s post hoc test. n=4-16 per group. **H)** High magnification hematoxylin-eosin stained tissue sections of the infarct zone of *Ackr3^fl/fl^* and *Ackr3^ΔLyve^*^1^ mice 3-, 6-, 28-days post LAD ligation. Bars, 50 µm. **I)** Cardiac fibrosis visualized by Masson’s trichrome staining in *Ackr3^fl/fl^* and *Ackr3^ΔLyve^*^1^ hearts 28-days after LAD ligation. Bars, 1 mm. Sections were collected 200µm from the beginning of the infarct zone towards the apex of the heart. Representative images from N=3-7 mice per group.

Given the observed cardiac phenotype in *Ackr3^ΔLyve^*^1^ mice we sought to investigate how lymphatic loss of ACKR3 affects cardiac health after ischemic cardiac injury. We found that *Ackr3^ΔLyve^*^1^ mice displayed significantly better survival post-LAD ligation than their sibling *Ackr3^fl/fl^* controls (Figure 4D). Assessment of systolic cardiac function after ischemic cardiac injury using M-mode echocardiography did not reveal statistically significant differences between *Ackr3^fl/fl^*and *Ackr3^ΔLyve^*^1^ mice (Supplementary Table 1). Nevertheless, we noticed that approximately half of *Ackr3^ΔLyve^*^1^ mice displayed remarkably better ejection fraction and fractional shortening than *Ackr3^fl/fl^* control mice, especially at 28 days post ligation (Supplementary Figure 5). Assessment of heart-to-body weight ratios at different timepoints post-LAD ligation revealed dynamic changes in the relative heart weights of control *Ackr3^fl/fl^* mice, but not *Ackr3^ΔLyve^*^1^ mice, after ischemic heart injury (Figure 4E). As expected, 3 days post-LAD ligation, relative heart weights of *Ackr3^fl/fl^*mice were significantly increased, likely due to acute edema formation, which was resolved by 6 days post ligation. Over a long-term 28-day recovery period, the heart weights of control *Ackr3^fl/fl^* mice were significantly higher than those of uninjured *Ackr3^fl/fl^* mice, potentiating the development of chronic cardiac edema in these mice. In contrast, no significant changes were observed in the heart-to-body weight ratios of *Ackr3^ΔLyve^*^1^ mice after LAD ligation, suggesting that mice with lymphatic loss of ACKR3 are protected against the development of cardiac edema post ischemic heart injury. Furthermore, dynamic changes in spleen weights were observed in both *Ackr3^fl/fl^* and *Ackr3^ΔLyve^*^1^ mice, with no difference between *Ackr3^ΔLyve^*^1^ and *Ackr3^fl/fl^* groups at any timepoint (Figure 4F). No change was observed in relative kidney weights (Figure 4G). Histological analysis confirmed the development of cardiac edema at 3-days post ligation in the peri-infarct zone of *Ackr3^fl/fl^* mice but not in *Ackr3^ΔLyve^*^1^ mice (Figure 4H). Masson’s trichrome staining revealed no obvious differences in fibrosis formation between the two genotypes (Figure 4I).

### 5.4 Loss of ACKR3 in LYVE1-positive cells promotes lymphangiogenesis and leads to an alteration of inflammatory molecular pathways after ischemic heart injury

The observation that *Ackr3^ΔLyve^*^1^ mice were protected against the formation of cardiac edema after LAD ligation suggests that lymphatic loss of ACKR3 may also have an effect on the cardiac lymphatic vasculature upon ischemic cardiac injury. Thus, we characterized the effects of lymphatic loss of ACKR3 on the lymphatic network of the heart after LAD ligation.

First, to define the transcriptomic changes within cardiac LECs due to LAD ligation, we established a method to collect CD68-/LYVE1+ cardiac LECs and CD68+ cardiac macrophages from mouse hearts (Supplementary Figures 6A,B). This allowed us to perform expression analysis on cardiac LECs and macrophages isolated from the hearts of *Ackr3^fl/fl^* and *Ackr3^ΔLyve^*^1^ mice at baseline and 7 days post LAD ligation. Comparison of differentially expressed genes in LECs upon LAD ligation in *Ackr3^fl/fl^* and *Ackr3^ΔLyve^*^1^ mice revealed 1397 differentially expressed genes that were specific to LAD-ligated *Ackr3^ΔLyve^*^1^ mice (Figures 5A,B, Supplementary Figure 6C, Supplementary Table 2). Gene ontology analysis revealed pathways associated with inflammatory and immune responses, as well as vesicle-mediated transport and chemosensory pathways as the most affected by loss of ACKR3 signaling in LECs after myocardial infarction (Supplementary Figures 6D-G). We also compared gene expression levels of these genes in cardiac LECs between LAD ligated *Ackr3^ΔLyve^*^1^ and *Ackr3^fl/fl^* mice (Figure 5C, Supplementary Table 3). Gene ontology analysis indicated negative regulation of reactive oxygen pathways and inhibition of heterochromatin assembly pathway upon LAD ligation in the cardiac lymphatics of *Ackr3^ΔLyve^*^1^ mice post myocardial infarction (Supplementary Figures 6H-J). Notably, we identified multiple differentially expressed genes that were previously associated with tissue remodeling (E.g. *Mmp3, Rab37, Prg4*) and endothelial cell function (E.g. *Angpt1, Itgb4, Saraf*) (Figures 5B,C). Collectively, these data support a tissue environment poised for remodeling in *Ackr3^ΔLyve^*^1^ mice.

**Figure 5.**
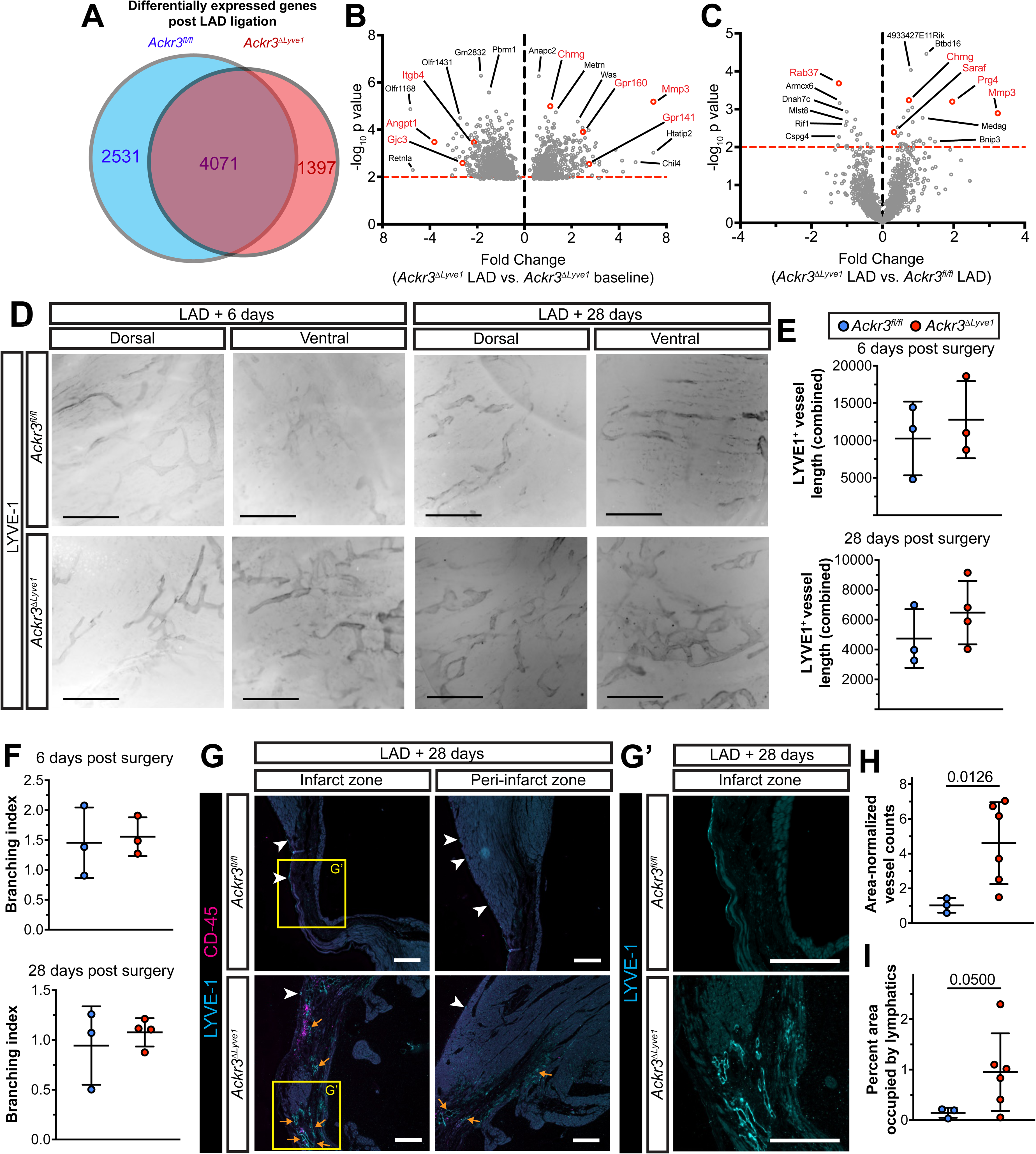
Lymphatic loss of ACKR3 alters ischemia-induced transcriptomic response in cardiac LECs and promotes lymphangiogenic response after ischemic cardiac injury. **A)** Venn diagram summarizing the transcriptomic differences in cardiac LECs isolated from male *Ackr3^fl/fl^* and *Ackr3^ΔLyve^*^1^ mice one week post LAD ligation. **B)** Differentially expressed genes in cardiac LECs upon LAD ligation that are specific to ACKR3-deficient mice. Positive fold change values represent upregulation in LAD ligated *Ackr3^ΔLyve^*^1^ LECs. **C)** Comparison of gene expression levels of identified ACKR3-specific LAD-triggered genes between LAD ligated *Ackr3^ΔLyve^*^1^ and *Ackr3^fl/fl^* mice. Positive fold change values represent upregulation in LAD ligated *Ackr3^ΔLyve^*^1^ LECs. **D)** Visualization of LYVE1-positive cardiac lymphatic vessels of male *Ackr3^fl/fl^* and *Ackr3^ΔLyve^*^1^ whole mount hearts 6- and 28-days post LAD ligation. Representative images of from 3-4 animals per group. Bars, 500 µm. **E,F)** Quantification of lymphatic vessel length **(E)** and branching index (number of branches divided by lymphatic vessel length) **(F)** per field of view. N=3-4 per group. Two-way ANOVA, Dunnett’s post-hoc test. **G)** LYVE1 (cyan) and CD45 (magenta) staining of the infarct zone and peri-infarct zones of male *Ackr3^fl/fl^*and *Ackr3^ΔLyve^*^1^ hearts 28-days post LAD ligation. Yellow arrows point at LYVE1-positive lymphatic structures in the injured tissues of the infarct and peri-infarct zone of hearts. White arrowheads point at epicardial lymphatic structures. Representative images from N=3-6 mice per group. Bars, 200 µm. **H,I)** Area-adjusted lymphatic vessel counts in the infarct zone **(H)** and percent area of the infarct zone occupied by lymphatic vessels **(I)** in male *Ackr3^fl/fl^*and *Ackr3^ΔLyve^*^1^ hearts 28-days post LAD ligation. N= 3-6 per group. Unpaired Welch’s unequal variances t-test.

Whole mount immunostaining of the hearts revealed no significant differences in the subepicardial lymphatic network in *Ackr3^ΔLyve^*^1^ mice, compared to *Ackr3^fl/fl^* mice, 6- and 28-days post LAD ligation (Figure 5D-F). However, immunofluorescence analysis of the cardiac tissue revealed a much denser lymphatic network in the deeper layers of the injured cardiac tissue of *Ackr3^ΔLyve^*^1^ hearts, compared to *Ackr3^fl/fl^*hearts, 28-days post ligation (Figures 5G-I, Supplementary Figure 7A). In contrast, LYVE1-positive lymphatic structures were only present in the epicardial layer of *Ackr3^fl/fl^* hearts (Figure 5G, white arrowheads). Interestingly, we observed a tendency for accumulation of leukocytes, including F4/80+ macrophages, adjacent to these lymphatic vessels within the deeper layers of the injured tissue areas of *Ackr3^ΔLyve^*^1^ hearts (Figure 6G, Supplementary Figures 7B,C). Taken together, this data suggests that deletion of ACKR3 in LYVE1-positive cells may impact immune cell clearance from the injured cardiac tissues.

**Figure 6.**
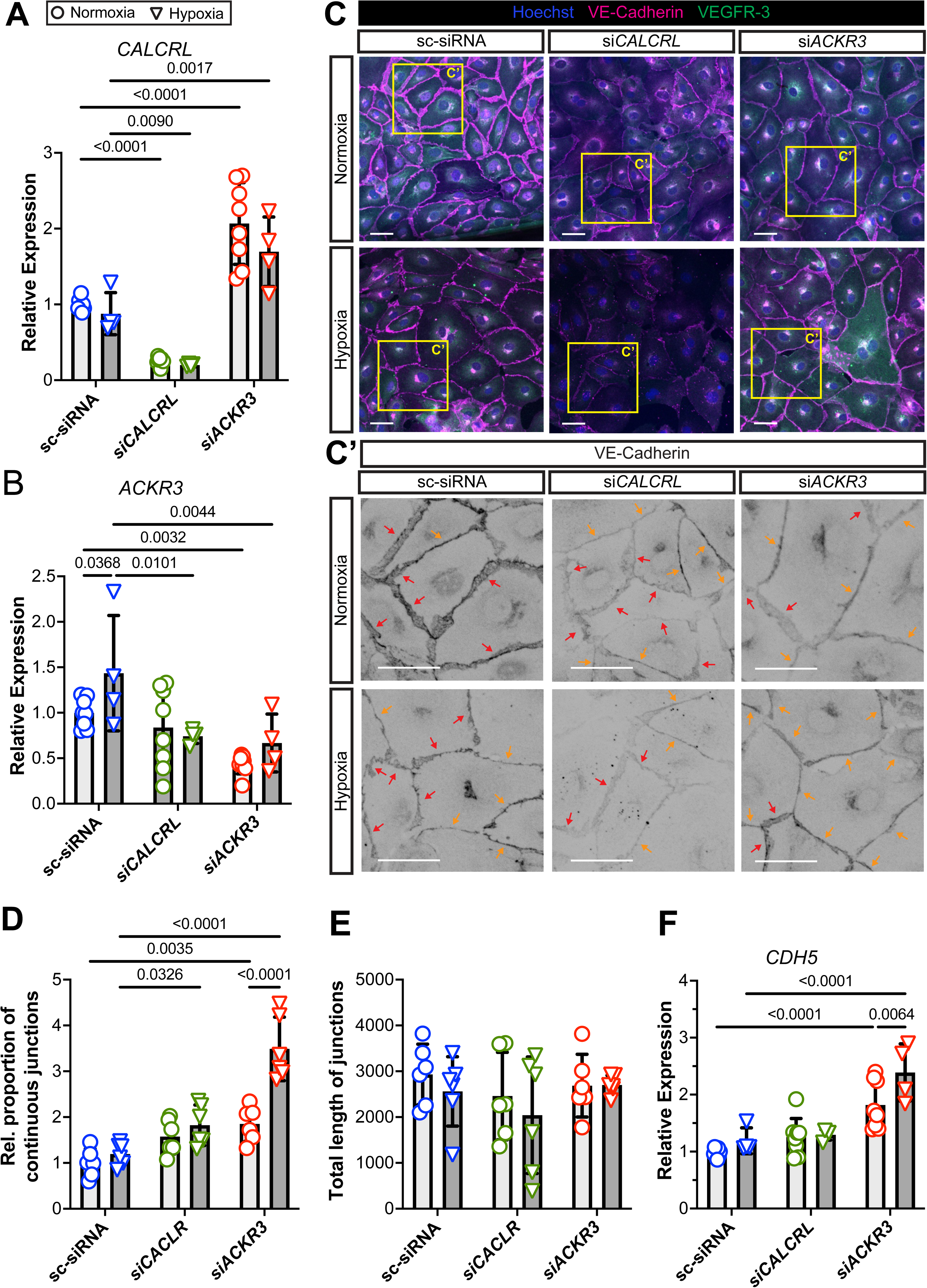
Hypoxia-induced tightening of lymphatic junctions is regulated by ACKR3. **A,B)** Relative gene expression levels of *CALCRL* **(A)** and *ACKR3* **(B)** in hLECs exposed to normoxia or hypoxia for 24h after treatment with non-targeting scramble (sc-)siRNA, *CALCRL*-targeting (*siCALCRL*) or *ACKR3*-targeting (*siACKR3*) siRNA. 1= average gene expression level in sc-siRNA treated cells under normoxia. N=4-8 per experimental group. Two-way ANOVA, Dunnett’s post-hoc test. **C)** VE-cadherin and VEGFR3 immunostaining of hLECs exposed to normoxia or hypoxia for 24h after treatment with control scramble (sc-), *CALCRL*- or *ACKR3*-targeting siRNA. Yellow and red arrows point to continuous and discontinuous VE-cadherin junction segments, respectively. Representative images from 6 biological replicates are shown. Bars, 20 µm. **D,E)** Relative proportion of continuous VE-cadherin junctions **(D)** and total length of VE-cadherin junctions regardless the type of junction dynamics **(E)** in hLECs exposed to normoxia or hypoxia for 24h after treatment with control scramble (sc-), *CALCRL*- or *ACKR3*-targeting siRNA. For relative proportion, 1 = average proportion of continuous VE-cadherin junctions in sc-siRNA treated normoxic hLECs. Two-way ANOVA, Dunnett’s post-hoc test. N=6 per group. **F)** Relative gene expression levels of *CDH5* in hLECs exposed to normoxia or hypoxia for 24h after treatment with non-targeting scramble (sc-)siRNA, *CALCRL*-targeting (*siCALCRL*) or *ACKR3*-targeting (*siACKR3*) siRNA. 1= average gene expression level in sc-siRNA treated cells under normoxia. N=4-8 per experimental group. Two-way ANOVA, Dunnett’s post-hoc test.

Since a subpopulation of macrophages also express LYVE1 and therefore lack ACKR3 expression in *Ackr3^ΔLyve^*^1^ mice, we sought to characterize the transcriptomic changes in macrophages isolated from the hearts of *Ackr3^fl/fl^*and *Ackr3^ΔLyve^*^1^ mice at baseline and 7 days post LAD ligation. Comparison of differentially expressed genes in CD68+ macrophages upon LAD ligation in *Ackr3^fl/fl^*and *Ackr3^ΔLyve^*^1^ mice revealed 1333 differentially expressed genes that were specific to LAD-ligated *Ackr3^ΔLyve^*^1^ mice (Supplementary Figures 8A,B, Supplementary Tables 4,5). Analysis of these differentially expressed genes revealed genes with well-established roles in modulating immune responses (E.g. *Il6*, *Ccl17, Ccr9*, *Cd226, Il1rl1, Serpinb2*) (Supplementary Figures 8C,D, Supplementary Tables 4,5). Gene ontology analysis of the differentially expressed genes further confirmed the activation of immune response pathways (Supplementary Figures 8E,F).

### 5.5 ACKR3 regulates hypoxia-induced rearrangement of lymphatic cellular junctions

In addition to promoting lymphangiogenesis under physiological and pathophysiological conditions^5,23,24^, AM signaling also orchestrates a reorganization of VE-cadherin cellular junctions in LECs^25,26^. Intriguingly, we found that hypoxia results in a time-dependent reorganization of VE-cadherin junctions in cultured hLECs (Supplementary Figures 9A,B), similarly to AM-treated LECs^25,26^. We also detected a modest, but statistically significant induction of *CDH5* gene expression levels in hLECs as soon as 6h after exposure to hypoxia (Supplementary Figure 9C). Since the availability of AM ligand is regulated by the scavenger activity of ACKR3^8^, we hypothesized that ACKR3 may also affect LEC intercellular junctions under hypoxic conditions, either through or independent of AM signaling. To test this hypothesis, we employed siRNA-induced gene silencing of *CALCRL* (*siCALCRL*), encoding the canonical AM receptor CLR, and *ACKR3* (*siACKR3*) in hLECs maintained under hypoxic or normoxic conditions. Interestingly, ACKR3 downregulation leads to a statistically significant induction of *CALCRL* gene expression levels under both normoxic and hypoxic conditions (Figure 6A), suggesting that ACKR3 not only titrates AM levels but also modulates AM-CLR signaling in LECs. Moreover, downregulation of *CALCRL* prevented the hypoxia-induced upregulation of *ACKR3* expression (Figure 6B), suggesting a bi-directional regulatory mechanism between AM and ACKR3 signaling in LECs. At the cellular physiological level, downregulation of both *CALCRL* and *ACKR3* increased the proportion of continuous VE-cadherin junctions, with ACKR3 exhibiting a more prominent role in regulating lymphatic cellular junction dynamics and *CDH5* gene expression levels under hypoxic conditions (Figures 6C-F, Supplementary Figures 9D-F). si*CALCRL* treatment also modestly affected the presence of lymphatic VE-cadherin junctions (Figure 6E, Supplementary Figure 9F) without affecting *CDH5* gene expression levels (Figure 6F). Together, these data demonstrate that AM peptide interacts with both signaling and scavenging receptors to govern the hypoxia-induced changes in VE-cadherin lymphatic cellular junctions.

## 6. Discussion

In this study, we show that expression and activation of ACKR3 in lymphatics is important for regulating the cardiac lymphatic response after ischemic heart injury. Spatial activation of ACKR3 signaling in the heart upon LAD ligation was characterized utilizing the novel ACKR3-TangoGFP mice in which GFP expression reports ligand activation of ACKR3. We detected a localized activation of ACKR3 signaling in the lymphatic vessels adjacent to the injury site after LAD ligation. Notably, in mice where the ratio of lymphatic / non lymphatic GFP signal ratio was lower in the peri-infarct zone, this ratio was high in the infarct zone, reporting that the ACKR3 signal activation in cardiac lymphatics is inversely proportional with the distance from the site of ischemic injury. Those mice with lower relative lymphatic GFP ratio in the infarcted tissue exhibited a higher infarct zone with severe tissue damage and lymphatic rarefaction. In these mice, high ACKR3 signaling intensity was observed in the lymphatics in the peri-infarct zone. These dynamics in lymphatic ACKR3 signaling suggest that local hypoxic conditions induce ACKR3 signaling in lymphatic vessels adjacent to the infarct zone outside of the necrotic tissue area. By crossing the ACKR3-TangoGFP strain to *Ramp3^-/-^* mice, we revealed that this ischemia-induced increase in lymphatic ACKR3 signaling activity is hindered by loss of RAMP3. Collectively, these data validate a unique in vivo reporter tool capable of defining sites of induced ACKR3 activation and indicates that RAMP3 is required for the ischemia-induced activation of ACKR3 in lymphatics upon LAD ligation. Other molecular mechanisms contributing to the upregulation of ACKR3 in LECs upon hypoxia and leading to the activation of ACKR3 signaling in lymphatics are yet to be discovered. Surprisingly, despite that RAMP3 may also heteromerize with CLR, forming a potent receptor complex for AM^27^, we found that downregulation of RAMP3 in hLECs leads to prolonged maintenance of AM-induced ERK phosphorylation, likely due to enhancing AM ligand availability by limiting ACKR3 recycling^10^. This data indicates the complex role of RAMP3 in modulating AM signaling in LECs.

Prior studies revealed that adrenomedullin plays an important role in cardiac development. *Adm^-/-^* mice exhibit developmental heart defects at E13.5, before they die in utero due to cardiovascular complications^28^. In contrast, global *Ackr3^-/-^* mouse embryos with uninhibited adrenomedullin signaling and adult *Adm^hi/hi^* mice overexpressing adrenomedullin have bigger hearts than control littermates^5,8^. Similarly to *Adm^hi/hi^* and global *Ackr3^-/-^* mice, here we found that both female and male adult mice with lymphatic loss of ACKR3 display cardiomegaly in *Ackr3^ΔLyve^*^1^ mice. This underlines the role of lymphatic ACKR3 signaling in regulating the physiological cardiovascular effects of adrenomedullin. However, unlike *Ackr3^-/-^*embryos, adult female and male *Ackr3^ΔLyve^*^1^ mice display no increased cardiac lymphatic density. This is not surprising, since epicardial adrenomedullin expression peaks during embryonic development, while the development of cardiac lymphatic vasculature ends only 2 weeks post birth in mice. Thus, despite the defective regulation of adrenomedullin signaling in *Ackr3^ΔLyve^*^1^ mice, this probably has less effect on the development of mature cardiac lymphatic network. Notably, only female adult *Adm^hi/hi^* mice display increased lymphatic counts in the heart, male *Adm^hi/hi^* have comparable cardiac lymphatic density to wild type littermates^5^, despite the constitutive expression of adrenomedullin in these mice, supporting that enhanced adrenomedullin signaling does not necessary lead to uninhibited lymphatic development. Prior human data, however, showed that plasma adrenomedullin levels are elevated in patients with myocardial infarction^7^, and we found a hypoxia-induced upregulation of both *ADM* and *ACKR3* expression levels in cultured LECs. This re-activation of AM and ACKR3 signaling upon myocardial infarction potentiated a possible cardiac lymphatic phenotype in *Ackr3^ΔLyve^*^1^ mice challenged with ischemic cardiac injury by LAD ligation, in addition to the likely positive cardiac effects of uninhibited AM signaling.

*Ackr3^ΔLyve^*^1^ mice exhibit significantly better survival after LAD ligation, especially within the first couple days post injury. Furthermore, although control *Ackr3^fl/fl^*mice developed acute cardiac edema within 3 days post LAD ligation, similar to reports from prior studies^3,5,29,30^, *Ackr3^ΔLyve^*mice were protected from the development of cardiac edema, to an extent that is similar to that previously reported for *Adm^hi/hi^* mice which express 3-times higher AM levels^5^. Lymphatics of mice with no ACKR3 signaling are likely more sensitive to increased AM levels after myocardial infarction and this uninhibited AM signaling may impart cardioprotection in *Ackr3^ΔLyve^*^1^ mice after ischemic cardiac injury. Furthermore, assessment of the cardiac lymphatic network after LAD ligation revealed an enhanced cardiac lymphatic expansion in *Ackr3^ΔLyve^*^1^ mice, most notably deep in the infarcted area, affected by the highest degree of ischemia post LAD ligation. This observation confirms a crucial role for hypoxia-induced lymphatic ACKR3 in regulating the expansion of the cardiac lymphatic network post myocardial infarction. Accordingly, we also revealed an altered expression of genes associated with tissue remodeling and endothelial cell function, as well as immunomodulatory and inflammatory pathways in ACKR3-deficient cardiac LECs, compared to *Ackr3^fl/fl^* controls. A calculated limitation, and yet fortuitous advantage, of this study design is that a subpopulation of macrophages also express LYVE1. Thus, we were able to determine that deletion of ACKR3 in LYVE1+ macrophages affected the expression of genes and pathways involved in the modulation of immune responses in cardiac macrophages. Thus, the phenotypes observed in *Ackr3^ΔLyve^*^1^ mice post LAD ligation, especially leukocyte populations, are likely also affected by the subpopulation of macrophages that lack ACKR3 signaling. Notably, all of the currently available lymphatic-Cre lines are riddled with some degree of non-lymphatic off-target effects ^31^, which is a well-known limitation of lineage-specific in vivo approaches in this field of research.

In addition of promoting lymphangiogenesis, AM also regulates intercellular junctions in lymphatics, largely through reorganization of VE-cadherin junctions. We revealed that modulation of *ACKR3* gene expression levels play a prevalent role in inducing the formation of continuous VE-cadherin junctions and fine-tuning *CDH5* gene expression levels under hypoxic conditions. We have recently revealed the role of lymphatic VE-cadherin in the maintenance of cardiac lymphatic vessels and lymphangiogenesis after myocardial infarction, but loss of endothelial VE-cadherin does not affect systolic cardiac function^29^. Thus, modulation of lymphatic VE-cadherin may be another possible downstream underlying mechanism responsible, at least in part, for the lymphatic phenotype observed after ischemic cardiac injury in mice with lymphatic loss of ACKR3.

Collectively, our results demonstrate that lymphatic ACKR3 is spatially activated within the heart following ischemic injury and plays an important role in multiple mechanisms, including the regulation of a lymphangiogenic response and modulating lymphatic intercellular junctions. Since AM ligand levels are regulated by ACKR3 scavenging activity, and mice with lymphatic loss of ACKR3 result in similar phenotypes observed in AM overexpression and ACKR3 global knockout mice, these data bolster a cooperative role for AM and ACKR3 in lymphatics following ischemic heart injury.

## 7. Acknowledgements

The authors appreciate the contributions of current and former members of the Caron Lab. We also thank Dr. Pablo Ariel (UNC-CH Microscopy Services Laboratory), Dr. Mike Vernon (UNC-CH Functional Genomics Core), Dr. Jonathan Schisler (UNC-CH Department of Pharmacology), Dr. Michelle Itano and Dr. Tessa-Jonne Ropp (UNC-CH Neuroscience Microscopy Core), Wendy Salmon (UNC-CH Hooker Imaging Core), Dr. Haifeng Yin (UNC-CH MHI Cardiovascular Physiology and Phenotyping Core), Dr. Li Qian and Dr. Joan M. Taylor (UNC-CH Department of Pathology and Laboratory Medicine) and Dr. Kristy Red-Horse and members of the Red-Horse Lab (Stanford University Department of Biology).

The UNC Microscopy Services Laboratory and the UNC Hooker Imaging Core Facility are supported in part by P30 CA016086 Cancer Center Core Support Grant to the UNC Lineberger Comprehensive Cancer Center. Research involving light-sheet imaging reported in this publication was supported in part by the North Carolina Biotech Center Institutional Support Grant 2016-IDG-1016. UNC Neuroscience Microscopy Core (RRID:SCR_019060) is supported, in part, by funding from the NIH-NICHD Intellectual and Developmental Disabilities Research Center Support Grant P50 HD103573.

## 8. Sources of Funding

National Institutes of Health (NIH) grants from the National Heart, Lung, and Blood Institute (NHLBI) HL1290986 and National Institute of Diabetes and Digestive and Kidney Diseases DK119145 and an American Heart Association (AHA) Innovator Award to Kathleen M. Caron, an AHA Postdoctoral Fellowship 23POST1022945 to Laszlo Balint, a NIH National Cancer Institute T32CA071341 and a NHLBI F31 HL163885 Predoctoral Fellowship to D.S. Serafin.

## 9. Disclosures

None.

## Notes

### Competing Interest Statement

The authors have declared no competing interest.

