## Supplementary Figures 1-9 for "Lymphatic activation of ACKR3 signaling regulates lymphatic response after ischemic heart injury"

### Supplementary Figure 1.

Balint et al.

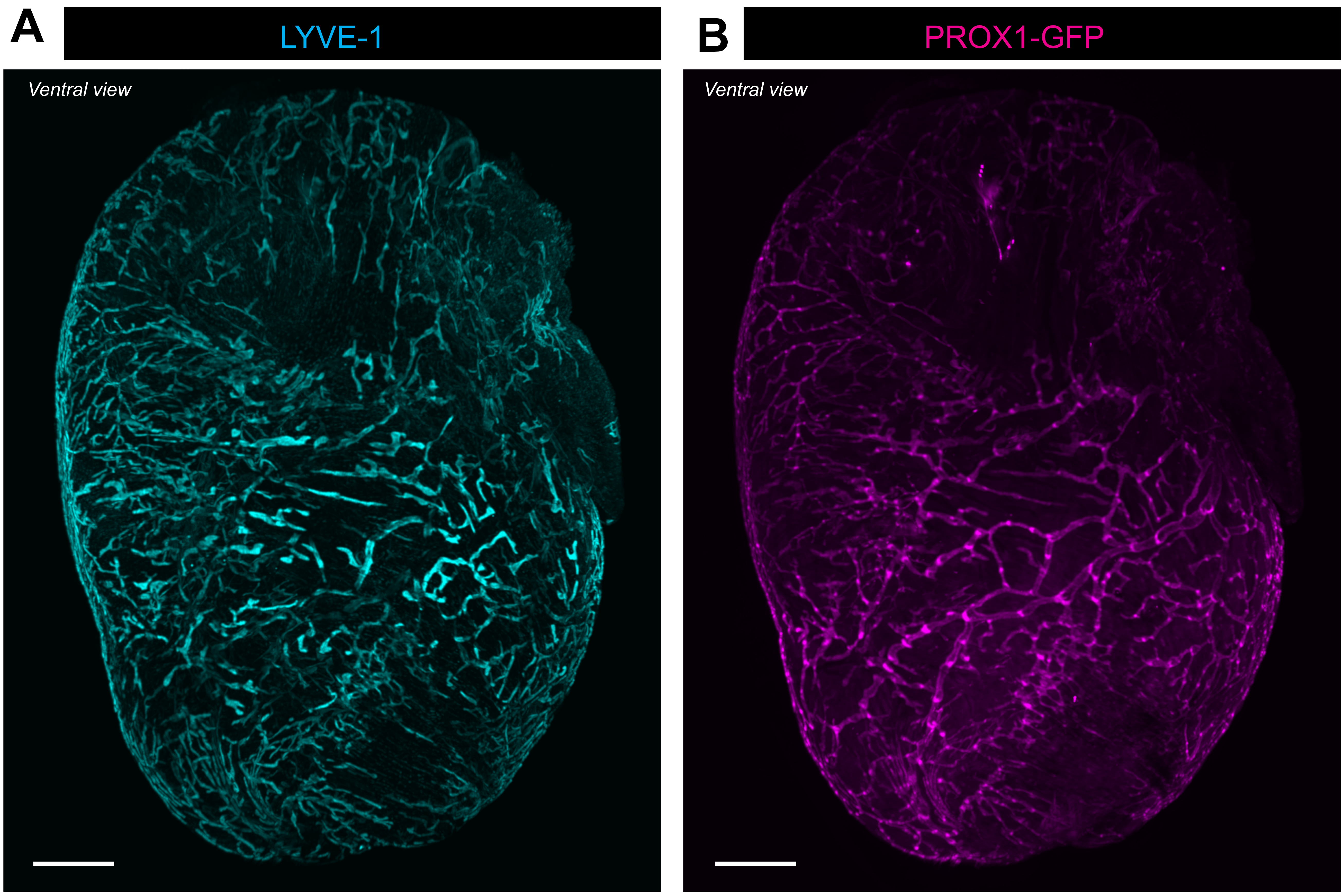

Supplementary Figure 2.

Balint et al.

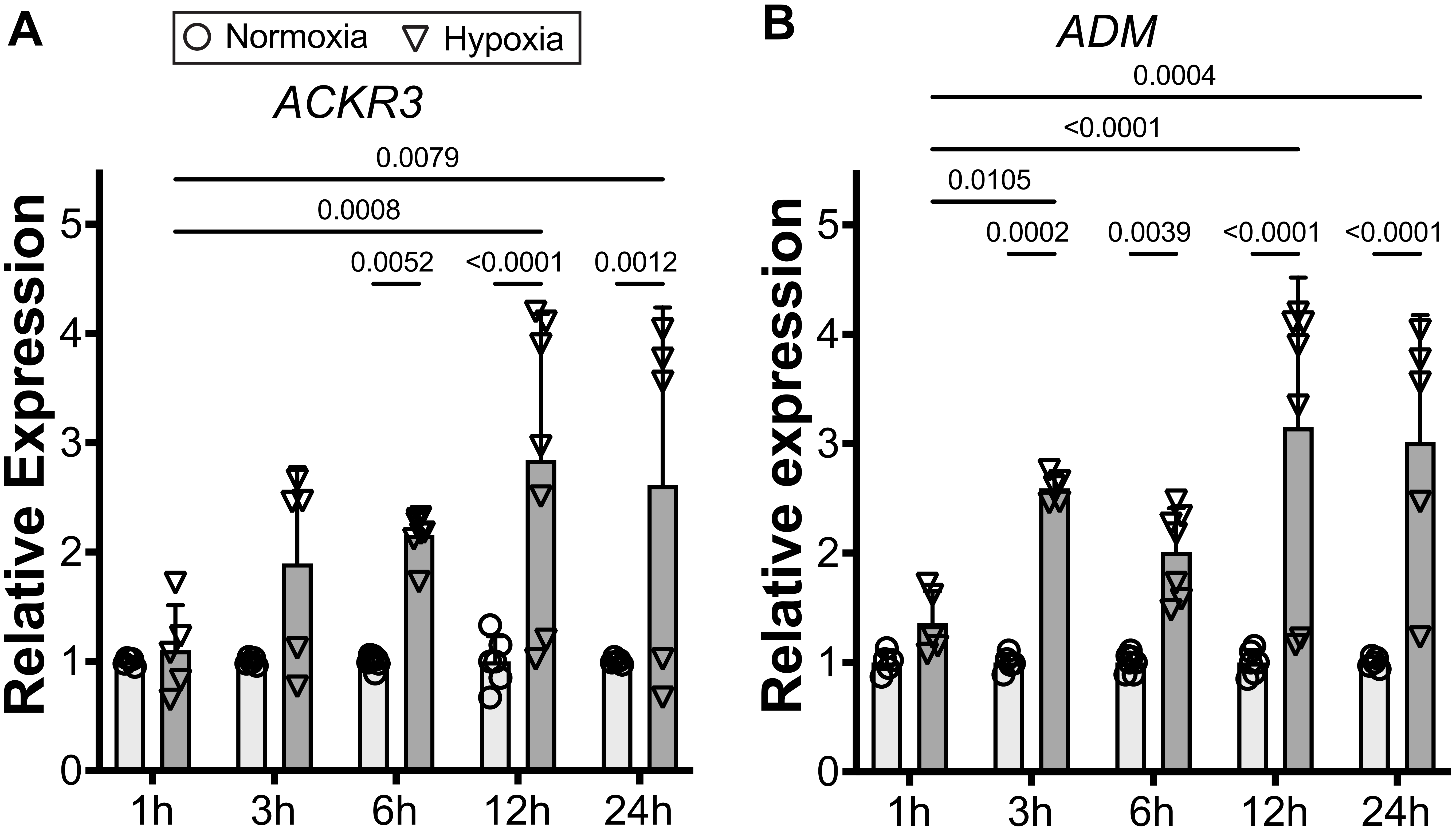

### Supplementary Figure 3.

Balint et al.

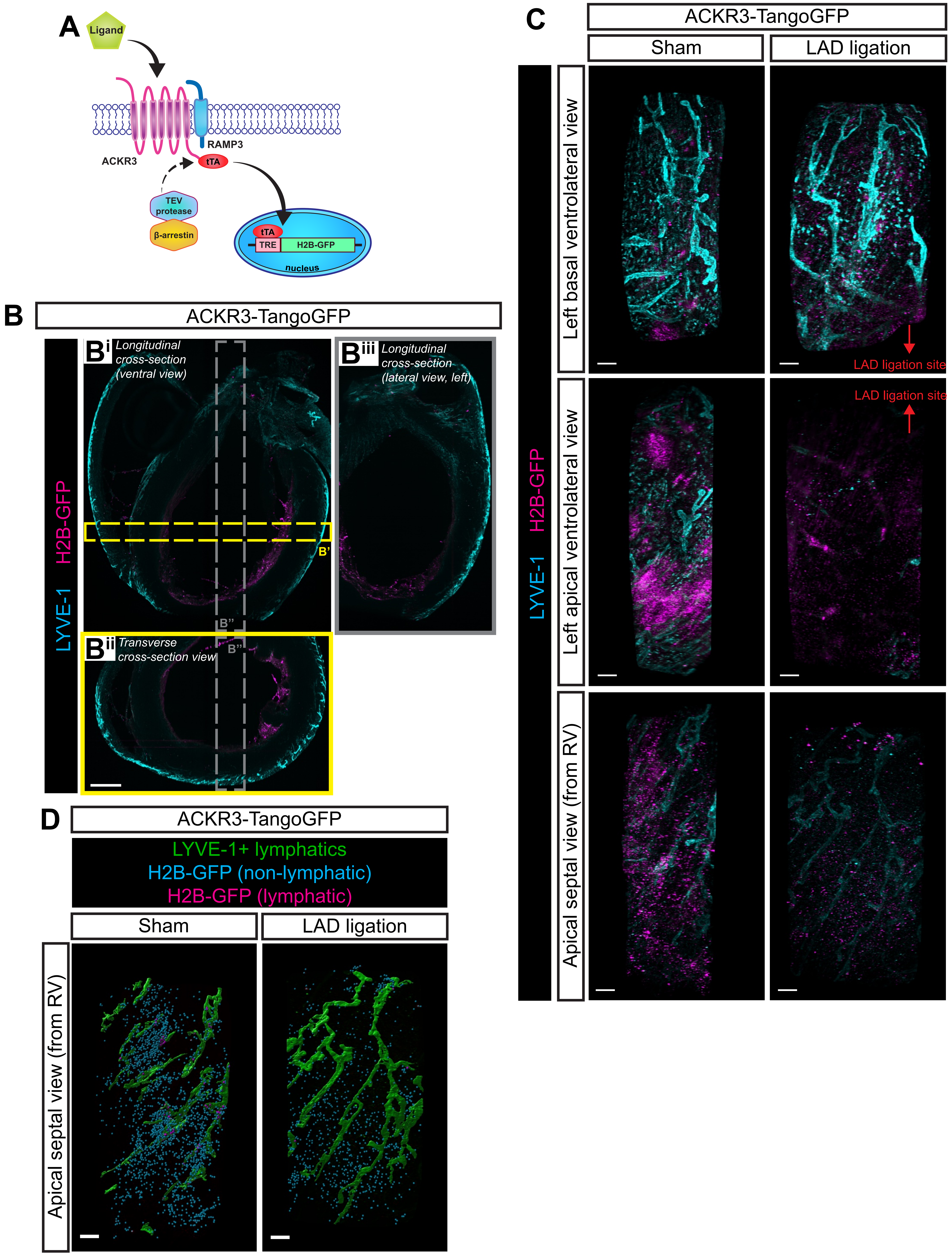

### Supplementary Figure 4.

Balint et al.

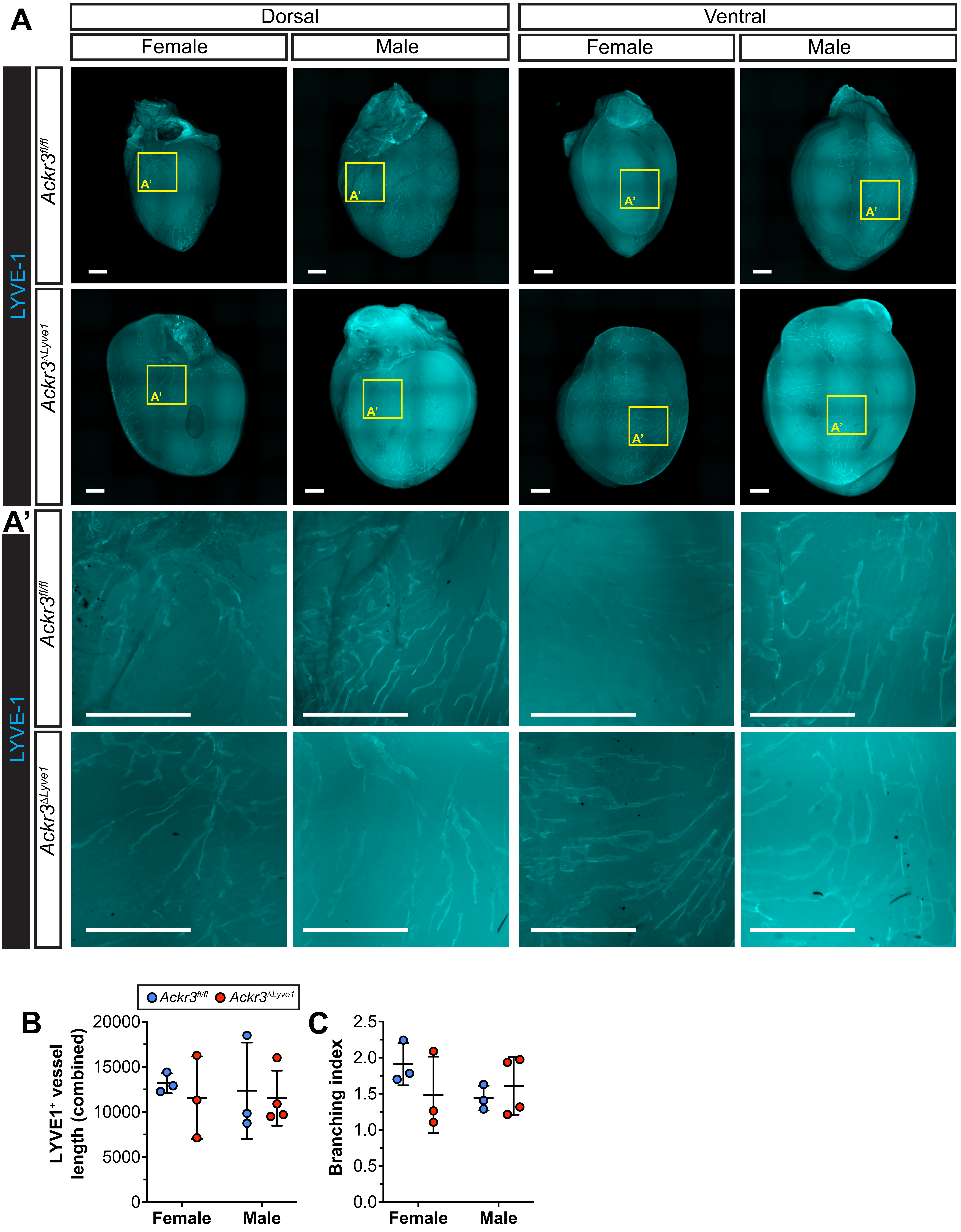

### Supplementary Figure 5.

Balint et al.

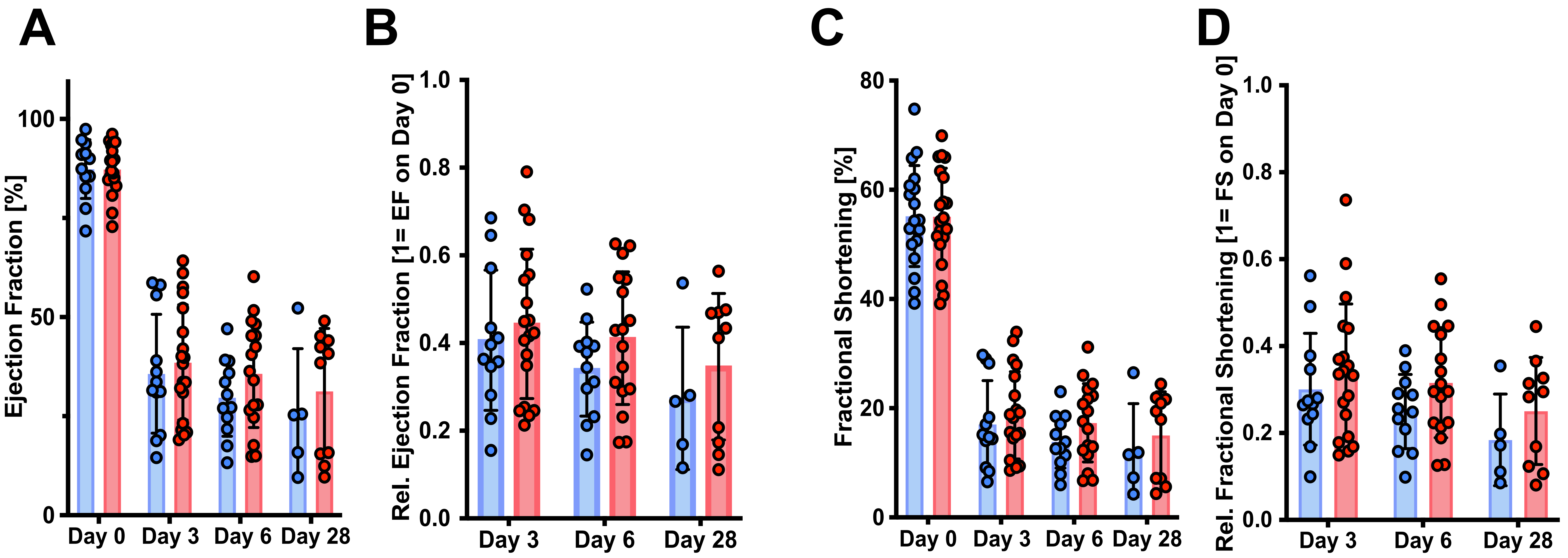

### Supplementary Figure 6.

Balint et al.

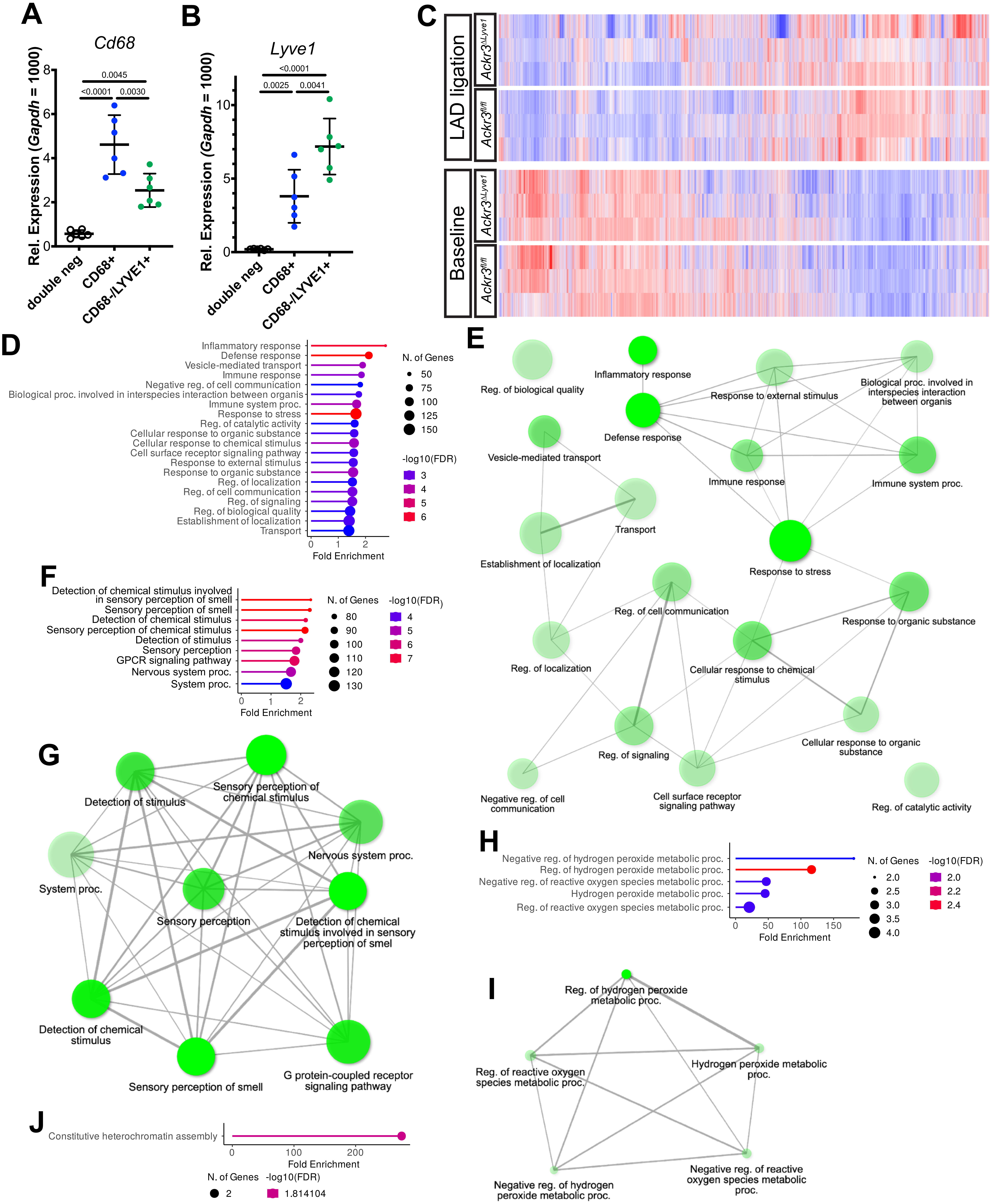

### Supplementary Figure 7.

Balint et al.

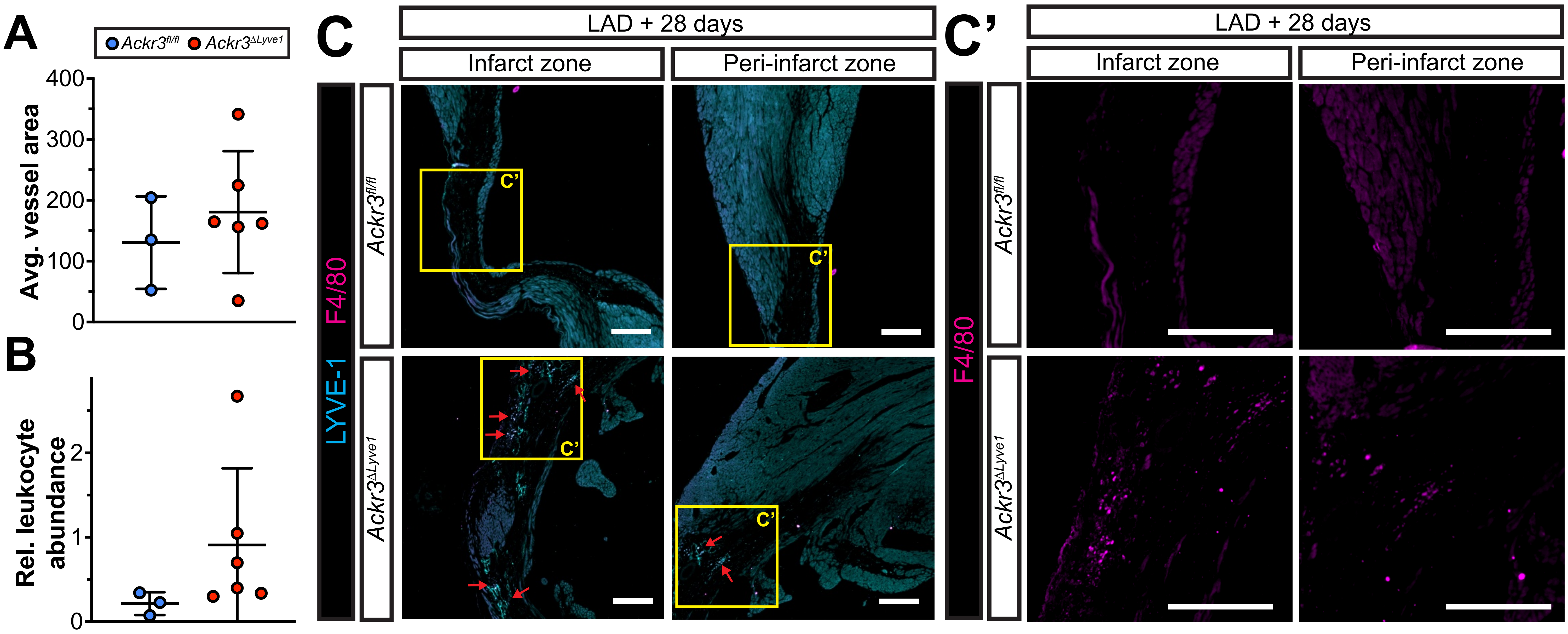

### Supplementary Figure 8.

Balint et al.

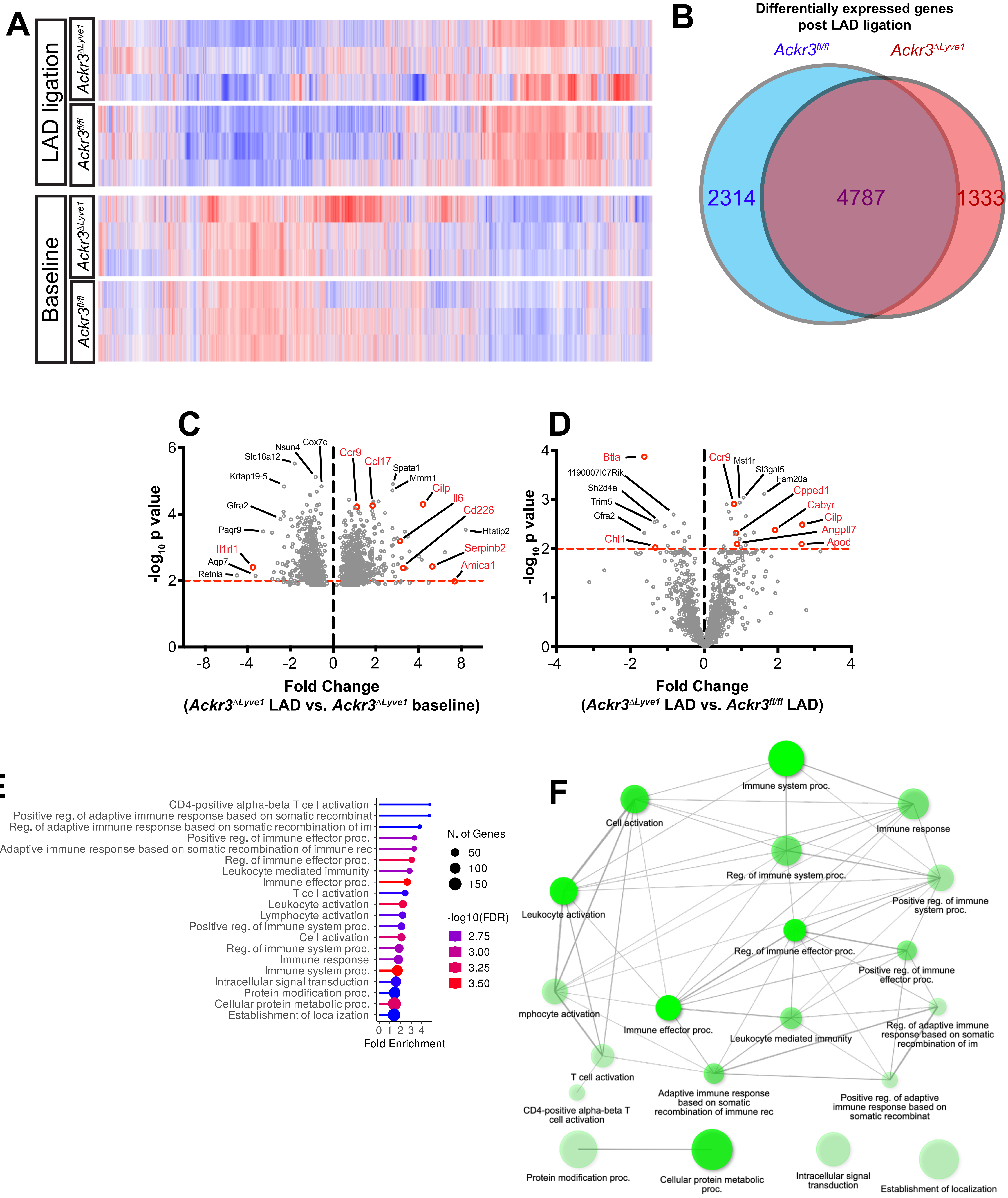

### Supplementary Figure 9.

Balint et al.

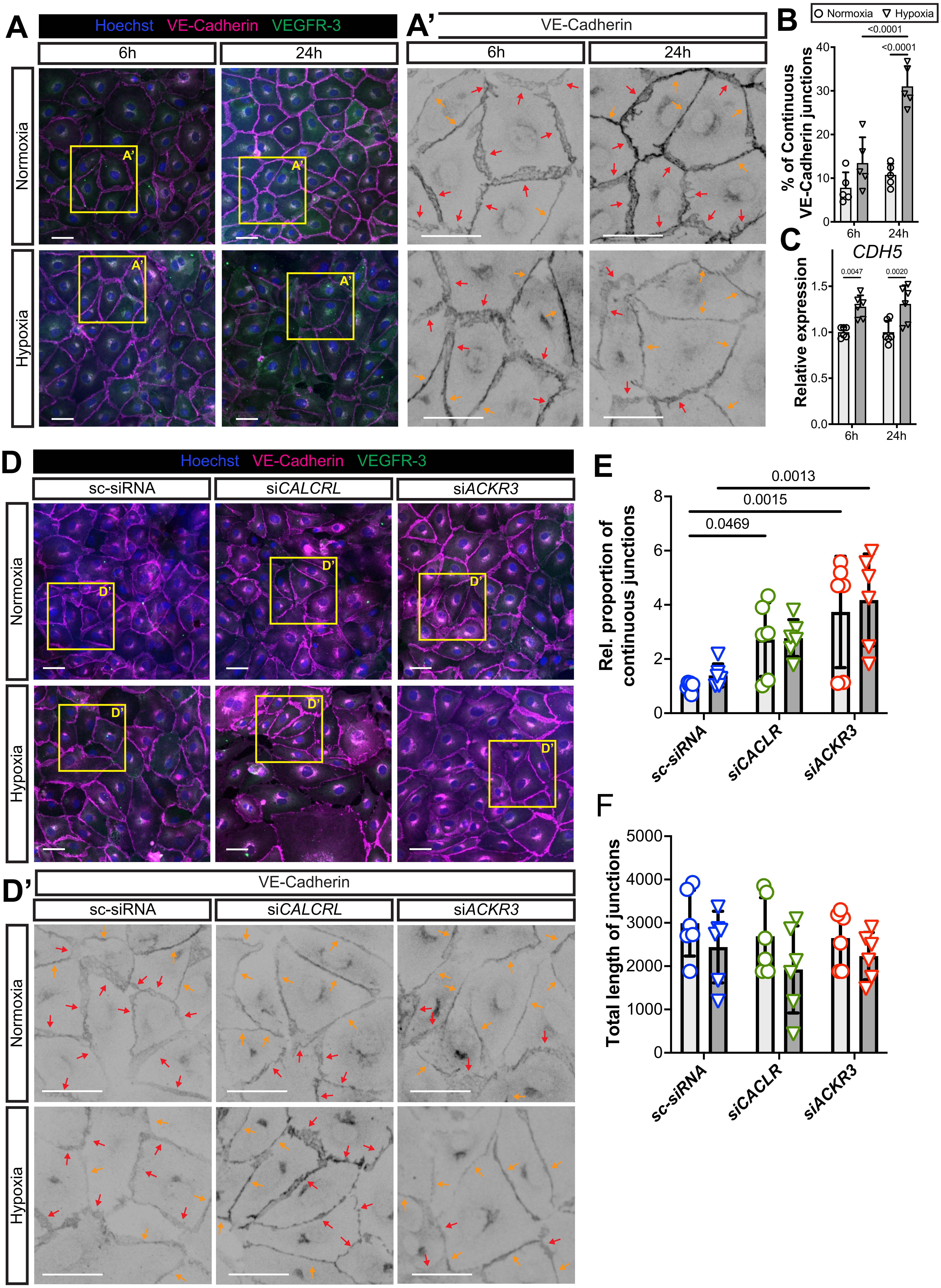

**Supplementary Figure 1.** Separated channels shown for lymphatic vessel endothelial hyaluronan receptor 1 (LYVE1) **(A)** and Prospero homeobox protein (PROX1) **(B)** positive cardiac lymphatic structures captured by light-sheet imaging of male PROX1-GFP mouse hearts immunostained with anti-LYVE1 and anti-GFP antibodies using the iDisco tissue clearing and staining protocol. Bars, 1000 µm.

**Supplementary Figure 2. A,B)** Fold changes in atypical chemokine receptor 3 (*ACKR3)* **(A)** and adrenomedullin (*ADM)* **(B)** mRNA levels in human primary lymphatic endothelial cells (hLEC) exposed to hypoxia or normoxia for 1h, 3h, 6h, 12h, 24h (1 = gene expression levels in normoxic cells at the corresponding time point). Housekeeping gene: actin beta (*ACTB)*. Biological replicates are shown. Two-way ANOVA, Dunnett's multiple comparisons test. N=5-7 per group.

**Supplementary Figure 3. A)** Explanation of the ACKR3-TangoGFP system. Adapted from *Kono et al., J Clin Invest 2014* [22]. **B)** Low-magnification expanded cross-section views showing ACKR3 activation in the endocardial layer of uninjured ACKR3-TangoGFP hearts, detected by H2B-GFP expression. Scale bar, 1000 μm. **C)** 3-D high-magnification fluorescent images showing LYVE1-positive lymphatics and ACKR3 activation detected by H2B-GFP signal in different volume segments of ACKR3-TangoGFP hearts 2 days post sham surgery or LAD ligation. Bars, 200 μm. D) 3-D rendered pseudocolored images showing LYVE1-positive cardiac lymphatic vessels (green), ACKR3 activation in LYVE1-negative (blue) and LYVE1-positive (magenta) cells in the apical septal wall facing the right ventricle of ACKR3-Tango-GFP mice 2 days post sham surgery or LAD ligation. Bars, 200 μm. Representative images from 3 or more animals per group are shown. Abbreviations: Atypical chemokine receptor 3, ACKR3; Receptor activity modifying protein 3, RAMP3; Tobacco Etch Virus, TEV; tetracycline-regulated transactivator, tTA; Tet response element, TRE; Histone H2B, H2B; Green fluorescent protein, GFP; Left anterior descending artery, LAD; Lymphatic vessel endothelial hyaluronan receptor 1, LYVE1

**Supplementary Figure 4. A)** Visualization of lymphatic vessel endothelial hyaluronan receptor 1 (LYVE1)-positive cardiac lymphatic vessels of *Ackr3*$^{fl/fl}$ and *Ackr3*$^{\Delta Lyve1}$ female and male whole mount hearts. Representative images from 3-4 animals per group are shown. Bars, 1000 µm. **B,C)** Quantification of lymphatic vessel length **(B)** and branching index (number of branches divided by lymphatic vessel length) **(C)** per field of view. N=3-4 per group. Two-way ANOVA, Dunnett's post-hoc test.

**Supplementary Figure 5. A-D)** Echocardiographic assessment of cardiac function of male *Ackr3*$^{fl/fl}$ and *Ackr3*$^{\Delta Lyve1}$ mice before left anterior descending artery (LAD) ligation and 3-, 6- and 28-days post LAD ligation. Changes in ejection fraction **(A)**, relative ejection fraction (ejection fraction on the corresponding day divided by ejection fraction on Day 0 in the same animal) **(B)**, fractional shortening **(C)**, and relative fractional shortening (fractional shortening on the corresponding day divided by fractional shortening on Day 0 in the same animal) **(D)** parameters are shown. N=5-20 mice per group. Mixed effects model of variance, Šídák's multiple comparison post-hoc test.

**Supplementary Figure 6. A,B)** Relative gene expression levels of cluster of differentiation 68 (*Cd68)* **(A)** and lymphatic vessel endothelial hyaluronan receptor 1 (*Lyve1)* **(B)** genes in cells isolated by anti-CD68 and anti-LYVE1 immunoprecipitation of *Ackr3*$^{fl/fl}$ and *Ackr3*$^{\Delta Lyve1}$ hearts. N= 6 per group. One-way ANOVA; Tukey's post hoc test. **C)** Heat maps showing gene expression patterns in CD68-negative LYVE1-positive lymphatic endothelial cells (LECs) isolated from the hearts of *Ackr3*$^{fl/fl}$ and *Ackr3*$^{\Delta Lyve1}$ male mice with or without left anterior descending artery (LAD) ligation. Upregulated and downregulated genes upon LAD ligation are shown in red and blue, respectively. Individual data from N=3 mice per experimental groups are shown. **D,E)** Gene Ontology annotation **(D)** and network analysis **(E)** of biological pathways upregulated in LECs isolated from *Ackr3*$^{\Delta Lyve1}$ hearts one week post LAD ligation compared to uninjured *Ackr3*$^{\Delta Lyve1}$ cardiac LECs. **F,G)** Gene Ontology annotation **(F)** and network analysis **(G)** of biological pathways downregulated in LECs isolated from *Ackr3*$^{\Delta Lyve1}$ hearts post LAD ligation compared to uninjured *Ackr3*$^{\Delta Lyve1}$ cardiac LECs. **H,I)** Gene Ontology annotation **(H)** and network analysis **(I)** of biological pathways upregulated in LECs isolated from *Ackr3*$^{\Delta Lyve1}$ hearts compared to *Ackr3*$^{fl/fl}$ LECs upon LAD ligation. **J)** Gene ontology annotation of downregulated biological pathways in LECs isolated from *Ackr3*$^{\Delta Lyve1}$ hearts compared to *Ackr3*$^{fl/fl}$ LECs upon LAD ligation. Images were generated with Shiny GO 0.8 using the "GO Biological Process" pathway database. For Gene Ontology annotation images, bar lengths correlate to -log10 false discovery rate (FDR), while the size of bar-terminating circles correlate to the number of genes found within the given pathway. For network analysis images, node sizes correlate to size of gene sets, intensity and opacity of green shading refers to enrichment of gene sets (transparent indicates lower degree of enrichment, opaque indicates higher degree of enrichment). Two pathways are considered connected if gene overlap between sets is >20% and is denoted by lines between nodes. Thickness of lines correlates to the degree of gene overlap between nodes.

**Supplementary Figure 7. A) A)** Average area of lymphatic vessels in the infarct zone of male *Ackr3*$^{fl/fl}$ and *Ackr3*$^{\Delta Lyve1}$ hearts 28 days post left anterior descending artery (LAD) ligation. N= 3-6 per group. Unpaired Welch's unequal variances t-test. **B)** Area-adjusted relative abundance of CD45-positive leukocytes per field of view (Number of CD45+ cells divided by infarcted area in the field of view) in the infarct zone of *Ackr3*$^{fl/fl}$ and *Ackr3*$^{\Delta Lyve1}$ male mice. N=3-6 per group. Mann-Whitney test. **C)** Lymphatic vessel endothelial hyaluronan receptor 1 (LYVE1, cyan)- and adhesion G protein-coupled receptor E1 (F4/80, magenta)-specific staining of the infarct zone and peri-infarct zones of male *Ackr3*$^{fl/fl}$ and *Ackr3*$^{\Delta Lyve1}$ whole mount hearts 28 days post LAD ligation. Red arrows point at F4/80-positive macrophages. Sections were collected 200μm from the beginning of the infarct zone towards the apex of the heart. Merged and separated F4/80 monochrome representative images from N=3-6 mice per group are shown. Bars, 200 μm.

**Supplementary Figure 8. A)** Heat maps showing gene expression patterns in cluster of differentiation 68 (CD68)-positive macrophages isolated from the hearts of *Ackr3*$^{fl/fl}$ and *Ackr3*$^{\Delta Lyve1}$ male mice with or without left anterior descending artery (LAD) ligation. Upregulated and downregulated genes upon LAD ligation are shown in red and blue, respectively. Individual data from N=3 mice per experimental groups are shown **B)** Venn diagram summarizing the transcriptomic differences in cardiac macrophages isolated from male *Ackr3*$^{fl/fl}$ and *Ackr3*$^{\Delta Lyve1}$ mice one week post LAD ligation. **C)** Differentially expressed genes in cardiac macrophages upon LAD ligation that are specific to ACKR3-deficient mice. Positive fold change values represent upregulation in LAD ligated *Ackr3*$^{\Delta Lyve1}$ macrophages. **D)** Comparison of gene expression levels of identified ACKR3-specific LAD-triggered genes between LAD ligated *Ackr3*$^{\Delta Lyve1}$ and *Ackr3*$^{fl/fl}$ mice. Positive fold change values represent upregulation in LAD ligated *Ackr3*$^{\Delta Lyve1}$ macrophages. **E,F)** Gene Ontology annotation **(E)** and network analysis **(F)** of biological pathways upregulated in CD68+ macrophages isolated from *Ackr3*$^{\Delta Lyve1}$ hearts one week post LAD ligation compared to uninjured *Ackr3*$^{\Delta Lyve1}$ cardiac macrophages. Images were generated with Shiny GO 0.8 using the "GO Biological Process" pathway database. For Gene Ontology annotation images, bar lengths correlate to -log10 false discovery rate (FDR), while the size of bar-terminating circles correlate to the number of genes found within the given pathway. For network analysis images, node sizes correlate to size of gene sets, intensity and opacity of green shading refers to enrichment of gene sets (transparent indicates lower degree of enrichment, opaque indicates higher degree of enrichment). Two pathways are considered connected if gene overlap between sets is >20% and is denoted by lines between nodes. Thickness of lines correlates to the degree of gene overlap between nodes.

**Supplementary Figure 9. A)** vascular endothelial (VE)-cadherin (magenta) and vascular endothelial growth factor receptor 3 (VEGFR3, green)-specific immunostaining of cultured human primary lymphatic endothelial cells (hLECs) exposed to normoxia or hypoxia for 6h or 24h. Yellow and red arrows point to continuous and discontinuous VE-cadherin junction segments, respectively. Representative merged and separated VE-cadherin monochrome inset images are shown from 5 biological replicates per group. Bars, 20 µm. **B)** Quantitative analysis of the percent ratio of continuous VE-cadherin junctions in hLECs exposed to normoxia or hypoxia for 6h, or 24h. Two-way ANOVA, Šídák's post-hoc test. N=5 per group. **C)** Fold changes in cadherin 5 (*CDH5)* mRNA levels in hLECs exposed to hypoxia for 6h, or 24h, compared to gene expression levels in time-matched hLECs cultured under normoxic conditions. Two-way ANOVA, Šídák's post-hoc test. N=6 per group. **D)** VE-cadherin (magenta) and VEGFR3 (green) immunostaining of hLECs exposed to normoxia or hypoxia for 6h after treatment with control scramble (sc-), calcitonin receptor like receptor (*CALCRL)*- or atypical chemokine receptor 3 (*ACKR3)*-targeting siRNA. Yellow and arrows point to continuous and discontinuous VE-cadherin junction segments, respectively. Representative merged and separated VE-cadherin monochrome inset images from 6 biological replicates are shown. Bars, 20 µm. **E,F)** Quantitative analysis of relative proportion of continuous VE-cadherin junctions **(E)** and total length of VE-cadherin junctions regardless the type of junction dynamics **(F)** in hLECs exposed to normoxia or hypoxia for 6h after treatment with control scramble (sc-), *CALCRL*- or *ACKR3*-targeting siRNA. For relative proportion, 1 = average proportion of continuous VE-cadherin junctions in sc-siRNA treated normoxic hLECs. Two-way ANOVA, Dunnett’s post-hoc test. N=6 per group.

**Supplementary Video 1.** 3-dimensional volumetric visualization of the cardiac lymphatic network of *Prox1*$^{GFP}$ mouse hearts immunostained with anti-LYVE1 (cyan) and anti-GFP (magenta) antibodies using the iDisco tissue clearing and staining protocol and light-sheet imaging. Lymphatic structures in the cardiac septal wall are pseudocolored with yellow color. Lymphatic vessels penetrating into the deep myocardial layers are shown in the high magnification views. Abbreviations: Prospero homeobox 1, PROX1; green fluorescent protein, GFP; lymphatic vessel endothelial hyaluronan receptor 1, LYVE1. Dynamic scale bar is shown.
