## Supplementary Tables 1-5 for "Lymphatic activation of ACKR3 signaling regulates lymphatic response after ischemic heart injury"

**Supplementary Table 1.** Echocardiogram analysis of *Ackr3*$^{fl/fl}$ and *Ackr3*$^{\Delta Lyve1}$ mice before and after LAD ligation.

| | | Baseline | | | 3 days post LAD ligation | | | 6 days post LAD ligation | | | 14 days post LAD ligation | | | 21 days post LAD ligation | | | 28 days post LAD ligation | | |
|---|---|---|---|---|---|---|---|---|---|---|---|---|---|---|---|---|---|---|---|
| | | *Ackr3*$^{fl/fl}$ | *Ackr3*$^{\Delta Lyve1}$ | t test p-value | *Ackr3*$^{fl/fl}$ | *Ackr3*$^{\Delta Lyve1}$ | t test p-value | *Ackr3*$^{fl/fl}$ | *Ackr3*$^{\Delta Lyve1}$ | t test p-value | *Ackr3*$^{fl/fl}$ | *Ackr3*$^{\Delta Lyve1}$ | t test p-value | *Ackr3*$^{fl/fl}$ | *Ackr3*$^{\Delta Lyve1}$ | t test p-value | *Ackr3*$^{fl/fl}$ | *Ackr3*$^{\Delta Lyve1}$ | t test p-value |
| Primary parameters | IVS;d | 1.08 ± 0.29 | 1.07 ± 0.26 | 0.9434 | 0.69 ± 0.36 | 0.70 ± 0.27 | 0.9112 | 0.55 ± 0.12 | 0.63 ± 0.26 | 0.2734 | 0.35 ± 0.09 | 0.56 ± 0.37 | 0.1153 | 0.33 ± 0.05 | 0.57 ± 0.43 | 0.1065 | 0.30 ± 0.04 | 0.54 ± 0.41 | 0.0979 |
| | IVS;s | 1.64 ± 0.28 | 1.62 ± 0.27 | 0.7796 | 0.77 ± 0.42 | 0.83 ± 0.36 | 0.6884 | 0.61 ± 0.13 | 0.75 ± 0.40 | 0.2108 | 0.36 ± 0.08 | 0.76 ± 0.64 | 0.0837 | 0.38 ± 0.17 | 0.73 ± 0.67 | 0.1538 | 0.29 ± 0.06 | 0.67 ± 0.66 | 0.1012 |
| | LVID;d | 2.39 ± 0.37 | 2.32 ± 0.47 | 0.6406 | 3.62 ± 0.79 | 3.77 ± 0.72 | 0.6145 | 4.38 ± 0.91 | 4.52 ± 0.82 | 0.6902 | 4.88 ± 0.76 | 4.41 ± 1.12 | 0.3634 | 4.91 ± 0.85 | 4.55 ± 1.24 | 0.5251 | 5.05 ± 0.99 | 4.82 ± 1.25 | 0.7112 |
| | LVID;s | 1.07 ± 0.27 | 1.05 ± 0.31 | 0.8423 | 3.05 ± 0.90 | 3.10 ± 0.82 | 0.8747 | 3.80 ± 0.95 | 3.78 ± 0.97 | 0.9556 | 4.28 ± 0.99 | 3.50 ± 1.44 | 0.2471 | 4.23 ± 1.14 | 3.88 ± 1.50 | 0.6316 | 4.47 ± 1.19 | 4.17 ± 1.44 | 0.6765 |
| | LVPW;d | 1.18 ± 0.28 | 1.15 ± 0.19 | 0.6471 | 1.13 ± 0.46 | 0.91 ± 0.32 | 0.1770 | 1.00 ± 0.55 | 0.75 ± 0.28 | 0.1761 | 0.81 ± 0.39 | 0.86 ± 0.29 | 0.8059 | 0.96 ± 0.37 | 0.71 ± 0.27 | 0.2199 | 0.94 ± 0.57 | 0.73 ± 0.29 | 0.4580 |
| | LVPW;s | 1.92 ± 0.26 | 1.90 ± 0.21 | 0.8893 | 1.45 ± 0.61 | 1.23 ± 0.43 | 0.2832 | 1.29 ± 0.62 | 1.07 ± 0.40 | 0.2792 | 1.23 ± 0.58 | 1.37 ± 0.56 | 0.6571 | 1.45 ± 0.49 | 1.02 ± 0.50 | 0.1454 | 1.39 ± 0.73 | 1.05 ± 0.51 | 0.3926 |
| | EF (%) | 86.54 ± 6.95 | 86.59 ± 6.91 | 0.9810 | 35.70 ± 14.99 | 38.56 ± 14.54 | 0.6058 | 29.69 ± 9.77 | 35.73 ± 13.62 | 0.1757 | 27.70 ± 13.55 | 44.97 ± 20.68 | 0.0774 | 31.17 ± 16.00 | 34.66 ± 21.05 | 0.7282 | 25.70 ± 16.32 | 31.31 ± 15.82 | 0.5434 |
| | FS (%) | 55.19 ± 9.25 | 55.10 ± 8.85 | 0.9766 | 17.05 ± 8.00 | 18.69 ± 7.99 | 0.5838 | 13.90 ± 4.87 | 17.32 ± 7.18 | 0.1380 | 13.12 ± 6.77 | 23.00 ± 11.40 | 0.0571 | 15.03 ± 8.16 | 17.09 ± 10.86 | 0.6886 | 12.28 ± 8.55 | 15.05 ± 7.95 | 0.5629 |
| | LV Mass | 95.11 ± 37.30 | 89.90 ± 36.87 | 0.6680 | 115.02 ± 30.67 | 105.03 ± 21.45 | 0.3372 | 123.59 ± 31.48 | 116.79 ± 34.46 | 0.5869 | 105.45 ± 27.79 | 111.79 ± 29.17 | 0.6922 | 123.75 ± 33.23 | 100.74 ± 29.71 | 0.2302 | 120.60 ± 41.49 | 116.63 ± 43.73 | 0.8678 |
| | LV Mass (Corrected) | 76.09 ± 29.84 | 71.92 ± 29.50 | 0.6680 | 92.02 ± 24.54 | 84.03 ± 17.16 | 0.3372 | 98.87 ± 25.18 | 93.44 ± 27.57 | 0.5869 | 84.36 ± 22.23 | 89.43 ± 23.34 | 0.6922 | 99.00 ± 26.58 | 80.59 ± 23.76 | 0.2302 | 96.48 ± 33.19 | 93.30 ± 34.99 | 0.8678 |
| | LV Vol;d | 20.66 ± 7.92 | 19.80 ± 9.65 | 0.7642 | 59.02 ± 29.67 | 63.90 ± 27.41 | 0.6506 | 92.00 ± 41.77 | 97.54 ± 38.80 | 0.7204 | 114.85 ± 41.37 | 96.02 ± 60.73 | 0.4942 | 117.28 ± 47.58 | 104.21 ± 62.21 | 0.6613 | 126.20 ± 55.54 | 118.17 ± 68.92 | 0.8130 |
| | LV Vol;s | 2.77 ± 1.90 | 2.75 ± 2.20 | 0.9760 | 41.13 ± 28.20 | 41.95 ± 24.86 | 0.9352 | 67.55 ± 37.86 | 67.10 ± 37.63 | 0.9751 | 87.39 ± 46.32 | 63.30 ± 63.49 | 0.4220 | 86.65 ± 53.77 | 78.52 ± 64.49 | 0.8020 | 98.76 ± 56.70 | 89.84 ± 69.59 | 0.7960 |
| | HR (BPM) | 606.27 ± 55.47 | 588.13 ± 73.31 | 0.3929 | 693.08 ± 40.27 | 686.90 ± 49.45 | 0.7066 | 671.83 ± 44.90 | 683.46 ± 53.17 | 0.5302 | 687.28 ± 28.61 | 691.06 ± 18.51 | 0.7977 | 702.26 ± 23.17 | 696.31 ± 23.40 | 0.6527 | 710.70 ± 33.58 | 682.29 ± 61.94 | 0.2708 |
| Calculated parameters | IVS;d/LVPW;d | 0.93 ± 0.26 | 0.95 ± 0.27 | 0.8067 | 0.66 ± 0.28 | 0.82 ± 0.35 | 0.1738 | 0.67 ± 0.27 | 0.89 ± 0.40 | 0.0886 | 0.54 ± 0.32 | 0.70 ± 0.43 | 0.4354 | 0.40 ± 0.22 | 0.90 ± 0.80 | 0.0928 | 0.25 ± 0.12 | 0.90 ± 1.16 | 0.1115 |
| | IVS;s/LVPW;s | 0.87 ± 0.15 | 0.86 ± 0.15 | 0.8645 | 0.59 ± 0.29 | 0.79 ± 0.50 | 0.1730 | 0.55 ± 0.21 | 0.78 ± 0.50 | 0.1084 | 0.37 ± 0.22 | 0.64 ± 0.55 | 0.2048 | 0.29 ± 0.16 | 0.94 ± 1.13 | 0.1057 | 0.29 ± 0.21 | 0.25 ± 0.22 | 0.7575 |
| | Cardiac Output | 29.34 ± 10.29 | 29.12 ± 13.28 | 0.9548 | 26.00 ± 8.68 | 31.83 ± 10.85 | 0.1100 | 36.61 ± 11.73 | 44.72 ± 11.63 | 0.0784 | 40.17 ± 9.13 | 47.24 ± 9.73 | 0.2015 | 43.80 ± 10.47 | 36.84 ± 17.44 | 0.3546 | 39.01 ± 16.27 | 41.92 ± 15.04 | 0.7465 |
| | Stroke Volume | 17.89 ± 6.95 | 17.04 ± 8.11 | 0.7324 | 17.88 ± 5.55 | 21.94 ± 8.04 | 0.1072 | 24.45 ± 7.61 | 30.44 ± 7.96 | 0.0513 | 27.47 ± 5.76 | 32.72 ± 7.11 | 0.1566 | 30.63 ± 6.76 | 25.69 ± 12.08 | 0.3294 | 27.44 ± 10.88 | 28.33 ± 9.72 | 0.8809 |

Data represented as a mean ± SD. LAD, left anterior descending ligation; BPM, beats per minutes; s, systolic; d, diastolic; IVS, interventricular septal; LVID, left ventricle internal diameter; LVPW, left ventricle posterior wall; EF, ejection fraction; FS, fractional shortening; LV, left ventricle. Two-tailed t test. P values are shown.

**Supplementary Table 2.**

Differentially expressed genes between CD68-/LYVE1+ cardiac lymphatic endothelial cells (LECs) isolated from the hearts of *Ackr3*$^{\Delta Lyve1}$ male mice with no cardiac injury and 7 days post left anterior descending artery (LAD) ligation. N=3 samples per group. One-way ANOVA. -log10 p-values and log2 fold change values are shown. Positive fold change values represent upregulation in LAD ligated *Ackr3*$^{\Delta Lyve1}$ LECs.

| Gene Symbol | -log10 p-value | log2 fold change |
|---|---|---|
| Mmp3 | 5.183983 | 2.726716 |
| Htatip2 | 3.025439 | 2.724875 |
| Olfr1168 | 4.869438 | -2.411608 |
| Retnla | 2.293451 | -2.361224 |
| Chil4 | 2.62656 | 2.349578 |
| Hal | 2.222628 | 2.083996 |
| Ttc39c | 2.692474 | 2.064403 |
| Tmem252 | 3.14968 | 1.926516 |
| Angpt1 | 3.478361 | -1.903621 |
| Car6 | 2.232979 | 1.735973 |
| Plac8 | 2.270009 | 1.67157 |
| Amy2a4 | 3.346124 | -1.619404 |
| Apod | 2.651672 | 1.584698 |
| Slpi | 2.521201 | 1.583476 |
| Zbtb16 | 1.941825 | 1.576271 |
| Mcpt4 | 2.073508 | -1.52655 |
| Reck | 3.124179 | -1.514481 |
| Ms4a8a | 3.433425 | 1.510825 |
| Amy2a3 | 3.634915 | -1.504036 |
| P2ry12 | 3.274645 | 1.490242 |
| Ppbp | 2.612244 | 1.477299 |
| Usp29 | 2.313737 | -1.465708 |
| Olfr1278 | 2.774097 | -1.455439 |
| Napsa | 2.866084 | 1.446659 |
| Gm2927 | 3.836868 | -1.441563 |
| Olfr899 | 3.239452 | -1.438016 |
| C5ar2 | 2.067868 | 1.432286 |
| Olfr1124 | 4.202663 | -1.405693 |
| Gm17727 | 2.854906 | -1.37244 |
| Gm6377 | 3.149512 | 1.371698 |
| Gpr141 | 2.543821 | 1.368897 |
| Zdhhc23 | 3.972332 | 1.359206 |
| Olfr1431 | 4.495895 | -1.357963 |
| Rgs18 | 3.115069 | 1.332215 |
| Olfr203 | 3.713566 | -1.321305 |

| | | |
|---|---|---|
| F7 | 2.45143 | 1.313693 |
| Gjc3 | 2.587454 | -1.308996 |
| Fgr | 3.265592 | 1.306192 |
| Hepacam2 | 3.060236 | -1.296534 |
| Hhipl1 | 2.5893 | 1.292487 |
| Hcst | 2.622442 | 1.291073 |
| Bst1 | 3.653084 | 1.281075 |
| Fgf23 | 3.915635 | 1.27422 |
| Slfn1 | 3.312761 | 1.271175 |
| Mt2 | 2.799491 | 1.262656 |
| Olfr510 | 2.74814 | -1.246256 |
| Gm10719 | 2.819989 | -1.240259 |
| Gpr160 | 3.90689 | 1.239795 |
| Ltb4r1 | 2.521988 | 1.234587 |
| Osm | 4.087006 | 1.229016 |
| Gp6 | 2.241037 | 1.226262 |
| Gm21738 | 2.504864 | -1.226052 |
| Arhgef37 | 2.661551 | 1.221054 |
| Olfr1247 | 4.230113 | -1.213459 |
| Psd3 | 2.592818 | -1.212576 |
| Gm9733 | 2.709349 | 1.20916 |
| Fjx1 | 2.498979 | 1.201872 |
| Tmem108 | 2.305164 | -1.201176 |
| Tmem182 | 2.858757 | -1.199035 |
| Slc2a1 | 3.544135 | 1.1934 |
| Gsr | 3.084751 | 1.189388 |
| Arhgap15 | 2.289169 | 1.186437 |
| Abi3bp | 2.239641 | -1.178256 |
| Olfr1489 | 2.073075 | -1.174304 |
| Insl6 | 3.215053 | 1.172206 |
| Sobp | 3.650099 | -1.167152 |
| Nfam1 | 3.459668 | 1.165867 |
| Stc2 | 2.461376 | -1.160546 |
| Olfr482 | 2.502953 | -1.148674 |
| Plxnc1 | 3.188564 | -1.147704 |
| Nav3 | 2.21935 | -1.144386 |
| Pdk1 | 2.120629 | 1.131879 |
| Il17ra | 2.993064 | 1.126676 |
| Was | 4.342063 | 1.125651 |
| Gm2022 | 3.772471 | -1.124176 |
| 6030408B16Rik | 2.534361 | 1.122302 |
| Olfr153 | 2.867065 | -1.120942 |
| Lrp8 | 2.380511 | 1.117442 |

| | | |
|---|---|---|
| Gm11084 | 2.312544 | -1.113867 |
| Kcnc2 | 2.243519 | -1.113414 |
| Podn | 3.011722 | -1.106469 |
| Bnip3 | 3.544532 | 1.104806 |
| Paqr9 | 2.091229 | -1.089722 |
| Krt222 | 2.787046 | -1.086369 |
| Tmod4 | 2.067572 | -1.084656 |
| Mrc1 | 2.858619 | 1.084493 |
| Olfr1152 | 2.850178 | -1.08369 |
| 5430427O19Rik | 2.548945 | 1.075923 |
| Olfr615 | 2.78229 | -1.072071 |
| Eif4a2 | 2.276497 | -1.069922 |
| Itgb4 | 3.464574 | -1.069771 |
| Ptgs2 | 2.090561 | -1.066358 |
| Azin2 | 2.36981 | 1.065496 |
| Olfr292 | 2.813945 | -1.057291 |
| Arhgap28 | 3.316785 | -1.056382 |
| Ros1 | 3.668229 | -1.055758 |
| Mkrn1 | 2.836374 | 1.055425 |
| Slitrk6 | 2.909791 | -1.052492 |
| Aox1 | 2.014677 | -1.052485 |
| Olfr727 | 3.486219 | -1.051887 |
| Fgf11 | 2.070267 | -1.049714 |
| Cd14 | 3.778302 | 1.047078 |
| F13a1 | 2.421058 | 1.046037 |
| D130040H23Rik | 2.981258 | -1.045597 |
| Olfr469 | 3.877224 | -1.039819 |
| Tmem192 | 2.919864 | 1.037333 |
| Ptprj | 3.239605 | 1.034624 |
| Gm2777 | 3.051116 | -1.033349 |
| Tlr2 | 3.542962 | 1.033053 |
| Adgrg2 | 2.502737 | 1.03215 |
| Slc2a3 | 2.563405 | 1.031134 |
| Cd33 | 2.105742 | 1.028824 |
| Eya2 | 2.007178 | 1.026928 |
| Ccl21a | 2.530188 | -1.026701 |
| Irs1 | 2.910448 | -1.025844 |
| Spata1 | 3.481847 | 1.025249 |
| Lum | 2.024087 | 1.023688 |
| Lrrc4c | 2.137122 | -1.023241 |
| Sema5a | 3.323287 | -1.020449 |
| Tmem40 | 2.739671 | 1.019659 |
| Gm11034 | 2.122751 | -1.018072 |

| | | |
|---|---|---|
| Gm20873 | 4.13664 | -1.018 |
| Igfbp3 | 3.1215 | 1.017109 |
| Apba1 | 2.782906 | -1.016225 |
| Cnr2 | 2.348693 | 1.014291 |
| Slc9a9 | 2.693895 | 1.011431 |
| Klf12 | 2.015059 | -1.010515 |
| Olfr167 | 3.690809 | -1.006025 |
| Snx32 | 3.527442 | -1.004192 |
| P2rx1 | 2.79556 | 1.00394 |
| Gm2933 | 2.448668 | -1.000656 |
| Tra2a | 2.483294 | -0.996786 |
| Gm17535 | 2.690892 | -0.995513 |
| Olfr502 | 3.097418 | -0.992312 |
| Fpr2 | 2.61338 | 0.986804 |
| Agrn | 2.004114 | -0.9862 |
| Pcolce2 | 2.929187 | 0.983576 |
| Vmn1r57 | 4.205757 | -0.98305 |
| Smyd1 | 2.087818 | -0.981728 |
| Gm13242 | 2.072047 | -0.980523 |
| Gm8281 | 3.306567 | -0.977097 |
| Cdh3 | 2.684779 | 0.971612 |
| Cd300ld | 2.271252 | 0.969572 |
| Gm2046 | 3.248886 | -0.968364 |
| Cdh11 | 2.458438 | -0.967146 |
| Ckmt2 | 2.055849 | -0.960371 |
| Angptl1 | 2.889134 | -0.96026 |
| Izumo3 | 2.672719 | -0.959918 |
| Bcl2 | 3.143842 | -0.950804 |
| Gm10256 | 3.125705 | -0.950424 |
| Trpv4 | 3.592456 | 0.950028 |
| Srd5a1 | 2.306862 | 0.949542 |
| Olfr936 | 2.232206 | -0.948444 |
| Zfp827 | 2.321598 | -0.946716 |
| Arhgap22 | 1.993979 | 0.945982 |
| 4930402H24Rik | 2.243439 | -0.944641 |
| Olfr1480 | 2.401028 | -0.944424 |
| Gem | 3.290837 | -0.94164 |
| Masp1 | 2.126789 | 0.940046 |
| Vmn2r8 | 4.44843 | -0.939302 |
| Srek1 | 3.443679 | -0.939129 |
| Plvap | 2.592431 | -0.937962 |
| Gm13304 | 2.289097 | -0.936433 |
| Ccl21c | 2.289097 | -0.936433 |

| | | |
|---|---|---|
| Ahnak2 | 2.357669 | -0.935882 |
| Gm16513 | 4.160026 | -0.934826 |
| Nrros | 3.645734 | 0.933777 |
| Vmn1r158 | 1.919006 | -0.930548 |
| Gm2832 | 6.282439 | -0.919142 |
| Tlr5 | 2.411826 | 0.918043 |
| Ppp1r12b | 2.225782 | -0.916071 |
| Clec2f | 2.091728 | -0.912696 |
| Prr15 | 1.96461 | -0.905074 |
| Doc2b | 2.02332 | 0.904172 |
| Tbc1d15 | 1.989887 | 0.901935 |
| Olfr294 | 2.195239 | -0.901626 |
| Tmem104 | 2.930069 | 0.898054 |
| Plaur | 1.943453 | 0.897852 |
| Nt5m | 2.284317 | 0.896326 |
| Sgpp2 | 2.769625 | -0.89621 |
| Apcdd1 | 2.285501 | -0.895993 |
| Olfr1450 | 2.925787 | -0.89243 |
| Gpm6b | 3.452436 | 0.889769 |
| Ticam2 | 3.21585 | 0.888539 |
| Tarm1 | 2.051659 | 0.888016 |
| Prss35 | 3.431546 | 0.886535 |
| Gm1987 | 2.395937 | -0.885262 |
| Synm | 3.961809 | -0.884551 |
| Crlf1 | 3.521347 | 0.883277 |
| Ogt | 2.595969 | -0.883253 |
| St8sia2 | 2.553917 | -0.880262 |
| Fibin | 3.143202 | 0.878953 |
| Phospho2 | 2.697656 | 0.875277 |
| LOC100861707 | 2.819931 | -0.874946 |
| Arhgap6 | 2.104569 | -0.874545 |
| Sdpr | 2.447036 | -0.873742 |
| Gm3755 | 2.733493 | -0.872088 |
| Prox1 | 2.784019 | -0.871039 |
| Atp2a1 | 2.861445 | -0.870195 |
| Tpst1 | 3.014791 | -0.870163 |
| Tifa | 2.543854 | 0.86845 |
| Lpar1 | 3.424752 | 0.862804 |
| Cspg4 | 3.479916 | -0.862701 |
| Nr2f6 | 3.912052 | -0.861805 |
| Gm21951 | 1.915667 | -0.861122 |
| Gm7008 | 3.674753 | 0.860581 |
| Mamdc2 | 3.534635 | -0.86016 |

| | | |
|---|---|---|
| Fdxacb1 | 2.715858 | 0.858737 |
| Gucy1a3 | 3.055837 | -0.857965 |
| Ccl4 | 1.99748 | 0.857217 |
| Ppid | 2.156502 | -0.855495 |
| P4ha2 | 2.730044 | 0.853261 |
| Olfr1298 | 3.137522 | -0.851735 |
| Ecm1 | 2.902566 | 0.851551 |
| Vnn3 | 2.880038 | 0.850239 |
| P2rx7 | 2.34287 | 0.849655 |
| Camk2a | 2.707367 | -0.849191 |
| 1700029J07Rik | 2.429857 | -0.848798 |
| Trim34b | 2.06214 | -0.847652 |
| Olfr1112 | 1.952939 | -0.843389 |
| Olfr1277 | 3.209941 | -0.83971 |
| Gm3376 | 2.305602 | -0.838048 |
| Tspan18 | 2.103933 | -0.837612 |
| Nr4a2 | 2.784532 | -0.836603 |
| 4930503L19Rik | 2.60837 | -0.836255 |
| Vmn1r129 | 2.088505 | -0.835771 |
| Krt8 | 2.003072 | 0.833076 |
| Stxbp4 | 1.978811 | -0.82786 |
| Zfp616 | 2.353235 | -0.825468 |
| Stap2 | 2.486756 | -0.825159 |
| Cobl | 2.126079 | -0.824882 |
| Cyp3a41b | 2.456113 | -0.824238 |
| Enpp3 | 2.082534 | -0.822959 |
| Tmpo | 3.968458 | -0.817893 |
| Zfp14 | 2.43205 | -0.816247 |
| N4bp2 | 2.450837 | -0.816125 |
| Map2k4 | 2.014662 | 0.812909 |
| Huwe1 | 2.618376 | -0.812268 |
| Creg2 | 2.539806 | 0.810591 |
| Qsox1 | 3.092846 | 0.80934 |
| Tcea3 | 2.061578 | -0.808031 |
| Spint2 | 3.118493 | -0.807157 |
| Bcl6 | 2.037448 | 0.806357 |
| Gdf11 | 2.429789 | -0.803665 |
| Vmn1r173 | 2.578674 | -0.801945 |
| E330017A01Rik | 3.458828 | -0.801465 |
| Tubb1 | 1.940615 | 0.801407 |
| Egln3 | 1.967103 | 0.800032 |
| Samd5 | 2.048903 | -0.799833 |
| Neto2 | 2.454467 | -0.799079 |

| | | |
|---|---|---|
| Iqgap2 | 2.087611 | 0.79854 |
| Fhl1 | 1.998773 | -0.796332 |
| Rrad | 2.336272 | -0.795925 |
| Plxdc1 | 3.096381 | -0.795817 |
| Ndn | 2.662084 | -0.795443 |
| Psmd9 | 2.151989 | 0.79417 |
| Gm14346 | 2.284572 | -0.793571 |
| Olfr1216 | 2.722994 | -0.792914 |
| Mptx1 | 2.730735 | -0.792639 |
| Cdkl5 | 2.350278 | -0.791806 |
| Zfp329 | 2.847207 | -0.790847 |
| Hes6 | 2.057769 | 0.789554 |
| Bin2 | 2.759488 | 0.789003 |
| 1110057P08Rik | 2.273281 | -0.788894 |
| Cd300lf | 2.229311 | 0.788753 |
| Sytl2 | 2.933663 | -0.787984 |
| Snca | 3.106037 | 0.787232 |
| Olfr1463 | 2.134432 | -0.787031 |
| Krtcap3 | 4.537719 | -0.785517 |
| Olfr1312 | 2.17539 | -0.785124 |
| Gpr84 | 2.972544 | 0.784864 |
| Snx24 | 2.086328 | 0.78447 |
| Vmn1r86 | 2.905892 | -0.783281 |
| Baz1b | 3.85888 | -0.781603 |
| Igfbp5 | 1.990791 | -0.781528 |
| Adam4 | 2.700216 | -0.781351 |
| Ccdc7b | 4.214546 | -0.779848 |
| Igfbp6 | 2.103047 | -0.77841 |
| Rit2 | 3.158041 | -0.777679 |
| Itgb2l | 2.696612 | 0.776357 |
| BC048502 | 2.814574 | -0.772147 |
| Wsb2 | 2.343375 | 0.770744 |
| Fgf10 | 2.132943 | -0.770601 |
| Ocln | 2.638805 | -0.770584 |
| Olfr1465 | 3.485062 | -0.770288 |
| Cnp | 2.919893 | 0.769213 |
| Adck4 | 2.457022 | 0.767358 |
| Olfml1 | 3.399599 | 0.766883 |
| Olfr1180 | 3.274088 | -0.766341 |
| LOC100503048 | 2.420556 | -0.761336 |
| Il10ra | 2.599963 | 0.760792 |
| 6430573F11Rik | 4.011937 | -0.760502 |
| Ccrl2 | 2.38549 | 0.757749 |

| | | |
|---|---|---|
| Gm13298 | 3.205458 | -0.756639 |
| Fbp1 | 2.069113 | 0.756408 |
| Arhgap42 | 3.195665 | -0.756152 |
| Inpp4b | 2.188006 | -0.755811 |
| Olfr1384 | 5.578169 | -0.75363 |
| Gpm6a | 1.954931 | -0.753579 |
| Aldh1a1 | 2.361352 | 0.752723 |
| Dennd2d | 2.148237 | -0.752543 |
| Klhl2 | 2.074054 | 0.752457 |
| Olfr1131 | 2.962255 | -0.750495 |
| Fbn2 | 3.353939 | 0.750195 |
| Fsd1l | 3.102333 | -0.749835 |
| Olfr193 | 3.735015 | -0.74974 |
| Nav1 | 2.361695 | -0.748161 |
| Zdhhc14 | 3.130376 | 0.747637 |
| Pbrm1 | 5.570001 | -0.747319 |
| Nsg1 | 2.031487 | 0.746115 |
| Olfr948 | 2.901391 | -0.745452 |
| Ppp1r16b | 2.111406 | -0.744695 |
| Hnrnph1 | 3.225825 | -0.742248 |
| Akr1c19 | 2.177978 | -0.740936 |
| Pigg | 2.433121 | -0.740116 |
| Zfp532 | 2.495116 | -0.740107 |
| Olfr492 | 1.942292 | -0.733615 |
| Gm5885 | 2.133898 | -0.732417 |
| Chid1 | 3.025054 | 0.732113 |
| Lpp | 2.539645 | -0.731383 |
| Prune2 | 3.072664 | 0.730444 |
| Pik3cg | 1.957373 | 0.730236 |
| Olfr968 | 2.869834 | -0.730236 |
| Ccr3 | 1.95481 | 0.730192 |
| Ssc5d | 3.152529 | 0.729609 |
| Ptprz1 | 2.159684 | -0.729305 |
| Fads2 | 1.964778 | -0.728504 |
| Adamts1 | 2.119541 | -0.72684 |
| Epsti1 | 2.1382 | 0.726613 |
| Olfr618 | 3.177696 | -0.726264 |
| 0610037L13Rik | 3.629359 | -0.724214 |
| Med8 | 3.501383 | 0.723698 |
| Olfr1471 | 2.934794 | -0.722055 |
| Tmem207 | 2.349329 | -0.721959 |
| Olfr1285 | 3.03482 | -0.721906 |
| Ddhd1 | 2.125034 | 0.72132 |

| | | |
|---|---|---|
| Borcs6 | 1.981092 | 0.719858 |
| Bambi | 2.578547 | -0.719806 |
| Tas2r125 | 2.313473 | -0.716271 |
| Cbx5 | 2.453965 | -0.715893 |
| Sox6 | 2.142841 | -0.714971 |
| Pih1d3 | 4.373738 | -0.714909 |
| Zbtb21 | 3.834165 | -0.713379 |
| Olfr474 | 2.41172 | -0.712957 |
| Gm5097 | 2.665273 | -0.712358 |
| Bdnf | 2.107521 | 0.712252 |
| Exoc4 | 2.517318 | -0.710904 |
| Fam208a | 2.441696 | -0.710287 |
| Olfr342 | 2.36377 | -0.709167 |
| Rora | 2.553663 | -0.708655 |
| Atl1 | 2.785559 | -0.706093 |
| Gm10600 | 2.215428 | -0.705721 |
| Olfr954 | 2.369018 | -0.704677 |
| 4930404N11Rik | 3.578727 | 0.704482 |
| Jmjd1c | 2.72132 | -0.703721 |
| Sphk1 | 2.112589 | 0.703623 |
| Olfr470 | 3.202272 | -0.703499 |
| Tmco5 | 3.559332 | -0.702994 |
| Shroom3 | 2.364023 | 0.699809 |
| Baalc | 2.470802 | 0.699347 |
| Tppp | 2.476467 | -0.69901 |
| Tmem189 | 2.099756 | 0.69853 |
| Prg4 | 2.085347 | 0.697987 |
| Fam166a | 2.194949 | -0.697952 |
| Eif4g3 | 4.200291 | -0.696065 |
| Slc36a1 | 2.272207 | 0.69562 |
| Olfr1061 | 2.159928 | -0.694729 |
| Fam120b | 2.47811 | -0.694541 |
| Dmd | 2.584169 | -0.692838 |
| Kcnmb1 | 2.047513 | -0.692534 |
| Gm3750 | 2.180686 | -0.692374 |
| Vmn2r72 | 2.177082 | -0.691597 |
| Dock4 | 2.306682 | -0.691391 |
| Saa4 | 2.245353 | 0.690641 |
| Csgalnact2 | 3.097553 | 0.689111 |
| Pcdhb9 | 1.921155 | 0.68887 |
| Nisch | 2.630953 | -0.688691 |
| P4ha3 | 2.297983 | 0.687777 |
| Rif1 | 3.621929 | -0.687267 |

| | | |
|---|---|---|
| Trem3 | 1.95124 | 0.686102 |
| Rttn | 3.132406 | -0.685859 |
| Nkd2 | 2.239584 | 0.684056 |
| Mettl13 | 2.536198 | 0.680612 |
| Gm16429 | 4.099664 | -0.679163 |
| Gm7971 | 4.099664 | -0.679163 |
| Cttnbp2 | 3.384154 | -0.679154 |
| Lpin3 | 1.92578 | 0.679117 |
| Eid2b | 2.157726 | 0.678207 |
| Olfr481 | 3.251622 | -0.67652 |
| Ero1l | 2.027711 | 0.675888 |
| Rps13 | 2.358542 | 0.675256 |
| Medag | 3.442971 | 0.674723 |
| Tubb3 | 2.594787 | 0.674506 |
| Gm4832 | 2.070041 | 0.673538 |
| Gm13247 | 3.639933 | -0.673348 |
| Chn1 | 2.400767 | -0.672398 |
| Galnt11 | 2.914809 | -0.672299 |
| Metap1d | 2.220494 | -0.672045 |
| Dclk2 | 3.295798 | -0.671574 |
| Vezt | 2.086259 | -0.670967 |
| Slc2a9 | 2.884127 | 0.668936 |
| Ocm | 3.292613 | 0.665884 |
| Gm2964 | 2.143406 | -0.665047 |
| Vmn2r68 | 2.337613 | -0.664901 |
| Ddx56 | 3.207925 | 0.664501 |
| Sel1l | 2.07833 | 0.663117 |
| Vmn1r29 | 2.919081 | -0.662862 |
| Gm6465 | 2.750491 | -0.662853 |
| Metrn | 4.732899 | 0.662762 |
| Lao1 | 1.922142 | 0.662023 |
| Chmp2b | 2.926663 | 0.661111 |
| Cercam | 2.935389 | 0.660235 |
| D16Ertd472e | 2.022305 | 0.659788 |
| Asph | 3.289593 | -0.6586 |
| Cyp3a41a | 2.864212 | -0.657869 |
| Sirt5 | 2.929844 | -0.657466 |
| Ncam1 | 2.055295 | -0.657292 |
| Olfr1098 | 2.881441 | -0.656332 |
| Tug1 | 3.602735 | -0.656158 |
| Dhdh | 2.166321 | -0.654417 |
| Kbtbd12 | 1.961193 | -0.653794 |
| Alms1 | 3.088897 | -0.653583 |

| | | |
|---|---|---|
| Gyg | 2.354596 | 0.652555 |
| Noc4l | 2.100839 | 0.652427 |
| Fstl3 | 2.728553 | 0.652298 |
| Astn2 | 2.784653 | -0.652216 |
| Mllt3 | 3.089339 | -0.651931 |
| Gtf2i | 2.618445 | -0.649202 |
| AA474331 | 2.101463 | 0.648898 |
| Brwd1 | 2.579416 | -0.646974 |
| Zfp948 | 2.250096 | -0.646771 |
| Zfp663 | 3.494276 | 0.645923 |
| Gm17449 | 2.483835 | -0.644807 |
| Ankrd29 | 3.354701 | 0.644419 |
| Gm10922 | 2.43921 | -0.644345 |
| Tnfaip8 | 2.488459 | 0.643949 |
| Fbln2 | 3.336894 | 0.643016 |
| Lin7a | 3.269709 | -0.641981 |
| Lgalsl | 2.368927 | 0.639936 |
| Olfr930 | 2.630476 | -0.639621 |
| Tmem2 | 2.124464 | -0.639371 |
| Pard3b | 3.273791 | -0.638482 |
| Zfp280d | 2.611893 | -0.638 |
| Nkx2-1 | 2.760145 | 0.637592 |
| Gm3115 | 2.192575 | -0.637564 |
| Pradc1 | 2.064205 | 0.637332 |
| AA414768 | 2.316344 | 0.636191 |
| Sval3 | 3.149645 | -0.634733 |
| Gm6455 | 2.253076 | -0.633952 |
| Tuft1 | 2.244087 | 0.633403 |
| Olfr1132 | 2.381555 | -0.631095 |
| Emilin3 | 2.774432 | 0.630899 |
| 4921511C20Rik | 3.871316 | -0.63034 |
| Cphx2 | 3.227963 | -0.628839 |
| Gm15737 | 2.511611 | -0.628699 |
| Gm884 | 2.596127 | -0.628176 |
| 4931414P19Rik | 3.100106 | 0.627812 |
| Gm10521 | 2.741667 | 0.62728 |
| Ada | 2.420658 | 0.627046 |
| Gm20877 | 2.095607 | -0.626439 |
| Mak | 1.981458 | 0.625934 |
| Parvb | 4.244492 | 0.625916 |
| Slc7a3 | 3.620249 | -0.625785 |
| Mcoln2 | 4.08524 | 0.625373 |
| Zkscan14 | 2.605589 | 0.625214 |

| | | |
|---|---|---|
| Bicc1 | 3.454747 | 0.624297 |
| Gm7970 | 2.46208 | -0.623221 |
| Rgs13 | 2.098263 | -0.623005 |
| Crtac1 | 2.058548 | 0.622199 |
| Slc25a23 | 2.278137 | -0.622199 |
| Ube3c | 3.983247 | -0.622012 |
| Gm5861 | 2.991596 | -0.621337 |
| Ankrd26 | 3.479857 | -0.619037 |
| Tmem154 | 2.340195 | 0.618972 |
| Pabpc1l | 2.746277 | 0.617336 |
| Rph3al | 2.748479 | -0.616885 |
| D1Ertd622e | 3.18274 | 0.616772 |
| Ifnz | 2.312218 | 0.61536 |
| Olfr1196 | 2.28504 | -0.615153 |
| Rarres2 | 2.564831 | -0.614597 |
| Col4a2 | 2.475732 | 0.61438 |
| Olfr1141 | 2.675103 | -0.614248 |
| Plk3 | 2.396682 | -0.614012 |
| Ppm1l | 1.973226 | -0.612966 |
| Trim37 | 3.013014 | -0.61257 |
| Gm8127 | 2.925516 | -0.611427 |
| Trnp1 | 2.541176 | 0.610955 |
| Csf1r | 2.407767 | 0.610407 |
| Luzp2 | 2.654895 | -0.610209 |
| Gm6337 | 2.234717 | -0.61019 |
| Efhb | 1.944331 | -0.609972 |
| Gm20026 | 3.517059 | 0.609538 |
| Olfr994 | 2.353133 | -0.609329 |
| Chic1 | 2.03676 | -0.608242 |
| Plekhb1 | 2.4753 | -0.608005 |
| Olfr703 | 3.292054 | 0.607778 |
| Smad7 | 2.123544 | -0.60774 |
| Olfr242 | 2.049507 | -0.607636 |
| Gm15448 | 2.855815 | 0.606347 |
| Top1 | 2.450309 | -0.602419 |
| Gm9944 | 2.146178 | -0.6024 |
| Tsc22d4 | 2.940615 | 0.600679 |
| Zfp672 | 2.060271 | 0.60066 |
| Gm16432 | 2.161863 | -0.600412 |
| Med22 | 2.007028 | 0.599984 |
| Fosl2 | 2.930969 | 0.598899 |
| Gtf2h3 | 2.037688 | -0.598499 |
| Ikbkg | 1.978125 | 0.598241 |

| | | |
|---|---|---|
| Nlk | 2.709807 | -0.598232 |
| Wfdc18 | 2.152844 | 0.598136 |
| Nrxn3 | 2.954759 | 0.597841 |
| Xcr1 | 1.958532 | -0.597765 |
| Slc35f1 | 2.443898 | -0.597069 |
| Lrrc10b | 2.824349 | 0.596935 |
| Esf1 | 4.771535 | -0.596859 |
| Apoe | 2.314689 | 0.596 |
| Parp14 | 1.91817 | -0.595981 |
| Hs6st2 | 2.399092 | 0.595494 |
| Gm8356 | 2.393736 | -0.595494 |
| Vmn1r39 | 3.243973 | -0.595103 |
| Timm21 | 2.001075 | -0.594969 |
| Ice2 | 2.149664 | -0.594577 |
| Il9 | 2.839766 | -0.594453 |
| Inppl1 | 2.563032 | -0.593268 |
| Wisp1 | 2.300564 | 0.592742 |
| Rom1 | 2.446181 | 0.59212 |
| Gda | 4.08199 | 0.591612 |
| Cgref1 | 2.756367 | 0.590271 |
| Cyp8b1 | 2.420348 | 0.589102 |
| Tm2d3 | 2.508432 | 0.588699 |
| Pclo | 1.99889 | -0.588267 |
| Cyp3a11 | 2.075308 | -0.587778 |
| Htr1b | 2.44025 | 0.58724 |
| Ptrh1 | 1.951624 | 0.586251 |
| F12 | 2.451462 | 0.585943 |
| Oxnad1 | 3.388994 | -0.585914 |
| Birc3 | 1.920623 | 0.585357 |
| Pkp3 | 2.15144 | 0.585155 |
| Ace | 2.510593 | 0.585078 |
| Cluh | 2.489397 | -0.583798 |
| Gm15850 | 2.711632 | 0.583192 |
| Gmcl1 | 2.844661 | -0.583172 |
| Gm5801 | 2.764446 | 0.582874 |
| Prkab2 | 2.59456 | -0.582373 |
| Ccdc96 | 3.788412 | 0.582325 |
| Fam187b | 1.960586 | 0.582151 |
| Trpc7 | 2.046164 | 0.582074 |
| Sdr16c6 | 2.165909 | -0.581881 |
| Gm15107 | 2.96194 | -0.581554 |
| Hars | 2.936441 | 0.580724 |
| Ube2cbp | 2.699605 | 0.58056 |

| | | |
|---|---|---|
| Gm5724 | 3.095123 | -0.579972 |
| P2ry14 | 2.437334 | -0.579055 |
| Tctn3 | 2.466946 | -0.578977 |
| Rpp38 | 2.670472 | 0.577953 |
| Acbd4 | 2.864276 | -0.577886 |
| Vmn1r234 | 2.2802 | -0.577576 |
| Ssxb10 | 2.857789 | -0.57747 |
| Git1 | 2.109252 | 0.577431 |
| Pllp | 2.451677 | 0.576909 |
| Zkscan4 | 2.451091 | 0.575554 |
| Bmper | 2.032676 | 0.575332 |
| Pcdhb16 | 1.928914 | 0.574712 |
| Stambp | 2.602182 | 0.573665 |
| Ppfibp2 | 1.938827 | 0.571852 |
| Snx18 | 2.911821 | 0.571395 |
| Tex101 | 2.728651 | 0.569929 |
| Scn3a | 2.067258 | -0.569413 |
| Clcc1 | 2.952585 | 0.567799 |
| Sgcd | 1.981387 | -0.567769 |
| Syk | 2.52971 | 0.567672 |
| Olfr724 | 2.142701 | -0.567643 |
| Lrsam1 | 2.664293 | -0.567039 |
| Def8 | 1.973839 | 0.566864 |
| Eif2a | 2.309106 | 0.566718 |
| Defa22 | 2.226476 | 0.56664 |
| Gm21767 | 3.36739 | -0.565958 |
| Gm21822 | 3.36739 | -0.565958 |
| Slc3a2 | 3.199316 | 0.565529 |
| Gm766 | 2.484832 | -0.565022 |
| Rfx3 | 2.388209 | -0.564632 |
| Amfr | 2.55258 | 0.564115 |
| Dus4l | 3.340947 | -0.563715 |
| Gm2244 | 2.053813 | -0.563597 |
| Ldlrad4 | 2.25654 | -0.563217 |
| Gm3278 | 1.987568 | -0.562777 |
| Parn | 2.518367 | 0.562719 |
| Prrc2a | 2.541914 | -0.562142 |
| Olfr1109 | 2.543119 | -0.561439 |
| Treml2 | 1.970539 | 0.561331 |
| Zeb2os | 1.953723 | 0.560725 |
| Pex11b | 1.935579 | 0.560177 |
| Fshb | 2.406789 | -0.559746 |
| Rfc1 | 3.982887 | -0.559365 |

| | | |
|---|---|---|
| Pla2g1b | 3.295487 | 0.559316 |
| Olfr745 | 2.376371 | -0.559237 |
| Celsr1 | 2.190014 | -0.558953 |
| Slc16a12 | 3.300998 | -0.558806 |
| Gbf1 | 2.738504 | -0.558032 |
| Gm10046 | 2.130159 | -0.557925 |
| Tcp10c | 2.335953 | -0.557503 |
| Mmaa | 2.016057 | -0.55716 |
| Zfyve28 | 2.293949 | 0.556709 |
| Anapc1 | 2.539753 | -0.555512 |
| Olfr373 | 2.262156 | -0.555021 |
| Lmo4 | 2.956013 | 0.55451 |
| Tecta | 3.636523 | 0.553832 |
| Paqr8 | 2.400844 | -0.553449 |
| Ces2a | 2.035463 | -0.553144 |
| Olfr350 | 2.129586 | -0.552761 |
| 2010111I01Rik | 2.635025 | -0.552633 |
| Cln6 | 2.064561 | 0.552485 |
| B9d2 | 2.018839 | 0.551669 |
| Gm21797 | 2.52238 | -0.551245 |
| Palm2 | 1.963215 | -0.550891 |
| Gfpt1 | 2.506266 | -0.550526 |
| Pgs1 | 2.469684 | 0.550408 |
| Irak2 | 2.178439 | 0.549847 |
| Gm10097 | 2.042712 | -0.549768 |
| Klhl36 | 2.106034 | 0.549413 |
| Olfr1188 | 1.969676 | 0.548131 |
| AI413582 | 2.460999 | 0.547894 |
| Dpp7 | 1.943133 | 0.546818 |
| Mcfd2 | 2.358853 | 0.546532 |
| Cr2 | 1.963615 | -0.546433 |
| Chrng | 4.991247 | 0.546324 |
| Dck | 1.945402 | 0.545197 |
| 1700008O03Rik | 2.337474 | 0.544812 |
| Tyw1 | 3.045976 | 0.54312 |
| Gm21891 | 3.04029 | -0.542327 |
| Gm21828 | 3.04029 | -0.542327 |
| Gm21725 | 3.04029 | -0.542327 |
| Gm21904 | 3.04029 | -0.542327 |
| Gm21764 | 3.04029 | -0.542327 |
| Gm21852 | 3.04029 | -0.542327 |
| Dalrd3 | 4.517352 | -0.541931 |
| Nfe2 | 2.404474 | 0.541812 |

| | | |
|---|---|---|
| Cep70 | 3.984611 | -0.541792 |
| Ptges3l | 1.937546 | 0.541584 |
| Atf7ip | 2.071693 | -0.541247 |
| Cfap20 | 2.552859 | -0.541039 |
| Ces1c | 2.140046 | -0.540622 |
| Ncs1 | 3.246064 | 0.539551 |
| Mdh1 | 2.728632 | -0.539333 |
| Mtss1 | 2.437548 | -0.538707 |
| Klhl32 | 2.241274 | -0.538697 |
| Gm10339 | 2.255648 | -0.538508 |
| H13 | 1.951671 | 0.53834 |
| Nudt4 | 3.48178 | -0.53827 |
| Sp110 | 2.319453 | 0.538051 |
| Gm17174 | 3.448006 | -0.538032 |
| Dusp5 | 2.341778 | -0.537902 |
| Olfr1331 | 1.988895 | -0.53662 |
| Itpripl2 | 2.741554 | 0.535555 |
| Parp8 | 2.844433 | -0.535525 |
| Dner | 2.655486 | -0.535436 |
| Slc12a8 | 2.428486 | 0.535376 |
| Vmn1r44 | 2.868866 | -0.534689 |
| Ppp1r18 | 2.912765 | 0.534261 |
| Foxf1 | 1.956889 | 0.533683 |
| Serpini2 | 1.996889 | -0.533603 |
| Prss36 | 2.410504 | -0.533384 |
| Olfr38 | 2.416496 | -0.532826 |
| Spata6 | 3.508763 | 0.532706 |
| F10 | 2.047296 | 0.532487 |
| Lace1 | 2.291397 | -0.532277 |
| Cntn1 | 2.832296 | -0.531619 |
| Vmn2r104 | 2.664297 | -0.531519 |
| Sult2a7 | 2.091063 | -0.531259 |
| Itch | 2.705721 | 0.53053 |
| Armcx6 | 2.688626 | -0.529441 |
| Gm21800 | 1.99892 | -0.529171 |
| Slc47a1 | 2.370636 | -0.528721 |
| Cerkl | 2.125289 | -0.528201 |
| Cdkal1 | 2.050117 | -0.527821 |
| Borcs5 | 1.917308 | 0.52719 |
| Tesc | 3.274438 | 0.526239 |
| Snrk | 2.025393 | -0.526069 |
| Apbb2 | 2.030661 | -0.525989 |
| Nr2f2 | 1.999974 | -0.525548 |

| | | |
|---|---|---|
| Gm3106 | 2.780099 | -0.524906 |
| Gm8267 | 3.037037 | -0.524545 |
| Htra3 | 2.663762 | 0.523973 |
| Olfr1487 | 3.382888 | -0.523281 |
| Trp53bp1 | 2.335421 | -0.523251 |
| Zfp804a | 2.14719 | -0.52308 |
| Smarca5 | 4.391691 | -0.522608 |
| Fam71f1 | 2.655759 | 0.522568 |
| BC017643 | 3.140371 | 0.522528 |
| Gm14496 | 2.305005 | -0.522146 |
| Tmprss11g | 2.216004 | -0.521523 |
| D830013O20Rik | 3.7621 | -0.521071 |
| Adgrl2 | 1.95474 | -0.520729 |
| Slc6a14 | 2.900292 | 0.520669 |
| Rgs12 | 2.0547 | 0.520628 |
| Rxfp2 | 2.566584 | 0.520568 |
| Trim10 | 1.977365 | -0.520347 |
| Zfp955b | 3.036803 | -0.518857 |
| Pstpip2 | 2.238146 | 0.518696 |
| Gm10972 | 3.295458 | -0.518303 |
| Slc24a2 | 2.16296 | -0.517628 |
| Gm13084 | 2.397357 | -0.517276 |
| Bahcc1 | 2.547904 | -0.517276 |
| Krt5 | 2.077141 | 0.515995 |
| E130114P18Rik | 3.168237 | -0.514844 |
| Gm10323 | 2.98046 | -0.514844 |
| Gabre | 2.381347 | 0.514814 |
| Gpihbp1 | 2.344075 | 0.514622 |
| Celf3 | 2.580328 | 0.514016 |
| Hps6 | 3.519543 | 0.51342 |
| Pcdhb6 | 2.334869 | 0.513147 |
| Dnah7c | 3.00284 | -0.513076 |
| Slc16a13 | 1.936764 | -0.511923 |
| Sbno2 | 3.193259 | 0.511154 |
| Ptger1 | 2.524462 | 0.511073 |
| Tax1bp1 | 2.533116 | 0.511063 |
| Slc6a15 | 1.948446 | -0.510516 |
| Rac3 | 2.074521 | 0.510111 |
| Frmd6 | 2.798469 | -0.50922 |
| Tmem151a | 2.544196 | 0.509118 |
| Scand1 | 3.185772 | 0.508966 |
| Casc3 | 3.622949 | -0.508885 |
| Gm26616 | 2.331011 | 0.508317 |

| | | |
|---|---|---|
| Braf | 2.862352 | -0.507891 |
| Smco2 | 2.420352 | -0.507079 |
| Mcpt8 | 1.991306 | -0.506643 |
| Gabrb3 | 2.360653 | 0.506155 |
| Plaa | 2.085239 | 0.505982 |
| Rnf113a1 | 2.019486 | 0.505515 |
| Gm906 | 2.017986 | -0.504966 |
| Parva | 2.031528 | -0.504702 |
| Slc35f6 | 2.382462 | 0.504254 |
| Mmadhc | 4.350178 | 0.504173 |
| B3gnt3 | 1.911131 | 0.504081 |
| Cds1 | 2.041137 | 0.504071 |
| Nln | 3.908949 | 0.503664 |
| Olfr357 | 2.640514 | -0.503379 |
| Uhrf2 | 2.483277 | -0.502463 |
| Gm16367 | 2.350099 | -0.502066 |
| Slc19a2 | 2.730989 | 0.501006 |
| Cyp39a1 | 2.254009 | -0.500068 |
| Gm17641 | 1.929301 | 0.499445 |
| Ncstn | 2.50537 | 0.498721 |
| Thoc2 | 2.216994 | -0.497566 |
| Tcp11x2 | 2.750044 | -0.496902 |
| Capn1 | 3.699676 | 0.496585 |
| Treh | 3.685882 | 0.496493 |
| Olfr1206 | 2.158949 | -0.496104 |
| Srxn1 | 2.068787 | 0.495746 |
| Msl3l2 | 2.243766 | -0.495173 |
| Gm21814 | 2.84274 | -0.495091 |
| Tmem82 | 2.525698 | 0.494559 |
| Il6st | 2.080933 | -0.494282 |
| Utp18 | 2.202089 | 0.494119 |
| Klhl23 | 2.141204 | -0.492469 |
| Urad | 2.918264 | 0.492202 |
| Tcp10b | 2.324614 | -0.492151 |
| Trib1 | 2.173761 | 0.492079 |
| Rtel1 | 3.298509 | -0.491925 |
| Kif1b | 2.24035 | -0.491884 |
| Dsg4 | 2.816611 | -0.491822 |
| Vwa8 | 2.560128 | -0.491648 |
| Usp20 | 2.37372 | -0.491042 |
| Bri3 | 2.04532 | 0.49056 |
| Zbtb48 | 2.534327 | -0.489183 |
| Cyp2d26 | 2.718526 | 0.488978 |

| | | |
|---|---|---|
| Gm4787 | 2.304021 | -0.488906 |
| Cdk17 | 2.698 | -0.488628 |
| Nabp2 | 2.079454 | 0.488587 |
| Myo7b | 2.188815 | 0.488525 |
| Itga11 | 2.774944 | 0.488196 |
| Pde8b | 2.889414 | -0.488145 |
| Rest | 2.310738 | 0.487949 |
| Vat1 | 2.197572 | 0.486529 |
| Kif13b | 2.03422 | -0.486302 |
| Arhgef12 | 2.318197 | -0.48623 |
| Nipal1 | 2.113384 | 0.48586 |
| Casp1 | 1.920869 | 0.485684 |
| Smu1 | 2.601376 | 0.485643 |
| Gpi1 | 3.197159 | 0.485252 |
| Gm21964 | 2.159594 | 0.484953 |
| Scaper | 2.201403 | -0.484076 |
| Ddhd2 | 2.27898 | -0.483282 |
| Vmn2r90 | 3.024525 | -0.482611 |
| Lman1l | 2.409088 | 0.482601 |
| Dnal1 | 1.963958 | -0.481485 |
| Gm21736 | 3.147188 | -0.481082 |
| Vmn2r50 | 2.279344 | -0.480544 |
| Hao2 | 3.518173 | -0.480151 |
| Olfr1008 | 2.705053 | -0.480131 |
| Vmn2r35 | 2.464011 | -0.479934 |
| Muc15 | 3.20452 | -0.4798 |
| Hilpda | 1.927287 | 0.479593 |
| Gpr150 | 2.541281 | 0.479293 |
| Gon4l | 3.12147 | -0.478982 |
| Dnajc3 | 1.916286 | 0.47891 |
| Oog1 | 2.207709 | -0.478061 |
| Agtrap | 2.879245 | 0.476869 |
| Mvd | 2.022888 | 0.476589 |
| Tcl1b2 | 2.363635 | 0.476247 |
| 9430038I01Rik | 2.15592 | 0.476081 |
| AW551984 | 2.072924 | -0.475697 |
| Epcam | 2.43152 | -0.475583 |
| B830017H08Rik | 3.148929 | 0.475261 |
| Hook2 | 2.707215 | 0.475137 |
| Gpx5 | 2.035796 | -0.474649 |
| Otud4 | 2.268108 | -0.474576 |
| Hs1bp3 | 4.082709 | -0.47414 |
| Ttc14 | 2.014737 | -0.474067 |

| | | |
|---|---|---|
| Rasgef1a | 2.311982 | -0.473766 |
| Gm17577 | 2.432627 | -0.473683 |
| Gm17467 | 2.432627 | -0.473683 |
| Sphkap | 2.411806 | -0.473569 |
| Adamts19 | 2.132807 | -0.473267 |
| Mst1r | 2.872879 | 0.472498 |
| Qrsl1 | 2.298311 | 0.472467 |
| 4931408C20Rik | 2.329384 | 0.471957 |
| Msc | 2.578982 | 0.471874 |
| Gm270 | 2.629163 | -0.471562 |
| Spsb4 | 2.282231 | 0.471531 |
| Oxa1l | 2.452747 | 0.47126 |
| Adar | 2.299811 | -0.471208 |
| Cyp4f40 | 1.917592 | 0.470698 |
| Gm19475 | 1.940592 | 0.470511 |
| Zbtb9 | 2.009341 | -0.470417 |
| Slc39a14 | 4.06316 | 0.470032 |
| Mlst8 | 2.439561 | -0.469167 |
| Crispld2 | 2.102354 | 0.468364 |
| Zcchc16 | 2.018121 | 0.467791 |
| Ddx52 | 2.183631 | 0.467457 |
| H2-Ea-ps | 2.17278 | 0.466591 |
| C1qtnf4 | 2.157436 | 0.466131 |
| Sdr39u1 | 2.01552 | -0.465912 |
| Olfr314 | 3.065509 | 0.46586 |
| Defa-rs2 | 2.971628 | 0.465557 |
| Olfr810 | 2.974473 | -0.463936 |
| Lins1 | 2.831791 | -0.463727 |
| Sfrp2 | 2.492808 | 0.462681 |
| Mapk8 | 2.161395 | -0.462618 |
| Tbc1d30 | 1.917039 | 0.462021 |
| V1ra8 | 2.37194 | -0.461497 |
| Nrp1 | 2.574005 | -0.46113 |
| Cst7 | 2.518129 | 0.460984 |
| Ybx2 | 2.068364 | -0.459463 |
| Glipr2 | 2.296759 | 0.458749 |
| Fut8 | 2.027016 | 0.457888 |
| Gjb4 | 2.246834 | 0.457752 |
| Rars2 | 1.930983 | 0.456995 |
| Prss32 | 2.620107 | 0.456817 |
| Atp13a2 | 2.260566 | 0.456817 |
| Speer4c | 2.196912 | -0.456522 |
| Zfp397 | 2.044576 | -0.456259 |

| | | |
|---|---|---|
| Vmn2r91 | 2.12325 | -0.456217 |
| C1qtnf7 | 2.229686 | 0.456165 |
| Mageb3 | 3.616161 | -0.455997 |
| Olfr1368 | 2.117288 | -0.455355 |
| Tctex1d1 | 3.167174 | -0.455018 |
| Rbm25 | 2.958485 | -0.454597 |
| Slc4a2 | 2.470239 | -0.454429 |
| Klre1 | 2.47505 | -0.453902 |
| Pgls | 2.106751 | 0.45269 |
| Spata33 | 1.92619 | -0.451604 |
| Cdh12 | 2.053116 | -0.451572 |
| Gm17324 | 2.09972 | 0.45133 |
| Cilp2 | 2.419443 | -0.450623 |
| Gm21866 | 2.433339 | -0.450412 |
| Sptbn4 | 2.023734 | 0.45039 |
| D430019H16Rik | 2.195246 | 0.450348 |
| Oscar | 2.680702 | 0.450084 |
| Lrp6 | 2.259069 | -0.449662 |
| Olfr726 | 1.949949 | -0.449609 |
| Diablo | 2.566821 | -0.449588 |
| Tas2r102 | 2.927883 | 0.449408 |
| Ormdl1 | 2.867836 | 0.449017 |
| Mlh1 | 2.085631 | -0.448922 |
| Fbxl13 | 2.814328 | -0.448869 |
| Med23 | 2.087659 | -0.448859 |
| Mylk2 | 2.091865 | 0.448256 |
| A430035B10Rik | 2.001483 | 0.448182 |
| Klhl18 | 2.742826 | 0.44814 |
| Lactbl1 | 3.43326 | 0.448002 |
| Gm17305 | 2.667436 | -0.447505 |
| Try10 | 2.884253 | -0.447029 |
| Drosha | 2.999388 | -0.446362 |
| Grem2 | 2.011768 | 0.446097 |
| Kcmf1 | 2.08249 | 0.445907 |
| Nek3 | 2.175822 | -0.445547 |
| Pomk | 2.340747 | 0.445271 |
| Mybpc2 | 2.057602 | 0.445102 |
| Pde4a | 3.426501 | -0.444137 |
| Btbd8 | 2.05719 | -0.443893 |
| B4galt3 | 2.014349 | -0.443861 |
| Eif3f | 2.235011 | 0.443076 |
| Zfp503 | 2.727806 | 0.442535 |
| Ube2o | 1.964546 | 0.442238 |

| | | |
|---|---|---|
| 4933427E11Rik | 4.529343 | 0.442004 |
| Siah1b | 2.135418 | 0.441653 |
| Sec24a | 2.106532 | -0.44109 |
| Exoc3l | 2.11598 | -0.440697 |
| Idh2 | 2.013325 | -0.439347 |
| Atm | 2.162737 | -0.439208 |
| Zfp955a | 2.525438 | -0.439091 |
| Isoc1 | 2.562809 | -0.439017 |
| Atat1 | 2.13604 | -0.438921 |
| Fkbp9 | 2.421368 | -0.438634 |
| Tsks | 2.246117 | 0.43858 |
| Gsdmcl1 | 2.165667 | 0.438282 |
| Laptm4b | 3.173894 | -0.43825 |
| Gm3147 | 1.9831 | -0.438048 |
| Sycp1 | 3.666516 | -0.43791 |
| Smok2a | 1.957811 | -0.437675 |
| Gm7980 | 2.344261 | -0.437174 |
| Nxnl1 | 2.786793 | 0.436801 |
| Gfod1 | 1.973087 | -0.436652 |
| Wdr48 | 1.973214 | -0.436439 |
| Sh3bp1 | 2.383448 | 0.436375 |
| Prkcd | 2.688142 | 0.435426 |
| Xkr9 | 2.423081 | -0.435234 |
| Fut10 | 2.148634 | -0.435127 |
| Anks4b | 3.371744 | 0.434145 |
| Zfp87 | 1.923713 | 0.434092 |
| Depdc5 | 1.953626 | -0.433867 |
| Btbd16 | 2.971494 | 0.433312 |
| Gm7225 | 2.007374 | 0.433269 |
| Ccdc160 | 2.099119 | -0.433098 |
| E130309D02Rik | 2.329601 | 0.433013 |
| 4930453H23Rik | 2.221351 | 0.432778 |
| Slc30a8 | 2.79263 | -0.432682 |
| Olfr652 | 2.083293 | -0.432649 |
| Defb43 | 2.02772 | 0.432607 |
| Tulp3 | 2.5159 | 0.432543 |
| Rab3ip | 1.961328 | 0.43219 |
| Samhd1 | 2.107487 | 0.43219 |
| Cyp2c39 | 2.60841 | -0.431163 |
| Jakmip1 | 1.922745 | 0.431035 |
| Tmem240 | 1.919908 | 0.430596 |
| Disp1 | 2.176795 | -0.430532 |
| Qtrtd1 | 2.337776 | -0.430532 |

| | | |
|---|---|---|
| Sdhaf2 | 3.476813 | 0.430521 |
| Cpeb4 | 2.374043 | -0.430082 |
| Gm9955 | 2.924643 | 0.429482 |
| Vmn1r55 | 2.207119 | -0.429343 |
| Iqcc | 3.049218 | -0.429236 |
| Cd2ap | 2.394155 | -0.428389 |
| Mccc2 | 1.910996 | -0.428207 |
| Ccdc36 | 3.079991 | -0.427274 |
| Smc6 | 2.076818 | 0.426501 |
| Yeats4 | 2.012723 | 0.426029 |
| Irf2bp1 | 2.005986 | 0.42518 |
| Spdye4b | 2.324708 | 0.425158 |
| Rab11fip5 | 2.282016 | 0.425116 |
| Exoc1 | 2.081483 | 0.424879 |
| Olfr801 | 2.081402 | -0.424632 |
| Rnase6 | 2.74494 | 0.424374 |
| Qsox2 | 2.327876 | -0.424309 |
| Zbtb1 | 2.369529 | 0.423664 |
| Rps27rt | 2.149201 | 0.42149 |
| Phf13 | 2.125487 | 0.421296 |
| Olfr1195 | 1.954372 | -0.421102 |
| Psmd4 | 2.620633 | 0.420736 |
| Mphosph8 | 2.634533 | -0.420714 |
| Scgb2b7 | 2.462835 | -0.420013 |
| Wscd2 | 1.949365 | 0.419528 |
| Angptl7 | 1.925549 | 0.419463 |
| Carm1 | 2.168093 | -0.418805 |
| Ilf3 | 3.639958 | -0.418622 |
| Rad54l2 | 1.993992 | -0.41833 |
| Tulp4 | 2.37184 | -0.418017 |
| Nsf | 1.970986 | 0.417693 |
| Olfr934 | 2.072373 | -0.417131 |
| Ascc2 | 2.584574 | -0.416894 |
| Tnk1 | 2.035046 | -0.41631 |
| Stat5b | 2.800308 | -0.415943 |
| Speer6-ps1 | 2.062907 | -0.415769 |
| Cacng4 | 2.926659 | 0.415726 |
| 2700049A03Rik | 2.253829 | -0.415694 |
| Olfr486 | 2.254769 | -0.415477 |
| Prtn3 | 2.195412 | 0.415369 |
| Vps13d | 2.092524 | -0.415001 |
| Rfxank | 1.977444 | -0.414893 |
| Gm14920 | 2.502903 | 0.414753 |

| | | |
|---|---|---|
| Tpcn1 | 2.290658 | -0.414352 |
| Psmf1 | 1.936768 | 0.413681 |
| Klk10 | 2.210924 | 0.413215 |
| Lpar3 | 3.378955 | 0.412706 |
| Kctd4 | 2.203078 | 0.412597 |
| Ino80c | 2.27487 | 0.412359 |
| Strbp | 2.366563 | -0.412359 |
| Ints6 | 2.686768 | -0.412142 |
| Mtmr4 | 2.492059 | 0.412034 |
| Slc31a2 | 2.091054 | 0.411979 |
| Fam58b | 2.814271 | 0.411914 |
| Gm16367 | 2.106852 | -0.411405 |
| Gm16367 | 2.106852 | -0.411405 |
| Ago1 | 2.694864 | -0.411405 |
| Gm17584 | 2.272343 | -0.411057 |
| Lzic | 2.084263 | 0.411047 |
| Uqcc1 | 2.940471 | -0.410591 |
| Ankrd27 | 1.996863 | -0.410515 |
| Hnrnpul2 | 2.436969 | -0.410157 |
| Olfr1217 | 2.627828 | -0.410124 |
| Ffar3 | 2.232346 | -0.408527 |
| Cd4 | 2.101478 | -0.408408 |
| Uso1 | 2.677353 | 0.408234 |
| Micall1 | 2.675006 | -0.407679 |
| Ovol3 | 1.955178 | 0.40757 |
| Deaf1 | 3.250167 | -0.406635 |
| Olfr747 | 2.462134 | -0.406591 |
| Gorasp2 | 2.263685 | 0.405971 |
| Mterf3 | 2.256737 | -0.405818 |
| Mob1b | 2.116582 | 0.405143 |
| Slc25a30 | 2.126866 | 0.405121 |
| Lpar6 | 2.582773 | -0.404663 |
| Dap | 2.281378 | 0.404565 |
| F630003A18Rik | 1.990779 | 0.404533 |
| Sptlc1 | 2.032859 | 0.404129 |
| Dcp2 | 2.014684 | -0.4039 |
| Olfr1453 | 2.450123 | -0.403878 |
| Slc1a4 | 2.20275 | 0.403824 |
| BC035947 | 2.282782 | 0.403464 |
| Cypt15 | 2.506778 | 0.403191 |
| Papolg | 2.644489 | -0.403137 |
| 4933427D14Rik | 2.014134 | -0.40258 |
| H2-M3 | 2.110254 | 0.40234 |

| | | |
|---|---|---|
| Irak3 | 1.930443 | 0.402242 |
| Sis | 2.203288 | -0.40162 |
| Gm17768 | 2.072899 | -0.400767 |
| Mos | 2.419066 | 0.400757 |
| Lct | 2.152381 | 0.400068 |
| Ankk1 | 1.940891 | 0.399816 |
| Srr | 2.514854 | -0.398777 |
| Olfr576 | 1.955758 | -0.398701 |
| Nek9 | 2.616784 | -0.398318 |
| Ccdc71 | 3.4736 | 0.397474 |
| Ubac2 | 2.694591 | 0.397463 |
| Prdx5 | 2.345324 | 0.396981 |
| Cyp2c69 | 2.149831 | -0.396006 |
| Psg16 | 2.413156 | 0.395524 |
| Sco2 | 2.612197 | 0.395414 |
| Gm7247 | 2.183913 | -0.39537 |
| Tcl1b4 | 1.936607 | 0.395085 |
| Fcamr | 2.154078 | -0.395063 |
| Edem3 | 2.292385 | -0.395052 |
| 1190007I07Rik | 2.519908 | -0.39469 |
| Tdh | 2.528902 | -0.392713 |
| 1600002K03Rik | 1.957114 | 0.392427 |
| Sox21 | 2.111351 | -0.392065 |
| Cxcl1 | 2.412089 | -0.391889 |
| Zfp507 | 2.105559 | -0.391823 |
| Ptpn11 | 2.087639 | 0.391196 |
| Gm9994 | 2.432974 | -0.391141 |
| Ap5b1 | 1.999072 | 0.390866 |
| AI837181 | 3.360191 | 0.390745 |
| Pfn2 | 1.929356 | 0.39036 |
| Phox2a | 1.998535 | 0.389963 |
| Mospd3 | 2.541231 | 0.389776 |
| Gpalpp1 | 1.973128 | 0.389347 |
| Spryd7 | 3.658243 | 0.388928 |
| Zfp141 | 1.981029 | -0.388906 |
| Trappc5 | 2.204424 | 0.388708 |
| Syt12 | 2.361282 | 0.388663 |
| Hcn3 | 2.128528 | 0.388267 |
| Krt31 | 2.096828 | 0.388068 |
| Cggbp1 | 2.457867 | 0.38755 |
| Rab37 | 2.076928 | -0.387462 |
| Dnmbp | 1.937828 | -0.387385 |
| Prm3 | 2.288701 | 0.386292 |

| | | |
|---|---|---|
| Srsf2 | 4.643743 | -0.386105 |
| Slc35e2 | 2.081531 | -0.385553 |
| Lrriq4 | 1.92041 | -0.38521 |
| Il4 | 2.267819 | 0.384857 |
| Glyat | 2.150147 | -0.384536 |
| Camkmt | 2.718571 | -0.383829 |
| Afap1l2 | 2.612366 | -0.383541 |
| Fam71e2 | 2.721651 | 0.383198 |
| Pin1rt1 | 2.027262 | 0.382601 |
| Vmn1r37 | 2.340463 | 0.382512 |
| Olfr1499 | 2.247872 | -0.382435 |
| Gm11709 | 2.148689 | 0.382413 |
| Ttc13 | 2.268099 | 0.381826 |
| Olfr93 | 2.164118 | -0.381228 |
| Chst3 | 2.746241 | 0.380718 |
| Ltk | 1.985181 | 0.380596 |
| Ccdc42 | 1.996626 | 0.38012 |
| Prkg2 | 2.236765 | -0.380098 |
| Smcp | 2.086741 | -0.379466 |
| Tdrd1 | 2.877866 | -0.379288 |
| 4933421I07Rik | 2.106062 | 0.378811 |
| Rabep1 | 2.561374 | -0.378767 |
| Taf7 | 2.229088 | -0.378501 |
| Lyar | 2.609362 | -0.378301 |
| Rpp25l | 2.864813 | 0.377879 |
| Diexf | 3.825132 | -0.377835 |
| Speer4f2 | 1.984179 | 0.377202 |
| Prdx6b | 2.429388 | -0.377013 |
| Pdp1 | 2.103197 | 0.376546 |
| Arl8a | 2.018937 | 0.375735 |
| Amigo1 | 3.731738 | -0.375612 |
| Iqcf5 | 2.074907 | -0.375212 |
| Snai1 | 2.266278 | 0.375156 |
| Jmjd4 | 2.256799 | -0.374444 |
| Actn3 | 2.875884 | 0.374366 |
| Olfr1318 | 2.138325 | -0.373665 |
| Adam15 | 2.07568 | 0.373387 |
| Tktl2 | 2.161955 | 0.373231 |
| Sprr3 | 1.950949 | 0.373164 |
| 1700017N19Rik | 2.326841 | -0.372384 |
| Gstt1 | 2.052744 | -0.372061 |
| S1pr4 | 2.008258 | 0.371358 |
| Arhgap44 | 2.568823 | 0.371135 |

| | | |
|---|---|---|
| Sry | 2.051345 | -0.371035 |
| Ldlrad2 | 2.038009 | 0.370399 |
| Mier1 | 2.443724 | 0.369863 |
| Ncln | 2.101092 | 0.369651 |
| Thap4 | 1.987222 | 0.369617 |
| Atg4d | 2.12301 | 0.369517 |
| Cnga1 | 1.930328 | 0.36717 |
| Lpin2 | 3.311672 | 0.36708 |
| Med13 | 2.138929 | -0.36708 |
| Cmtm8 | 2.107526 | 0.366588 |
| Pip4k2a | 2.34955 | 0.366566 |
| Gjb6 | 1.952585 | 0.365894 |
| Serpinb12 | 2.076044 | 0.365726 |
| AW011738 | 2.069781 | -0.365525 |
| Casd1 | 2.517856 | -0.364931 |
| Sapcd2 | 2.076447 | -0.364864 |
| Rbm22 | 2.640201 | 0.364808 |
| Kcng1 | 1.926934 | 0.364135 |
| Trp53i13 | 2.229133 | -0.363967 |
| Prph | 2.316352 | 0.362868 |
| Unc5c | 2.254689 | 0.362083 |
| Cd109 | 2.284414 | 0.361959 |
| Wfdc5 | 2.017365 | 0.360713 |
| A430089I19Rik | 2.29302 | -0.36069 |
| Olfr834 | 2.470576 | -0.3606 |
| Map1s | 2.418373 | 0.360465 |
| Rnf181 | 2.591272 | 0.360432 |
| Vamp3 | 2.677663 | 0.360038 |
| Etv5 | 2.392223 | 0.3596 |
| Pcsk1n | 2.138111 | 0.35951 |
| Vps37c | 1.978993 | 0.358126 |
| Zfp263 | 2.139493 | -0.358115 |
| Fkbp3 | 2.156262 | -0.357608 |
| Micalcl | 2.733728 | -0.357608 |
| Mbd6 | 2.374387 | -0.357417 |
| Grm8 | 2.174671 | 0.357124 |
| Zfp112 | 2.41763 | -0.356696 |
| Ang6 | 2.101273 | -0.356245 |
| Tango6 | 2.386708 | -0.356076 |
| Urod | 2.203103 | 0.355366 |
| Rnf40 | 2.103751 | -0.355343 |
| Nudt13 | 2.16359 | -0.354768 |
| 1700008J07Rik | 1.976069 | -0.354599 |

| | | |
|---|---|---|
| Narfl | 1.92054 | 0.354509 |
| Pqbp1 | 2.465537 | 0.35417 |
| 4930567H17Rik | 2.470943 | 0.353877 |
| Esrrg | 1.963954 | -0.353854 |
| Mfap2 | 2.886377 | 0.353843 |
| Pank4 | 2.328027 | -0.353007 |
| Gm839 | 2.130235 | -0.352657 |
| Gdf9 | 2.460225 | 0.352148 |
| Cenpv | 2.141738 | -0.351199 |
| Rnf38 | 2.188813 | -0.351097 |
| Raver2 | 2.211822 | -0.350475 |
| Olfr687 | 1.982036 | -0.350395 |
| Olfr1289 | 1.909904 | -0.349886 |
| D17Wsu92e | 2.341039 | 0.349739 |
| Cyp4f41-ps | 1.93345 | 0.349229 |
| Ifngr1 | 2.007449 | 0.348878 |
| Sox15 | 1.978852 | -0.348697 |
| Xpnpep3 | 2.188444 | -0.347405 |
| Zzef1 | 2.396617 | -0.34711 |
| Scn5a | 2.668514 | -0.346384 |
| Enpp7 | 2.081452 | 0.346236 |
| Upf1 | 2.381353 | -0.346021 |
| Slc23a3 | 2.152202 | -0.345635 |
| Pkmyt1 | 1.938679 | -0.344988 |
| Commd5 | 2.123656 | 0.344658 |
| Serpinb3d | 2.679335 | -0.344397 |
| Ovca2 | 2.535098 | 0.344385 |
| Pdia5 | 2.334754 | 0.344272 |
| Smarcd3 | 2.273921 | -0.34409 |
| 2010106E10Rik | 2.668542 | -0.343988 |
| Tspan10 | 2.367394 | -0.343351 |
| Olfr466 | 2.326894 | -0.342953 |
| Lmf1 | 2.272844 | 0.342646 |
| Gm16513 | 1.933503 | -0.342612 |
| Gm16513 | 1.933503 | -0.342612 |
| Caln1 | 2.134214 | 0.342521 |
| Gm15217 | 2.192536 | -0.341268 |
| Taf1 | 2.302521 | -0.339833 |
| Rptn | 2.410435 | 0.339514 |
| Cphx3 | 2.083226 | -0.339411 |
| Zfp951 | 2.603311 | -0.338875 |
| Traf4 | 2.535303 | -0.338202 |
| Fam195b | 3.605958 | 0.337734 |

| | | |
|---|---|---|
| Lysmd4 | 2.297985 | 0.337528 |
| Whsc1l1 | 3.544882 | -0.336957 |
| Pah | 2.125526 | 0.336923 |
| Rnf115 | 3.130522 | 0.336729 |
| Ing3 | 2.192672 | -0.335975 |
| Pik3cb | 2.245186 | -0.335918 |
| Gm8842 | 2.286347 | 0.335861 |
| Gmpr2 | 1.948855 | 0.335518 |
| Gm16513 | 2.175198 | -0.335129 |
| Rnf187 | 4.260761 | -0.334625 |
| Gm4302 | 1.948242 | 0.334088 |
| Mat2b | 2.047607 | 0.333824 |
| Gm8817 | 2.168029 | 0.333813 |
| Oxsm | 1.98691 | 0.333721 |
| Cirh1a | 3.561192 | -0.333492 |
| Wbscr16 | 3.259084 | -0.333355 |
| Pttg1 | 1.93931 | -0.333103 |
| D030025P21Rik | 2.116744 | 0.332278 |
| Gm14408 | 1.946065 | -0.332267 |
| Fzr1 | 2.757777 | 0.331877 |
| Senp1 | 2.295654 | -0.331258 |
| Ficd | 2.820721 | 0.331029 |
| Atl2 | 1.934685 | -0.330444 |
| Olfr1335 | 2.013239 | 0.330317 |
| Heatr3 | 2.036817 | -0.329101 |
| Pgrmc2 | 2.312136 | -0.328101 |
| Olfr1408 | 1.935628 | -0.327595 |
| Plekho2 | 2.10256 | 0.327377 |
| Rhoq | 2.101652 | 0.327262 |
| Cox7b2 | 2.179409 | -0.327078 |
| Asb6 | 2.547994 | 0.327066 |
| Vmn2r106 | 2.321243 | -0.32702 |
| Thoc3 | 1.937602 | 0.326928 |
| Olfr805 | 2.296986 | -0.326744 |
| Sppl2a | 1.928726 | 0.326721 |
| Stat3 | 2.629208 | 0.325835 |
| Pdss2 | 2.465346 | -0.325467 |
| Icmt | 2.626231 | -0.325099 |
| Ptpn23 | 2.111494 | 0.324845 |
| Tsr2 | 3.396409 | -0.324384 |
| Gm10382 | 2.042564 | 0.32382 |
| Atp13a3 | 2.647902 | 0.323347 |
| Bnip3l | 2.748476 | 0.32202 |

| | | |
|---|---|---|
| Lypd5 | 2.053068 | -0.321155 |
| Hemk1 | 2.038267 | -0.321132 |
| Spats1 | 2.127735 | -0.320716 |
| Olfr686 | 2.002362 | -0.32045 |
| Gm11360 | 1.989081 | -0.319791 |
| Cpt1a | 3.207385 | -0.319525 |
| Ccndbp1 | 4.081035 | 0.319421 |
| Znrf4 | 1.971697 | 0.319329 |
| Rab35 | 2.447087 | 0.318137 |
| Sgcz | 1.980306 | -0.317779 |
| Fam129b | 1.914338 | 0.31625 |
| Olfr1221 | 2.911708 | -0.314337 |
| Msx1 | 2.068322 | 0.313524 |
| Unc79 | 2.998751 | 0.313455 |
| Olfr126 | 2.389934 | -0.312921 |
| Igip | 2.008166 | -0.312584 |
| Ggt5 | 2.006136 | 0.31105 |
| Tldc1 | 2.006638 | 0.310957 |
| Dcaf6 | 2.279985 | -0.310643 |
| Slc39a7 | 2.085745 | 0.31055 |
| Kin | 2.023867 | 0.309584 |
| Ap5s1 | 2.267365 | 0.309397 |
| Mc3r | 2.236276 | 0.309362 |
| Mtch2 | 2.227557 | 0.309048 |
| Bmyc | 2.119626 | 0.308943 |
| 4933434E20Rik | 2.222841 | -0.308093 |
| Stxbp5l | 2.077872 | -0.307743 |
| Gm13213 | 2.117083 | 0.306741 |
| C330006A16Rik | 2.115992 | -0.306029 |
| Kifap3 | 2.066192 | 0.305971 |
| Serinc2 | 2.099577 | 0.305679 |
| Dazap2 | 2.507311 | 0.305352 |
| Siglec15 | 2.020959 | 0.30513 |
| Pkig | 2.052968 | 0.304978 |
| Vmn1r177 | 2.13918 | -0.303845 |
| Tbpl1 | 1.947245 | 0.303459 |
| Anapc2 | 6.26143 | 0.303377 |
| Hormad1 | 1.971652 | -0.303214 |
| Bud13 | 2.04249 | 0.302699 |
| Sde2 | 2.476947 | -0.301553 |
| Lypd3 | 2.091412 | 0.301318 |
| Spred1 | 2.198565 | 0.300135 |
| Pdlim2 | 2.861435 | 0.298177 |

| | | |
|---|---|---|
| Dlx3 | 2.022257 | 0.298013 |
| Rab5b | 2.883054 | 0.297708 |
| Cyp20a1 | 2.026947 | -0.29678 |
| Pdrg1 | 1.965925 | -0.29678 |
| Gm5927 | 1.956677 | -0.296264 |
| Gins4 | 2.006299 | 0.295182 |
| Gnas | 2.064004 | -0.294794 |
| Iqcj | 1.935153 | -0.294759 |
| Creb1 | 2.294858 | -0.294618 |
| Sim2 | 2.228463 | 0.2945 |
| Ube4b | 2.585398 | -0.292582 |
| Kti12 | 1.919536 | -0.292205 |
| Nedd4l | 4.037186 | -0.291132 |
| Map2k2 | 2.320333 | 0.289587 |
| Fam120a | 2.90977 | -0.288583 |
| Usp4 | 2.234493 | 0.288252 |
| Dtd1 | 1.922047 | -0.287271 |
| Evpl | 2.463776 | 0.285852 |
| Natd1 | 2.20891 | 0.284846 |
| Scnm1 | 1.963695 | -0.284052 |
| Pgap2 | 2.169261 | 0.28289 |
| Gnl2 | 2.313586 | 0.282772 |
| Vmn1r53 | 2.187363 | 0.282653 |
| Oraov1 | 2.143235 | 0.282618 |
| Vmn1r213 | 1.914025 | -0.282535 |
| Plin3 | 2.533164 | 0.281645 |
| Fam175b | 2.561034 | 0.280707 |
| Dctn4 | 2.57902 | 0.279448 |
| Gm17374 | 2.095632 | -0.278675 |
| Smo | 2.499783 | -0.278532 |
| Nup98 | 2.336312 | -0.27802 |
| Zdhhc2 | 2.333631 | 0.277937 |
| Ech1 | 2.078613 | -0.277009 |
| Snap23 | 2.182545 | 0.276211 |
| Agtr1b | 2.681411 | 0.274733 |
| Mrgprd | 2.21016 | 0.271713 |
| Gorasp1 | 2.020871 | -0.271485 |
| Fam89a | 2.075536 | 0.270876 |
| Khdrbs2 | 2.274823 | 0.270601 |
| Tmem209 | 2.456997 | -0.269189 |
| Hapln4 | 2.071867 | 0.268866 |
| Lemd3 | 1.924617 | -0.268854 |
| Mdk | 1.935359 | -0.26859 |

| | | |
|---|---|---|
| 4930596D02Rik | 1.938604 | -0.268111 |
| Mta2 | 1.970697 | -0.267907 |
| Efcab8 | 2.289549 | 0.266625 |
| Ilk | 2.666321 | -0.266541 |
| Mageb18 | 2.074565 | -0.266421 |
| Aurkaip1 | 3.746744 | 0.266169 |
| Fbxo8 | 1.995081 | 0.264212 |
| Trim71 | 2.030115 | 0.263107 |
| Ccdc50 | 2.857323 | -0.26277 |
| Capn2 | 2.603476 | -0.261759 |
| Cmtr1 | 2.247753 | 0.261627 |
| Nono | 4.352428 | -0.260411 |
| Psmd1 | 2.215284 | 0.260279 |
| Mrm1 | 2.900516 | 0.258555 |
| Ubxn4 | 2.315538 | 0.257662 |
| 2310057M21Rik | 2.043568 | -0.257276 |
| Cdc123 | 2.762811 | 0.256757 |
| Cd320 | 2.039493 | 0.256564 |
| Plekha3 | 2.298386 | 0.254594 |
| Psmc2 | 2.239372 | 0.253203 |
| Pnpt1 | 2.591045 | -0.253045 |
| Ate1 | 2.416825 | -0.252416 |
| Arhgef19 | 1.98829 | -0.250671 |
| Sgsm1 | 1.928917 | -0.24913 |
| Ier5 | 2.166297 | 0.24896 |
| Rpl6 | 3.154615 | 0.248778 |
| Tecr | 2.282578 | -0.248753 |
| Rabggtb | 2.066299 | -0.248207 |
| Nrbp1 | 2.075573 | 0.247563 |
| Lrrc57 | 2.022395 | 0.243779 |
| Phb2 | 1.941403 | 0.242828 |
| Eps15 | 1.933637 | 0.242548 |
| Wdr70 | 2.340354 | -0.241584 |
| Rpl8 | 2.651286 | 0.240925 |
| Ints1 | 2.452836 | -0.240693 |
| Nadk | 2.07635 | 0.240192 |
| Ube2j1 | 2.16837 | 0.238885 |
| Acad8 | 1.92092 | -0.238383 |
| Ccar2 | 2.430391 | -0.237405 |
| Zfp148 | 2.12908 | -0.23694 |
| Rps27a | 2.388997 | 0.232833 |
| Larp4b | 2.799018 | -0.230928 |
| Ap3b1 | 2.771217 | 0.230314 |

| | | |
|---|---|---|
| Osbpl9 | 2.286531 | 0.227914 |
| Gm6934 | 2.10963 | 0.22763 |
| Hsp90ab1 | 1.914692 | -0.226607 |
| Gm5640 | 2.132123 | -0.226521 |
| Rbbp9 | 2.199352 | -0.224312 |
| Fam50a | 1.994369 | 0.223385 |
| Tial1 | 2.812073 | -0.221221 |
| Spop | 2.231666 | 0.216523 |
| Shisa5 | 2.342974 | 0.208692 |
| Kat5 | 2.395281 | 0.205768 |
| Hapln3 | 2.211101 | -0.203226 |
| Sfr1 | 2.118128 | 0.202136 |
| Ythdf3 | 1.918185 | 0.202023 |
| Eif3c | 1.98338 | 0.200404 |
| Ywhaz | 2.68638 | 0.199336 |
| Man2c1 | 2.001044 | -0.196834 |
| Rpl19 | 1.982654 | 0.193797 |
| Nucb1 | 2.706671 | 0.191866 |
| Gna12 | 2.044685 | 0.191247 |
| Cdc42se2 | 2.492067 | 0.190703 |
| Ssh1 | 2.174672 | -0.188515 |
| Exosc10 | 1.949864 | -0.18816 |
| Arl5a | 2.765414 | -0.1874 |
| Champ1 | 2.057207 | 0.187223 |
| Saraf | 2.770149 | 0.180428 |
| Glg1 | 2.218449 | -0.163099 |
| Srp54b | 2.020267 | -0.131023 |
| Eef1a1 | 2.29031 | 0.11462 |
| Gm1821 | 1.966359 | -0.093452 |

**Supplementary Table 3.**

Differentially expressed genes between CD68-/LYVE1+ cardiac lymphatic endothelial cells (LECs) isolated from the hearts of *Ackr3*$^{fl/fl}$ and *Ackr3*$^{\Delta Lyve1}$ male mice 7 days post left anterior descending artery (LAD) ligation. N=3 samples per group. One-way ANOVA. -log10 p-values and log2 fold change values are shown. Only genes that are differentially expressed in *Ackr3*$^{\Delta Lyve1}$ LECs upon LAD ligation are shown. Positive fold change values represent upregulation in LAD ligated *Ackr3*$^{\Delta Lyve1}$ LECs.

| Gene Symbol | -log10 p-value | log2 fold change |
|---|---|---|
| Mmp3 | 2.891279 | 1.619009 |
| Chil4 | 1.108801 | 1.227279 |
| Apod | 1.651444 | 1.102826 |
| Retnla | 0.808569 | -1.085418 |
| Prg4 | 3.199079 | 0.980091 |
| Tmem252 | 1.277554 | 0.964162 |
| Arhgef37 | 1.928203 | 0.951633 |
| Zbtb16 | 0.886936 | 0.881563 |
| Car6 | 0.774572 | 0.788894 |
| Gjc3 | 1.308014 | -0.781553 |
| Slpi | 0.934312 | 0.752774 |
| 6030408B16Rik | 1.479435 | 0.746029 |
| Bnip3 | 2.144117 | 0.736934 |
| Mcpt4 | 0.763742 | -0.726552 |
| Hal | 0.538647 | -0.721408 |
| Doc2b | 1.441417 | 0.701629 |
| Amy2a4 | 1.062353 | -0.672489 |
| Itgb4 | 1.921685 | -0.669499 |
| Fibin | 2.226231 | 0.667829 |
| Amy2a3 | 1.254947 | -0.65796 |
| Pcolce2 | 1.74153 | 0.657393 |
| Gpr160 | 1.734465 | 0.647066 |
| Ms4a8a | 1.123634 | 0.639779 |
| Dhdh | 2.051678 | -0.627784 |
| Angpt1 | 0.809977 | -0.625551 |
| Btbd16 | 4.458185 | 0.616706 |
| Rab37 | 3.680267 | -0.614832 |
| Cspg4 | 2.267246 | -0.610237 |
| Armcx6 | 3.160106 | -0.603777 |
| Ecm1 | 1.842827 | 0.598232 |
| Zdhhc23 | 1.396107 | 0.594921 |
| Eya2 | 0.952207 | 0.590396 |
| Tarm1 | 1.174147 | 0.586097 |
| Spata1 | 1.705289 | 0.584106 |

| | | |
|---|---|---|
| Snca | 2.088258 | 0.575148 |
| Hhipl1 | 0.878256 | 0.574654 |
| Hepacam2 | 1.041945 | -0.571754 |
| F13a1 | 1.080939 | 0.571745 |
| Medag | 2.769615 | 0.563763 |
| Irs1 | 1.300146 | -0.554913 |
| F7 | 0.773173 | 0.55451 |
| Arhgap6 | 1.105757 | -0.541604 |
| Fads2 | 1.280115 | -0.529551 |
| Nkx2-1 | 2.161265 | 0.526459 |
| Paqr9 | 0.783655 | -0.525317 |
| Sphk1 | 1.426047 | 0.524164 |
| Emilin3 | 2.177882 | 0.521835 |
| Rif1 | 2.577159 | -0.519149 |
| Reck | 0.757337 | -0.517981 |
| Rarres2 | 2.021258 | -0.511195 |
| Ttc39c | 0.421194 | 0.507536 |
| Prss35 | 1.678571 | 0.505383 |
| Mlst8 | 2.671347 | -0.504 |
| Dnah7c | 2.935512 | -0.503664 |
| Qsox1 | 1.670352 | 0.501495 |
| Tmod4 | 0.732307 | -0.501302 |
| Cluh | 2.037799 | -0.5002 |
| Cerkl | 1.970223 | -0.498608 |
| Egln3 | 1.039284 | 0.498251 |
| Htatip2 | 0.324815 | 0.494958 |
| Fbp1 | 1.163134 | 0.492592 |
| Klhl2 | 1.162766 | 0.489009 |
| Tuft1 | 1.583877 | 0.48692 |
| Gpm6b | 1.583726 | 0.48323 |
| Igfbp3 | 1.16628 | 0.47952 |
| P4ha2 | 1.269075 | 0.477408 |
| Ocm | 2.153346 | 0.47442 |
| Timm21 | 1.472114 | -0.472904 |
| Lpin3 | 1.121889 | 0.454744 |
| F10 | 1.654164 | 0.453576 |
| Baalc | 1.372769 | 0.448721 |
| Napsa | 0.60088 | 0.445716 |
| Cd33 | 0.67449 | 0.440729 |
| D130040H23Rik | 0.944828 | -0.438208 |
| Rxfp2 | 2.048142 | 0.437111 |
| Slc16a12 | 2.426057 | -0.435671 |
| Gsr | 0.818319 | 0.435597 |

| | | |
|---|---|---|
| Gdf11 | 1.067955 | -0.434145 |
| Nrxn3 | 1.950348 | 0.43204 |
| Irak3 | 2.118312 | 0.431655 |
| 4930402H24Rik | 0.781633 | -0.430649 |
| Sema5a | 1.070799 | -0.428711 |
| Fpr2 | 0.855977 | 0.427295 |
| Pstpip2 | 1.731055 | 0.426909 |
| Neto2 | 1.05745 | -0.425158 |
| Mlh1 | 1.933365 | -0.423815 |
| Agrn | 0.642362 | -0.42276 |
| Cypt15 | 2.660288 | 0.422599 |
| Bcl2 | 1.081368 | -0.421458 |
| Lpar1 | 1.358579 | 0.421178 |
| 4930567H17Rik | 3.044451 | 0.41833 |
| Plxnc1 | 0.835391 | -0.415304 |
| Ltb4r1 | 0.589128 | 0.414937 |
| Lgalsl | 1.315791 | 0.411264 |
| Olfr1278 | 0.515578 | -0.409255 |
| Gm2927 | 0.741217 | -0.407103 |
| Fam71f1 | 1.918419 | 0.406417 |
| Chic1 | 1.182634 | -0.405818 |
| Ogt | 0.916634 | -0.404554 |
| Synm | 1.476756 | -0.404543 |
| Trim37 | 1.761969 | -0.403606 |
| Lao1 | 0.985404 | 0.403388 |
| Defa22 | 1.415831 | 0.4033 |
| Cbx5 | 1.140631 | -0.402842 |
| Ero1l | 1.005639 | 0.401467 |
| Plxdc1 | 1.265499 | -0.401128 |
| Bst1 | 0.797711 | 0.399007 |
| Fgf10 | 0.879012 | -0.398613 |
| Cenpv | 2.517306 | -0.398099 |
| Trpc7 | 1.227323 | 0.397737 |
| 4933427E11Rik | 4.03227 | 0.397168 |
| Fbln2 | 1.801667 | 0.395754 |
| Irak2 | 1.40418 | 0.395545 |
| Crtac1 | 1.118239 | 0.39514 |
| Olfr1431 | 0.931751 | -0.393713 |
| Hcst | 0.538763 | 0.392823 |
| 4930503L19Rik | 0.937952 | -0.388145 |
| Ccl21a | 0.689033 | -0.387671 |
| Zfp663 | 1.832987 | 0.387263 |
| Scn3a | 1.231684 | -0.387065 |

| | | |
|---|---|---|
| Gm13242 | 0.590173 | -0.384182 |
| Tspan10 | 2.726932 | -0.38384 |
| Apba1 | 0.755018 | -0.381062 |
| Arhgap28 | 0.861252 | -0.378345 |
| Cyp2d26 | 1.951205 | 0.378079 |
| Epcam | 1.798617 | -0.377723 |
| Gm7008 | 1.275206 | 0.377357 |
| Gtf2h3 | 1.095326 | -0.377246 |
| Prss36 | 1.524804 | -0.377113 |
| Cgref1 | 1.531987 | 0.376657 |
| Tpst1 | 0.987623 | -0.373409 |
| Zfp141 | 1.874815 | -0.373041 |
| Spint2 | 1.129225 | -0.371882 |
| Lpp | 1.026037 | -0.37051 |
| Chrng | 3.236521 | 0.369595 |
| C5ar2 | 0.344188 | 0.367975 |
| Hps6 | 2.320322 | 0.366163 |
| Tulp3 | 2.010354 | 0.363765 |
| Gsdmcl1 | 1.691133 | 0.363575 |
| 0610037L13Rik | 1.499198 | -0.361768 |
| Pard3b | 1.575917 | -0.361364 |
| Ndn | 0.927463 | -0.360252 |
| Slc39a14 | 2.963199 | 0.358813 |
| Gpr141 | 0.430995 | 0.358509 |
| Pla2g1b | 1.87475 | 0.35843 |
| Gm2832 | 2.192474 | -0.358284 |
| Krt222 | 0.637702 | -0.358216 |
| Atm | 1.648468 | -0.357845 |
| Rfc1 | 2.317952 | -0.35771 |
| Mamdc2 | 1.131994 | -0.357259 |
| Ankrd29 | 1.573782 | 0.356944 |
| Trp53bp1 | 1.404196 | -0.35673 |
| Olfml1 | 1.253667 | 0.355028 |
| Gjb6 | 1.872652 | 0.354531 |
| Ube2o | 1.441343 | 0.350916 |
| Gm13304 | 0.613556 | -0.350135 |
| Ccl21c | 0.613556 | -0.350135 |
| Stxbp4 | 0.622357 | -0.349807 |
| Msl3l2 | 1.409169 | -0.349116 |
| Slitrk6 | 0.672322 | -0.348867 |
| Qsox2 | 1.79578 | -0.348221 |
| Trpv4 | 0.973144 | 0.348017 |
| Olfr1216 | 0.907483 | -0.347031 |

| | | |
|---|---|---|
| Zkscan14 | 1.192946 | 0.346872 |
| Pex11b | 1.013811 | 0.346747 |
| Sgpp2 | 0.778367 | -0.345147 |
| Zfp827 | 0.597391 | -0.343067 |
| Tbc1d15 | 0.539246 | 0.340038 |
| Gtf2i | 1.107707 | -0.339696 |
| Ice2 | 1.011131 | -0.339149 |
| Tmpo | 1.293339 | -0.338647 |
| Ptprz1 | 0.769669 | -0.338419 |
| Ovol3 | 1.515046 | 0.336798 |
| Vmn1r55 | 1.593373 | -0.335792 |
| Pfn2 | 1.57451 | 0.335266 |
| Olfr1335 | 2.049779 | 0.334866 |
| Eid2b | 0.834827 | 0.334637 |
| Tmem40 | 0.622126 | 0.334523 |
| Enpp7 | 1.989306 | 0.334511 |
| Isoc1 | 1.789813 | -0.333344 |
| Galnt11 | 1.152837 | -0.332416 |
| Srek1 | 0.880787 | -0.331304 |
| Vmn1r37 | 1.934372 | 0.330593 |
| Rab3ip | 1.362838 | 0.329445 |
| Ccrl2 | 0.780444 | 0.32902 |
| Jmjd1c | 0.99083 | -0.328871 |
| Inpp4b | 0.714443 | -0.328354 |
| Grm8 | 1.94233 | 0.327687 |
| Tcl1b4 | 1.498612 | 0.326192 |
| Pkmyt1 | 1.799016 | -0.3261 |
| Trim34b | 0.570595 | -0.324845 |
| Cyp4f40 | 1.163625 | 0.324834 |
| Gm2022 | 0.748252 | -0.324523 |
| Alms1 | 1.232148 | -0.323658 |
| Borcs6 | 0.675121 | 0.323255 |
| 1190007I07Rik | 1.943206 | -0.323093 |
| Tra2a | 0.55155 | -0.32239 |
| Oxnad1 | 1.563084 | -0.320126 |
| Ssc5d | 1.066446 | 0.319375 |
| F12 | 1.089644 | 0.31904 |
| Fgf23 | 0.643122 | 0.317512 |
| Hnrnph1 | 1.059357 | -0.317188 |
| D17Wsu92e | 2.057443 | 0.31669 |
| Gfpt1 | 1.188268 | -0.314035 |
| Tcl1b2 | 1.355958 | 0.313478 |
| Dclk2 | 1.220076 | -0.312816 |

| | | |
|---|---|---|
| Tmem182 | 0.481406 | -0.312003 |
| Zkscan4 | 1.078061 | 0.310957 |
| 4930596D02Rik | 2.349123 | -0.310596 |
| Olfr242 | 0.824491 | -0.308955 |
| Pbrm1 | 1.99163 | -0.308139 |
| Tubb3 | 0.912059 | 0.307906 |
| Nisch | 0.899343 | -0.307732 |
| Fdxacb1 | 0.692461 | 0.307685 |
| Anapc1 | 1.15731 | -0.307533 |
| B4galt3 | 1.223276 | -0.306181 |
| Chst3 | 2.072718 | 0.305702 |
| Slc3a2 | 1.441768 | 0.305469 |
| Map2k4 | 0.542515 | 0.305025 |
| Ace | 1.053704 | 0.305025 |
| Mak | 0.751054 | 0.304523 |
| Birc3 | 0.793595 | 0.303857 |
| Prox1 | 0.683862 | -0.303085 |
| Psg16 | 1.699476 | 0.302875 |
| Gjb4 | 1.292142 | 0.302418 |
| Scand1 | 1.629178 | 0.302067 |
| Gpihbp1 | 1.148085 | 0.301553 |
| Gm6377 | 0.428215 | 0.30078 |
| Zbtb21 | 1.266763 | -0.299913 |
| Lzic | 1.359373 | 0.298623 |
| H2-Ea-ps | 1.189438 | 0.297567 |
| Adck4 | 0.691363 | 0.297262 |
| D430019H16Rik | 1.25942 | 0.297215 |
| Gpr150 | 1.346035 | 0.296463 |
| Unc79 | 2.783267 | 0.295065 |
| Pah | 1.783496 | 0.295029 |
| Gbf1 | 1.169845 | -0.293312 |
| Atl1 | 0.865785 | -0.293041 |
| Aldh1a1 | 0.664664 | 0.291945 |
| Ifnz | 0.849538 | 0.291639 |
| Gm17449 | 0.855034 | -0.290401 |
| Tspan18 | 0.510914 | -0.290306 |
| Osm | 0.629002 | 0.289787 |
| Oscar | 1.503611 | 0.289622 |
| Pcdhb6 | 1.086549 | 0.28961 |
| Cpeb4 | 1.401723 | -0.289126 |
| Metap1d | 0.712821 | -0.288181 |
| Gm26616 | 1.090481 | 0.288016 |
| Gm5885 | 0.609259 | 0.287496 |

| | | |
|---|---|---|
| Nipal1 | 1.039283 | 0.286515 |
| Stap2 | 0.60541 | -0.286313 |
| Prrc2a | 1.031656 | -0.285722 |
| Wfdc5 | 1.471225 | 0.284822 |
| Olfr373 | 0.922145 | -0.284336 |
| Git1 | 0.81286 | 0.284206 |
| Clcc1 | 1.182703 | 0.283104 |
| Pik3cg | 0.550059 | -0.283056 |
| Slc6a14 | 1.303107 | 0.282926 |
| Cdkal1 | 0.881405 | -0.281977 |
| Gm1987 | 0.520382 | -0.281526 |
| Cep70 | 1.759958 | -0.280897 |
| Htra3 | 1.165232 | 0.280743 |
| Olfr1132 | 0.804676 | -0.280719 |
| Olfr203 | 0.489472 | -0.279697 |
| Camkmt | 1.793209 | -0.278484 |
| Mos | 1.490253 | 0.277794 |
| Gda | 1.582432 | 0.277259 |
| Clec2f | 0.426617 | -0.275937 |
| Rtel1 | 1.564728 | -0.275198 |
| Bicc1 | 1.193396 | 0.274924 |
| Olfr314 | 1.548128 | 0.274804 |
| Mc3r | 1.907767 | 0.274077 |
| Smco2 | 1.063863 | -0.273981 |
| Dus4l | 1.312296 | -0.273922 |
| Slfn1 | 0.440827 | 0.273408 |
| Psmd9 | 0.517755 | 0.273337 |
| Krt5 | 0.882792 | 0.273241 |
| Prtn3 | 1.253705 | 0.273194 |
| Slc2a1 | 0.513097 | 0.272859 |
| Stat3 | 2.09227 | 0.272823 |
| Afap1l2 | 1.670775 | -0.272453 |
| Gm10719 | 0.381271 | -0.27145 |
| Gm8817 | 1.645698 | 0.271127 |
| Gm21814 | 1.284857 | -0.270601 |
| Zfp503 | 1.427705 | 0.270266 |
| Acbd4 | 1.046977 | -0.269943 |
| Bri3 | 0.913252 | 0.269548 |
| Pabpc1l | 0.908003 | 0.26847 |
| Gm13298 | 0.81268 | -0.267044 |
| Mylk2 | 1.032865 | 0.265209 |
| 1700008O03Rik | 0.890158 | 0.265089 |
| Unc5c | 1.489535 | 0.264981 |

| | | |
|---|---|---|
| Fosl2 | 0.993076 | 0.264068 |
| Cmtm8 | 1.355056 | 0.263455 |
| Mkrn1 | 0.451385 | 0.262806 |
| Vmn1r129 | 0.447208 | -0.262541 |
| Zfp951 | 1.868831 | -0.262517 |
| Dck | 0.724965 | -0.261916 |
| Slc25a23 | 0.713683 | -0.26182 |
| Lace1 | 0.882397 | -0.261314 |
| Cobl | 0.459908 | -0.261302 |
| Arl8a | 1.239289 | 0.26123 |
| Plekha3 | 2.363164 | 0.260146 |
| Lrrc10b | 0.93661 | 0.259712 |
| Masp1 | 0.386616 | 0.25952 |
| Olfr294 | 0.419753 | -0.259339 |
| Igfbp5 | 0.457787 | -0.259339 |
| Olfr153 | 0.414698 | -0.25882 |
| Slc25a30 | 1.163306 | 0.258362 |
| Uqcc1 | 1.610432 | -0.258061 |
| Chmp2b | 0.839144 | 0.257566 |
| Ncs1 | 1.239419 | 0.257518 |
| Krtcap3 | 1.109714 | -0.256902 |
| Pigg | 0.592108 | -0.25683 |
| Plvap | 0.46361 | -0.256274 |
| 1700029J07Rik | 0.492772 | -0.255924 |
| Gm3376 | 0.475023 | -0.255803 |
| Foxf1 | 0.727036 | 0.255791 |
| Eif4a2 | 0.344866 | -0.255742 |
| Nav1 | 0.562297 | -0.255416 |
| Fhl1 | 0.438444 | -0.254667 |
| Ifngr1 | 1.311898 | 0.254062 |
| Bdnf | 0.5299 | 0.25359 |
| Gm16367 | 1.099297 | -0.253469 |
| Gm16367 | 1.099297 | -0.253469 |
| Ficd | 2.000786 | 0.253251 |
| Pcdhb9 | 0.503126 | 0.252779 |
| Gm6337 | 0.683198 | -0.252173 |
| Gm2046 | 0.550599 | -0.251689 |
| Cdk17 | 1.116063 | -0.251046 |
| Lins1 | 1.260092 | -0.250622 |
| Tsc22d4 | 0.922556 | 0.250125 |
| Rps27rt | 1.063272 | 0.249555 |
| Nfam1 | 0.458948 | 0.24947 |
| Gm14920 | 1.271877 | 0.249008 |

| | | |
|---|---|---|
| Trib1 | 0.868138 | 0.248559 |
| Tktl2 | 1.254442 | 0.248498 |
| Snx24 | 0.450523 | 0.248061 |
| Oxa1l | 1.033554 | 0.246676 |
| Samd5 | 0.428358 | -0.246445 |
| Cmtr1 | 2.077683 | 0.246311 |
| Pde4a | 1.614292 | -0.246238 |
| Olfr899 | 0.323684 | -0.246141 |
| Gm9955 | 1.407767 | 0.245398 |
| Fbn2 | 0.771679 | 0.245228 |
| Olfr1217 | 1.332482 | -0.245216 |
| Fjx1 | 0.309727 | 0.244802 |
| Tas2r102 | 1.312667 | 0.243645 |
| Trnp1 | 0.743607 | 0.243486 |
| Mcpt8 | 0.74229 | -0.243413 |
| Mettl13 | 0.641753 | 0.243072 |
| Cyp2c39 | 1.217307 | -0.242596 |
| Frmd6 | 1.047766 | -0.242499 |
| Baz1b | 0.844464 | -0.242096 |
| Mcoln2 | 1.221029 | 0.241791 |
| Gm19475 | 0.789663 | 0.24145 |
| Gpi1 | 1.287791 | 0.241267 |
| Sel1l | 0.536873 | 0.241047 |
| Rest | 0.897254 | 0.240961 |
| Arhgap22 | 0.328975 | -0.240864 |
| Tnk1 | 0.973679 | -0.240791 |
| Rps13 | 0.594331 | 0.240766 |
| Gm16513 | 0.718047 | -0.239752 |
| AA474331 | 0.551999 | 0.238958 |
| Lman1l | 0.939427 | 0.238714 |
| Olfr1480 | 0.388739 | 0.238457 |
| Serpini2 | 0.674653 | -0.238053 |
| C330006A16Rik | 1.512869 | -0.237882 |
| Gm10256 | 0.502272 | -0.237613 |
| Fam71e2 | 1.45367 | 0.2376 |
| Chid1 | 0.678147 | -0.236633 |
| Spata6 | 1.229499 | 0.236633 |
| Tm2d3 | 0.736463 | 0.23531 |
| LOC100503048 | 0.504619 | -0.234538 |
| Hs1bp3 | 1.699289 | -0.234281 |
| Olfr1188 | 0.628836 | 0.234268 |
| Huwe1 | 0.501025 | -0.233925 |
| Olfr1487 | 1.188249 | -0.233839 |

| | | |
|---|---|---|
| Tmem151a | 0.899395 | 0.233618 |
| Il17ra | 0.378496 | 0.233581 |
| Sval3 | 0.840773 | -0.233152 |
| Asph | 0.828718 | -0.230806 |
| Vps13d | 0.948469 | -0.230474 |
| Zfp948 | 0.565334 | -0.230191 |
| Thoc2 | 0.784743 | -0.2296 |
| Gm5801 | 0.799853 | 0.229428 |
| Mtss1 | 0.777489 | -0.229354 |
| Smyd1 | 0.308697 | -0.229182 |
| Tango6 | 1.324855 | -0.228665 |
| Zfp87 | 0.807171 | 0.227938 |
| 4933434E20Rik | 1.487094 | -0.227679 |
| Bud13 | 1.394823 | 0.227618 |
| Gm5927 | 1.371762 | -0.227347 |
| Fgf11 | 0.279055 | -0.226977 |
| Ech1 | 1.589014 | -0.226373 |
| 4931408C20Rik | 0.867484 | 0.22588 |
| Ces1c | 0.663016 | -0.225793 |
| Mmaa | 0.59998 | -0.225731 |
| Zfp672 | 0.55561 | 0.225682 |
| Psmf1 | 0.854872 | 0.225559 |
| Klf12 | 0.282821 | -0.225522 |
| Diablo | 1.022621 | -0.225287 |
| Abi3bp | 0.258454 | -0.225065 |
| Slc16a13 | 0.641768 | -0.225053 |
| Fstl3 | 0.657791 | 0.223979 |
| Inppl1 | 0.696998 | -0.223682 |
| Exoc3l | 0.847712 | -0.22325 |
| Cds1 | 0.682568 | 0.223126 |
| Exoc4 | 0.535041 | -0.222211 |
| Lactbl1 | 1.391778 | 0.222162 |
| Fsd1l | 0.618881 | -0.221469 |
| Gm10046 | 0.611295 | -0.219859 |
| Hilpda | 0.674503 | 0.219748 |
| Stambp | 0.720096 | 0.219017 |
| Rrad | 0.421516 | -0.218918 |
| Tcp10b | 0.783673 | 0.218707 |
| Zdhhc2 | 1.693444 | 0.217876 |
| Olfr1463 | 0.388108 | 0.217342 |
| LOC100861707 | 0.446615 | -0.217069 |
| Snx32 | 0.471885 | -0.215989 |
| Olfr1141 | 0.663118 | -0.215592 |

| | | |
|---|---|---|
| Hapln4 | 1.538575 | 0.215057 |
| Mrm1 | 2.296679 | 0.214945 |
| Pcsk1n | 1.069123 | 0.214572 |
| Tecr | 1.873544 | -0.213976 |
| Ilf3 | 1.551322 | -0.213938 |
| Ankrd27 | 0.829791 | -0.213913 |
| Siglec15 | 1.251986 | 0.213577 |
| Pnpt1 | 2.081806 | -0.213528 |
| Ccndbp1 | 2.520714 | 0.213366 |
| Ube2cbp | 0.708678 | 0.212694 |
| Treh | 1.238744 | 0.212544 |
| Pdk1 | 0.240345 | 0.212158 |
| Luzp2 | 0.649517 | -0.212109 |
| Agtr1b | 1.918167 | 0.212009 |
| Psd3 | 0.266127 | -0.211698 |
| Cr2 | 0.550526 | -0.211436 |
| Spryd7 | 1.685324 | 0.210489 |
| Trim71 | 1.506439 | 0.210414 |
| Fam208a | 0.482242 | -0.20979 |
| Nav3 | 0.242848 | -0.208954 |
| Hcn3 | 0.923578 | 0.20868 |
| Ppp1r12b | 0.317708 | -0.208455 |
| Tbc1d30 | 0.65608 | 0.208143 |
| Ppid | 0.334228 | -0.207855 |
| Ldlrad2 | 0.935651 | 0.20778 |
| Cd14 | 0.459254 | 0.207718 |
| Olfr502 | 0.395909 | -0.207318 |
| Irf2bp1 | 0.76223 | 0.207218 |
| Ddhd2 | 0.731675 | -0.207156 |
| Ikbkg | 0.479475 | 0.206831 |
| Apoe | 0.561608 | 0.206481 |
| Oog1 | 0.714828 | 0.206343 |
| Msx1 | 1.178726 | 0.206243 |
| BC035947 | 0.927802 | 0.206181 |
| AI413582 | 0.666323 | 0.206143 |
| Gdf9 | 1.203098 | 0.205693 |
| Depdc5 | 0.714857 | -0.20558 |
| Iqcc | 1.158381 | -0.20528 |
| Cggbp1 | 1.051801 | 0.205167 |
| Usp29 | 0.184584 | -0.20513 |
| Tctex1d1 | 1.114495 | -0.20498 |
| Olfr1196 | 0.526847 | 0.204904 |
| Tax1bp1 | 0.744431 | 0.204354 |

| | | |
|---|---|---|
| Ccdc96 | 0.971307 | 0.203389 |
| Plk3 | 0.54858 | -0.203339 |
| Olfr1331 | 0.541208 | -0.202951 |
| Gm9733 | 0.264665 | 0.202913 |
| Sphkap | 0.774058 | -0.202625 |
| Shroom3 | 0.454995 | 0.202549 |
| Med8 | 0.657877 | 0.202248 |
| Vmn1r57 | 0.543329 | -0.201897 |
| Amfr | 0.647727 | 0.20186 |
| Olfr1298 | 0.469777 | -0.201496 |
| Vwa8 | 0.777343 | -0.20137 |
| Cd300ld | 0.289503 | -0.201283 |
| Rit2 | 0.531061 | -0.201283 |
| Urad | 0.892637 | 0.201245 |
| Mier1 | 1.084635 | 0.201195 |
| Syt12 | 0.977584 | 0.201119 |
| Gm2933 | 0.298456 | -0.200906 |
| Iqgap2 | 0.338023 | -0.200718 |
| Senp1 | 1.174733 | -0.200642 |
| Fam187b | 0.471259 | 0.199914 |
| Krt31 | 0.858833 | 0.199889 |
| Crispld2 | 0.669643 | 0.199789 |
| Gorasp2 | 0.873969 | 0.199751 |
| Fkbp3 | 0.985228 | -0.199638 |
| Olfr1152 | 0.311039 | -0.199474 |
| Agtrap | 0.906197 | 0.199336 |
| E130114P18Rik | 0.90689 | -0.199085 |
| Tmem192 | 0.333927 | 0.198381 |
| Gm21738 | 0.234762 | -0.198029 |
| Lrrc4c | 0.251463 | -0.198029 |
| Disp1 | 0.766321 | -0.197928 |
| Olfr703 | 0.7499 | 0.197526 |
| Vmn1r213 | 1.180106 | -0.197287 |
| 4921511C20Rik | 0.859347 | -0.197236 |
| Rgs12 | 0.55842 | 0.196783 |
| Sim2 | 1.296385 | 0.196267 |
| Kctd4 | 0.807494 | 0.195448 |
| Rgs13 | 0.448004 | -0.195285 |
| Oxsm | 0.965163 | 0.195285 |
| Gm15217 | 1.035129 | -0.195083 |
| Nek9 | 1.009178 | -0.194667 |
| Cnr2 | 0.271498 | 0.194554 |
| Gm4302 | 0.939317 | 0.194491 |

| | | |
|---|---|---|
| Slc2a9 | 0.558354 | 0.193847 |
| Gm15850 | 0.625109 | 0.193608 |
| Tesc | 0.879459 | 0.193507 |
| Rnase6 | 0.964042 | 0.193002 |
| P2rx7 | 0.332671 | 0.192914 |
| Gm6465 | 0.532849 | -0.19251 |
| Gm9994 | 0.94252 | -0.192459 |
| Mccc2 | 0.650814 | -0.192194 |
| Gm20873 | 0.477904 | -0.192156 |
| Prm3 | 0.89512 | 0.191841 |
| Mdh1 | 0.687476 | -0.191474 |
| Olfr93 | 0.855012 | -0.191045 |
| Gm11709 | 0.84383 | 0.190817 |
| Zdhhc14 | 0.513621 | 0.189945 |
| Olfr38 | 0.608486 | -0.189806 |
| Gon4l | 0.91989 | -0.189502 |
| 1600002K03Rik | 0.734131 | 0.189451 |
| Gm20026 | 0.761024 | 0.189375 |
| D1Ertd622e | 0.668293 | 0.189059 |
| Gm11084 | 0.231014 | -0.188907 |
| Arhgap42 | 0.513613 | -0.188844 |
| Top1 | 0.52195 | -0.188692 |
| Pin1rt1 | 0.78184 | 0.188528 |
| BC017643 | 0.8174 | 0.188401 |
| Zcchc16 | 0.594812 | 0.188122 |
| Olfr687 | 0.854201 | -0.187628 |
| Olfr342 | 0.405727 | -0.187527 |
| Csgalnact2 | 0.55458 | 0.187388 |
| Pqbp1 | 1.052316 | 0.187096 |
| D16Ertd472e | 0.380107 | 0.186729 |
| Cfap20 | 0.615743 | -0.186285 |
| Pgs1 | 0.581782 | 0.18626 |
| Nudt4 | 0.867094 | -0.185943 |
| Enpp3 | 0.29546 | -0.185689 |
| Eif3f | 0.694539 | 0.185385 |
| Lpar6 | 0.909195 | -0.184978 |
| Metrn | 0.943747 | 0.184737 |
| Atg4d | 0.83527 | 0.184674 |
| Ncstn | 0.664428 | 0.184636 |
| Fam89a | 1.228866 | 0.183264 |
| Heatr3 | 0.924351 | -0.183035 |
| Slc31a2 | 0.702515 | 0.182972 |
| Klhl36 | 0.484125 | 0.182476 |

| | | |
|---|---|---|
| Parva | 0.520143 | -0.182158 |
| Eif4g3 | 0.742645 | -0.181497 |
| Hook2 | 0.750836 | 0.181395 |
| Pik3cb | 0.980813 | -0.181192 |
| Ncam1 | 0.370365 | -0.17997 |
| Srsf2 | 1.835656 | -0.179384 |
| Cd4 | 0.694765 | -0.179193 |
| Astn2 | 0.501193 | -0.178874 |
| Parp8 | 0.661932 | -0.178683 |
| Tdrd1 | 1.056529 | -0.177675 |
| Pllp | 0.510932 | 0.177663 |
| Jakmip1 | 0.585334 | 0.177624 |
| Gm3106 | 0.655559 | -0.177101 |
| Lmo4 | 0.650742 | 0.177025 |
| Celf3 | 0.622493 | 0.176948 |
| Idh2 | 0.594135 | -0.176859 |
| Ces2a | 0.445539 | -0.17668 |
| Bnip3l | 1.237932 | 0.175952 |
| 4930453H23Rik | 0.662341 | 0.175518 |
| Lrsam1 | 0.559724 | -0.175454 |
| Slc4a2 | 0.691628 | -0.175352 |
| Mdk | 1.088617 | -0.175237 |
| B830017H08Rik | 0.842018 | 0.174803 |
| Saa4 | 0.364546 | 0.174483 |
| Ptges3l | 0.428644 | -0.174189 |
| Gnas | 1.016694 | -0.174163 |
| Cox7b2 | 0.933413 | -0.173997 |
| Rab5b | 1.427469 | 0.173869 |
| Gm17305 | 0.754115 | -0.173434 |
| Vps37c | 0.744788 | 0.173307 |
| Srr | 0.825408 | -0.173243 |
| Plekhb1 | 0.466645 | -0.173166 |
| Gm15737 | 0.452558 | -0.17282 |
| Cyp20a1 | 0.97791 | -0.172731 |
| Crlf1 | 0.416487 | 0.172462 |
| Ubxn4 | 1.353825 | 0.172129 |
| Rbm22 | 0.968002 | 0.171668 |
| 6430573F11Rik | 0.580887 | -0.171591 |
| Sis | 0.702099 | 0.171284 |
| Olfr292 | 0.262574 | 0.170989 |
| Zfp112 | 0.903045 | -0.170874 |
| Man2c1 | 1.657773 | -0.170874 |
| Gm6934 | 1.438237 | 0.170822 |

| | | |
|---|---|---|
| Phox2a | 0.658995 | 0.170797 |
| Cilp2 | 0.660984 | -0.170784 |
| Tmem104 | 0.332388 | 0.170617 |
| Olfr618 | 0.471483 | -0.170489 |
| Gm766 | 0.504029 | -0.170374 |
| Fbxl13 | 0.774815 | -0.170066 |
| Ascc2 | 0.778601 | -0.169681 |
| Ube3c | 0.737853 | -0.169322 |
| Micalcl | 1.013501 | 0.169322 |
| Parn | 0.508826 | 0.169143 |
| Gm16513 | 0.865874 | -0.168873 |
| Smarca5 | 1.045519 | -0.168655 |
| Ap5b1 | 0.645051 | 0.168411 |
| Ssh1 | 1.8704 | -0.168116 |
| Cdh3 | 0.272163 | 0.168103 |
| 2700049A03Rik | 0.667958 | -0.167782 |
| Nabp2 | 0.498916 | 0.167743 |
| Rnf113a1 | 0.461044 | 0.166857 |
| Gm16429 | 0.666446 | -0.166844 |
| Gm7971 | 0.666446 | -0.166844 |
| Carm1 | 0.627969 | -0.166188 |
| Mat2b | 0.800352 | 0.166124 |
| Trem3 | 0.302263 | 0.165725 |
| Tex101 | 0.528867 | 0.165623 |
| Mvd | 0.489648 | 0.16467 |
| Olfr1221 | 1.242964 | -0.164439 |
| Hemk1 | 0.826176 | -0.16422 |
| Adam15 | 0.688276 | 0.164207 |
| Dalrd3 | 0.992303 | -0.163885 |
| Gm20877 | 0.356395 | -0.163859 |
| E130309D02Rik | 0.634076 | 0.163769 |
| Nedd4l | 1.992099 | -0.163769 |
| Cyp3a11 | 0.38079 | -0.163344 |
| Nrbp1 | 1.187733 | 0.163331 |
| Sapcd2 | 0.704253 | -0.163215 |
| Zfp616 | 0.281966 | -0.163125 |
| Gpalpp1 | 0.613653 | 0.163112 |
| Nr2f6 | 0.448112 | -0.162623 |
| Tmem207 | 0.33013 | -0.162558 |
| Ccr3 | 0.274153 | 0.162519 |
| Zzef1 | 0.867494 | -0.162494 |
| Aox1 | 0.182433 | -0.162146 |
| Dazap2 | 1.073699 | 0.161578 |

| | | |
|---|---|---|
| Gm16513 | 0.700268 | -0.161101 |
| Gm16513 | 0.700268 | -0.161101 |
| Saraf | 2.392704 | 0.160701 |
| Pclo | 0.358549 | -0.160288 |
| Kcng1 | 0.63825 | 0.16003 |
| Atat1 | 0.552699 | -0.159784 |
| Gm4787 | 0.51413 | -0.158789 |
| Olfr934 | 0.568432 | -0.158789 |
| Taf1 | 0.824508 | -0.158027 |
| Snap23 | 1.027791 | 0.157613 |
| Gm3278 | 0.368277 | -0.157354 |
| Olfr727 | 0.295301 | -0.156979 |
| Adgrl2 | 0.398235 | -0.156953 |
| Gm13247 | 0.54129 | -0.156798 |
| Gm15448 | 0.476501 | 0.156655 |
| Actn3 | 0.903813 | 0.156332 |
| Slc36a1 | 0.317177 | 0.155581 |
| Muc15 | 0.723155 | -0.155205 |
| Pank4 | 0.773689 | -0.155205 |
| Kcmf1 | 0.508175 | 0.155115 |
| Tifa | 0.26798 | 0.154804 |
| Arhgap44 | 0.797501 | 0.154531 |
| Anks4b | 0.864 | 0.153715 |
| Brwd1 | 0.38629 | -0.153637 |
| Cttnbp2 | 0.480634 | -0.15352 |
| Hs6st2 | 0.398002 | 0.153299 |
| Atf7ip | 0.388076 | -0.15291 |
| Olfr1112 | 0.214442 | -0.152832 |
| Xpnpep3 | 0.724474 | -0.152495 |
| Sirt5 | 0.425973 | -0.152352 |
| Lpin2 | 1.048692 | 0.152313 |
| Thap4 | 0.603644 | 0.152093 |
| Xkr9 | 0.594152 | -0.151924 |
| Slc39a7 | 0.79672 | 0.151872 |
| Gm10972 | 0.653363 | -0.151703 |
| Ino80c | 0.594925 | 0.151326 |
| Olfr810 | 0.67229 | -0.15104 |
| Rom1 | 0.399873 | 0.150703 |
| Hnrnpul2 | 0.638656 | -0.150599 |
| Gm2244 | 0.358391 | 0.150378 |
| Itch | 0.506287 | 0.150066 |
| Camk2a | 0.281264 | -0.149975 |
| Casp1 | 0.40205 | -0.149454 |

| | | |
|---|---|---|
| Sbno2 | 0.629298 | 0.149285 |
| Slc9a9 | 0.224723 | 0.148635 |
| Zfp263 | 0.656381 | -0.148466 |
| Palm2 | 0.346704 | -0.148101 |
| 4931414P19Rik | 0.462133 | 0.148088 |
| Sult2a7 | 0.384879 | 0.148049 |
| Olfr745 | 0.408358 | -0.148036 |
| Kin | 0.749514 | 0.147984 |
| D830013O20Rik | 0.72827 | -0.14775 |
| Kcnc2 | 0.168889 | 0.147737 |
| Cdh11 | 0.215126 | -0.147229 |
| Dock4 | 0.302929 | -0.147202 |
| Sfr1 | 1.382205 | 0.146851 |
| Grem2 | 0.457528 | 0.146786 |
| Psmd4 | 0.639605 | 0.146056 |
| Prkab2 | 0.415952 | -0.146003 |
| Celsr1 | 0.37105 | -0.146003 |
| Paqr8 | 0.409465 | -0.145638 |
| Wbscr16 | 1.102894 | -0.145495 |
| Klre1 | 0.542922 | -0.145417 |
| Pih1d3 | 0.562124 | -0.145377 |
| Cpt1a | 1.144036 | -0.145208 |
| Gorasp1 | 0.869949 | -0.145208 |
| Zfyve28 | 0.387093 | -0.145143 |
| Stc2 | 0.171068 | 0.144908 |
| Tmem209 | 1.071164 | -0.144477 |
| Ptrh1 | 0.309278 | -0.144268 |
| Krt8 | 0.207746 | 0.144138 |
| Wsb2 | 0.262488 | 0.144007 |
| Slc7a3 | 0.528384 | 0.143707 |
| Rnf181 | 0.757414 | 0.143406 |
| Nup98 | 0.962167 | -0.143354 |
| Prr15 | 0.184242 | -0.143341 |
| Dennd2d | 0.247875 | -0.143328 |
| Fam175b | 1.046014 | 0.143289 |
| Caln1 | 0.662413 | 0.143263 |
| Ang6 | 0.617342 | -0.142923 |
| Wdr48 | 0.445198 | -0.142649 |
| Gnl2 | 0.921217 | 0.142257 |
| Tmem189 | 0.262582 | 0.14206 |
| Gem | 0.280765 | -0.141551 |
| 1700008J07Rik | 0.575347 | -0.141263 |
| Lypd3 | 0.754638 | 0.141197 |

| | | |
|---|---|---|
| Lin7a | 0.447147 | -0.141171 |
| Rab11fip5 | 0.523866 | 0.141132 |
| Speer4f2 | 0.532183 | 0.14108 |
| Uso1 | 0.649543 | 0.140962 |
| Olfr1489 | 0.1406 | -0.14091 |
| AA414768 | 0.318168 | 0.140543 |
| Dtd1 | 0.733684 | -0.14053 |
| Mcfd2 | 0.390505 | 0.140425 |
| Rora | 0.30552 | -0.140321 |
| Zfp329 | 0.297052 | -0.140307 |
| AI837181 | 0.873986 | 0.139941 |
| Spop | 1.243174 | 0.139758 |
| Msc | 0.511327 | 0.139718 |
| Rttn | 0.386864 | -0.139378 |
| Was | 0.298206 | 0.139339 |
| Mfap2 | 0.836013 | 0.139024 |
| Dnajc3 | 0.371234 | 0.138539 |
| Sptbn4 | 0.42126 | 0.138251 |
| Izumo3 | 0.217605 | -0.138133 |
| Gm10097 | 0.330953 | -0.138107 |
| Gm10600 | 0.263353 | -0.13808 |
| Uhrf2 | 0.447287 | -0.13808 |
| Noc4l | 0.274809 | 0.13774 |
| Gabrb3 | 0.419403 | 0.137543 |
| Olfr930 | 0.347034 | 0.137228 |
| Gm3115 | 0.292696 | -0.137228 |
| Sry | 0.540942 | -0.137018 |
| Hes6 | 0.210678 | 0.135443 |
| Tsks | 0.46853 | 0.135141 |
| St8sia2 | 0.224443 | -0.134812 |
| Cst7 | 0.487534 | 0.134037 |
| Azin2 | 0.165906 | 0.133419 |
| Gpr84 | 0.294283 | 0.133419 |
| Ybx2 | 0.399912 | -0.133103 |
| Prss32 | 0.508692 | 0.133051 |
| Atp13a3 | 0.809677 | 0.132945 |
| Klhl23 | 0.377324 | -0.132656 |
| Gucy1a3 | 0.269107 | -0.132419 |
| Ilk | 1.05195 | -0.132327 |
| Slc19a2 | 0.466754 | 0.132024 |
| Slc1a4 | 0.495286 | 0.131906 |
| Gm8281 | 0.247622 | -0.131761 |
| Gm17584 | 0.498566 | -0.131761 |

| | | |
|---|---|---|
| Raver2 | 0.596454 | -0.131695 |
| Nono | 1.893004 | -0.131589 |
| Dcp2 | 0.451175 | -0.131313 |
| Nr4a2 | 0.25052 | -0.131194 |
| Cxcl1 | 0.559828 | 0.131155 |
| Laptm4b | 0.643156 | -0.130786 |
| Vamp3 | 0.693944 | 0.130681 |
| Lysmd4 | 0.644064 | 0.130549 |
| Fam120b | 0.277141 | -0.129956 |
| Sdr39u1 | 0.371534 | -0.129811 |
| 4933421I07Rik | 0.50325 | 0.129665 |
| Fam120a | 1.005332 | -0.129151 |
| Adgrg2 | 0.174043 | 0.129125 |
| H13 | 0.297806 | 0.128439 |
| Pdss2 | 0.708658 | -0.128109 |
| Lypd5 | 0.596958 | 0.127831 |
| Cyp2c69 | 0.475422 | -0.127646 |
| Dnal1 | 0.340778 | 0.127607 |
| Khdrbs2 | 0.828912 | 0.127514 |
| Gm10922 | 0.291827 | 0.127382 |
| Evpl | 0.834545 | 0.127237 |
| Ubac2 | 0.587417 | 0.126471 |
| Tecta | 0.527004 | 0.126431 |
| Adam4 | 0.251799 | -0.126127 |
| Pcdhb16 | 0.265894 | 0.125968 |
| Utp18 | 0.3615 | 0.125863 |
| Sdpr | 0.200656 | -0.125783 |
| Kbtbd12 | 0.230124 | -0.125373 |
| Arhgap15 | 0.132534 | 0.125175 |
| C1qtnf7 | 0.401002 | -0.125043 |
| Pdrg1 | 0.606335 | -0.123547 |
| Qrsl1 | 0.387088 | 0.123004 |
| 2310057M21Rik | 0.7547 | -0.122712 |
| Lum | 0.137463 | 0.122606 |
| Ptgs2 | 0.134682 | -0.122368 |
| Samhd1 | 0.392834 | 0.121692 |
| Smc6 | 0.393503 | 0.121599 |
| N4bp2 | 0.208914 | -0.121466 |
| Rpl8 | 1.061687 | 0.120976 |
| Ppp1r16b | 0.202087 | -0.120896 |
| Ddx52 | 0.364902 | 0.12075 |
| Snrk | 0.293135 | 0.120458 |
| Champ1 | 1.137268 | 0.120445 |

| | | |
|---|---|---|
| Tyw1 | 0.418972 | -0.120339 |
| Rbm25 | 0.508103 | -0.11998 |
| Xcr1 | 0.241403 | -0.119462 |
| Bambi | 0.24896 | -0.119383 |
| Hao2 | 0.56687 | -0.119303 |
| Fcamr | 0.436699 | -0.118958 |
| Serpinb12 | 0.463141 | 0.118626 |
| Tulp4 | 0.444869 | -0.118572 |
| Vmn1r173 | 0.216973 | 0.118426 |
| Gm11034 | 0.138514 | 0.118386 |
| Tlr2 | 0.218789 | 0.118346 |
| Olfr481 | 0.333498 | -0.118267 |
| Lrp6 | 0.384895 | -0.118094 |
| Gm3755 | 0.206885 | -0.118067 |
| Eif3c | 0.970563 | 0.117921 |
| Ube2j1 | 0.83694 | 0.117655 |
| Gm10521 | 0.305887 | 0.117509 |
| Podn | 0.170551 | -0.117243 |
| Ago1 | 0.502377 | -0.116045 |
| Mphosph8 | 0.476678 | -0.115979 |
| Gm17641 | 0.284412 | 0.115739 |
| Gyg | 0.24769 | 0.115726 |
| Angptl7 | 0.349613 | 0.115206 |
| Ap5s1 | 0.604169 | 0.11518 |
| Adamts19 | 0.331279 | -0.115126 |
| Olfr1312 | 0.184132 | 0.114993 |
| Nudt13 | 0.480114 | -0.114674 |
| Med22 | 0.233504 | 0.11446 |
| Rac3 | 0.291629 | 0.114207 |
| Bcl6 | 0.166402 | 0.113714 |
| Mmadhc | 0.639558 | 0.113714 |
| Natd1 | 0.644794 | 0.113687 |
| Parvb | 0.468604 | 0.113647 |
| Mageb18 | 0.660425 | 0.113607 |
| Ltk | 0.399105 | 0.113567 |
| Ube4b | 0.7292 | -0.11334 |
| Sco2 | 0.496573 | 0.1133 |
| Vmn1r29 | 0.290221 | 0.11306 |
| Olfr1408 | 0.46774 | -0.112927 |
| Gm16367 | 0.329466 | -0.112834 |
| Chn1 | 0.2353 | -0.1125 |
| Cacng4 | 0.518834 | 0.1125 |
| Nln | 0.556074 | 0.112487 |

| | | |
|---|---|---|
| Zfp397 | 0.322335 | -0.112113 |
| Olfr1384 | 0.507207 | -0.111485 |
| Ppp1r18 | 0.37108 | 0.111459 |
| Gm21891 | 0.378746 | 0.111058 |
| Gm21828 | 0.378746 | 0.111058 |
| Gm21725 | 0.378746 | 0.111058 |
| Gm21904 | 0.378746 | 0.111058 |
| Gm21764 | 0.378746 | 0.111058 |
| Gm21852 | 0.378746 | 0.111058 |
| Ptprj | 0.185208 | 0.110858 |
| Itga11 | 0.391108 | 0.11047 |
| Sytl2 | 0.230153 | -0.11031 |
| Rab35 | 0.593971 | 0.110176 |
| Defb43 | 0.333406 | 0.109949 |
| Spdye4b | 0.387615 | 0.109829 |
| Wscd2 | 0.33232 | 0.109561 |
| Sh3bp1 | 0.382621 | 0.109414 |
| Ddhd1 | 0.187252 | 0.109133 |
| Tmem108 | 0.112712 | -0.108999 |
| Srd5a1 | 0.146685 | -0.108973 |
| Phf13 | 0.356041 | 0.108919 |
| Slc30a8 | 0.450136 | -0.108866 |
| Slc6a15 | 0.259505 | 0.108826 |
| Jmjd4 | 0.435372 | -0.108518 |
| Upf1 | 0.50703 | -0.108357 |
| Olfr1318 | 0.412554 | -0.108183 |
| Tug1 | 0.344296 | -0.107875 |
| Znrf4 | 0.463525 | 0.107782 |
| Prdx5 | 0.411601 | 0.106831 |
| H2-M3 | 0.364354 | 0.106563 |
| Gm15107 | 0.320412 | 0.106295 |
| Olfr167 | 0.206797 | -0.106027 |
| Olfr1471 | 0.243359 | -0.105919 |
| Sox21 | 0.372489 | -0.105651 |
| Fut8 | 0.296002 | -0.105624 |
| Rfxank | 0.325527 | -0.10541 |
| Vnn3 | 0.195951 | 0.105356 |
| Olfr1131 | 0.232412 | -0.105101 |
| Vmn2r8 | 0.269468 | -0.104551 |
| Mt2 | 0.120677 | 0.104538 |
| Olfr1465 | 0.262575 | -0.104337 |
| Gm17727 | 0.111157 | 0.103853 |
| Nrp1 | 0.35962 | -0.103585 |

| | | |
|---|---|---|
| Ccdc42 | 0.356064 | 0.103075 |
| Akr1c19 | 0.173458 | 0.102954 |
| Tmprss11g | 0.264258 | 0.102309 |
| Olfr1285 | 0.241024 | -0.102201 |
| B9d2 | 0.226314 | -0.10204 |
| Plaur | 0.124 | 0.101153 |
| Tcp11x2 | 0.339705 | -0.101126 |
| Olfr805 | 0.481109 | -0.101018 |
| Ahnak2 | 0.13933 | -0.100897 |
| Vmn1r86 | 0.206567 | 0.100803 |
| Il6st | 0.259382 | -0.100184 |
| Osbpl9 | 0.757026 | 0.099968 |
| Mrgprd | 0.579384 | 0.099915 |
| Olfr486 | 0.343337 | 0.099686 |
| Atp2a1 | 0.177589 | -0.099565 |
| Mbd6 | 0.434799 | -0.099484 |
| Kcnmb1 | 0.170301 | -0.099282 |
| Tdh | 0.408348 | -0.099012 |
| Rnf115 | 0.619796 | 0.098824 |
| Anapc2 | 1.714893 | 0.098568 |
| Sgcd | 0.205922 | -0.098298 |
| Rars2 | 0.260183 | -0.098298 |
| Olfr482 | 0.11279 | 0.097813 |
| Ldlrad4 | 0.232016 | -0.097705 |
| Pkig | 0.450069 | 0.097557 |
| Creg2 | 0.168327 | 0.097368 |
| Ffar3 | 0.337279 | -0.097368 |
| Speer4c | 0.289332 | -0.097193 |
| Nsg1 | 0.151573 | 0.097166 |
| Cd320 | 0.553457 | 0.096856 |
| Smcp | 0.34406 | -0.096788 |
| Olfr126 | 0.498308 | -0.09641 |
| Cnp | 0.200123 | 0.095978 |
| Mtmr4 | 0.364019 | 0.09587 |
| Fam58b | 0.41007 | 0.095695 |
| Mrc1 | 0.131634 | 0.095425 |
| Olfr1098 | 0.236541 | -0.095411 |
| Bmyc | 0.441635 | 0.09502 |
| Tsr2 | 0.67034 | -0.094371 |
| Atp13a2 | 0.28565 | 0.094128 |
| Gm8842 | 0.421519 | 0.093939 |
| Tmem240 | 0.262628 | 0.093749 |
| Nucb1 | 1.043661 | 0.093574 |

| | | |
|---|---|---|
| Rptn | 0.433246 | 0.093141 |
| Gpx5 | 0.242223 | -0.092383 |
| Ormdl1 | 0.358902 | 0.092356 |
| 2010106E10Rik | 0.46012 | 0.091314 |
| Smarcd3 | 0.391748 | -0.091192 |
| Plaa | 0.225685 | 0.091097 |
| Olfr1289 | 0.324259 | -0.090962 |
| Cyp8b1 | 0.215547 | 0.090935 |
| Trim10 | 0.206122 | -0.090311 |
| Spsb4 | 0.263005 | 0.090068 |
| Kti12 | 0.39984 | -0.08958 |
| Col4a2 | 0.205621 | 0.089498 |
| Prdx6b | 0.362289 | -0.089146 |
| Ppfibp2 | 0.178642 | 0.089064 |
| Ankk1 | 0.271946 | 0.088874 |
| Olfr1168 | 0.084542 | -0.088847 |
| Ints1 | 0.646765 | -0.088766 |
| Drosha | 0.358604 | -0.088467 |
| Ptger1 | 0.256642 | 0.088413 |
| Gm17768 | 0.285902 | -0.088318 |
| Olfr936 | 0.112595 | 0.088142 |
| Papolg | 0.354028 | -0.087517 |
| P2ry12 | 0.095214 | 0.08749 |
| Scgb2b7 | 0.313939 | 0.08749 |
| B3gnt3 | 0.199618 | 0.087341 |
| Thoc3 | 0.339215 | 0.087205 |
| Rasgef1a | 0.253719 | -0.086947 |
| Cyp3a41a | 0.209726 | -0.086784 |
| Sfrp2 | 0.278411 | 0.086498 |
| Vmn1r158 | 0.098625 | -0.08605 |
| Bmper | 0.176549 | 0.085574 |
| Try10 | 0.329659 | 0.085452 |
| Larp4b | 0.743662 | -0.085221 |
| Serpinb3d | 0.419846 | 0.084785 |
| 2010111I01Rik | 0.231569 | -0.084772 |
| Hsp90ab1 | 0.513402 | 0.084677 |
| Mob1b | 0.272355 | 0.084418 |
| Ada | 0.18375 | 0.084391 |
| Mst1r | 0.300815 | 0.084078 |
| Il4 | 0.305756 | -0.083656 |
| Fut10 | 0.249398 | -0.083288 |
| Gins4 | 0.375603 | 0.083247 |
| Oraov1 | 0.422419 | 0.08322 |

| | | |
|---|---|---|
| Gm21767 | 0.279717 | 0.083179 |
| Gm21822 | 0.279717 | 0.083179 |
| Cdh12 | 0.228514 | -0.083138 |
| Pradc1 | 0.1535 | 0.082907 |
| V1ra8 | 0.253163 | -0.08288 |
| P2ry14 | 0.198504 | -0.082852 |
| Lrp8 | 0.092726 | 0.082798 |
| Gm17374 | 0.417846 | -0.082757 |
| Ppbp | 0.07402 | -0.082634 |
| Gm17577 | 0.248569 | -0.08213 |
| Gm17467 | 0.248569 | -0.08213 |
| Spats1 | 0.349252 | -0.081571 |
| Vezt | 0.143257 | -0.081462 |
| Dlx3 | 0.360454 | 0.080835 |
| A430035B10Rik | 0.217517 | 0.080685 |
| Sox15 | 0.290454 | -0.080658 |
| Olfr1368 | 0.222779 | -0.080153 |
| Zfp532 | 0.146187 | -0.079648 |
| Ankrd26 | 0.245686 | -0.079416 |
| Klk10 | 0.256494 | -0.079211 |
| Gm4832 | 0.136974 | 0.079047 |
| Adamts1 | 0.128243 | 0.079006 |
| Vmn2r90 | 0.28599 | -0.079006 |
| Tcp10c | 0.188061 | -0.078705 |
| Pdp1 | 0.271609 | 0.078664 |
| Gm21800 | 0.173373 | -0.07846 |
| Olfr948 | 0.163167 | -0.078337 |
| Parp14 | 0.145597 | -0.078227 |
| Srxn1 | 0.191535 | -0.078063 |
| Klhl18 | 0.28011 | 0.07805 |
| Rfx3 | 0.186474 | -0.07779 |
| Apbb2 | 0.175157 | -0.077749 |
| Rpl19 | 0.58151 | 0.07764 |
| Mybpc2 | 0.214719 | 0.077612 |
| Fgr | 0.096031 | 0.077475 |
| Olfr686 | 0.309611 | -0.077407 |
| Lrrc57 | 0.439351 | 0.077393 |
| Defa-rs2 | 0.286152 | 0.077352 |
| Cyp39a1 | 0.200957 | -0.076833 |
| Itgb2l | 0.142135 | 0.076641 |
| Otud4 | 0.213887 | -0.076532 |
| Gm13213 | 0.338139 | 0.076299 |
| Phospho2 | 0.123586 | 0.076258 |

| | | |
|---|---|---|
| P4ha3 | 0.140602 | 0.076244 |
| Ppm1l | 0.140365 | 0.076244 |
| Med23 | 0.210714 | -0.076176 |
| Trappc5 | 0.262515 | 0.076149 |
| Spred1 | 0.358468 | 0.076025 |
| Itpripl2 | 0.219199 | -0.075984 |
| Olfr726 | 0.196225 | -0.075601 |
| Ccdc36 | 0.319575 | -0.075478 |
| Gm5861 | 0.198068 | -0.075341 |
| Sdhaf2 | 0.353959 | 0.074629 |
| Zfp955b | 0.245715 | -0.074615 |
| Prph | 0.290886 | 0.074478 |
| Acad8 | 0.409205 | -0.074396 |
| Gm6455 | 0.147291 | 0.074355 |
| Ccdc71 | 0.388515 | 0.0743 |
| Tial1 | 0.658999 | -0.074245 |
| Rgs18 | 0.085888 | 0.074177 |
| Fam50a | 0.457212 | 0.073957 |
| Angptl1 | 0.114415 | -0.073875 |
| Sycp1 | 0.362125 | -0.073861 |
| Tmco5 | 0.198318 | -0.073765 |
| Zfp507 | 0.239388 | -0.073601 |
| Olfr954 | 0.134507 | -0.073354 |
| Adar | 0.207112 | -0.073148 |
| Olfr1109 | 0.184313 | 0.072888 |
| Olfr470 | 0.17603 | 0.072833 |
| Olfr576 | 0.215615 | 0.072504 |
| Taf7 | 0.257302 | -0.072298 |
| Mptx1 | 0.131387 | -0.072243 |
| Tpcn1 | 0.236102 | -0.072106 |
| Wisp1 | 0.155975 | 0.071955 |
| Wdr70 | 0.466569 | -0.071872 |
| Gm17535 | 0.099197 | -0.071241 |
| Sgsm1 | 0.367917 | 0.0712 |
| Olfr510 | 0.07893 | 0.071062 |
| Cyp3a41b | 0.111993 | -0.070774 |
| Olfr724 | 0.150158 | -0.070389 |
| Fam166a | 0.120859 | -0.070046 |
| Gmpr2 | 0.252906 | 0.069991 |
| Vmn2r106 | 0.303683 | -0.069427 |
| Psmc2 | 0.39949 | -0.06896 |
| Esrrg | 0.234355 | -0.068836 |
| Olfr1453 | 0.244838 | 0.068767 |

| | | |
|---|---|---|
| Mterf3 | 0.2255 | -0.068712 |
| Sox6 | 0.112546 | -0.068533 |
| Gm8127 | 0.176746 | -0.068437 |
| Ttc14 | 0.168268 | -0.068079 |
| Capn2 | 0.437227 | -0.068079 |
| Fzr1 | 0.342983 | -0.067914 |
| Vmn2r68 | 0.12974 | -0.067721 |
| Gm839 | 0.245935 | 0.067157 |
| Csf1r | 0.144383 | 0.066813 |
| Ate1 | 0.414957 | -0.066785 |
| Glg1 | 0.667471 | -0.066606 |
| Bin2 | 0.120545 | 0.066013 |
| Vmn2r72 | 0.113421 | -0.065958 |
| Zbtb1 | 0.213344 | 0.065903 |
| Spata33 | 0.164409 | -0.065752 |
| Htr1b | 0.149693 | -0.065669 |
| Pgrmc2 | 0.281321 | -0.065614 |
| Nsf | 0.182938 | 0.065586 |
| Olfr1206 | 0.162541 | -0.065572 |
| Iqcf5 | 0.216165 | -0.065379 |
| Tbpl1 | 0.262275 | 0.065297 |
| Dsg4 | 0.204918 | 0.064469 |
| Gm16432 | 0.128145 | 0.064234 |
| Rpl6 | 0.528478 | 0.064207 |
| Aurkaip1 | 0.583451 | 0.064027 |
| Vmn2r91 | 0.171245 | -0.063917 |
| Cdc42se2 | 0.576654 | 0.063655 |
| Slc24a2 | 0.149658 | -0.063544 |
| Olfr1008 | 0.198793 | 0.063351 |
| Gm9944 | 0.124783 | -0.063296 |
| Gm13084 | 0.162774 | -0.06313 |
| Cercam | 0.147201 | 0.062785 |
| Qtrtd1 | 0.194153 | -0.06244 |
| Pdlim2 | 0.363436 | 0.062066 |
| Etv5 | 0.24162 | 0.061693 |
| Slc35f6 | 0.160357 | 0.061043 |
| Gm2777 | 0.08976 | -0.060947 |
| Gm7225 | 0.163747 | 0.060933 |
| Wfdc18 | 0.119975 | -0.060518 |
| Nt5m | 0.080805 | 0.060407 |
| Plin3 | 0.333805 | -0.060324 |
| Dctn4 | 0.339867 | 0.059881 |
| Apcdd1 | 0.079748 | -0.059632 |

| | | |
|---|---|---|
| Mageb3 | 0.261288 | -0.059591 |
| Plekho2 | 0.226481 | -0.058746 |
| Mapk8 | 0.154777 | -0.058524 |
| Prkg2 | 0.199844 | 0.058469 |
| Il9 | 0.146783 | -0.057928 |
| Tldc1 | 0.224741 | 0.057443 |
| Ccdc160 | 0.158964 | -0.05736 |
| Kif13b | 0.135392 | -0.057249 |
| Slc12a8 | 0.140545 | -0.056833 |
| Usp4 | 0.267471 | -0.05675 |
| Treml2 | 0.111142 | 0.056708 |
| Gm3750 | 0.095604 | -0.056459 |
| Arhgef12 | 0.147848 | -0.055973 |
| Fshb | 0.130206 | -0.055918 |
| Zfp804a | 0.126639 | 0.055668 |
| Nlk | 0.133212 | -0.055529 |
| Tcea3 | 0.075503 | 0.055321 |
| 4930404N11Rik | 0.142616 | 0.055196 |
| Olfr801 | 0.154491 | 0.055196 |
| Smok2a | 0.141373 | 0.055112 |
| Nrros | 0.105759 | 0.055057 |
| 4933427D14Rik | 0.158151 | -0.054779 |
| Cphx3 | 0.197185 | 0.054501 |
| Vmn1r177 | 0.229004 | -0.054321 |
| 9430038I01Rik | 0.13608 | 0.053848 |
| Scaper | 0.135893 | -0.05382 |
| Gm11360 | 0.198169 | 0.053417 |
| Mllt3 | 0.129978 | 0.053236 |
| Efhb | 0.09351 | -0.053195 |
| Lpar3 | 0.238543 | 0.052958 |
| Mtch2 | 0.224631 | 0.05268 |
| Gm8356 | 0.1128 | -0.052513 |
| Vmn2r104 | 0.140371 | 0.052402 |
| Zfp955a | 0.165785 | -0.052277 |
| Ticam2 | 0.093638 | -0.052165 |
| Olfr747 | 0.176226 | -0.052082 |
| Trp53i13 | 0.18202 | -0.05179 |
| E330017A01Rik | 0.110429 | -0.051414 |
| Gm17324 | 0.133405 | -0.051177 |
| Vmn1r39 | 0.143799 | -0.051052 |
| Snx18 | 0.135961 | 0.050982 |
| Dpp7 | 0.100165 | 0.050801 |
| Exosc10 | 0.341981 | -0.050286 |

| | | |
|---|---|---|
| Fam195b | 0.304676 | 0.05023 |
| Slc35e2 | 0.153831 | 0.049937 |
| Hars | 0.131025 | 0.049784 |
| Gstt1 | 0.157041 | 0.049589 |
| Btbd8 | 0.128104 | -0.04931 |
| Sgcz | 0.18019 | 0.049073 |
| BC048502 | 0.090051 | -0.04878 |
| Whsc1l1 | 0.289016 | -0.048697 |
| Ttc13 | 0.161693 | -0.048362 |
| Sde2 | 0.230411 | -0.048362 |
| C1qtnf4 | 0.122347 | 0.047929 |
| Traf4 | 0.203215 | -0.047873 |
| Vmn2r35 | 0.131866 | 0.047594 |
| Speer6-ps1 | 0.131377 | -0.047147 |
| Gm3147 | 0.119812 | -0.047134 |
| Arhgef19 | 0.226994 | 0.047022 |
| Zfp280d | 0.099254 | -0.04677 |
| Commd5 | 0.164559 | 0.046631 |
| Shisa5 | 0.324416 | 0.046421 |
| Serinc2 | 0.18585 | 0.046379 |
| Ccar2 | 0.286344 | -0.046337 |
| Exoc1 | 0.125551 | -0.045946 |
| Casd1 | 0.175125 | -0.045681 |
| Dusp5 | 0.106147 | -0.045667 |
| Nfe2 | 0.107422 | 0.045555 |
| Il10ra | 0.078864 | -0.045303 |
| Gmcl1 | 0.112173 | 0.044618 |
| Cnga1 | 0.133856 | 0.04452 |
| Map2k2 | 0.202421 | 0.043778 |
| Borcs5 | 0.087619 | 0.043764 |
| Eps15 | 0.20997 | 0.043498 |
| Rnf38 | 0.152583 | -0.043443 |
| Myo7b | 0.1047 | -0.043148 |
| Hormad1 | 0.161926 | 0.042798 |
| Rabggtb | 0.21267 | 0.042784 |
| Gm21736 | 0.144149 | -0.042406 |
| Fkbp9 | 0.12612 | -0.042378 |
| Lmf1 | 0.15656 | 0.042126 |
| Igfbp6 | 0.058457 | 0.041313 |
| 5430427O19Rik | 0.047634 | -0.040514 |
| Srp54b | 0.422843 | -0.040387 |
| Rph3al | 0.091478 | -0.040303 |
| AW551984 | 0.095393 | -0.040247 |

| | | |
|---|---|---|
| Ocln | 0.069001 | -0.040051 |
| Cd300lf | 0.058472 | -0.040037 |
| Prkcd | 0.127084 | 0.039012 |
| Sptlc1 | 0.105097 | -0.037902 |
| Olfr350 | 0.076785 | -0.037509 |
| Ccdc7b | 0.095972 | -0.036989 |
| Eef1a1 | 0.505723 | 0.036918 |
| Nr2f2 | 0.075354 | 0.036764 |
| Asb6 | 0.156105 | 0.036679 |
| Gm1821 | 0.559833 | -0.036595 |
| Olfr474 | 0.06264 | 0.036327 |
| Syk | 0.082865 | -0.036159 |
| Olfr1277 | 0.066785 | -0.036102 |
| D030025P21Rik | 0.127841 | -0.03606 |
| Cdc123 | 0.215647 | 0.035708 |
| Gm10339 | 0.077517 | 0.035272 |
| Zfp148 | 0.183661 | 0.035202 |
| Sppl2a | 0.115916 | 0.03485 |
| Gm7970 | 0.069959 | -0.034652 |
| Cd2ap | 0.101882 | -0.034441 |
| Glyat | 0.104413 | -0.034357 |
| Olfr492 | 0.047941 | 0.034272 |
| Edem3 | 0.106628 | -0.034244 |
| Psmd1 | 0.164073 | -0.034004 |
| Mospd3 | 0.116224 | 0.033694 |
| F630003A18Rik | 0.090475 | 0.033624 |
| Sec24a | 0.086229 | 0.033596 |
| Atl2 | 0.109116 | 0.033286 |
| Olfr834 | 0.121938 | -0.033271 |
| Tas2r125 | 0.054239 | 0.032919 |
| Stat5b | 0.113576 | -0.032595 |
| Ccdc50 | 0.194746 | -0.032595 |
| Cphx2 | 0.082021 | -0.03258 |
| Pgls | 0.079755 | 0.032073 |
| Slc35f1 | 0.066697 | -0.031932 |
| Dap | 0.094745 | 0.031607 |
| Casc3 | 0.110338 | -0.031353 |
| Olfr469 | 0.054409 | -0.031339 |
| Gm21866 | 0.088142 | -0.03131 |
| Gm17174 | 0.096376 | -0.030576 |
| Olfr357 | 0.081606 | -0.030562 |
| Nek3 | 0.07899 | 0.030534 |
| Cirh1a | 0.168987 | -0.030449 |

| | | |
|---|---|---|
| Smu1 | 0.083311 | 0.030421 |
| Pdia5 | 0.110457 | -0.030378 |
| Nkd2 | 0.049678 | -0.029658 |
| Vmn1r234 | 0.060231 | 0.02963 |
| Nadk | 0.143986 | -0.029403 |
| Diexf | 0.150128 | 0.029234 |
| Strbp | 0.087687 | -0.029163 |
| Zeb2os | 0.05293 | 0.028654 |
| Dmd | 0.051546 | -0.027819 |
| Rpp38 | 0.063001 | -0.027324 |
| Ccl4 | 0.032874 | -0.02731 |
| Rpp25l | 0.105408 | 0.027168 |
| P2rx1 | 0.036121 | 0.026956 |
| Hapln3 | 0.166634 | 0.026956 |
| Sp110 | 0.0593 | -0.026843 |
| Ing3 | 0.09353 | 0.026743 |
| Rbbp9 | 0.143764 | -0.026233 |
| Rps27a | 0.147028 | 0.026049 |
| Gm21951 | 0.030057 | 0.025964 |
| Pkp3 | 0.048636 | 0.025638 |
| Dner | 0.061694 | -0.02493 |
| Eif2a | 0.051492 | -0.02483 |
| Yeats4 | 0.062333 | 0.02483 |
| Igip | 0.084583 | -0.024263 |
| Olfr193 | 0.05663 | -0.024249 |
| Gm14346 | 0.035008 | 0.024235 |
| Esf1 | 0.089843 | -0.023724 |
| Glipr2 | 0.060248 | 0.023412 |
| Pgap2 | 0.096023 | 0.023255 |
| Dcaf6 | 0.090377 | -0.023241 |
| Sobp | 0.033145 | 0.023085 |
| Narfl | 0.067279 | -0.023 |
| Tlr5 | 0.029271 | 0.022616 |
| Olfr1180 | 0.045318 | -0.02246 |
| Usp20 | 0.055104 | -0.022446 |
| Olfr615 | 0.027569 | -0.022247 |
| Slc47a1 | 0.050328 | 0.022204 |
| Gm8267 | 0.06241 | -0.022148 |
| Tnfaip8 | 0.042436 | 0.022133 |
| Gm10323 | 0.061546 | 0.021792 |
| Bahcc1 | 0.052486 | 0.021366 |
| Zbtb9 | 0.047749 | 0.021338 |
| Zfp14 | 0.03093 | 0.021067 |

| | | |
|---|---|---|
| Gabre | 0.048999 | 0.021011 |
| Olfr1247 | 0.033042 | -0.020982 |
| Icmt | 0.086802 | -0.020982 |
| Gm5640 | 0.107762 | 0.020954 |
| Slc23a3 | 0.067386 | 0.020598 |
| Smo | 0.09526 | -0.020356 |
| Fbxo8 | 0.083268 | 0.020314 |
| Gm14496 | 0.044611 | -0.019987 |
| Lct | 0.055619 | 0.019901 |
| Map1s | 0.065502 | -0.019118 |
| Gna12 | 0.112548 | 0.019004 |
| Rad54l2 | 0.046262 | -0.01852 |
| Def8 | 0.033196 | 0.018392 |
| Gm5097 | 0.032181 | -0.017779 |
| S1pr4 | 0.050358 | 0.017722 |
| Rnf187 | 0.108943 | -0.017637 |
| Olfr994 | 0.033736 | -0.017551 |
| Snai1 | 0.054147 | 0.017494 |
| Lemd3 | 0.067106 | -0.01738 |
| Capn1 | 0.06083 | 0.017323 |
| Ovca2 | 0.063193 | 0.01701 |
| Gfod1 | 0.039829 | 0.016881 |
| Rnf40 | 0.051738 | 0.016796 |
| Ssxb10 | 0.03921 | 0.016482 |
| Olfr652 | 0.040542 | -0.016325 |
| Ints6 | 0.052135 | -0.016225 |
| Iqcj | 0.056288 | 0.016083 |
| Gm7247 | 0.045489 | 0.016068 |
| Gm5724 | 0.040631 | -0.01604 |
| Gm21797 | 0.035865 | -0.015954 |
| Cln6 | 0.030352 | 0.015883 |
| Klhl32 | 0.032318 | 0.015455 |
| Ddx56 | 0.034745 | 0.015355 |
| Deaf1 | 0.055196 | 0.014455 |
| Insl6 | 0.018114 | 0.014341 |
| Olfr466 | 0.049283 | -0.014327 |
| Efcab8 | 0.06257 | -0.014127 |
| Stxbp5l | 0.049534 | -0.014098 |
| Cyp4f41-ps | 0.040229 | 0.013841 |
| Cntn1 | 0.035034 | -0.013712 |
| Epsti1 | 0.019965 | 0.013527 |
| Gm7980 | 0.036148 | -0.013498 |
| Olfr1124 | 0.017831 | -0.013412 |

| | | |
|---|---|---|
| Pip4k2a | 0.043065 | 0.013369 |
| Tctn3 | 0.027775 | 0.013312 |
| Vmn2r50 | 0.031197 | 0.013155 |
| Ggt5 | 0.044143 | 0.013112 |
| Amigo1 | 0.060307 | -0.012898 |
| Pde8b | 0.035695 | 0.012612 |
| Gm21964 | 0.028199 | 0.01254 |
| Gpm6a | 0.016542 | -0.012497 |
| Ros1 | 0.018883 | 0.012025 |
| Med13 | 0.034055 | 0.011481 |
| Cd109 | 0.035197 | 0.011109 |
| Kat5 | 0.064998 | -0.010909 |
| Braf | 0.029179 | -0.01088 |
| Ier5 | 0.048667 | 0.010851 |
| Kifap3 | 0.03669 | -0.010565 |
| Ywhaz | 0.071538 | 0.010565 |
| Fam129b | 0.032615 | -0.010322 |
| Ap3b1 | 0.061124 | 0.010279 |
| Ythdf3 | 0.051742 | 0.01025 |
| Sprr3 | 0.02746 | 0.010164 |
| Scnm1 | 0.036076 | 0.010021 |
| Scn5a | 0.037161 | -0.00992 |
| Mta2 | 0.037786 | 0.009863 |
| Sdr16c6 | 0.018077 | -0.009734 |
| Urod | 0.030189 | 0.009677 |
| Gp6 | 0.00864 | -0.009648 |
| Siah1b | 0.023314 | -0.009576 |
| Ckmt2 | 0.010255 | 0.009562 |
| A430089I19Rik | 0.02974 | 0.009376 |
| Tmem154 | 0.016983 | 0.009175 |
| Tmem2 | 0.015123 | 0.009118 |
| Olfr1450 | 0.013338 | 0.008731 |
| Micall1 | 0.027039 | 0.008573 |
| Zbtb48 | 0.019663 | -0.00787 |
| Ncln | 0.020326 | 0.007095 |
| Pomk | 0.018069 | -0.007009 |
| Tppp | 0.011716 | -0.00688 |
| Olfr1499 | 0.019744 | 0.006779 |
| Olfr1061 | 0.01036 | 0.006736 |
| Gm10382 | 0.019907 | 0.006233 |
| Slc2a3 | 0.007321 | 0.006205 |
| Gm884 | 0.011609 | -0.005903 |
| Gm906 | 0.011627 | -0.005774 |

| | | |
|---|---|---|
| Cdkl5 | 0.00798 | 0.005558 |
| Rhoq | 0.015329 | 0.004767 |
| Gm14408 | 0.013992 | -0.004696 |
| Ptpn23 | 0.014886 | 0.00458 |
| Tmem82 | 0.011049 | 0.004523 |
| Prune2 | 0.008624 | 0.004465 |
| Lrriq4 | 0.011316 | 0.004451 |
| Tubb1 | 0.005141 | -0.004207 |
| AW011738 | 0.011622 | 0.004106 |
| Pttg1 | 0.011673 | 0.003948 |
| Gm2964 | 0.006108 | 0.003847 |
| Gm270 | 0.009613 | -0.003645 |
| 1110057P08Rik | 0.004932 | -0.003516 |
| Plac8 | 0.002262 | 0.00343 |
| Dnmbp | 0.008675 | -0.003415 |
| Rabep1 | 0.010908 | -0.003386 |
| Vmn1r44 | 0.00807 | 0.003242 |
| Olfr1195 | 0.007044 | -0.003012 |
| Vmn1r53 | 0.010972 | -0.002868 |
| Vat1 | 0.005225 | -0.002364 |
| Kif1b | 0.004131 | 0.00186 |
| Phb2 | 0.007098 | -0.001759 |
| Smad7 | 0.002769 | -0.001615 |
| Nxnl1 | 0.004778 | -0.001615 |
| Creb1 | 0.004672 | -0.00124 |
| 1700017N19Rik | 0.002703 | -0.000894 |
| Lyar | 0.002422 | -0.00075 |
| Arl5a | 0.002635 | 0.000389 |
| Olfr968 | 0.000485 | 0.000274 |
| Ptpn11 | 0.00054 | -0.000202 |

**Supplementary Table 4.**

Differentially expressed genes between CD68+ macrophages isolated from the hearts of *Ackr3*$^{\Delta Lyve1}$ male mice with no cardiac injury and 7 days post left anterior descending artery (LAD) ligation. N=3 samples per group. One-way ANOVA. -log10 p-values and log2 fold change values are shown. Positive fold change values represent upregulation in LAD ligated *Ackr3*$^{\Delta Lyve1}$ macrophages.

| Gene Symbol | -log10 p-value | log2 fold change |
|---|---|---|
| Htatip2 | 3.533346 | 3.103608 |
| Amica1 | 1.979278 | 2.848085 |
| Retnlg | 2.855227 | 2.616769 |
| Serpinb2 | 2.426745 | 2.32224 |
| Retnla | 2.158303 | -2.253327 |
| Padi4 | 1.933514 | 2.234428 |
| Cilp | 4.296352 | 2.105453 |
| Olr1 | 2.633319 | 2.0732 |
| Trem1 | 2.728516 | 2.011209 |
| Il1rl1 | 2.400526 | -1.878745 |
| Aqp7 | 2.147451 | -1.817071 |
| Cxcr2 | 1.926711 | 1.792056 |
| Tmem252 | 2.877624 | 1.789045 |
| Slc7a2 | 3.329457 | 1.748994 |
| F5 | 2.369852 | 1.716591 |
| Arntl | 3.476889 | 1.675346 |
| Cd226 | 2.37976 | 1.644373 |
| Paqr9 | 3.480955 | -1.63957 |
| Tceanc | 2.685916 | 1.609684 |
| Clec1b | 2.688984 | 1.598227 |
| Car6 | 1.981628 | 1.584611 |
| Il6 | 3.188526 | 1.563295 |
| Slc7a11 | 2.497567 | 1.515849 |
| Fkbp14 | 3.273994 | 1.50402 |
| Serpinb10 | 1.923243 | 1.463753 |
| Ccdc122 | 2.123153 | 1.452516 |
| Hepacam2 | 3.443336 | -1.431013 |
| Cmah | 2.050429 | -1.42219 |
| Tiam2 | 3.242809 | 1.422066 |
| Ppbp | 2.467159 | 1.41224 |
| Spata1 | 4.905833 | 1.400828 |
| Hsd11b1 | 2.740339 | 1.398701 |
| Naip1 | 3.067094 | 1.395238 |
| Cemip | 2.268299 | 1.394558 |
| Tarm1 | 3.578819 | 1.388333 |
| Mmrn1 | 4.714961 | 1.38606 |

| | | |
|---|---|---|
| Pcdh12 | 2.255652 | -1.385613 |
| Ms4a8a | 3.052756 | 1.36945 |
| Ccnd2 | 2.53047 | -1.301102 |
| Ucp3 | 2.0292 | -1.293824 |
| Ms4a4d | 2.399379 | 1.238463 |
| Gfra2 | 3.849548 | -1.235004 |
| Clec7a | 2.332411 | 1.221716 |
| Apod | 1.891384 | 1.220807 |
| Osm | 4.038537 | 1.215865 |
| Nes | 2.262063 | -1.215592 |
| P2ry12 | 2.511809 | 1.201998 |
| Pdgfrl | 2.881854 | 1.19735 |
| Pram1 | 2.30303 | 1.19514 |
| Maob | 2.499939 | -1.18593 |
| F2rl2 | 2.478258 | 1.17282 |
| Mmp13 | 2.070716 | -1.162545 |
| 45538 | 4.0749 | -1.161385 |
| Pkhd1l1 | 2.038099 | -1.16083 |
| Eda2r | 2.231413 | -1.159642 |
| Asah2 | 1.949698 | -1.159022 |
| Kbtbd12 | 3.946783 | -1.152482 |
| Pdk1 | 2.167674 | 1.15106 |
| Uchl1 | 3.482907 | 1.14935 |
| Krtap19-5 | 4.833756 | -1.147294 |
| Atp1a2 | 3.006646 | -1.14338 |
| Nox4 | 3.153247 | 1.140373 |
| Slamf1 | 2.414341 | 1.131425 |
| Rwdd2a | 2.41184 | 1.119516 |
| Cabyr | 2.895984 | 1.118958 |
| Chchd10 | 2.462883 | -1.103142 |
| Pi16 | 2.728038 | 1.100648 |
| Ogfrl1 | 2.655755 | 1.100116 |
| Mapk6 | 2.655869 | 1.095162 |
| Gm13152 | 2.442235 | -1.083615 |
| Steap1 | 1.9094 | 1.079511 |
| D16Ertd472e | 3.699239 | 1.073238 |
| Azin2 | 2.383326 | 1.070224 |
| Lrp8 | 2.250796 | 1.069956 |
| Cxcl14 | 2.236142 | 1.064345 |
| Arl6 | 4.382383 | 1.063317 |
| Rrad | 3.335001 | -1.060407 |
| Cybrd1 | 2.561021 | -1.059245 |
| Gzma | 1.949377 | -1.05446 |

| | | |
|---|---|---|
| Tbxas1 | 1.908291 | 1.043289 |
| Gm16442 | 2.22236 | 1.037509 |
| Usp3 | 2.663468 | 1.034413 |
| Hhipl1 | 1.936532 | 1.032912 |
| Zfp30 | 2.20668 | -1.01867 |
| Lmod2 | 1.901983 | -1.01417 |
| Reck | 1.871782 | -1.013512 |
| Lpar4 | 2.171935 | -1.003307 |
| Ddx17 | 2.040428 | -0.989938 |
| Sytl2 | 3.839096 | -0.989328 |
| Adgrg2 | 2.349078 | 0.982291 |
| Gm1966 | 3.610269 | 0.98137 |
| Gm17530 | 1.919724 | -0.979279 |
| Klhl31 | 1.924446 | -0.977207 |
| Kcnk6 | 3.415177 | 0.976547 |
| Mt2 | 1.990757 | 0.968179 |
| BC048403 | 3.413791 | 0.965781 |
| Olfr488 | 3.175761 | 0.96431 |
| Stfa3 | 2.207436 | 0.963837 |
| Ikbkg | 3.583501 | 0.962675 |
| Crlf1 | 3.874766 | 0.960045 |
| Sgpp2 | 3.013955 | -0.959815 |
| Slc44a5 | 2.158091 | -0.958434 |
| Kctd12b | 2.297226 | -0.957877 |
| Eif4a2 | 1.966713 | -0.956703 |
| Dock7 | 2.482181 | 0.954784 |
| Tecrl | 3.630988 | -0.952945 |
| Masp1 | 2.164316 | 0.952721 |
| Olfr1276 | 2.259191 | 0.947344 |
| Rab27a | 4.381103 | 0.945248 |
| Gstk1 | 3.103651 | -0.942736 |
| Shkbp1 | 3.9864 | 0.937834 |
| Olfr1287 | 2.025932 | 0.937277 |
| Rtn1 | 2.608197 | -0.937141 |
| Me2 | 4.335914 | 0.936101 |
| Thbs2 | 2.470748 | 0.933573 |
| Ccl17 | 4.258077 | 0.92426 |
| Tnfrsf26 | 2.557013 | 0.918936 |
| Borcs6 | 2.711601 | 0.918478 |
| Slc9a4 | 2.458738 | 0.91847 |
| Bnip3 | 2.830129 | 0.91624 |
| Ssc5d | 4.095871 | 0.914878 |
| Prrc1 | 2.963467 | 0.913998 |

| | | |
|---|---|---|
| Vcam1 | 2.260543 | 0.90975 |
| Slfn8 | 2.876567 | 0.906906 |
| Rab43 | 2.664677 | 0.903177 |
| Slc16a12 | 5.524951 | -0.900235 |
| Ppid | 2.302974 | -0.89991 |
| Pcdh7 | 2.002081 | -0.899145 |
| Zfand4 | 2.65073 | 0.896435 |
| Wdr46 | 2.154217 | 0.886847 |
| Il6ra | 2.217788 | 0.885215 |
| Lgi1 | 2.517632 | -0.883339 |
| Tubb1 | 2.202694 | 0.882643 |
| Naga | 2.766468 | 0.877893 |
| Igf2r | 2.340232 | -0.877116 |
| Psmd9 | 2.440521 | 0.875387 |
| Aifm2 | 2.80877 | 0.873766 |
| D930015E06Rik | 3.167187 | 0.870858 |
| Cited2 | 2.868183 | 0.869635 |
| Mpp6 | 2.397619 | 0.86086 |
| Hmgcs2 | 2.137476 | -0.859119 |
| Adgrg6 | 2.173365 | 0.858053 |
| Arrdc3 | 2.721105 | 0.854075 |
| Bves | 2.896073 | -0.853157 |
| Aldh1a1 | 2.757578 | 0.850751 |
| AI607873 | 3.147293 | 0.848854 |
| Def8 | 3.247044 | 0.840298 |
| Inpp4b | 2.507046 | -0.840097 |
| Serpinb8 | 1.952386 | 0.837265 |
| Fshb | 3.902163 | -0.834695 |
| Adam3 | 2.496903 | 0.833829 |
| Cd24a | 2.323693 | 0.833173 |
| Gm6880 | 3.3589 | -0.83242 |
| Gucy2g | 3.637644 | -0.831837 |
| Gm17535 | 2.133276 | -0.830539 |
| Nt5c3 | 2.161451 | 0.830036 |
| Nt5m | 2.059355 | 0.827787 |
| Tigd2 | 2.374738 | 0.827014 |
| Fancb | 1.971075 | -0.826957 |
| Fam187b | 3.053605 | 0.824304 |
| Khdrbs3 | 2.386792 | -0.821718 |
| Spsb3 | 2.414767 | 0.82091 |
| Kif21a | 3.707604 | -0.819194 |
| Tra2a | 1.905791 | -0.813377 |
| Pcdh15 | 2.863685 | -0.811118 |

| | | |
|---|---|---|
| H13 | 3.25718 | 0.806976 |
| Ptprj | 2.375974 | 0.806217 |
| Snx7 | 3.182269 | 0.804111 |
| Impad1 | 2.528012 | 0.8024 |
| Homez | 1.975198 | 0.798017 |
| Kcnk3 | 3.488607 | -0.79516 |
| Nhp2l1 | 2.096154 | -0.795085 |
| Ogn | 2.273817 | 0.792064 |
| Nfe2 | 3.797245 | 0.789312 |
| Cfi | 2.444633 | 0.785433 |
| Smim20 | 2.159322 | 0.784077 |
| Atad5 | 2.113056 | 0.783616 |
| Fdxacb1 | 2.418 | 0.783457 |
| Tlr5 | 1.955335 | 0.781452 |
| Bcl6 | 1.954599 | 0.78136 |
| Pygb | 3.257874 | -0.780545 |
| Gm10775 | 3.758919 | -0.779495 |
| Reep1 | 2.871817 | -0.777468 |
| Gnl3 | 1.973246 | 0.771776 |
| Enpep | 2.226611 | -0.771083 |
| Fads2 | 2.106882 | -0.768239 |
| Ccnd1 | 2.597434 | -0.768231 |
| Mertk | 2.376766 | 0.767773 |
| Naa16 | 2.894766 | -0.765984 |
| Macrod1 | 2.396494 | -0.765076 |
| Rragc | 2.353975 | 0.763276 |
| Taok3 | 2.529313 | 0.761047 |
| Col5a2 | 2.165218 | 0.760528 |
| Hsdl2 | 2.468927 | -0.754708 |
| Asnsd1 | 2.017029 | 0.752894 |
| Rnf24 | 2.011359 | -0.750452 |
| Slc25a15 | 2.06413 | 0.749714 |
| Qsox1 | 2.813408 | 0.749208 |
| Hspb2 | 2.433372 | -0.748393 |
| Krt36 | 3.880665 | -0.74367 |
| Rnf113a2 | 2.8383 | 0.741834 |
| Baiap2 | 3.483662 | 0.738725 |
| 1700029J07Rik | 2.021719 | -0.736735 |
| Rnf135 | 3.273273 | 0.735462 |
| Lrrc20 | 2.931614 | -0.73528 |
| Rnf122 | 2.774595 | -0.73521 |
| Chpf | 1.915041 | 0.734439 |
| Zfp92 | 2.517853 | -0.733632 |

| | | |
|---|---|---|
| Chl1 | 2.295847 | -0.732547 |
| Abcd3 | 2.076794 | -0.732182 |
| Osgepl1 | 2.750183 | -0.729583 |
| Wdr73 | 3.080222 | 0.728086 |
| Ltc4s | 2.493229 | 0.727938 |
| Ky | 2.872328 | -0.72765 |
| Tbc1d4 | 2.599972 | -0.727467 |
| Hagh | 2.046783 | 0.725628 |
| Gcfc2 | 3.555612 | -0.725008 |
| Slc37a4 | 2.912907 | 0.721941 |
| Vmn1r42 | 2.16471 | 0.720489 |
| Cend1 | 2.682833 | -0.719735 |
| Zfp948 | 2.574438 | -0.718438 |
| Gm17404 | 2.365577 | -0.71807 |
| Slc2a12 | 4.049043 | -0.718053 |
| Tmem128 | 2.053178 | 0.716473 |
| Gli1 | 2.957424 | -0.71592 |
| Alg13 | 2.458317 | 0.714892 |
| Gp1ba | 1.888338 | 0.714839 |
| Drd2 | 2.298542 | -0.714602 |
| Lsm11 | 2.217979 | 0.714021 |
| Tnfrsf1b | 3.637148 | 0.713925 |
| Gm3182 | 2.856114 | -0.713916 |
| 6330416G13Rik | 2.237456 | 0.713749 |
| Hsp90aa1 | 2.99974 | -0.713546 |
| Mfap3 | 3.379053 | 0.713018 |
| Pf4 | 1.929859 | 0.712376 |
| Nt5c1b | 1.989789 | 0.711821 |
| P3h1 | 2.816161 | 0.709749 |
| Tmem143 | 2.300376 | -0.709017 |
| Hcn1 | 2.354514 | -0.707869 |
| Riok3 | 2.172994 | 0.706924 |
| Tmem5 | 2.249264 | 0.706623 |
| Gm8246 | 2.294536 | -0.705606 |
| Ercc6l2 | 2.482062 | -0.705589 |
| 4931422A03Rik | 2.652319 | -0.704544 |
| Kank4 | 2.703269 | -0.703145 |
| Gm21560 | 2.211919 | -0.703048 |
| Tmem251 | 2.40749 | 0.702507 |
| Slco2b1 | 2.011914 | -0.697391 |
| Hadha | 3.158402 | -0.6934 |
| Col6a6 | 2.586042 | -0.692731 |
| Agfg2 | 2.13723 | 0.692338 |

| | | |
|---|---|---|
| Lpar1 | 2.615523 | 0.691838 |
| Gm3182 | 2.387282 | -0.691364 |
| Vps52 | 2.570277 | -0.690936 |
| Ccl2 | 2.020146 | 0.69014 |
| Plxdc1 | 2.589201 | -0.68878 |
| Ap4b1 | 2.853593 | 0.68689 |
| A830080D01Rik | 2.075706 | -0.686765 |
| Clu | 2.969242 | -0.685348 |
| Itch | 3.665832 | 0.682789 |
| Treml2 | 2.534237 | 0.681539 |
| Olfr1462 | 2.528827 | 0.679748 |
| Fhad1 | 3.703561 | -0.678306 |
| Tspan5 | 2.412976 | 0.677287 |
| Map3k2 | 2.822553 | 0.67717 |
| Fam198b | 2.484577 | -0.675527 |
| Fibin | 2.25576 | 0.67465 |
| Lonrf2 | 3.562546 | -0.673104 |
| Sfrp5 | 2.037546 | -0.671275 |
| Gsta4 | 2.125159 | -0.671121 |
| Abtb2 | 2.624448 | 0.670976 |
| Map3k9 | 2.116713 | 0.669907 |
| Pfn3 | 3.230959 | -0.669299 |
| Idh3a | 2.146346 | -0.66791 |
| Gm8113 | 3.551728 | -0.667175 |
| Synpo | 1.938043 | 0.666602 |
| Rpl31 | 2.410418 | -0.666302 |
| Pea15a | 2.482737 | -0.666238 |
| Htra4 | 2.358598 | 0.66612 |
| Spryd3 | 3.447742 | 0.664428 |
| Pld1 | 2.619057 | 0.661722 |
| Olfr495 | 2.308827 | 0.661631 |
| Pde6b | 2.199124 | -0.661166 |
| Gm13212 | 2.265239 | 0.660144 |
| Arl5c | 2.013772 | 0.660089 |
| Spink2 | 2.330532 | 0.658838 |
| Cep78 | 2.134205 | 0.658774 |
| Nucb2 | 2.442506 | 0.658591 |
| Eif2ak1 | 2.572946 | 0.658243 |
| Ak8 | 2.831308 | 0.656817 |
| Arf2 | 2.699248 | 0.655517 |
| Ptpro | 1.980842 | 0.653216 |
| Irf6 | 2.070453 | -0.651105 |
| Mfsd9 | 2.361366 | 0.649533 |

| | | |
|---|---|---|
| Sbk2 | 3.808781 | -0.649045 |
| Alx3 | 3.747756 | -0.648668 |
| Fetub | 2.232094 | 0.647996 |
| Wdr62 | 3.836293 | 0.647591 |
| Pgs1 | 3.017456 | 0.646181 |
| Crispld2 | 3.154092 | 0.645951 |
| Nrp2 | 2.368752 | -0.645554 |
| Usp28 | 2.230743 | -0.644262 |
| Cyp1a1 | 2.703205 | -0.64417 |
| Slc33a1 | 3.068743 | 0.643976 |
| Prdx6 | 1.99221 | 0.640926 |
| Slc48a1 | 2.681868 | 0.639954 |
| Gm10339 | 2.804585 | -0.639251 |
| Sel1l | 1.979854 | 0.639093 |
| Palld | 2.515619 | -0.638732 |
| Pdzd4 | 2.498735 | -0.637193 |
| Tead4 | 3.247128 | -0.637054 |
| Micu2 | 2.604925 | 0.6349 |
| Gmpr | 3.397582 | -0.634472 |
| Gm5801 | 3.063437 | 0.633589 |
| Pira6 | 2.733742 | 0.633152 |
| Zfp365 | 2.917926 | -0.631691 |
| Sh2d6 | 3.049396 | -0.631216 |
| Mxra8 | 2.249178 | 0.63103 |
| Olfr338 | 2.584191 | 0.630461 |
| Camkk2 | 3.294454 | 0.629287 |
| Mst1r | 4.022101 | 0.628951 |
| Mmaa | 2.357882 | -0.628382 |
| Adss | 2.437657 | 0.62827 |
| Crat | 2.833238 | -0.62771 |
| Tfcp2 | 2.181806 | -0.624775 |
| Tm2d3 | 2.698642 | 0.623792 |
| Hsdl1 | 3.005747 | 0.621487 |
| Srcin1 | 2.242823 | -0.621102 |
| Lrrc23 | 2.703644 | -0.620023 |
| Olfr1038-ps | 2.43464 | 0.61961 |
| Medag | 3.105178 | 0.618765 |
| Arl14 | 2.170779 | -0.618521 |
| Adra1b | 2.030334 | -0.618436 |
| Bivm | 3.514193 | 0.618229 |
| Mis18a | 2.16824 | 0.617618 |
| 1190007I07Rik | 4.281997 | -0.617496 |
| Cytip | 2.451715 | 0.617026 |

| | | |
|---|---|---|
| Sprr2d | 2.567571 | -0.616621 |
| 4932429P05Rik | 1.988015 | -0.615115 |
| Ubxn6 | 2.540817 | 0.614851 |
| Lzts3 | 2.697399 | -0.614286 |
| Akap17b | 2.794141 | 0.61322 |
| 2510002D24Rik | 3.891878 | -0.61256 |
| Plekhd1 | 2.087814 | -0.612437 |
| Skap1 | 3.354534 | -0.611711 |
| Cxadr | 1.955021 | -0.609774 |
| Xpo6 | 3.514352 | 0.60915 |
| Bcl3 | 2.161789 | 0.608847 |
| 2410016O06Rik | 2.674273 | 0.608724 |
| Klhl40 | 2.37842 | -0.608488 |
| Ttc38 | 1.923269 | -0.608185 |
| Nlrx1 | 2.483774 | 0.607759 |
| Senp2 | 3.201184 | 0.607039 |
| Gpalpp1 | 3.445473 | 0.606954 |
| Gm128 | 2.918286 | -0.60685 |
| Olfr247 | 2.646278 | 0.606556 |
| Cybb | 2.332065 | 0.60521 |
| Cadm2 | 2.137458 | -0.604195 |
| Rbm12b1 | 2.374768 | -0.604147 |
| Trpv4 | 2.044161 | 0.603986 |
| Olfml1 | 2.534454 | 0.603578 |
| AA474331 | 1.908942 | 0.603255 |
| Ccdc185 | 3.179418 | -0.602609 |
| Asph | 2.953204 | -0.602381 |
| Zfp449 | 2.144935 | -0.601297 |
| 0610037L13Rik | 2.902441 | -0.600622 |
| Slc18a1 | 2.892437 | 0.600289 |
| Magea8 | 2.493396 | -0.59787 |
| Gpr84 | 2.103132 | 0.597717 |
| Dpp10 | 2.037058 | -0.597097 |
| Fnip1 | 1.992461 | 0.595647 |
| Nol11 | 3.324691 | 0.595255 |
| Pnma2 | 3.169147 | -0.594214 |
| 1700003F12Rik | 2.68193 | -0.594205 |
| Kif3b | 2.758175 | 0.593794 |
| Rara | 2.973659 | 0.593736 |
| Chka | 1.920099 | 0.593067 |
| Fkbp4 | 2.53239 | -0.59278 |
| Ddx11 | 1.985425 | -0.592703 |
| Lgals2 | 2.056037 | -0.592053 |

| | | |
|---|---|---|
| Neu2 | 3.626031 | -0.590875 |
| Gda | 4.074131 | 0.590578 |
| Trnt1 | 2.195676 | 0.590521 |
| 2700062C07Rik | 2.220588 | 0.590319 |
| Tuft1 | 2.046371 | 0.590204 |
| Ncstn | 3.08383 | 0.58939 |
| Nus1 | 2.688248 | 0.588095 |
| Sh2d4a | 2.278824 | -0.588075 |
| Aldh5a1 | 2.733712 | -0.587442 |
| A730061H03Rik | 2.573416 | -0.586366 |
| Gabra3 | 2.025263 | 0.585972 |
| Pop4 | 1.99518 | 0.585934 |
| Ndufs1 | 2.739011 | -0.585645 |
| Gm10197 | 1.926615 | 0.585107 |
| Ahcy | 1.882116 | 0.584587 |
| Plaa | 2.512512 | 0.584501 |
| Ttc12 | 1.917301 | -0.584424 |
| Ttc25 | 2.042338 | 0.584328 |
| Slc5a3 | 2.057009 | -0.582816 |
| Erbb4 | 2.807823 | -0.582045 |
| Gcsam | 1.86503 | 0.581997 |
| Ivd | 2.047739 | -0.581987 |
| Gckr | 2.146551 | 0.580994 |
| Aagab | 3.469378 | 0.580522 |
| 4930426L09Rik | 1.927416 | 0.58029 |
| Acss1 | 2.372043 | -0.579692 |
| Olfr571 | 2.252974 | -0.579682 |
| Fosl2 | 2.810482 | 0.57889 |
| Olfr753-ps1 | 2.648235 | -0.578659 |
| Asic5 | 2.799284 | -0.57861 |
| Fbxo34 | 2.683503 | 0.578523 |
| Ace | 2.473294 | 0.578243 |
| Unc13a | 2.813849 | 0.576 |
| Scn4a | 2.41379 | -0.575583 |
| Gm2837 | 2.275025 | 0.575448 |
| Polr3f | 1.880926 | 0.574799 |
| Ccl11 | 2.755253 | -0.573811 |
| Gtf2h3 | 1.92544 | -0.573345 |
| Hspa12a | 3.219861 | -0.571978 |
| Traf5 | 2.558211 | -0.571754 |
| Rbbp8nl | 2.162368 | -0.571745 |
| Angptl7 | 2.865944 | 0.571735 |
| Ctage5 | 2.452657 | 0.570725 |

| | | |
|---|---|---|
| Ciita | 2.448622 | -0.570716 |
| Engase | 2.094078 | 0.570162 |
| AW551984 | 2.611206 | -0.569063 |
| Zfp958 | 2.242646 | 0.568888 |
| Git1 | 2.066544 | 0.568489 |
| Gm9112 | 2.51758 | -0.568363 |
| Dcun1d1 | 3.34041 | 0.567954 |
| Rxrb | 1.931458 | 0.567906 |
| Prtn3 | 3.242653 | 0.567098 |
| Spata2l | 1.955021 | 0.564017 |
| Morc3 | 2.673465 | 0.563881 |
| Srrm3 | 3.432815 | -0.563441 |
| Wfdc1 | 2.323559 | -0.562181 |
| Cluh | 2.367634 | -0.56136 |
| Olfr632 | 2.022304 | -0.561106 |
| Klk1b22 | 2.045409 | -0.560989 |
| Rabepk | 1.977654 | 0.560382 |
| Smco1 | 1.915431 | -0.560089 |
| Fam20a | 1.935389 | 0.559795 |
| Tmem115 | 2.534305 | 0.559678 |
| Samd9l | 1.98147 | -0.559502 |
| Gm5901 | 1.987458 | -0.559423 |
| Mvd | 2.48921 | 0.559316 |
| G6bos | 3.305028 | -0.559218 |
| Dnd1 | 3.774399 | -0.558287 |
| Armc2 | 2.3604 | -0.557817 |
| Ccr9 | 4.22569 | 0.557454 |
| Sdcbp2 | 2.047243 | 0.557052 |
| Klhl36 | 2.14367 | 0.556905 |
| Map3k15 | 2.300895 | 0.556385 |
| Swap70 | 2.019148 | -0.555865 |
| D6Ertd527e | 2.289641 | -0.555384 |
| Tcte2 | 3.902281 | -0.554962 |
| Apobec3 | 2.244158 | -0.554166 |
| Clp1 | 3.029188 | 0.553803 |
| Ttc3 | 2.064487 | -0.552898 |
| Prkcb | 2.217262 | 0.552456 |
| Olfr1471 | 2.086053 | 0.5522 |
| Tac1 | 2.419689 | -0.552161 |
| Cx3cl1 | 2.322987 | -0.551521 |
| Zeb2 | 2.991183 | 0.551176 |
| Plod3 | 2.873735 | 0.54961 |
| Cyp2a4 | 3.442575 | -0.549472 |

| | | |
|---|---|---|
| Cd209c | 2.354525 | -0.549226 |
| Ccdc87 | 3.617578 | -0.549009 |
| Saxo2 | 2.066999 | -0.548792 |
| Tiam1 | 1.871905 | 0.548535 |
| Ppp1r21 | 3.071435 | 0.548486 |
| Hdac9 | 2.056703 | -0.548348 |
| Micu1 | 3.105335 | 0.548101 |
| Trim5 | 1.933439 | -0.547756 |
| Mpp4 | 3.516865 | -0.547637 |
| Atp5g1 | 2.333092 | -0.547568 |
| Znhit2 | 3.418245 | 0.547085 |
| Wfikkn2 | 2.710902 | -0.54667 |
| Ffar3 | 3.200626 | -0.544663 |
| Npnt | 2.282378 | -0.544416 |
| Klk15 | 2.604541 | -0.544011 |
| Tox3 | 2.507592 | -0.543991 |
| Ptger1 | 2.724762 | 0.542991 |
| Acat1 | 4.582321 | -0.542496 |
| Ccdc137 | 3.311379 | -0.542218 |
| Zfp551 | 2.403919 | -0.541773 |
| Elf4 | 3.439144 | 0.541654 |
| Ddit4l | 2.313123 | -0.540503 |
| Rac3 | 2.233754 | 0.539859 |
| Pvrl4 | 1.879058 | 0.53973 |
| Zfp772 | 2.254354 | 0.539392 |
| Samd8 | 3.253403 | 0.539323 |
| Zbtb1 | 3.1823 | 0.536789 |
| Ankrd45 | 2.345667 | -0.53656 |
| Mkl1 | 2.632583 | 0.53655 |
| Maf | 2.049118 | 0.536123 |
| Nsa2 | 2.802292 | 0.536113 |
| Xpnpep2 | 2.972634 | -0.536073 |
| Ech1 | 4.574319 | -0.534988 |
| Notch2 | 1.912336 | 0.534679 |
| Spata13 | 2.247072 | -0.53437 |
| Pigk | 3.107308 | 0.533972 |
| Cntnap1 | 2.081063 | -0.533224 |
| Tax1bp1 | 2.672378 | 0.533195 |
| Fam101a | 3.057191 | -0.532177 |
| Mxi1 | 2.421673 | 0.531379 |
| B230217C12Rik | 2.130905 | -0.531069 |
| Frmd4b | 2.364756 | 0.53047 |
| Ascc1 | 1.865026 | 0.529901 |

| | | |
|---|---|---|
| Smim5 | 4.091112 | -0.529721 |
| Nlgn3 | 2.957645 | -0.529601 |
| Wdr11 | 1.878092 | -0.529561 |
| Top1 | 2.069046 | -0.528691 |
| Tbc1d13 | 3.239873 | 0.528241 |
| Olfr1415 | 2.272012 | -0.52686 |
| Tctex1d1 | 3.759578 | -0.52664 |
| Yipf3 | 3.430427 | 0.526359 |
| Slc36a3 | 2.489542 | -0.526049 |
| Flot2 | 2.012382 | 0.525387 |
| Irak3 | 2.722804 | 0.525107 |
| A930033H14Rik | 3.176661 | -0.524084 |
| Olfr449 | 2.322005 | 0.523943 |
| Nfyb | 3.053942 | 0.523542 |
| Ctf1 | 3.137691 | -0.522488 |
| Phf10 | 2.030153 | 0.521985 |
| Zbtb9 | 2.300687 | -0.521845 |
| Dpysl5 | 2.456523 | -0.521091 |
| Gm21800 | 1.957318 | -0.520799 |
| Gm5129 | 2.924471 | -0.520528 |
| C1ra | 2.37358 | 0.520488 |
| Sphk2 | 3.649279 | 0.520085 |
| Inadl | 2.652951 | -0.519995 |
| Pld6 | 2.271947 | -0.51919 |
| Gm10330 | 2.128111 | -0.518908 |
| Trim37 | 2.447951 | -0.518807 |
| Igdcc4 | 3.13652 | -0.517588 |
| Anxa9 | 2.124247 | 0.517528 |
| Ache | 3.706971 | -0.517508 |
| 1700123L14Rik | 2.283396 | -0.517457 |
| Pdhb | 2.4757 | -0.517407 |
| Gm3633 | 2.407232 | -0.516035 |
| Itfg2 | 2.696012 | -0.515844 |
| Eci1 | 2.38961 | -0.515723 |
| Map2k3 | 2.92839 | 0.51544 |
| Mettl6 | 2.231003 | 0.515299 |
| Gm3618 | 2.491083 | -0.514834 |
| Zfp60 | 2.010872 | -0.514753 |
| Tmem120b | 2.352542 | -0.514713 |
| Atp5o | 2.549626 | -0.514531 |
| Lce1f | 2.749924 | -0.513248 |
| Foxi3 | 2.3004 | -0.513238 |
| Brix1 | 1.945663 | 0.512955 |

| | | |
|---|---|---|
| Grik2 | 2.541621 | -0.512692 |
| Pex3 | 1.976735 | 0.512257 |
| Epor | 3.460889 | -0.51167 |
| Tnfsf13b | 2.644679 | 0.510932 |
| Bpifa3 | 3.493316 | -0.510435 |
| Ecsit | 3.789799 | -0.509777 |
| Vmn2r37 | 2.122514 | -0.508976 |
| Serpina9 | 1.935007 | -0.508946 |
| H2-M3 | 2.846179 | 0.508672 |
| Cyc1 | 3.079086 | -0.507992 |
| 1110002L01Rik | 3.070169 | -0.50779 |
| D930048N14Rik | 2.500911 | -0.507648 |
| Ddx43 | 2.182335 | -0.507607 |
| Stxbp3 | 2.304113 | 0.507404 |
| Bnc2 | 1.98328 | 0.50713 |
| Chst12 | 2.570048 | 0.507089 |
| Acvr1 | 1.991847 | -0.506815 |
| Etfb | 2.969267 | -0.506663 |
| Panx2 | 2.422367 | -0.505728 |
| Irx2 | 2.236403 | -0.50523 |
| Bdh2 | 1.93877 | 0.504712 |
| 5031439G07Rik | 2.138208 | 0.504142 |
| Gm10251 | 2.018162 | -0.503603 |
| Gm7225 | 2.440353 | 0.503562 |
| A930017M01Rik | 1.943221 | -0.503552 |
| Wdr48 | 2.375163 | -0.503155 |
| Mettl18 | 4.182138 | 0.503013 |
| Prss54 | 2.056538 | -0.502738 |
| Epgn | 2.100807 | -0.502616 |
| Uhrf1bp1l | 2.012699 | 0.502473 |
| Sec14l2 | 2.303558 | -0.50176 |
| Socs4 | 2.264681 | 0.50175 |
| Lrrc29 | 2.233546 | -0.50173 |
| Olfr483 | 2.336273 | 0.500955 |
| Pex12 | 2.54606 | 0.50071 |
| Spryd4 | 2.428775 | 0.500455 |
| 1700025C18Rik | 2.276896 | -0.499507 |
| Tnfrsf8 | 3.118337 | -0.498588 |
| Cxcl11 | 1.899754 | -0.497382 |
| Tex261 | 2.481928 | 0.496728 |
| Cdkn2b | 2.833949 | 0.496544 |
| Gm829 | 2.216012 | -0.496278 |
| Snx25 | 2.84645 | -0.495572 |

| | | |
|---|---|---|
| Agk | 2.216192 | -0.495327 |
| Palb2 | 2.602182 | 0.494743 |
| Slc1a4 | 2.850507 | 0.494651 |
| Id2 | 2.171537 | 0.494385 |
| Tex22 | 2.445024 | -0.49418 |
| Cyp2d26 | 2.754426 | 0.494139 |
| Brwd3 | 3.182982 | -0.49375 |
| Slc4a2 | 2.742718 | -0.493709 |
| Aatf | 1.889047 | 0.493688 |
| Ankrd28 | 3.636234 | 0.493401 |
| Olfr1121 | 3.34566 | -0.493084 |
| Zfand2b | 2.147561 | 0.492469 |
| Pwp2 | 2.50441 | 0.492325 |
| Gm14501 | 3.238542 | -0.49172 |
| Dennd4c | 2.346218 | 0.491289 |
| Cebpz | 2.671806 | 0.491012 |
| Abcb7 | 2.031357 | 0.490396 |
| Olfr536 | 3.045903 | -0.4898 |
| Cyb5r1 | 1.911226 | -0.489276 |
| Vmn2r38 | 2.038907 | -0.489111 |
| Ypel1 | 2.995111 | -0.48908 |
| Pgr | 1.929825 | 0.48907 |
| Dusp2 | 1.903865 | 0.488762 |
| Vstm4 | 1.89083 | -0.48868 |
| Kif2a | 3.294806 | 0.488422 |
| Olfr985 | 2.136586 | 0.488093 |
| Cfap221 | 2.987264 | -0.48796 |
| Ptpn4 | 2.506031 | -0.487764 |
| Vwa8 | 2.523713 | -0.486127 |
| Fmnl2 | 2.447003 | 0.485684 |
| Ovgp1 | 3.443119 | -0.485602 |
| Epm2a | 2.693569 | -0.485478 |
| D10Wsu102e | 2.098111 | 0.484025 |
| Oxnad1 | 2.68267 | -0.483664 |
| Cand1 | 2.927067 | 0.482301 |
| 4930555F03Rik | 2.434118 | -0.482208 |
| Slit3 | 2.785811 | -0.482063 |
| Sec63 | 3.020139 | 0.482053 |
| Tbx19 | 2.792983 | -0.47981 |
| Atp11a | 2.083488 | -0.479634 |
| Smim1 | 2.408003 | -0.479262 |
| Oit1 | 2.269116 | -0.478599 |
| Dnal1 | 1.942943 | -0.477574 |

| | | |
|---|---|---|
| Epn1 | 2.047253 | 0.477429 |
| Wdyhv1 | 2.228064 | 0.476361 |
| Vasp | 3.048265 | -0.476288 |
| Kcnh1 | 1.968426 | -0.475707 |
| Gm9945 | 2.710012 | -0.47549 |
| Nrde2 | 3.068518 | 0.474753 |
| Tmem170b | 3.53311 | 0.474285 |
| 45545 | 2.14528 | 0.473922 |
| Samhd1 | 2.374705 | 0.473849 |
| Aar2 | 3.147161 | 0.473839 |
| Dpcr1 | 2.758802 | -0.473683 |
| Adrbk1 | 2.176244 | 0.4736 |
| Wif1 | 2.796652 | -0.473465 |
| Atp8a1 | 2.431512 | 0.473247 |
| Sfrp2 | 2.564866 | 0.473184 |
| Zfp647 | 3.154314 | -0.472904 |
| Gm9611 | 2.054968 | -0.472748 |
| Fzd3 | 2.117514 | -0.472446 |
| E130112N10Rik | 3.544721 | -0.47228 |
| Ankrd42 | 2.071105 | -0.472269 |
| 9030624G23Rik | 1.861416 | -0.472165 |
| Btla | 1.972712 | -0.471468 |
| Smyd3 | 2.697537 | 0.471396 |
| Mr1 | 2.366143 | 0.471063 |
| Prr22 | 2.623354 | -0.470958 |
| Pcdhac1 | 2.785548 | -0.470532 |
| Celsr2 | 2.339317 | -0.468761 |
| Dhx38 | 2.417067 | 0.468635 |
| Myh14 | 2.770776 | -0.468375 |
| Aoc2 | 2.205983 | -0.467978 |
| G0s2 | 2.479096 | -0.467749 |
| T2 | 1.971559 | -0.467593 |
| Aif1l | 2.323483 | -0.467467 |
| Ier5l | 2.506159 | -0.467353 |
| Fam207a | 2.115146 | 0.466998 |
| Btbd10 | 1.873271 | 0.466768 |
| Iqcj | 3.446898 | -0.466465 |
| Ifrd2 | 2.189198 | -0.466309 |
| Psmc1 | 2.315069 | 0.466162 |
| Pdc | 2.558438 | -0.465786 |
| Vipr1 | 2.148158 | -0.465619 |
| Fkbp3 | 3.011222 | -0.465598 |
| Dleu7 | 2.5403 | -0.465567 |

| | | |
|---|---|---|
| Smpd5 | 2.115846 | -0.465191 |
| Olfr643 | 2.929718 | -0.464679 |
| Olfr1394 | 2.140994 | -0.46424 |
| Tmem25 | 2.550242 | -0.46403 |
| Src | 2.396005 | -0.46403 |
| Angpt4 | 1.941498 | -0.463821 |
| Gm10081 | 1.912783 | 0.463748 |
| Fut8 | 2.057896 | 0.463193 |
| Rab27b | 2.236588 | 0.463026 |
| Olfr1219 | 2.263759 | 0.462995 |
| Vmn1r41 | 2.052482 | 0.46289 |
| Sectm1a | 2.586072 | -0.462702 |
| Spock1 | 2.498129 | -0.462565 |
| Pla2r1 | 3.014983 | -0.462346 |
| Exosc6 | 2.298214 | 0.462199 |
| Cldn34b4 | 2.334387 | -0.462157 |
| Rimbp3 | 2.344109 | -0.461445 |
| Sp9 | 3.054464 | -0.460743 |
| Olfr983 | 2.651641 | 0.46069 |
| Scn2b | 2.124828 | -0.460376 |
| Wdr7 | 2.801599 | 0.460009 |
| Apof | 2.10952 | -0.459537 |
| Mtr | 2.406496 | 0.459505 |
| Vars2 | 2.575687 | -0.459222 |
| 4930539E08Rik | 2.459121 | -0.458949 |
| Srr | 2.996354 | -0.458907 |
| Prkci | 2.1842 | 0.458077 |
| Mtg1 | 2.126738 | 0.457636 |
| Lrrc40 | 1.874281 | 0.456459 |
| Mcts2 | 2.48759 | -0.456459 |
| Pgm3 | 2.029381 | 0.456249 |
| Gm14180 | 2.394584 | -0.456039 |
| 4931408C20Rik | 2.228222 | 0.456028 |
| Pigp | 2.870188 | -0.455923 |
| Rad52 | 3.351443 | -0.454923 |
| 1700049L16Rik | 1.970425 | 0.45485 |
| Vsir | 2.530315 | 0.454734 |
| Ino80b | 2.100801 | 0.453955 |
| Tfip11 | 2.289179 | 0.453839 |
| 4930423O20Rik | 2.337466 | -0.45366 |
| Fgfbp1 | 2.582741 | -0.453217 |
| Pkm | 2.27832 | 0.452838 |
| C1qtnf2 | 3.153654 | -0.452722 |

| | | |
|---|---|---|
| Gm5124 | 2.281277 | -0.452363 |
| Snrnp40 | 2.745121 | 0.451942 |
| Uqcrb | 3.476506 | -0.451741 |
| Rrp7a | 2.249347 | 0.451625 |
| Slc35e2 | 2.552617 | -0.451583 |
| Amph | 2.042805 | -0.451541 |
| Krtap22-2 | 2.702905 | -0.451119 |
| Rnf138rt1 | 2.076291 | -0.450105 |
| Lyzl6 | 2.519759 | -0.449926 |
| 4933405O20Rik | 2.717446 | 0.449292 |
| Ikbkap | 2.649824 | 0.448827 |
| Ube2d2b | 2.68877 | 0.448679 |
| Faxc | 3.154935 | -0.448584 |
| Zbtb21 | 2.170036 | -0.448499 |
| Sat1 | 2.107888 | 0.447886 |
| Il2ra | 3.203563 | 0.447791 |
| 4930444P10Rik | 3.119515 | -0.44723 |
| Dmrtc1a | 2.671749 | -0.447135 |
| Cdc37l1 | 2.903785 | -0.446807 |
| Ogfod3 | 1.886264 | 0.446764 |
| D3Ertd751e | 1.995026 | -0.44633 |
| Uroc1 | 2.289842 | -0.445547 |
| 2310061I04Rik | 2.382178 | -0.445388 |
| Prpf18 | 2.823084 | -0.445261 |
| 4931414P19Rik | 2.003198 | 0.445091 |
| Slc25a36 | 2.516657 | 0.444996 |
| Brat1 | 3.910257 | -0.443532 |
| Rbm10 | 1.974043 | -0.443225 |
| Pgc | 1.930865 | -0.442705 |
| Trim61 | 2.116989 | -0.442673 |
| Gm13298 | 1.979427 | -0.442439 |
| Fgf8 | 2.043792 | -0.4421 |
| Sapcd1 | 3.184841 | -0.442089 |
| Pcolce | 2.056068 | 0.441877 |
| Cort | 2.518235 | -0.441707 |
| Galnt13 | 2.496795 | 0.441579 |
| Mier1 | 3.046537 | 0.441292 |
| Chst2 | 2.436235 | -0.440963 |
| Srprb | 2.696448 | 0.440283 |
| Actl11 | 1.931069 | -0.440155 |
| Mt3 | 2.191719 | -0.440049 |
| Man2a1 | 2.756491 | 0.439559 |
| Sgpp1 | 1.907423 | 0.439495 |

| | | |
|---|---|---|
| Cacng6 | 2.054485 | -0.438985 |
| Dixdc1 | 1.895906 | -0.438974 |
| Fcamr | 2.466525 | -0.438772 |
| Catip | 1.931755 | -0.438612 |
| Obox6 | 2.021565 | -0.438197 |
| Pdrg1 | 3.218353 | -0.438016 |
| Anapc4 | 2.458335 | 0.437494 |
| Hddc3 | 3.350236 | -0.436652 |
| Prss41 | 1.916272 | -0.436578 |
| Mir339 | 2.171447 | -0.436567 |
| Dcp2 | 2.23 | -0.436514 |
| Kifap3 | 3.231345 | 0.436311 |
| Iglon5 | 2.522011 | -0.436215 |
| Zc2hc1b | 2.054096 | -0.435991 |
| Jph4 | 2.348183 | -0.435842 |
| Vav2 | 1.938246 | 0.435767 |
| Scgb1b7 | 3.819805 | -0.435661 |
| Gtpbp3 | 2.171315 | -0.434775 |
| Angptl8 | 1.939212 | -0.434337 |
| Smu1 | 2.254943 | 0.434327 |
| 45356 | 2.097661 | 0.433558 |
| Spats1 | 3.112824 | -0.433515 |
| Psme2b | 1.878466 | -0.433483 |
| Fgf16 | 2.104775 | -0.433045 |
| Vmn1r37 | 2.738613 | 0.432927 |
| St3gal4 | 3.300939 | 0.432906 |
| Tor3a | 3.409677 | 0.432874 |
| Omp | 3.540085 | -0.432756 |
| Urah | 2.307431 | -0.432735 |
| Rsg1 | 1.892756 | -0.432682 |
| Coq2 | 2.268887 | -0.432072 |
| Neu1 | 1.88975 | 0.431634 |
| Plekhf1 | 3.815195 | 0.431612 |
| Calca | 1.887713 | -0.43143 |
| Gpi1 | 2.761512 | 0.430521 |
| Slc9a1 | 3.973352 | 0.430307 |
| Mus81 | 2.25615 | -0.429503 |
| Parvb | 2.727059 | 0.429418 |
| Ccdc39 | 2.274232 | -0.429204 |
| Slc39a5 | 2.737774 | -0.428539 |
| Cox15 | 2.119188 | 0.42841 |
| Stat6 | 4.015103 | 0.428357 |
| Ppargc1b | 1.968123 | -0.428153 |

| | | |
|---|---|---|
| Tmem175 | 2.02443 | 0.427949 |
| Gabrb3 | 1.893507 | 0.427595 |
| Egr3 | 2.606562 | -0.426479 |
| Pitx1 | 2.963615 | -0.426082 |
| Prap1 | 2.674468 | -0.42605 |
| Gab2 | 2.149206 | 0.425749 |
| Gm9140 | 3.105967 | -0.425094 |
| Gm9970 | 2.884406 | -0.42418 |
| Nanos3 | 1.913761 | -0.424051 |
| Ccdc96 | 2.58481 | -0.423955 |
| Ska2 | 2.669621 | -0.42346 |
| Usp35 | 2.56718 | -0.423395 |
| Olfr312 | 2.207664 | -0.423395 |
| Speer4b | 1.95105 | -0.423245 |
| Cdc5l | 2.031151 | 0.423137 |
| Zfp654 | 2.703131 | 0.422933 |
| Ferd3l | 2.031945 | 0.422922 |
| Tars | 2.327511 | 0.422847 |
| Wdr60 | 2.504772 | 0.422674 |
| Tdh | 2.768243 | -0.421985 |
| Cmas | 2.198157 | 0.421888 |
| 5730522E02Rik | 2.11862 | -0.421727 |
| Hpcal1 | 2.548553 | 0.421458 |
| Tnnc2 | 2.510324 | -0.421199 |
| Mutyh | 2.002777 | -0.420768 |
| Tnk1 | 2.063058 | -0.42066 |
| Wdr5b | 2.755154 | -0.42024 |
| Sptbn5 | 2.326236 | 0.420013 |
| Sall4 | 2.191491 | -0.419809 |
| Sec23b | 3.379794 | 0.419636 |
| Rasal1 | 3.286483 | -0.419377 |
| Rbm43 | 1.866969 | 0.419345 |
| Tmem14a | 2.191707 | -0.418708 |
| Tdp2 | 2.602924 | -0.41832 |
| Bri3bp | 2.473454 | 0.418114 |
| Meaf6 | 2.250411 | 0.418017 |
| Cyp2g1 | 2.883887 | -0.417736 |
| Ranbp9 | 2.110296 | 0.41765 |
| Gm6356 | 2.424065 | -0.417618 |
| Ccdc90b | 3.322218 | 0.417294 |
| Rnf186 | 2.145644 | -0.416969 |
| Mob3c | 2.945387 | -0.416299 |
| Car10 | 2.313758 | -0.41591 |

| | | |
|---|---|---|
| Cd38 | 1.926692 | 0.41512 |
| 4933402J07Rik | 2.298433 | -0.414255 |
| Arl8a | 2.291797 | 0.414071 |
| Krtap31-1 | 1.871107 | -0.413951 |
| Fkbp15 | 2.582744 | 0.413822 |
| Ddx51 | 3.490417 | -0.413735 |
| F630003A18Rik | 2.050136 | 0.413659 |
| Raph1 | 2.003133 | -0.41315 |
| Rab2a | 2.835344 | 0.413063 |
| Usp45 | 2.23047 | 0.412782 |
| Plod1 | 2.245195 | 0.412218 |
| Jmjd7 | 1.909925 | -0.412153 |
| Emc7 | 1.918491 | 0.411828 |
| Wfdc3 | 1.915663 | -0.411643 |
| Heatr3 | 2.710569 | -0.411025 |
| Sebox | 1.901197 | -0.409777 |
| A330041J22Rik | 2.352981 | -0.409722 |
| Plcxd3 | 2.101632 | -0.409581 |
| Nsun4 | 5.120194 | -0.409179 |
| Aph1b | 2.158278 | 0.409016 |
| 3300002I08Rik | 2.694655 | 0.40894 |
| Kremen2 | 2.242222 | -0.408494 |
| Vegfb | 3.43013 | -0.408331 |
| Armc10 | 2.485092 | 0.407864 |
| Kirrel3 | 2.422897 | -0.407538 |
| Vwc2 | 2.34048 | -0.407494 |
| St3gal5 | 2.162061 | 0.407287 |
| Marcksl1 | 4.134222 | -0.405883 |
| Olfr384 | 3.552502 | -0.405785 |
| Tnfrsf19 | 2.087563 | -0.405644 |
| Slc25a17 | 3.132372 | 0.4056 |
| Tssk3 | 2.29918 | -0.405219 |
| Mad2l1bp | 2.757608 | 0.404849 |
| H2-M5 | 2.262127 | -0.404838 |
| Akap14 | 2.794877 | -0.404827 |
| Alg2 | 2.883253 | 0.404554 |
| Olfr1511 | 3.100516 | -0.40402 |
| Fbxo28 | 2.244204 | 0.404009 |
| Abce1 | 2.313923 | 0.403824 |
| Pqbp1 | 2.905854 | 0.403737 |
| Rbm22 | 2.992893 | 0.403595 |
| Usf2 | 1.958757 | 0.403453 |
| Adra1a | 1.872947 | 0.402951 |

| | | |
|---|---|---|
| Cggbp1 | 2.58019 | 0.402646 |
| Wapl | 2.046336 | 0.402613 |
| Chmp4b | 2.143029 | 0.402427 |
| Scnm1 | 3.059392 | -0.402351 |
| Tbc1d10c | 2.667016 | -0.401543 |
| Slc45a2 | 1.97949 | -0.401379 |
| Gm3127 | 2.205916 | -0.401314 |
| Kcp | 2.066522 | -0.40115 |
| Tox | 2.146607 | -0.400308 |
| Olfr458 | 3.146571 | -0.398985 |
| 9330159N05Rik | 2.001085 | -0.398974 |
| Swt1 | 2.316031 | -0.398744 |
| Zkscan8 | 1.894786 | -0.39858 |
| Slc38a7 | 2.369745 | 0.398558 |
| Stam2 | 2.213595 | 0.39846 |
| Try5 | 1.887529 | -0.397737 |
| Ifngr1 | 2.380481 | 0.397693 |
| Lrrd1 | 1.882523 | 0.396949 |
| Flg | 3.024037 | -0.396729 |
| Fam73b | 1.887623 | -0.396587 |
| Cycs | 2.473515 | -0.396554 |
| Frem1 | 2.897028 | -0.396006 |
| Prr30 | 3.229007 | -0.39594 |
| Dnajc8 | 2.777447 | 0.395677 |
| Ubqln1 | 2.312755 | 0.395644 |
| 1700086D15Rik | 1.86824 | -0.395524 |
| Tcl1b4 | 1.936798 | 0.395107 |
| Olfr747 | 2.372944 | 0.395041 |
| Map1s | 2.714717 | 0.39492 |
| Cntfr | 1.96599 | -0.394778 |
| Pcbp2 | 2.764068 | 0.394624 |
| Ankrd27 | 1.895298 | -0.394624 |
| Gm20346 | 2.791989 | -0.393229 |
| S1pr5 | 2.151432 | 0.392647 |
| Rwdd1 | 2.173913 | 0.392603 |
| Cep192 | 2.169099 | 0.392383 |
| Ncln | 2.269984 | 0.392262 |
| Phactr3 | 2.1364 | -0.39168 |
| Lrrc8d | 2.101748 | 0.390404 |
| Wdr92 | 1.882643 | 0.38993 |
| Pkmyt1 | 2.27231 | -0.389479 |
| Gm1979 | 1.921637 | -0.38928 |
| Zfp286 | 2.233624 | -0.389203 |

| | | |
|---|---|---|
| Kcnj10 | 2.670922 | -0.389203 |
| Dnah7b | 2.050561 | -0.388862 |
| Thrb | 2.380921 | -0.388686 |
| Yipf1 | 1.926252 | 0.388465 |
| Cpped1 | 2.013008 | 0.388179 |
| Uso1 | 2.510874 | 0.388013 |
| Fbxw15 | 2.543258 | 0.38691 |
| Klk1b27 | 2.02174 | -0.386789 |
| Hinfp | 1.907802 | -0.386601 |
| Mob4 | 2.437626 | 0.386469 |
| Cox20 | 2.405237 | -0.386381 |
| Wdr83 | 4.679158 | -0.385464 |
| Erich5 | 2.462786 | -0.385232 |
| Gm9969 | 2.103784 | -0.385111 |
| Gm26571 | 2.321493 | -0.385089 |
| Mapt | 1.922599 | -0.384846 |
| Zfp626 | 2.166687 | -0.384558 |
| Dnajc7 | 2.440349 | -0.384481 |
| Stat5b | 2.535625 | -0.384326 |
| Rundc1 | 2.630281 | 0.384182 |
| Cep120 | 2.056806 | -0.38416 |
| Smap2 | 3.109273 | 0.384072 |
| Siglecg | 2.375706 | -0.383939 |
| Lurap1 | 1.957673 | -0.383884 |
| Usp40 | 2.492822 | -0.383818 |
| Piga | 1.860776 | 0.383652 |
| Pabpc5 | 2.47871 | -0.383453 |
| Mia | 1.89237 | -0.383431 |
| Nlrp4f | 1.872902 | -0.382081 |
| Ankib1 | 2.780544 | 0.381704 |
| 4921530L21Rik | 2.12683 | 0.380486 |
| Dmc1 | 2.631518 | -0.380452 |
| Zfp599 | 4.268235 | -0.38043 |
| Gadd45a | 2.167191 | 0.380342 |
| Npy5r | 2.133765 | -0.379255 |
| Ankrd44 | 1.994691 | -0.378711 |
| Nfxl1 | 3.09133 | -0.378678 |
| Prrxl1 | 2.300212 | -0.378334 |
| Tmem248 | 2.217655 | 0.377612 |
| 4932416K20Rik | 1.872487 | -0.377368 |
| Rab21 | 1.966975 | 0.377135 |
| Rexo4 | 2.081305 | -0.377068 |
| Cdh9 | 1.966629 | -0.37669 |

| | | |
|---|---|---|
| Idh3b | 1.93088 | -0.376379 |
| Ccdc24 | 4.264539 | -0.376168 |
| Vps51 | 2.136954 | -0.376113 |
| Gm9930 | 1.929135 | -0.376001 |
| Rab3d | 2.451892 | 0.375557 |
| Ythdf2 | 2.791983 | 0.37549 |
| Mapkapk2 | 2.240295 | 0.375468 |
| Ptbp3 | 4.066443 | 0.375423 |
| Fgfr4 | 1.863939 | -0.375067 |
| Mdga1 | 2.302971 | -0.374288 |
| Vta1 | 2.66333 | 0.373999 |
| Pck2 | 2.411772 | -0.373888 |
| Ccdc190 | 2.430929 | -0.373709 |
| Slc25a30 | 1.911428 | 0.373554 |
| Sry | 2.06469 | -0.372874 |
| 4930415O20Rik | 2.034385 | -0.372116 |
| E130309D02Rik | 1.90947 | 0.371938 |
| Mphosph8 | 2.245835 | -0.371336 |
| Bgn | 2.853977 | 0.37128 |
| Plb1 | 2.61195 | -0.37099 |
| Clcn3 | 1.881835 | 0.370812 |
| Ctla2a | 4.436164 | 0.370678 |
| Pnkd | 2.21868 | -0.370644 |
| Prg3 | 2.23829 | -0.370153 |
| Tbpl1 | 2.516303 | 0.369707 |
| Myl6b | 2.330251 | -0.369662 |
| Grem1 | 2.540813 | -0.369617 |
| Mboat4 | 2.233602 | -0.369182 |
| Abcf3 | 2.647725 | -0.368768 |
| Gm5460 | 2.218377 | -0.368645 |
| Park2 | 2.411573 | -0.3685 |
| Krt16 | 2.227707 | -0.368422 |
| Slc39a14 | 3.056892 | 0.368087 |
| Aifm3 | 2.936186 | -0.367986 |
| Sox9 | 2.023121 | 0.367908 |
| Gm5134 | 1.972887 | -0.367673 |
| Rbm12 | 3.314396 | 0.367539 |
| Cfap157 | 2.843033 | -0.367293 |
| Gm6401 | 1.926531 | 0.366588 |
| Rmnd1 | 2.130642 | -0.36651 |
| Homer3 | 2.226215 | -0.366073 |
| Eef2k | 2.074134 | 0.365894 |
| Lrp10 | 2.674546 | 0.365693 |

| | | |
|---|---|---|
| Cyp4f15 | 1.922828 | -0.36492 |
| Gm13889 | 2.540976 | -0.364158 |
| Dab1 | 1.945211 | -0.363732 |
| Gjd3 | 1.920374 | -0.363698 |
| Ninl | 2.360294 | -0.363014 |
| G3bp1 | 1.905742 | -0.363003 |
| Strn3 | 1.928788 | -0.362969 |
| Sipa1l3 | 2.095001 | -0.362756 |
| Mtmr14 | 2.314244 | 0.362442 |
| Trafd1 | 2.091984 | 0.362206 |
| 1700026L06Rik | 2.656172 | -0.361948 |
| Fbxo38 | 2.477337 | 0.361768 |
| Mphosph9 | 1.973352 | -0.361768 |
| Naa30 | 2.703607 | 0.361701 |
| Efcc1 | 2.49292 | -0.36096 |
| Rpap3 | 2.581382 | 0.360937 |
| Hand2 | 2.503837 | -0.35942 |
| Hoxa11os | 2.342736 | -0.359375 |
| Snd1 | 2.251216 | 0.359263 |
| Chst14 | 3.173931 | 0.359218 |
| 4930550C14Rik | 1.990932 | -0.358678 |
| Mterf1a | 2.145491 | 0.35852 |
| Diablo | 1.908308 | -0.357788 |
| 9930022D16Rik | 1.985001 | -0.357743 |
| Arfip1 | 2.505947 | 0.357496 |
| Pcsk1 | 2.276029 | -0.356538 |
| Serpina11 | 2.678478 | -0.356358 |
| Rras2 | 3.030046 | 0.356313 |
| Apex1 | 1.950728 | 0.355885 |
| Mesp2 | 2.158539 | -0.355219 |
| 4933411K16Rik | 2.658498 | -0.355118 |
| 4930544G11Rik | 2.640757 | -0.354892 |
| Rad1 | 2.74861 | 0.354746 |
| Scgb1b21 | 2.173477 | -0.354509 |
| Ogdh | 2.771033 | -0.35426 |
| Krtap4-9 | 2.515907 | -0.35417 |
| Sox30 | 2.254236 | -0.353594 |
| Tfe3 | 1.932468 | 0.353041 |
| Lrrc15 | 2.542151 | -0.352499 |
| Trap1 | 2.350141 | -0.352182 |
| Glod5 | 1.880302 | -0.352047 |
| Mospd1 | 2.284785 | 0.351922 |
| Zfp865 | 1.953127 | -0.351357 |

| | | |
|---|---|---|
| D930020B18Rik | 2.583604 | 0.349796 |
| Pias1 | 2.279717 | 0.349671 |
| Zfp622 | 1.976941 | 0.348878 |
| Efr3a | 2.271027 | -0.348459 |
| H2afy3 | 2.038765 | -0.348414 |
| BC089597 | 2.187745 | -0.348142 |
| Olfr67 | 1.918206 | 0.346373 |
| Tnfrsf13c | 2.035076 | -0.346327 |
| Serpina3i | 2.372502 | -0.345658 |
| Msto1 | 1.886735 | 0.345022 |
| Fgf21 | 2.255533 | -0.344272 |
| Usp44 | 1.924263 | -0.344238 |
| Krt73 | 1.95122 | -0.344147 |
| Gtf3c4 | 2.117597 | -0.343988 |
| Map3k4 | 1.90023 | -0.343738 |
| Tm9sf3 | 2.393151 | 0.343726 |
| Ndufa5 | 2.465936 | -0.343021 |
| Atf7ip2 | 2.408428 | -0.34293 |
| Fut9 | 2.092375 | -0.342725 |
| Hoxb3 | 2.586982 | -0.342407 |
| Ap5z1 | 2.128186 | -0.342316 |
| Knop1 | 2.829287 | 0.341451 |
| Actg1 | 1.874545 | -0.34128 |
| Hyal1 | 3.729319 | -0.340539 |
| Emc3 | 1.926817 | 0.340448 |
| Fam25c | 1.893932 | -0.339605 |
| Pafah2 | 2.19908 | -0.339491 |
| Mad2l2 | 2.093129 | -0.338898 |
| Pdia5 | 2.285258 | 0.338601 |
| Krtap19-3 | 2.773351 | -0.338373 |
| Plekho2 | 2.190201 | 0.33778 |
| 2310033P09Rik | 2.458915 | -0.337745 |
| Ralb | 3.349832 | 0.337254 |
| D17Wsu92e | 2.229484 | 0.336775 |
| Thoc3 | 2.012067 | 0.336398 |
| Acap3 | 2.133583 | -0.336043 |
| Nphp1 | 2.908607 | -0.335232 |
| Prp2 | 2.165026 | -0.335106 |
| Gm1965 | 2.404683 | -0.334946 |
| Sap30bp | 1.996617 | 0.334374 |
| Nthl1 | 2.353522 | 0.334088 |
| Ndnl2 | 2.315902 | 0.333893 |
| Gorasp1 | 2.633028 | -0.333298 |

| | | |
|---|---|---|
| Ccdc142 | 1.955523 | -0.333206 |
| Crbn | 2.633049 | 0.33221 |
| Ccdc157 | 2.177555 | -0.330076 |
| Invs | 1.866595 | -0.329744 |
| Skida1 | 1.878165 | -0.32948 |
| Fubp3 | 2.073737 | -0.329181 |
| AI837181 | 2.725048 | 0.328986 |
| Adh6-ps1 | 1.946173 | -0.328963 |
| Kcnb1 | 2.575587 | -0.32894 |
| Ankrd13b | 2.614058 | -0.328607 |
| Ipmk | 2.324176 | -0.328572 |
| Rarres1 | 2.03606 | -0.328538 |
| Bnip3l | 2.803478 | 0.327181 |
| Neurod6 | 1.909971 | -0.326583 |
| Creld1 | 2.05787 | -0.32656 |
| Olfr76 | 2.468248 | -0.326514 |
| 1700052K11Rik | 1.993795 | -0.326399 |
| Rab35 | 2.52691 | 0.32625 |
| Cfap206 | 1.91606 | -0.326227 |
| Igsf9 | 2.277315 | -0.32602 |
| Slc7a6os | 1.894224 | -0.325513 |
| Gm11360 | 2.035117 | -0.325398 |
| Lhfpl1 | 2.336066 | -0.32496 |
| Gm8674 | 1.881732 | 0.324926 |
| Evx1 | 2.114964 | -0.324603 |
| 3110082J24Rik | 4.294656 | -0.324142 |
| Mcur1 | 2.493743 | 0.323635 |
| Adam10 | 2.761783 | 0.323163 |
| Olfr620 | 1.891161 | 0.323116 |
| Gldc | 1.872941 | -0.323082 |
| Chsy1 | 2.006978 | 0.32284 |
| Teddm3 | 2.087382 | -0.32202 |
| Gfra4 | 1.923014 | -0.322009 |
| Chchd7 | 1.862156 | -0.321928 |
| Fbxo46 | 1.943289 | 0.321778 |
| 6430571L13Rik | 2.111339 | -0.321455 |
| 1700016D08Rik | 2.055142 | -0.321201 |
| Mrps9 | 2.058105 | 0.320196 |
| Pkp2 | 2.193935 | -0.319618 |
| Zfp108 | 1.92692 | -0.319572 |
| Yae1d1 | 3.100996 | -0.318913 |
| Yy2 | 2.130483 | 0.317756 |
| Tmprss7 | 2.353449 | 0.317443 |

| | | |
|---|---|---|
| Gm13238 | 2.494893 | -0.317385 |
| Dcaf12 | 2.472556 | 0.31735 |
| Ppp1r3d | 2.259434 | -0.317292 |
| Utp3 | 2.009048 | 0.317177 |
| Ccl27b | 1.975211 | -0.316829 |
| Ccl27b | 1.975211 | -0.316829 |
| Mcm8 | 2.303833 | 0.316702 |
| Irf2bp2 | 1.865845 | 0.316493 |
| Nkx2-5 | 1.892288 | -0.315914 |
| Ndrg3 | 2.292273 | 0.315566 |
| Id4 | 3.59194 | -0.314813 |
| Gm10936 | 2.280708 | 0.314256 |
| Tspan17 | 2.385741 | 0.313791 |
| Fmn2 | 1.921819 | -0.313745 |
| Cd151 | 1.918286 | -0.313548 |
| Rhox3f | 2.042939 | -0.31299 |
| Ampd1 | 2.57458 | 0.312561 |
| Gm7932 | 2.156039 | -0.311677 |
| Eef2kmt | 2.100684 | 0.311352 |
| Arih1 | 3.010472 | 0.311212 |
| Ap5s1 | 2.284307 | 0.311189 |
| Cwc27 | 2.1313 | -0.311178 |
| Odam | 2.057808 | -0.31055 |
| Xcl1 | 1.983614 | -0.310422 |
| Vmn1r45 | 2.067074 | -0.309758 |
| Scfd1 | 1.934495 | 0.308804 |
| Wdr26 | 1.947779 | 0.308559 |
| Pklr | 2.313301 | -0.306577 |
| Iqcc | 1.98561 | -0.306437 |
| Fig4 | 2.05942 | 0.306391 |
| 4933434E20Rik | 2.202798 | -0.305959 |
| Stat3 | 2.42292 | 0.305527 |
| Thoc7 | 2.23876 | 0.304651 |
| Mybbp1a | 3.111481 | 0.30464 |
| Zfp747 | 1.931247 | 0.304488 |
| Ubl7 | 2.108552 | 0.304418 |
| Drg2 | 2.187599 | 0.304266 |
| Ltbr | 2.90014 | 0.303915 |
| Smagp | 2.209095 | -0.303892 |
| 5530401A14Rik | 1.890118 | 0.30388 |
| Csmd1 | 2.089804 | -0.303763 |
| Crtap | 2.28686 | 0.302921 |
| Gm6498 | 2.498912 | -0.302886 |

| | | |
|---|---|---|
| D930007J09Rik | 1.876227 | -0.302734 |
| Dffa | 1.896501 | -0.301389 |
| Nomo1 | 2.845813 | -0.301154 |
| Lta4h | 2.038205 | -0.300698 |
| Eif3c | 3.308614 | 0.300698 |
| Smtnl1 | 1.862409 | -0.300077 |
| Faap20 | 3.67382 | 0.299292 |
| Zfp33b | 2.48709 | -0.298236 |
| Idnk | 1.89029 | -0.297802 |
| Lyg1 | 2.208589 | -0.296639 |
| E130304I02Rik | 2.251686 | -0.29624 |
| Tab2 | 2.652338 | 0.295417 |
| Mmd | 1.975067 | -0.293111 |
| Slc1a7 | 2.196082 | -0.293041 |
| Mettl9 | 2.171388 | 0.29224 |
| Chchd3 | 2.216481 | -0.291167 |
| Lancl2 | 1.932989 | 0.290542 |
| Sprtn | 1.961471 | 0.289563 |
| Emc1 | 2.631531 | 0.289244 |
| Urm1 | 2.652332 | 0.288878 |
| Cdca4 | 2.01454 | 0.288193 |
| Plrg1 | 3.48654 | 0.286739 |
| Avp | 2.144155 | -0.286526 |
| Pced1b | 2.449577 | -0.28642 |
| Zbtb7b | 2.16542 | 0.286408 |
| Smarca5-ps | 1.87031 | 0.286408 |
| Actrt3 | 1.948658 | -0.286053 |
| Ubqln2 | 1.933033 | 0.28597 |
| Porcn | 4.014308 | -0.285556 |
| Dalrd3 | 2.089554 | -0.285083 |
| Ulk3 | 2.039913 | -0.284633 |
| C030039L03Rik | 3.034707 | -0.284408 |
| Tmem125 | 2.369772 | -0.283294 |
| Ss18l1 | 2.065909 | -0.28295 |
| Fbxl15 | 2.002984 | 0.282523 |
| Osbpl9 | 2.991004 | 0.282072 |
| Olfr192 | 2.30054 | 0.28187 |
| Etnk2 | 2.383434 | -0.28117 |
| Sfxn3 | 2.981383 | 0.28098 |
| Thap7 | 2.70727 | 0.28022 |
| Tmub1 | 2.033812 | 0.279863 |
| Gnb2 | 3.811868 | 0.279507 |
| Mmadhc | 2.130688 | 0.278853 |

| | | |
|---|---|---|
| Apba3 | 2.298581 | 0.27733 |
| Gpx6 | 2.027429 | -0.27689 |
| Wfdc6b | 1.867718 | -0.274721 |
| Dpagt1 | 2.605657 | 0.27459 |
| Fem1c | 2.453275 | 0.273755 |
| Cox7c | 4.83886 | -0.272692 |
| Tapbp | 2.44906 | -0.272692 |
| Gm136 | 1.868061 | 0.272489 |
| Spag6l | 2.375919 | -0.272298 |
| Ppp2cb | 1.911853 | 0.271581 |
| Tex15 | 2.032735 | -0.270888 |
| Ccdc71 | 2.14739 | 0.270074 |
| Slc6a9 | 1.949624 | -0.269153 |
| Snip1 | 2.001049 | -0.268794 |
| Plpp7 | 2.305884 | -0.2675 |
| Uqcrc2 | 2.085637 | -0.267464 |
| Gm8994 | 1.884067 | 0.265149 |
| 1700028J19Rik | 2.239045 | -0.264849 |
| Hdac1 | 2.071 | 0.264741 |
| Phb2 | 2.157677 | 0.263155 |
| Olfr1469 | 1.921543 | 0.262217 |
| Csnk2b | 3.344357 | 0.261784 |
| Dhx36 | 2.095138 | 0.261687 |
| Fer1l6 | 2.262432 | -0.261278 |
| Cdyl | 1.912719 | 0.260038 |
| Fank1 | 2.562687 | -0.258254 |
| Rfk | 2.008194 | 0.25689 |
| Slu7 | 2.354553 | 0.256781 |
| Dazap2 | 2.00024 | 0.256516 |
| Dvl2 | 2.543695 | -0.254001 |
| Pigo | 1.905033 | -0.253421 |
| Cfl2 | 2.164959 | -0.253094 |
| Rabif | 2.411316 | 0.252585 |
| Slc9a8 | 2.783415 | -0.252307 |
| Radil | 1.953923 | -0.251737 |
| Gtf2a1l | 1.900344 | -0.251022 |
| Mdfic | 2.329519 | 0.250852 |
| Gtf2a1 | 2.085889 | 0.250683 |
| 4930480E11Rik | 1.917452 | 0.250367 |
| Ppm1g | 2.022863 | 0.249542 |
| Ptms | 3.308599 | -0.248693 |
| Olfr380 | 3.760148 | 0.248341 |
| Unc79 | 2.233231 | -0.248183 |

| | | |
|---|---|---|
| Fpgs | 2.146625 | -0.248134 |
| Ube2d1 | 2.436407 | 0.24777 |
| Ei24 | 2.130183 | -0.246518 |
| Hsd11b2 | 2.014416 | -0.245313 |
| Ccdc181 | 2.238035 | -0.24496 |
| Etv6 | 3.053899 | 0.243608 |
| Fam134a | 2.930739 | 0.243438 |
| Gm8325 | 3.131152 | 0.242791 |
| Ddi2 | 1.909795 | 0.240204 |
| Tbcc | 2.617468 | 0.239215 |
| Acad8 | 1.929316 | -0.239166 |
| Msl3 | 3.150541 | 0.23787 |
| Suclg2 | 2.579958 | -0.237246 |
| Znhit6 | 2.675485 | 0.235163 |
| Tfpt | 2.230928 | 0.235127 |
| Prkar1a | 2.849618 | 0.234832 |
| Gm21863 | 1.885666 | -0.234023 |
| Cpz | 2.205621 | -0.233765 |
| Ptpru | 2.190347 | -0.231064 |
| Napg | 2.258823 | -0.224892 |
| Slc35a4 | 2.117951 | 0.224571 |
| Agtr1b | 2.066608 | -0.224349 |
| Rdx | 2.04747 | 0.223793 |
| Abhd10 | 1.875851 | -0.221085 |
| Wwp2 | 1.87887 | 0.220862 |
| Dnajc14 | 1.985782 | 0.219872 |
| Shoc2 | 2.794625 | 0.219624 |
| F13b | 2.281027 | -0.217293 |
| Tada1 | 2.337591 | 0.216747 |
| Foxp4 | 2.143445 | -0.214635 |
| Rabac1 | 2.046686 | 0.214448 |
| Tecr | 1.867699 | -0.213478 |
| Slc39a9 | 3.000383 | 0.209765 |
| Dnttip2 | 1.889316 | 0.209229 |
| Pfdn2 | 2.117267 | 0.206518 |
| Gtf2f1 | 2.37163 | 0.202123 |
| Kat5 | 2.318885 | 0.200642 |
| Tmem41b | 2.897309 | -0.197035 |
| Tceb3 | 2.178329 | 0.196959 |
| Myl12a | 1.986287 | -0.196393 |
| Nono | 3.080869 | -0.192737 |
| Elf1 | 1.90526 | 0.190906 |
| Akt1s1 | 2.436742 | -0.190804 |

| | | |
|---|---|---|
| Cdc42se2 | 2.408213 | 0.185664 |
| Ywhaz | 2.305361 | 0.176782 |
| Zfp341 | 2.132498 | -0.165442 |
| Rps16 | 3.290496 | 0.164297 |
| Rps6 | 2.12776 | 0.150937 |
| Slbp | 1.911992 | -0.144268 |
| Rpl13a | 1.87196 | 0.084813 |

**Supplementary Table 5.**

Differentially expressed genes between CD68 + macrophages isolated from the hearts of *Ackr3*$^{fl/fl}$ and *Ackr3*$^{\Delta Lyve1}$ male mice 7 days post left anterior descending artery (LAD) ligation. N=3 samples per group. One-way ANOVA. -log10 p-values and log2 fold change values are shown. Only genes that are differentially expressed in *Ackr3*$^{\Delta Lyve1}$ macrophages upon LAD ligation are shown. Positive fold change values represent upregulation in LAD ligated *Ackr3*$^{\Delta Lyve1}$ macrophages.

| Gene Symbol | -log10 p-value | log2 fold change |
|---|---|---|
| Retnla | 1.320215 | -1.559922 |
| Amica1 | 0.749499 | 1.384625 |
| Il1rl1 | 1.560603 | -1.355744 |
| Cilp | 2.488439 | 1.328446 |
| Apod | 2.095519 | 1.319502 |
| Tmem252 | 1.333546 | 0.995159 |
| Serpinb10 | 1.095474 | 0.964029 |
| Cabyr | 2.381127 | 0.95787 |
| Tiam2 | 1.841945 | 0.911531 |
| Serpinb2 | 0.688191 | 0.905212 |
| Hepacam2 | 1.884366 | -0.887057 |
| Fkbp14 | 1.539153 | 0.836393 |
| Hhipl1 | 1.4323 | 0.824727 |
| Gfra2 | 2.321433 | -0.81501 |
| Fam20a | 3.115284 | 0.813746 |
| Btla | 3.872707 | -0.81359 |
| Slc7a2 | 1.184692 | 0.790413 |
| Paqr9 | 1.336514 | -0.779856 |
| Retnlg | 0.534026 | 0.738716 |
| Nox4 | 1.780583 | 0.729427 |
| Steap1 | 1.095071 | 0.714663 |
| Ogn | 1.980896 | 0.712763 |
| Ucp3 | 0.883801 | -0.698014 |
| Maob | 1.222861 | -0.69232 |
| Trim5 | 2.538562 | -0.675599 |
| Pi16 | 1.429445 | 0.67276 |
| 4930426L09Rik | 2.33006 | 0.670922 |
| Chl1 | 2.023749 | -0.66502 |
| Naip1 | 1.15942 | 0.66451 |
| Stfa3 | 1.309893 | 0.652023 |
| Cmah | 0.707396 | -0.644484 |
| Sh2d4a | 2.557249 | -0.643459 |
| Aldh1a1 | 1.886194 | 0.633896 |
| Eda2r | 0.985576 | -0.630713 |
| Tarm1 | 1.203912 | 0.597011 |

| | | |
|---|---|---|
| Olr1 | 0.493523 | 0.587442 |
| Bnip3 | 1.527122 | 0.571017 |
| Crispld2 | 2.663955 | 0.563168 |
| Sdcbp2 | 2.073873 | 0.562562 |
| Hsd11b1 | 0.804305 | 0.556738 |
| Olfml1 | 2.274948 | 0.554805 |
| Htatip2 | 0.371969 | 0.55221 |
| Nes | 0.778956 | -0.549245 |
| Gm829 | 2.456309 | -0.537564 |
| Tbc1d4 | 1.751934 | -0.537425 |
| Fibin | 1.659705 | 0.534848 |
| St3gal5 | 3.039236 | 0.533224 |
| Cfi | 1.462891 | 0.532636 |
| Uchl1 | 1.278701 | 0.528881 |
| Car6 | 0.454976 | 0.525127 |
| Qsox1 | 1.759723 | 0.521413 |
| Tceanc | 0.5899 | 0.515188 |
| Abtb2 | 1.861596 | 0.514966 |
| Git1 | 1.810785 | 0.514451 |
| Abcd3 | 1.282383 | -0.510942 |
| Atp1a2 | 1.03709 | -0.509817 |
| Vcam1 | 1.020048 | 0.502473 |
| Fetub | 1.593422 | 0.502443 |
| Ace | 2.035934 | 0.497628 |
| A730061H03Rik | 2.059452 | -0.493401 |
| Mst1r | 2.938152 | 0.481196 |
| Gabrb3 | 2.2003 | 0.479365 |
| Lpar1 | 1.610963 | 0.477833 |
| Lsm11 | 1.28017 | 0.473174 |
| Nhp2l1 | 1.040509 | -0.472228 |
| Cxadr | 1.387378 | -0.472145 |
| Inpp4b | 1.161029 | -0.470969 |
| 45538 | 1.291019 | -0.468948 |
| Rab27a | 1.802177 | 0.457521 |
| Enpep | 1.097178 | -0.454702 |
| Crlf1 | 1.499528 | 0.452943 |
| Ms4a8a | 0.701317 | 0.450612 |
| Aqp7 | 0.3401 | -0.449197 |
| Angptl7 | 2.09675 | 0.447495 |
| Rrad | 1.073945 | -0.445144 |
| Fshb | 1.759196 | -0.44075 |
| Rnf24 | 0.975613 | -0.43858 |
| Clec7a | 0.592115 | -0.438154 |

| | | |
|---|---|---|
| Mxra8 | 1.363443 | 0.433397 |
| Cpped1 | 2.317154 | 0.432414 |
| Nfe2 | 1.739149 | 0.423298 |
| Ascc1 | 1.375531 | 0.42247 |
| Gm1966 | 1.212849 | -0.421124 |
| Slc16a12 | 2.318188 | -0.42052 |
| 1190007I07Rik | 2.694782 | -0.416245 |
| Ccl17 | 1.563782 | 0.41315 |
| Olfr983 | 2.296869 | 0.411621 |
| Adgrg2 | 0.728514 | 0.409939 |
| Mmp13 | 0.512657 | -0.408918 |
| Ikbkg | 1.1827 | 0.408125 |
| Atad5 | 0.873388 | 0.406319 |
| Ccr9 | 2.914535 | 0.405143 |
| Pdgfrl | 0.68303 | 0.404805 |
| Zbtb21 | 1.868956 | -0.400385 |
| Krtap19-5 | 1.303748 | -0.397748 |
| Ercc6l2 | 1.154297 | -0.396981 |
| Ogfrl1 | 0.681978 | 0.396675 |
| 4931422A03Rik | 1.236663 | -0.395765 |
| Usp28 | 1.135297 | -0.38906 |
| Medag | 1.70556 | 0.388145 |
| Arl6 | 1.230041 | 0.387605 |
| Wdr11 | 1.232219 | -0.387241 |
| Asph | 1.666853 | -0.387065 |
| Trpv4 | 1.117165 | 0.385354 |
| Gm16442 | 0.58797 | 0.384182 |
| Plxdc1 | 1.185589 | -0.382302 |
| Ms4a4d | 0.500408 | 0.381649 |
| Ccnd2 | 0.495391 | -0.381582 |
| Spata1 | 0.941676 | 0.376513 |
| Cxcl14 | 0.554749 | 0.375134 |
| Acss1 | 1.32177 | -0.373409 |
| Idh3a | 0.977823 | -0.372094 |
| Asic5 | 1.573285 | -0.371938 |
| Vps52 | 1.124748 | -0.371035 |
| Nucb2 | 1.109704 | 0.364494 |
| Sprtn | 2.614896 | 0.361645 |
| Slc1a4 | 1.904228 | 0.361443 |
| Tnfrsf26 | 0.735114 | 0.361297 |
| Olfr338 | 1.232342 | 0.3606 |
| Gstk1 | 0.871187 | -0.359915 |
| Gckr | 1.121809 | 0.358205 |

| | | |
|---|---|---|
| Pkhd1l1 | 0.426214 | -0.357698 |
| Asah2 | 0.409229 | -0.357665 |
| Cd24a | 0.743321 | 0.355986 |
| Tmem143 | 0.908154 | -0.35452 |
| Ccnd1 | 0.924522 | -0.353956 |
| Gm17530 | 0.490726 | -0.352646 |
| Akap14 | 2.344683 | -0.352386 |
| Thbs2 | 0.661354 | 0.348198 |
| Slc44a5 | 0.556617 | -0.348051 |
| Tigd2 | 0.739266 | 0.346021 |
| Gm128 | 1.397043 | -0.345396 |
| Col5a2 | 0.735428 | 0.340403 |
| Mmrn1 | 0.790632 | 0.339924 |
| Rtn1 | 0.670816 | -0.338544 |
| Sptbn5 | 1.746146 | 0.337711 |
| Chpf | 0.672086 | 0.33722 |
| Naga | 0.77217 | 0.336375 |
| Cyp2d26 | 1.660954 | 0.335026 |
| Ttc25 | 0.959987 | 0.333538 |
| 6330416G13Rik | 0.80425 | 0.333092 |
| Ska2 | 1.954305 | -0.332588 |
| 4930480E11Rik | 2.75298 | 0.33143 |
| Ddx17 | 0.473295 | -0.330914 |
| Slco2b1 | 0.735768 | -0.330249 |
| Gabra3 | 0.934178 | 0.329893 |
| Borcs6 | 0.692423 | 0.329491 |
| Fgfbp1 | 1.700249 | -0.329135 |
| Il6 | 0.410422 | 0.328101 |
| Slc18a1 | 1.301463 | 0.326606 |
| Usp3 | 0.574021 | 0.326399 |
| Bdh2 | 1.073864 | 0.325571 |
| Tnfsf13b | 1.460812 | 0.325306 |
| Slc25a30 | 1.583158 | 0.324419 |
| Spata13 | 1.149381 | -0.32382 |
| Ptger1 | 1.379723 | 0.323635 |
| 0610037L13Rik | 1.288517 | -0.323474 |
| Sec14l2 | 1.28189 | -0.323255 |
| Olfr1287 | 0.488432 | -0.322851 |
| Pygb | 1.022703 | -0.32232 |
| Fmnl2 | 1.421078 | 0.322124 |
| Irak3 | 1.429208 | 0.32142 |
| Tspan5 | 0.888452 | 0.321028 |
| Hmgcs2 | 0.570119 | -0.320126 |

| | | |
|---|---|---|
| Tnfrsf8 | 1.759204 | -0.31897 |
| Lpar4 | 0.468797 | -0.317385 |
| Smpd5 | 1.267066 | -0.317223 |
| Ddx43 | 1.161892 | -0.317119 |
| Kcnk6 | 0.770569 | 0.314024 |
| Trim37 | 1.25525 | -0.313884 |
| Spryd3 | 1.275868 | 0.307895 |
| Cfap221 | 1.646255 | -0.307755 |
| Fosl2 | 1.221381 | 0.307615 |
| Cdh9 | 1.497643 | -0.307265 |
| F5 | 0.2515 | 0.306787 |
| Igf2r | 0.573092 | -0.306006 |
| Kcnh1 | 1.072785 | -0.303132 |
| Prtn3 | 1.440703 | 0.302723 |
| Arih1 | 2.90128 | 0.301974 |
| D16Ertd472e | 0.706587 | 0.301892 |
| Klhl31 | 0.404353 | -0.301658 |
| Pde6b | 0.766245 | -0.301564 |
| Agfg2 | 0.698414 | 0.301061 |
| Rabepk | 0.846307 | 0.298506 |
| Ddx11 | 0.783808 | -0.297274 |
| Ankrd45 | 1.06399 | -0.297097 |
| P3h1 | 0.88313 | 0.296311 |
| Chka | 0.752606 | 0.295935 |
| Map3k15 | 0.986847 | 0.295805 |
| Spata2l | 0.819026 | 0.29577 |
| Cdkn2b | 1.435731 | 0.295312 |
| Vmn1r37 | 1.655555 | 0.294171 |
| Eci1 | 1.123405 | -0.292982 |
| Tbxas1 | 0.355671 | -0.292511 |
| Gm7225 | 1.173777 | 0.290672 |
| D17Wsu92e | 1.833253 | 0.290224 |
| Arl14 | 0.783153 | -0.289634 |
| D930020B18Rik | 1.976612 | 0.284408 |
| Map3k9 | 0.658453 | 0.280683 |
| Pcolce | 1.115878 | 0.280434 |
| Hadha | 0.959809 | -0.280362 |
| Gjd3 | 1.351585 | -0.279982 |
| Camkk2 | 1.145791 | 0.279959 |
| Hsdl2 | 0.653829 | -0.27921 |
| Aifm2 | 0.614247 | 0.27827 |
| Tiam1 | 0.749092 | 0.278044 |
| Actrt3 | 1.869657 | -0.277259 |

| | | |
|---|---|---|
| Lrrc20 | 0.803509 | -0.276949 |
| Palld | 0.822602 | -0.276711 |
| Gp1ba | 0.529597 | 0.276509 |
| Grem1 | 1.727773 | -0.275198 |
| Olfr1462 | 0.754426 | 0.2749 |
| Dhx38 | 1.186633 | 0.274733 |
| Baiap2 | 0.957984 | 0.274637 |
| 2410016O06Rik | 0.92333 | 0.273778 |
| Crat | 0.936449 | -0.272382 |
| 5530401A14Rik | 1.632776 | 0.272334 |
| Unc79 | 2.51503 | -0.272226 |
| Ech1 | 2.019994 | -0.271043 |
| Ttc3 | 0.78794 | -0.270254 |
| Ssc5d | 0.850153 | 0.270015 |
| Man2a1 | 1.451378 | 0.269464 |
| Nrp2 | 0.733131 | -0.268937 |
| Etfb | 1.295336 | -0.268913 |
| Ahcy | 0.661231 | 0.268626 |
| Sgpp2 | 0.55729 | -0.267943 |
| Zfp33b | 2.154457 | -0.266829 |
| Slc37a4 | 0.775863 | 0.266289 |
| Sec63 | 1.382321 | 0.264717 |
| Lrp8 | 0.355548 | 0.264428 |
| Mmaa | 0.737651 | -0.263936 |
| Epor | 1.481332 | -0.2639 |
| AI837181 | 2.046473 | 0.263239 |
| Psmd9 | 0.492897 | 0.263095 |
| Fbxo38 | 1.625123 | 0.262493 |
| Bcl6 | 0.455312 | -0.261796 |
| Pgr | 0.827485 | 0.260977 |
| Gm13298 | 0.971547 | -0.260917 |
| Cyb5r1 | 0.818574 | -0.260893 |
| Swap70 | 0.722654 | -0.258989 |
| Pcdh12 | 0.252727 | -0.258338 |
| Ppargc1b | 0.992205 | -0.257566 |
| Def8 | 0.682892 | 0.257481 |
| Stat3 | 1.93871 | 0.257469 |
| Gm9112 | 0.873489 | -0.257324 |
| A830080D01Rik | 0.557495 | -0.25724 |
| Inadl | 1.036855 | -0.256455 |
| Tmem170b | 1.607368 | 0.255344 |
| Pgc | 0.914117 | 0.254292 |
| Vasp | 1.334178 | -0.252864 |

| | | |
|---|---|---|
| Spink2 | 0.643815 | 0.251992 |
| Agk | 0.889383 | -0.250974 |
| Oxnad1 | 1.114948 | -0.249445 |
| Lgi1 | 0.469277 | -0.249348 |
| Rara | 0.943911 | 0.249178 |
| Slc9a4 | 0.435605 | 0.249142 |
| Zfand4 | 0.48345 | 0.248656 |
| BC048403 | 0.572494 | 0.248073 |
| Gpi1 | 1.336779 | 0.248 |
| Chchd10 | 0.341908 | -0.246299 |
| Ppid | 0.41258 | -0.24608 |
| Gda | 1.342383 | 0.245471 |
| Ptpro | 0.533359 | -0.245033 |
| Arntl | 0.287304 | 0.244984 |
| Spsb3 | 0.482912 | 0.244838 |
| Lmod2 | 0.290976 | 0.24206 |
| Mospd1 | 1.381986 | 0.241181 |
| Ferd3l | 0.951345 | 0.24073 |
| Fads2 | 0.448818 | -0.2404 |
| Slfn8 | 0.494953 | -0.239936 |
| Rab27b | 0.922222 | 0.239312 |
| Kifap3 | 1.480092 | 0.238298 |
| Plekhd1 | 0.588173 | -0.237967 |
| Tfcp2 | 0.595018 | -0.236793 |
| Yipf1 | 0.982278 | 0.235764 |
| E130309D02Rik | 1.031472 | 0.235592 |
| Ciita | 0.74572 | -0.234673 |
| Gm1965 | 1.499824 | -0.234366 |
| Wdr46 | 0.370338 | 0.234244 |
| Gm13238 | 1.671168 | -0.233827 |
| Ap4b1 | 0.680174 | 0.233348 |
| Ankib1 | 1.445116 | 0.231629 |
| Gm9930 | 1.001772 | -0.231285 |
| Ppp2cb | 1.540567 | 0.231101 |
| Il2ra | 1.356461 | 0.230953 |
| Hddc3 | 1.475684 | -0.230842 |
| Traf5 | 0.757864 | -0.229908 |
| Fam198b | 0.587988 | -0.229379 |
| Psmc1 | 0.892933 | 0.229046 |
| Foxi3 | 0.779251 | -0.228973 |
| Hsdl1 | 0.796725 | 0.227914 |
| Slc39a14 | 1.645953 | 0.227778 |
| Fbxo34 | 0.77279 | 0.227273 |

| | | |
|---|---|---|
| Cadm2 | 0.577116 | -0.227261 |
| Sfrp2 | 0.962498 | 0.226866 |
| Scn4a | 0.693022 | 0.226496 |
| Rsg1 | 0.789847 | 0.226299 |
| Slc7a11 | 0.213271 | 0.226237 |
| Wdr60 | 1.08727 | 0.225929 |
| Heatr3 | 1.225823 | -0.22546 |
| Vmn1r45 | 1.350166 | -0.225337 |
| Pcdh15 | 0.522608 | -0.224645 |
| Dab1 | 1.013059 | -0.224188 |
| Ltbr | 1.957937 | 0.223645 |
| Gorasp1 | 1.555471 | -0.223101 |
| Oit1 | 0.814747 | -0.223089 |
| Glod5 | 1.015511 | -0.223002 |
| Gm5124 | 0.878555 | -0.222112 |
| Parvb | 1.137398 | 0.221815 |
| Olfr67 | 1.048897 | 0.221357 |
| Mertk | 0.454436 | 0.221085 |
| Rnf113a2 | 0.569129 | 0.220999 |
| Pgs1 | 0.728844 | 0.220974 |
| S1pr5 | 0.991391 | 0.22059 |
| Zkscan8 | 0.852924 | -0.220578 |
| Nlgn3 | 0.925601 | -0.22002 |
| Kctd12b | 0.331018 | -0.219946 |
| Pdk1 | 0.250253 | 0.219463 |
| Bgn | 1.429379 | 0.218843 |
| 2510002D24Rik | 1.033107 | -0.21867 |
| Zfp622 | 1.051636 | 0.218347 |
| Rmnd1 | 1.061151 | -0.218124 |
| Aldh5a1 | 0.728632 | -0.217566 |
| Ptpn4 | 0.850125 | -0.217206 |
| Hsp90aa1 | 0.618784 | -0.216635 |
| Ccdc39 | 0.908421 | -0.21656 |
| Pkmyt1 | 1.034133 | -0.216436 |
| 3300002I08Rik | 1.158856 | 0.216225 |
| Fancb | 0.333288 | -0.214672 |
| Zfp948 | 0.516293 | -0.214635 |
| Gcfc2 | 0.721585 | -0.214411 |
| Ampd1 | 1.567005 | 0.214137 |
| Pabpc5 | 1.139051 | 0.213864 |
| Fzd3 | 0.728979 | 0.213677 |
| Ino80b | 0.762225 | 0.21359 |
| Mvd | 0.683667 | 0.212669 |

| | | |
|---|---|---|
| Mad2l2 | 1.11243 | -0.211685 |
| Snx25 | 0.918782 | -0.211237 |
| Pcdhac1 | 0.9537 | -0.209803 |
| Cd38 | 0.768189 | 0.209803 |
| Zfp30 | 0.278401 | -0.209354 |
| Nlrx1 | 0.598269 | 0.209179 |
| Cntnap1 | 0.592183 | -0.208779 |
| Mapt | 0.842694 | -0.208767 |
| 4933405O20Rik | 0.978426 | 0.208255 |
| Olfr488 | 0.422802 | 0.207905 |
| Cacng6 | 0.748778 | 0.207306 |
| Tra2a | 0.314668 | -0.206693 |
| Usp40 | 1.090161 | -0.206331 |
| Rbm22 | 1.239239 | 0.205993 |
| Clcn3 | 0.850892 | 0.205905 |
| Acat1 | 1.373209 | -0.205693 |
| Atp5o | 0.749161 | -0.205655 |
| Plaa | 0.622666 | 0.20553 |
| Bivm | 0.829545 | 0.205005 |
| Olfr1469 | 1.380068 | 0.204892 |
| Fgf21 | 1.125627 | 0.204842 |
| Piga | 0.798021 | 0.204817 |
| Slc9a1 | 1.561273 | 0.204554 |
| Ankrd42 | 0.671164 | -0.204128 |
| Gab2 | 0.799612 | 0.204041 |
| Olfr384 | 1.475004 | -0.204003 |
| Trafd1 | 0.965825 | 0.203915 |
| Gm8994 | 1.323401 | 0.203878 |
| Prrxl1 | 1.001431 | 0.203389 |
| Rras2 | 1.45682 | 0.202863 |
| Gtf2h3 | 0.480004 | -0.202487 |
| Slc2a12 | 0.788242 | -0.202386 |
| Actg1 | 0.925897 | -0.202374 |
| D3Ertd751e | 0.684191 | -0.201383 |
| Wfdc3 | 0.731017 | -0.20132 |
| Gm5801 | 0.671612 | 0.201195 |
| Nsun4 | 2.240393 | -0.201119 |
| Pnkd | 0.974776 | -0.200831 |
| Myl6b | 1.028693 | -0.20068 |
| Scgb1b7 | 1.426973 | -0.200655 |
| Msto1 | 0.907034 | -0.200504 |
| Rxrb | 0.480689 | 0.200316 |
| Tmem248 | 0.939922 | 0.199236 |

| | | |
|---|---|---|
| Pea15a | 0.496719 | -0.198758 |
| Zfp626 | 0.88943 | -0.198268 |
| Ccdc96 | 0.926755 | 0.196443 |
| Il6ra | 0.306142 | 0.195965 |
| Tmprss7 | 1.234225 | 0.1956 |
| Bri3bp | 0.893952 | 0.195272 |
| Sectm1a | 0.81632 | -0.195007 |
| Wdr62 | 0.807413 | 0.194995 |
| Napg | 1.872419 | -0.194932 |
| Ifngr1 | 0.910275 | 0.193974 |
| Htra4 | 0.456677 | 0.193709 |
| Tmem5 | 0.403981 | 0.193519 |
| Igsf9 | 1.133185 | -0.193494 |
| Prrc1 | 0.382722 | 0.192737 |
| Slc4a2 | 0.782737 | -0.192434 |
| Tdp2 | 0.924282 | -0.192371 |
| Olfr1394 | 0.656168 | -0.192308 |
| Homer3 | 0.934062 | -0.191689 |
| Kbtbd12 | 0.384242 | -0.191032 |
| Zfp92 | 0.423304 | -0.190956 |
| Samd8 | 0.826753 | 0.190425 |
| G0s2 | 0.743352 | -0.190021 |
| Pfn3 | 0.610609 | -0.189173 |
| Lta4h | 1.088052 | -0.188566 |
| Kif21a | 0.542995 | -0.188186 |
| Tac1 | 0.574332 | -0.188008 |
| 4921530L21Rik | 0.823171 | 0.187844 |
| Dalrd3 | 1.192194 | -0.187641 |
| Sytl2 | 0.440995 | -0.187464 |
| Mboat4 | 0.897378 | 0.187172 |
| Itch | 0.675547 | 0.187134 |
| 1700123L14Rik | 0.585418 | -0.18702 |
| Riok3 | 0.373771 | 0.186855 |
| Tmem14a | 0.741557 | -0.18683 |
| Palb2 | 0.707571 | 0.186412 |
| Pdia5 | 1.026498 | 0.186323 |
| Stxbp3 | 0.600626 | 0.185714 |
| Smim1 | 0.677055 | -0.185651 |
| Fbxo46 | 0.92541 | 0.185613 |
| T2 | 0.569053 | -0.185029 |
| Ubxn6 | 0.512717 | 0.18461 |
| Unc13a | 0.619814 | 0.184433 |
| Mmd | 1.057646 | -0.184356 |

| | | |
|---|---|---|
| Arfip1 | 1.034567 | 0.184141 |
| Fbxw15 | 0.941665 | 0.183734 |
| Fam187b | 0.422164 | 0.183353 |
| Slc36a3 | 0.609033 | -0.183201 |
| Cggbp1 | 0.89935 | 0.182718 |
| Snx7 | 0.452719 | 0.182641 |
| Wfdc1 | 0.517292 | -0.18226 |
| Sfxn3 | 1.691357 | 0.181115 |
| Tctex1d1 | 0.941415 | -0.18095 |
| Cend1 | 0.431234 | 0.180784 |
| Slc48a1 | 0.498312 | 0.180059 |
| Hpcal1 | 0.817882 | 0.179944 |
| Kcp | 0.703076 | -0.17988 |
| Ppp1r3d | 1.052274 | -0.179384 |
| G6bos | 0.736392 | -0.178696 |
| Masp1 | 0.244287 | 0.178224 |
| Raph1 | 0.647335 | -0.178211 |
| Vwa8 | 0.659679 | -0.177969 |
| Gm6401 | 0.724799 | 0.177382 |
| Gtpbp3 | 0.651849 | -0.177293 |
| C1ra | 0.561757 | 0.176859 |
| Rfk | 1.214459 | 0.176667 |
| Rbm43 | 0.583621 | -0.176284 |
| Fut8 | 0.562914 | 0.17594 |
| Mettl6 | 0.530381 | 0.175748 |
| 4933434E20Rik | 1.044699 | -0.175454 |
| 1700016D08Rik | 0.91018 | -0.175403 |
| Psme2b | 0.556737 | -0.174956 |
| Uso1 | 0.865931 | 0.174905 |
| Zfp958 | 0.466277 | 0.174841 |
| Gmpr | 0.624296 | -0.174828 |
| Cytip | 0.456895 | 0.174163 |
| Eif4a2 | 0.216097 | -0.173575 |
| 1700049L16Rik | 0.542108 | 0.173473 |
| Zfp286 | 0.752692 | -0.173076 |
| Zfp551 | 0.524901 | -0.172999 |
| Thoc3 | 0.818039 | 0.172449 |
| Gm17404 | 0.359768 | -0.172385 |
| Rwdd2a | 0.213654 | 0.171975 |
| Mad2l1bp | 0.88316 | 0.171899 |
| Nt5c1b | 0.307627 | 0.171873 |
| 1700029J07Rik | 0.298311 | -0.171373 |
| Bnc2 | 0.466369 | 0.171232 |

| | | |
|---|---|---|
| Zfp60 | 0.460665 | -0.170348 |
| Ttc38 | 0.357648 | -0.170271 |
| H2afy3 | 0.774518 | -0.169745 |
| Yipf3 | 0.775832 | 0.169476 |
| Rdx | 1.396472 | 0.168116 |
| Tm2d3 | 0.473202 | 0.167743 |
| Rnf122 | 0.395005 | -0.167692 |
| Cluh | 0.472628 | -0.167191 |
| Zfp747 | 0.860121 | 0.16714 |
| Rabac1 | 1.461304 | 0.166651 |
| Prss41 | 0.527744 | -0.166561 |
| Engase | 0.408047 | -0.166266 |
| Pvrl4 | 0.3939 | -0.166137 |
| Naa16 | 0.385565 | -0.16561 |
| A930033H14Rik | 0.693286 | -0.165417 |
| Lonrf2 | 0.56652 | -0.165378 |
| Slamf1 | 0.201506 | 0.165198 |
| Cycs | 0.766659 | -0.165159 |
| Polr3f | 0.361348 | 0.165018 |
| Grik2 | 0.561151 | -0.164992 |
| Serpina3i | 0.880978 | 0.164979 |
| Hsd11b2 | 1.181785 | -0.164966 |
| Top1 | 0.438707 | -0.164761 |
| Rpap3 | 0.905344 | 0.164529 |
| F2rl2 | 0.196619 | 0.164284 |
| Vwc2 | 0.692971 | -0.164246 |
| Ddx51 | 1.047777 | -0.164078 |
| Gnl3 | 0.26147 | 0.163975 |
| Adrbk1 | 0.526135 | 0.163628 |
| Rab2a | 0.824711 | 0.162996 |
| Yae1d1 | 1.291109 | -0.162945 |
| Rarres1 | 0.790321 | 0.162661 |
| Tnfrsf13c | 0.736007 | -0.162623 |
| Vars2 | 0.644296 | -0.162507 |
| Epn1 | 0.484526 | 0.162133 |
| Apex1 | 0.676412 | -0.161926 |
| Gm136 | 0.924818 | 0.161862 |
| Wdr48 | 0.523131 | -0.161759 |
| Aagab | 0.645033 | 0.161088 |
| Nus1 | 0.482117 | 0.160959 |
| Tubb1 | 0.240683 | 0.160881 |
| Notch2 | 0.388533 | 0.160662 |
| Cxcr2 | 0.095576 | 0.160533 |

| | | |
|---|---|---|
| Vipr1 | 0.517608 | -0.160443 |
| Tmem120b | 0.496672 | -0.160133 |
| Clec1b | 0.143953 | 0.159926 |
| Taok3 | 0.325244 | -0.159797 |
| Arf2 | 0.412693 | 0.158259 |
| G3bp1 | 0.623357 | -0.158001 |
| Magea8 | 0.425958 | 0.1576 |
| Clp1 | 0.573703 | 0.157432 |
| Gli1 | 0.40074 | 0.156863 |
| Acad8 | 1.08968 | -0.156526 |
| Ctage5 | 0.440369 | 0.156448 |
| Reep1 | 0.349048 | -0.156189 |
| Aif1l | 0.536997 | 0.155995 |
| Tmem115 | 0.464588 | 0.155943 |
| Fam101a | 0.601633 | -0.155697 |
| Ogdh | 0.929881 | -0.155684 |
| Drd2 | 0.310338 | -0.155607 |
| C1qtnf2 | 0.770062 | -0.155568 |
| Gm10197 | 0.335652 | 0.155425 |
| Suclg2 | 1.474775 | -0.155127 |
| Smyd3 | 0.614539 | 0.155127 |
| Impad1 | 0.293828 | 0.155115 |
| Slc25a15 | 0.263553 | 0.154946 |
| Ccdc122 | 0.125324 | 0.154467 |
| Adra1a | 0.516888 | 0.153909 |
| Xpo6 | 0.578263 | 0.153818 |
| Pkp2 | 0.820897 | 0.153766 |
| Anxa9 | 0.423349 | 0.153702 |
| Aatf | 0.401322 | 0.153624 |
| Fkbp4 | 0.423337 | -0.153611 |
| Ccdc142 | 0.68597 | -0.152936 |
| Trap1 | 0.766628 | -0.152651 |
| Slc45a2 | 0.542029 | -0.152534 |
| Pex12 | 0.520462 | 0.151846 |
| Rab35 | 0.907943 | 0.151573 |
| Synpo | 0.278997 | 0.151521 |
| Drg2 | 0.854415 | 0.151144 |
| Tcl1b4 | 0.534156 | 0.150937 |
| Ppbp | 0.143538 | -0.150885 |
| Lancl2 | 0.797291 | -0.150612 |
| Ankrd13b | 0.922712 | -0.150417 |
| Adra1b | 0.314989 | 0.149923 |
| Tead4 | 0.482787 | -0.149624 |

| | | |
|---|---|---|
| Pqbp1 | 0.77428 | 0.148986 |
| Mier1 | 0.721617 | 0.148609 |
| Map2k3 | 0.561303 | 0.148127 |
| Cyc1 | 0.603485 | -0.148101 |
| Usp35 | 0.630124 | 0.14771 |
| Gm5460 | 0.649452 | -0.14745 |
| Diablo | 0.580599 | -0.147359 |
| Slc9a8 | 1.371379 | -0.147176 |
| Tbc1d10c | 0.699307 | -0.147059 |
| Wdr73 | 0.37712 | 0.146994 |
| D930015E06Rik | 0.310108 | 0.146668 |
| 4932429P05Rik | 0.301742 | 0.14629 |
| Tecrl | 0.316214 | -0.145234 |
| Tex261 | 0.482813 | 0.144986 |
| Gm8674 | 0.635582 | 0.144908 |
| Chchd3 | 0.866218 | -0.144647 |
| Prdx6 | 0.279722 | 0.142975 |
| Vav2 | 0.438573 | -0.142675 |
| H13 | 0.338566 | 0.142609 |
| AW551984 | 0.418056 | 0.142505 |
| Wdr83 | 1.3631 | -0.142492 |
| Cort | 0.557171 | -0.1424 |
| Plekho2 | 0.686677 | -0.142322 |
| Pigp | 0.612187 | -0.142322 |
| Zfp654 | 0.634142 | 0.142309 |
| Arrdc3 | 0.262882 | 0.142034 |
| Mis18a | 0.313695 | 0.141917 |
| Cox20 | 0.630185 | -0.141877 |
| Ube2d1 | 1.158039 | 0.141786 |
| Dnd1 | 0.63181 | -0.141786 |
| Dcp2 | 0.496645 | -0.141537 |
| Mt3 | 0.482993 | 0.141498 |
| Bnip3l | 0.921275 | 0.141459 |
| Fam25c | 0.582724 | 0.141067 |
| Slc5a3 | 0.317341 | -0.140739 |
| Lgals2 | 0.310627 | -0.140556 |
| Ccdc190 | 0.655392 | -0.140203 |
| Gm3633 | 0.425827 | -0.139784 |
| Rab43 | 0.235552 | 0.139312 |
| Ifrd2 | 0.438597 | -0.139155 |
| Treml2 | 0.314563 | -0.13905 |
| Rasal1 | 0.76722 | -0.138854 |
| Gtf2f1 | 1.43972 | 0.138697 |

| | | |
|---|---|---|
| Ccdc185 | 0.459029 | -0.138474 |
| Nthl1 | 0.723276 | 0.138461 |
| Iglon5 | 0.546391 | -0.138369 |
| Rpl31 | 0.305422 | -0.138015 |
| Ddi2 | 0.903584 | 0.137936 |
| Ubqln1 | 0.564501 | 0.1377 |
| Gpalpp1 | 0.491821 | 0.137569 |
| Smap2 | 0.797419 | 0.137071 |
| Gm10330 | 0.365801 | 0.136979 |
| Olfr380 | 1.781231 | 0.136769 |
| 1700052K11Rik | 0.617531 | -0.136323 |
| 4931408C20Rik | 0.447067 | 0.136283 |
| Slc39a9 | 1.709709 | 0.135601 |
| Olfr1471 | 0.328139 | -0.135574 |
| Lyg1 | 0.771392 | -0.135535 |
| Sox9 | 0.529609 | 0.135075 |
| Adam3 | 0.233546 | -0.134431 |
| Ap5z1 | 0.607292 | -0.134313 |
| Olfr643 | 0.565666 | -0.134287 |
| Armc10 | 0.564433 | 0.134195 |
| Kirrel3 | 0.550417 | -0.134168 |
| Shkbp1 | 0.323047 | 0.133958 |
| Hinfp | 0.463578 | -0.1338 |
| 4933411K16Rik | 0.722978 | -0.133695 |
| Krt36 | 0.417197 | -0.133603 |
| Ky | 0.312461 | -0.133603 |
| Gm8113 | 0.434002 | -0.13359 |
| Dock7 | 0.196423 | 0.133458 |
| Slbp | 1.715955 | -0.133024 |
| Cebpz | 0.472437 | 0.132932 |
| Omp | 0.754189 | -0.132801 |
| Tmem128 | 0.229518 | 0.132406 |
| Cdyl | 0.763443 | 0.131537 |
| Nlrp4f | 0.450867 | -0.131194 |
| Cited2 | 0.245555 | 0.130944 |
| Pcbp2 | 0.636903 | 0.130865 |
| Tars | 0.48101 | 0.129375 |
| Tfip11 | 0.431214 | 0.129112 |
| Dnajc7 | 0.566517 | -0.128689 |
| Cmas | 0.452582 | 0.128637 |
| Wfikkn2 | 0.401084 | 0.128584 |
| Rwdd1 | 0.490272 | 0.128518 |
| Fbxl15 | 0.689776 | 0.127871 |

| | | |
|---|---|---|
| Ctf1 | 0.490371 | -0.127805 |
| Celsr2 | 0.415504 | -0.127343 |
| Rad1 | 0.702485 | 0.127263 |
| D6Ertd527e | 0.328869 | 0.127171 |
| Znhit2 | 0.503263 | 0.127092 |
| Olfr985 | 0.360615 | 0.126986 |
| Nono | 1.800508 | -0.126722 |
| Tax1bp1 | 0.399598 | 0.126484 |
| Srrm3 | 0.48107 | 0.125916 |
| Ddit4l | 0.339147 | -0.125902 |
| AA474331 | 0.247312 | 0.125453 |
| Gzma | 0.132048 | -0.125387 |
| Gm3618 | 0.383958 | -0.125056 |
| Mob3c | 0.598146 | -0.125016 |
| Spats1 | 0.597893 | 0.124368 |
| Brat1 | 0.748824 | -0.124116 |
| Pira6 | 0.322363 | -0.12409 |
| Spock1 | 0.43593 | -0.123984 |
| Tdh | 0.543258 | -0.123759 |
| Slc33a1 | 0.352285 | 0.123468 |
| Vegfb | 0.71201 | -0.123282 |
| Etnk2 | 0.784815 | -0.122686 |
| Lrrc40 | 0.331461 | -0.12254 |
| Gm8325 | 1.282332 | 0.122434 |
| Iqcc | 0.579604 | -0.122315 |
| Vmn2r38 | 0.328415 | -0.122275 |
| Apba3 | 0.765485 | 0.122156 |
| Galnt13 | 0.453429 | 0.122143 |
| Prkar1a | 1.20067 | 0.121851 |
| Mkl1 | 0.372332 | 0.121745 |
| D930007J09Rik | 0.551736 | -0.12144 |
| Bcl3 | 0.263221 | 0.121347 |
| Mt2 | 0.141936 | 0.120816 |
| Utp3 | 0.550727 | 0.120657 |
| Arl8a | 0.44502 | 0.120644 |
| Ctla2a | 1.069661 | 0.120511 |
| Sel1l | 0.227434 | 0.120471 |
| Mtg1 | 0.363893 | -0.120365 |
| Rab3d | 0.537015 | 0.120179 |
| Gm6498 | 0.723354 | 0.119794 |
| Kcnj10 | 0.556696 | -0.119688 |
| Ap5s1 | 0.635213 | 0.119662 |
| Phf10 | 0.293741 | 0.119516 |

| | | |
|---|---|---|
| Mxi1 | 0.339324 | 0.119489 |
| Sfrp5 | 0.217494 | -0.11921 |
| Me2 | 0.307184 | 0.118878 |
| Dixdc1 | 0.338074 | 0.118719 |
| Prkci | 0.364234 | 0.118134 |
| Trem1 | 0.080917 | 0.118041 |
| Pfdn2 | 0.997406 | 0.117974 |
| Neurod6 | 0.489805 | -0.117881 |
| 45356 | 0.374159 | 0.117841 |
| Khdrbs3 | 0.195147 | -0.117775 |
| Dmrtc1a | 0.45541 | -0.117629 |
| Dcun1d1 | 0.423963 | 0.117629 |
| Krtap22-2 | 0.453456 | -0.11719 |
| Gm10081 | 0.312015 | 0.116711 |
| Wif1 | 0.438635 | 0.116591 |
| Thrb | 0.478964 | -0.116458 |
| Mus81 | 0.398919 | -0.116178 |
| 3110082J24Rik | 1.170083 | -0.116032 |
| Dnal1 | 0.303038 | -0.115979 |
| Gm13152 | 0.142447 | -0.115912 |
| Rexo4 | 0.433149 | -0.115819 |
| B230217C12Rik | 0.289196 | -0.115686 |
| Lzts3 | 0.302826 | -0.115619 |
| Wdr5b | 0.494278 | -0.114993 |
| Pld1 | 0.267138 | 0.114967 |
| Angpt4 | 0.310475 | 0.114953 |
| Arl5c | 0.210037 | -0.11482 |
| Shoc2 | 1.186514 | 0.114767 |
| Lrrc23 | 0.297243 | -0.114727 |
| Tnk1 | 0.367631 | 0.114167 |
| Gm2837 | 0.273305 | -0.113767 |
| Alg13 | 0.225514 | -0.113274 |
| Apof | 0.330864 | 0.11262 |
| Tmub1 | 0.597116 | 0.11222 |
| Gm8246 | 0.212279 | -0.11206 |
| Lrp10 | 0.553579 | 0.111859 |
| Plod1 | 0.39444 | 0.110924 |
| Aph1b | 0.380049 | 0.110136 |
| Spryd4 | 0.330273 | 0.109842 |
| Gm5129 | 0.377592 | -0.109762 |
| Mettl9 | 0.584476 | 0.109668 |
| Hdac9 | 0.251448 | -0.109267 |
| Yy2 | 0.511466 | 0.109173 |

| | | |
|---|---|---|
| Prss54 | 0.2788 | -0.109173 |
| Olfr483 | 0.31518 | 0.109093 |
| Znhit6 | 0.959354 | 0.108772 |
| Olfr1415 | 0.28632 | -0.108357 |
| 4933402J07Rik | 0.389071 | 0.108277 |
| Pdhb | 0.317274 | -0.108277 |
| Rbm10 | 0.309407 | -0.10817 |
| Chsy1 | 0.466246 | 0.108009 |
| Radil | 0.626283 | 0.107902 |
| Tor3a | 0.548651 | 0.107728 |
| Gm11360 | 0.465823 | -0.107648 |
| Ndrg3 | 0.544335 | 0.107541 |
| Tcte2 | 0.460943 | -0.107313 |
| Mrps9 | 0.47704 | 0.106991 |
| Tnfrsf19 | 0.357932 | 0.106791 |
| Osm | 0.182708 | 0.106375 |
| Osgepl1 | 0.226979 | -0.106348 |
| Sh2d6 | 0.29855 | -0.106335 |
| Iqcj | 0.495295 | -0.106268 |
| Ube2d2b | 0.400958 | 0.106201 |
| Pdzd4 | 0.243108 | 0.106201 |
| Fkbp3 | 0.429464 | -0.106187 |
| Map3k2 | 0.253566 | 0.106107 |
| Wfdc6b | 0.522228 | -0.105986 |
| Xpnpep2 | 0.353486 | -0.105785 |
| Agtr1b | 0.747387 | -0.105329 |
| Atf7ip2 | 0.499171 | -0.105168 |
| 1700086D15Rik | 0.324835 | 0.104712 |
| Itfg2 | 0.330784 | -0.10439 |
| Faap20 | 0.934215 | -0.104323 |
| Prr30 | 0.553516 | -0.103867 |
| Prpf18 | 0.412779 | -0.103719 |
| Tmem125 | 0.616091 | 0.103437 |
| Mfsd9 | 0.218015 | 0.103195 |
| Vps51 | 0.3851 | -0.103128 |
| Ubl7 | 0.4952 | 0.102806 |
| Phactr3 | 0.363871 | 0.102658 |
| Stat6 | 0.623516 | 0.101946 |
| Krt73 | 0.388857 | 0.101919 |
| Chst2 | 0.352691 | -0.101825 |
| Akap17b | 0.268814 | 0.101704 |
| Mettl18 | 0.526874 | 0.101408 |
| Wdyhv1 | 0.292696 | 0.101274 |

| | | |
|---|---|---|
| Urm1 | 0.654828 | 0.101112 |
| Ccdc71 | 0.573702 | 0.100762 |
| Mpp6 | 0.154823 | 0.100547 |
| Ecsit | 0.456854 | -0.100453 |
| Idh3b | 0.337933 | 0.100359 |
| Cybrd1 | 0.12927 | -0.10013 |
| Vstm4 | 0.240576 | -0.100009 |
| Dennd4c | 0.291404 | 0.099995 |
| Igdcc4 | 0.363653 | 0.099982 |
| Ppm1g | 0.592344 | 0.099861 |
| Ccdc157 | 0.443077 | 0.099699 |
| Pitx1 | 0.436327 | -0.099686 |
| Zfand2b | 0.265564 | 0.099376 |
| Samd9l | 0.212144 | -0.099363 |
| Ndufs1 | 0.270837 | -0.099309 |
| 2700062C07Rik | 0.219511 | -0.098932 |
| Ncstn | 0.300057 | 0.098783 |
| Gm21863 | 0.590894 | 0.098581 |
| Gm3182 | 0.193711 | -0.098447 |
| Scnm1 | 0.476399 | -0.098258 |
| Gm5901 | 0.209956 | -0.098244 |
| A930017M01Rik | 0.232115 | -0.097948 |
| Pcdh7 | 0.122411 | 0.097907 |
| Saxo2 | 0.220773 | 0.097638 |
| 1700003F12Rik | 0.255325 | 0.097557 |
| 9330159N05Rik | 0.313087 | -0.097314 |
| Fcamr | 0.337961 | -0.096936 |
| Vmn1r41 | 0.266062 | 0.096842 |
| Serpina11 | 0.475797 | -0.096815 |
| Car10 | 0.337395 | -0.096451 |
| Smco1 | 0.198559 | 0.096437 |
| Olfr1038-ps | 0.218568 | 0.096329 |
| Plrg1 | 0.830073 | 0.095776 |
| Thap7 | 0.644524 | 0.095344 |
| Ptbp3 | 0.68628 | 0.094993 |
| Eef2k | 0.349045 | 0.094858 |
| Ier5l | 0.308739 | -0.094736 |
| Gnb2 | 0.938642 | 0.094668 |
| Knop1 | 0.514242 | 0.094317 |
| 1110002L01Rik | 0.335527 | -0.09356 |
| Gm3182 | 0.207564 | -0.09356 |
| Stat5b | 0.391377 | -0.093479 |
| Gm9611 | 0.24873 | -0.093411 |

| | | |
|---|---|---|
| Ache | 0.390557 | 0.091842 |
| Ovgp1 | 0.390424 | 0.091815 |
| Sry | 0.325608 | 0.091693 |
| Chst14 | 0.523394 | 0.091544 |
| Ccl11 | 0.251978 | -0.091192 |
| Tecr | 0.59544 | -0.091111 |
| 4931414P19Rik | 0.253167 | 0.091002 |
| Fgf8 | 0.259714 | 0.090948 |
| Mcur1 | 0.45743 | -0.090217 |
| Mob4 | 0.357363 | 0.090149 |
| Urah | 0.294868 | 0.090095 |
| Pcsk1 | 0.367357 | -0.089688 |
| Tssk3 | 0.316274 | 0.089593 |
| Pias1 | 0.376244 | 0.089552 |
| Etv6 | 0.81325 | 0.089539 |
| Nt5c3 | 0.129234 | -0.089498 |
| Cyp2a4 | 0.324254 | -0.089471 |
| Ankrd44 | 0.298662 | -0.089078 |
| Zfp647 | 0.35226 | 0.08863 |
| Rabif | 0.594415 | 0.088535 |
| Atp11a | 0.231673 | -0.088345 |
| Armc2 | 0.216705 | -0.088196 |
| Id2 | 0.230233 | 0.087762 |
| Fam134a | 0.756082 | 0.087639 |
| Aar2 | 0.345338 | 0.087558 |
| Macrod1 | 0.150951 | -0.087422 |
| Cep192 | 0.302056 | -0.087395 |
| Pwp2 | 0.262506 | 0.087341 |
| Cyp2g1 | 0.368197 | 0.087313 |
| Marcksl1 | 0.563205 | -0.087042 |
| Dpp10 | 0.172516 | -0.086906 |
| Park2 | 0.357886 | -0.086784 |
| Chmp4b | 0.287136 | 0.08677 |
| Mir339 | 0.262369 | 0.086471 |
| Brwd3 | 0.326128 | 0.086281 |
| Zc2hc1b | 0.248119 | 0.085996 |
| Nol11 | 0.269826 | 0.085833 |
| Mtr | 0.268687 | 0.085737 |
| Cdc37l1 | 0.331967 | -0.085425 |
| Rrp7a | 0.256242 | 0.08537 |
| Gm10339 | 0.207166 | -0.084881 |
| Gpr84 | 0.172277 | 0.084799 |
| Pex3 | 0.194447 | 0.084595 |

| | | |
|---|---|---|
| Sebox | 0.244111 | -0.084541 |
| Tnfrsf1b | 0.232367 | -0.084132 |
| Nfyb | 0.281663 | 0.083955 |
| Cd209c | 0.206289 | 0.083438 |
| Anapc4 | 0.281044 | 0.083329 |
| Nomo1 | 0.5184 | -0.083316 |
| Nt5m | 0.114413 | 0.083016 |
| Tm9sf3 | 0.365045 | 0.082743 |
| Nanos3 | 0.228556 | -0.082362 |
| Ss18l1 | 0.401285 | -0.082267 |
| Fank1 | 0.55684 | 0.082089 |
| Plekhf1 | 0.438801 | 0.081912 |
| Ptms | 0.766933 | -0.081858 |
| Fmn2 | 0.327531 | -0.081858 |
| Gucy2g | 0.187838 | -0.081749 |
| Btbd10 | 0.198534 | -0.081721 |
| Tmem25 | 0.264226 | -0.081585 |
| Evx1 | 0.341071 | 0.081381 |
| Cox7c | 1.063156 | -0.081258 |
| Cx3cl1 | 0.196332 | -0.081149 |
| Pla2r1 | 0.309835 | -0.08104 |
| Dnttip2 | 0.528427 | 0.080767 |
| Dnah7b | 0.263439 | -0.08074 |
| Tmem251 | 0.152503 | 0.080712 |
| Pigo | 0.413147 | -0.080194 |
| Mpp4 | 0.289504 | -0.079798 |
| C030039L03Rik | 0.562952 | -0.079566 |
| Gm7932 | 0.355212 | -0.079566 |
| Scgb1b21 | 0.305066 | 0.07947 |
| Hagh | 0.12542 | 0.079402 |
| Smu1 | 0.246016 | -0.079197 |
| Ninl | 0.318004 | -0.079006 |
| Gm3127 | 0.26423 | -0.079006 |
| Adgrg6 | 0.108938 | 0.079006 |
| Krt16 | 0.295408 | 0.078938 |
| Col6a6 | 0.16043 | -0.078664 |
| Ikbkap | 0.271662 | 0.07835 |
| Flot2 | 0.175499 | 0.078337 |
| Uroc1 | 0.237864 | -0.078036 |
| Usp44 | 0.276255 | 0.078022 |
| Swt1 | 0.271863 | -0.077407 |
| Myh14 | 0.265467 | 0.077393 |
| Brix1 | 0.171728 | 0.076997 |

| | | |
|---|---|---|
| Crbn | 0.382303 | 0.076983 |
| Mdfic | 0.482163 | -0.076846 |
| Dpysl5 | 0.207073 | -0.076833 |
| Zfp449 | 0.153047 | -0.075765 |
| Hdac1 | 0.393816 | -0.075697 |
| Ivd | 0.152371 | -0.075683 |
| Slu7 | 0.461674 | 0.075409 |
| Wwp2 | 0.445988 | 0.075012 |
| Frem1 | 0.327384 | -0.074999 |
| Flg | 0.338862 | 0.074615 |
| Ccdc137 | 0.252369 | -0.074204 |
| Tbpl1 | 0.305828 | 0.074122 |
| Map1s | 0.303583 | 0.074108 |
| Tnnc2 | 0.259138 | 0.073779 |
| Mdga1 | 0.273679 | 0.073382 |
| Efcc1 | 0.307083 | 0.073148 |
| Try5 | 0.207823 | -0.071996 |
| Rimbp3 | 0.211738 | -0.071927 |
| Cep120 | 0.233221 | 0.071859 |
| Ccdc87 | 0.261725 | -0.071818 |
| 1700025C18Rik | 0.187396 | -0.071708 |
| Azin2 | 0.083021 | 0.07157 |
| Osbpl9 | 0.488339 | 0.071543 |
| Exosc6 | 0.20624 | -0.071529 |
| Gpx6 | 0.339884 | -0.071488 |
| Emc1 | 0.414097 | 0.071447 |
| Avp | 0.34257 | -0.071337 |
| Wdr26 | 0.279375 | 0.06995 |
| Prr22 | 0.22196 | -0.06995 |
| Obox6 | 0.189761 | 0.069661 |
| Dmc1 | 0.28557 | -0.069592 |
| Lrrc15 | 0.302378 | 0.069455 |
| Kif2a | 0.261285 | 0.069097 |
| Ppp1r21 | 0.212431 | 0.069097 |
| Elf1 | 0.488166 | 0.068863 |
| Fam207a | 0.179154 | 0.068244 |
| Zbtb7b | 0.326637 | 0.068079 |
| Rnf138rt1 | 0.183356 | -0.068065 |
| Clu | 0.156138 | -0.06801 |
| Mapk6 | 0.083972 | -0.067955 |
| Fam73b | 0.194728 | -0.0679 |
| Eef2kmt | 0.285392 | -0.067818 |
| F630003A18Rik | 0.197915 | 0.06746 |

| | | |
|---|---|---|
| 4930444P10Rik | 0.265411 | -0.067088 |
| Olfr1121 | 0.252448 | -0.066909 |
| Actl11 | 0.172663 | -0.066757 |
| Dpcr1 | 0.218693 | -0.066744 |
| Ak8 | 0.153071 | 0.066565 |
| Cyp1a1 | 0.149144 | -0.066234 |
| Mmadhc | 0.320492 | 0.066082 |
| Dnajc14 | 0.400891 | 0.065834 |
| Neu1 | 0.170285 | 0.065738 |
| Ncln | 0.22314 | 0.065531 |
| 2310033P09Rik | 0.287247 | -0.06549 |
| Smim20 | 0.097234 | 0.065421 |
| D10Wsu102e | 0.161895 | 0.065159 |
| Siglecg | 0.235317 | -0.064773 |
| Vta1 | 0.26879 | 0.064455 |
| Apobec3 | 0.145119 | -0.064345 |
| Cwc27 | 0.270468 | 0.064124 |
| Frmd4b | 0.157943 | 0.063738 |
| Mcm8 | 0.282085 | 0.063655 |
| Emc7 | 0.175083 | 0.063531 |
| Rad52 | 0.260515 | -0.063282 |
| Dleu7 | 0.192889 | -0.062909 |
| Tex22 | 0.173447 | -0.062771 |
| Pafah2 | 0.24428 | 0.06273 |
| Cemip | 0.051784 | 0.062274 |
| Atp8a1 | 0.179584 | -0.062232 |
| Tox | 0.193371 | 0.06179 |
| Pdc | 0.189537 | -0.061652 |
| Csnk2b | 0.49989 | 0.061624 |
| Sgpp1 | 0.155758 | 0.061486 |
| AI607873 | 0.115409 | 0.061348 |
| Kat5 | 0.478578 | 0.061334 |
| Rundc1 | 0.24086 | 0.061002 |
| Ranbp9 | 0.178016 | 0.060808 |
| Dazap2 | 0.300378 | 0.06049 |
| Ffar3 | 0.189839 | -0.060227 |
| D930048N14Rik | 0.163301 | -0.060103 |
| Catip | 0.151827 | -0.059425 |
| Zfp865 | 0.197758 | -0.059411 |
| Snip1 | 0.277104 | -0.0593 |
| Cldn34b4 | 0.166943 | -0.058871 |
| Tuft1 | 0.112852 | 0.058746 |
| Erbb4 | 0.150577 | -0.058538 |

| | | |
|---|---|---|
| Bpifa3 | 0.21563 | -0.058455 |
| Kif3b | 0.144234 | 0.058316 |
| Gm5134 | 0.182586 | -0.057596 |
| Lrrc8d | 0.178924 | 0.057263 |
| Rbbp8nl | 0.118658 | -0.057083 |
| Mutyh | 0.155613 | -0.056681 |
| Krtap31-1 | 0.148449 | 0.05632 |
| Rab21 | 0.17214 | 0.056264 |
| Ptprj | 0.086741 | -0.056264 |
| Zfp599 | 0.361638 | -0.055959 |
| Mterf1a | 0.193806 | 0.055459 |
| Asnsd1 | 0.080244 | 0.055459 |
| P2ry12 | 0.058225 | -0.05539 |
| Mr1 | 0.153587 | -0.055196 |
| Cep78 | 0.09629 | 0.054973 |
| Wdr92 | 0.155341 | -0.05496 |
| Gm10936 | 0.235139 | 0.054682 |
| Ndnl2 | 0.221021 | 0.054487 |
| Olfr1276 | 0.067187 | 0.054237 |
| Crtap | 0.24282 | 0.054043 |
| Npnt | 0.123289 | -0.053973 |
| Npy5r | 0.175068 | 0.053959 |
| Pgm3 | 0.135669 | -0.053904 |
| Ccdc90b | 0.235346 | -0.053709 |
| Idnk | 0.207091 | -0.053695 |
| Eif3c | 0.348204 | 0.053654 |
| Uqcrb | 0.223956 | -0.053654 |
| Tbx19 | 0.169005 | -0.053626 |
| Cdc5l | 0.145421 | -0.053125 |
| Uhrf1bp1l | 0.117505 | -0.052527 |
| Pklr | 0.233657 | 0.052485 |
| Gtf3c4 | 0.188122 | -0.052388 |
| Neu2 | 0.166291 | 0.052388 |
| Padi4 | 0.023359 | 0.05236 |
| Slc35a4 | 0.312109 | 0.052319 |
| Tab2 | 0.276391 | 0.052277 |
| Epm2a | 0.156675 | -0.052277 |
| Samhd1 | 0.142456 | -0.051804 |
| Invs | 0.173737 | 0.051581 |
| Meaf6 | 0.155679 | 0.051525 |
| 4930544G11Rik | 0.215649 | -0.051163 |
| Slc35e2 | 0.15622 | -0.050606 |
| Sec23b | 0.220639 | 0.050397 |

| | | |
|---|---|---|
| 4930539E08Rik | 0.147099 | -0.05023 |
| Naa30 | 0.206682 | 0.049366 |
| Fig4 | 0.195957 | 0.049366 |
| Myl12a | 0.321648 | -0.04924 |
| Kremen2 | 0.150978 | -0.049157 |
| Hand2 | 0.192298 | 0.048989 |
| Alg2 | 0.19072 | 0.048934 |
| Ipmk | 0.198836 | 0.04885 |
| Ulk3 | 0.208202 | -0.048655 |
| Ndufa5 | 0.198213 | -0.048571 |
| Slc6a9 | 0.212374 | 0.048432 |
| Tada1 | 0.323594 | -0.048208 |
| Kcnk3 | 0.104469 | -0.048152 |
| Prkcb | 0.104049 | -0.048041 |
| Slc38a7 | 0.158779 | -0.048027 |
| Sprr2d | 0.10405 | 0.047832 |
| Zfp772 | 0.107486 | 0.047706 |
| Sap30bp | 0.164983 | -0.047524 |
| Xcl1 | 0.177891 | 0.047343 |
| Tox3 | 0.114501 | -0.046882 |
| Mia | 0.132823 | 0.046868 |
| Tceb3 | 0.326503 | 0.046589 |
| Adam10 | 0.224956 | 0.046589 |
| Tex15 | 0.208688 | -0.046533 |
| Slc25a17 | 0.193911 | 0.046491 |
| Slc1a7 | 0.203601 | -0.046449 |
| Pkm | 0.127703 | 0.046393 |
| Pck2 | 0.165408 | 0.046058 |
| Homez | 0.060593 | 0.045946 |
| Olfr632 | 0.089772 | 0.04575 |
| Sat1 | 0.118757 | 0.045639 |
| Plb1 | 0.177301 | -0.045597 |
| 9030624G23Rik | 0.099854 | 0.045177 |
| Tspan17 | 0.195741 | 0.045107 |
| Ccl27b | 0.163488 | 0.045038 |
| Ccl27b | 0.163488 | 0.045038 |
| Prp2 | 0.16587 | -0.044982 |
| Plod3 | 0.121812 | 0.044926 |
| Lurap1 | 0.129991 | -0.04487 |
| F13b | 0.28802 | -0.044772 |
| Adss | 0.090569 | 0.044716 |
| Cfap157 | 0.189192 | 0.044618 |
| Pld6 | 0.104586 | -0.04452 |

| | | |
|---|---|---|
| 5031439G07Rik | 0.10256 | -0.044492 |
| Fgfr4 | 0.12667 | 0.04445 |
| Msl3 | 0.346212 | 0.044002 |
| Skida1 | 0.144554 | 0.043666 |
| Fhad1 | 0.1181 | -0.043582 |
| A330041J22Rik | 0.136538 | -0.043457 |
| Kank4 | 0.084883 | 0.043457 |
| Olfr571 | 0.089582 | -0.043429 |
| Rps16 | 0.569752 | 0.043373 |
| Src | 0.119609 | 0.043106 |
| Lyzl6 | 0.129073 | -0.043022 |
| Ythdf2 | 0.173696 | -0.042994 |
| 4930555F03Rik | 0.115605 | 0.04291 |
| Rac3 | 0.094645 | 0.042812 |
| Cdca4 | 0.174638 | -0.042756 |
| Klhl36 | 0.088079 | 0.042672 |
| Acvr1 | 0.091013 | 0.042336 |
| Tlr5 | 0.056121 | -0.042182 |
| Zfp365 | 0.098575 | 0.042112 |
| Emc3 | 0.136073 | 0.041944 |
| Tapbp | 0.214739 | -0.041565 |
| Gfra4 | 0.142806 | -0.041467 |
| Ptpru | 0.233007 | 0.041159 |
| Pdrg1 | 0.156871 | -0.040836 |
| Adh6-ps1 | 0.137896 | -0.04071 |
| 1700026L06Rik | 0.162436 | 0.040682 |
| Rnf135 | 0.089183 | -0.040682 |
| Klk15 | 0.101126 | -0.040654 |
| Zfp108 | 0.140533 | -0.040514 |
| Cox15 | 0.108982 | -0.040233 |
| Mtmr14 | 0.141523 | 0.040191 |
| Ei24 | 0.204507 | 0.040149 |
| Lrrc29 | 0.095539 | -0.040135 |
| Snrnp40 | 0.127623 | -0.039967 |
| Mapkapk2 | 0.130651 | 0.03977 |
| Dpagt1 | 0.213994 | -0.039742 |
| Gcsam | 0.069165 | 0.039616 |
| Fdxacb1 | 0.061573 | 0.039209 |
| Nsa2 | 0.104239 | 0.038858 |
| Ywhaz | 0.314645 | 0.038844 |
| Cd151 | 0.134693 | 0.038408 |
| Sapcd1 | 0.142384 | 0.038169 |
| Ankrd27 | 0.101763 | 0.037902 |

| | | |
|---|---|---|
| Pced1b | 0.18114 | -0.03772 |
| E130304I02Rik | 0.161117 | 0.037621 |
| Ubqln2 | 0.146787 | -0.037593 |
| Nfxl1 | 0.159281 | -0.036989 |
| Abcf3 | 0.142282 | -0.036974 |
| Srr | 0.124043 | -0.036848 |
| Gm6880 | 0.071767 | -0.036848 |
| Pigk | 0.106977 | 0.036412 |
| Akt1s1 | 0.278838 | 0.036356 |
| Ogfod3 | 0.084072 | 0.036145 |
| Gm17535 | 0.047686 | -0.035948 |
| Dnajc8 | 0.132597 | 0.035863 |
| Hoxa11os | 0.125348 | -0.035427 |
| Slit3 | 0.10527 | 0.035413 |
| Ralb | 0.185802 | -0.03523 |
| Ccdc24 | 0.209215 | -0.035188 |
| Vmn2r37 | 0.076654 | -0.034568 |
| Irf2bp2 | 0.115394 | 0.034469 |
| Pop4 | 0.06235 | 0.034399 |
| Smtnl1 | 0.121865 | -0.034371 |
| Pf4 | 0.049311 | 0.034357 |
| Serpina9 | 0.070476 | -0.034286 |
| Gsta4 | 0.056112 | 0.03399 |
| 4930423O20Rik | 0.090784 | -0.033441 |
| Olfr620 | 0.109583 | 0.033243 |
| Nrde2 | 0.108683 | 0.033187 |
| Zfp341 | 0.259373 | -0.032919 |
| Fut9 | 0.110394 | -0.032877 |
| Atp5g1 | 0.071542 | 0.032425 |
| Olfr247 | 0.07108 | 0.032341 |
| Fnip1 | 0.057203 | -0.03227 |
| Hcn1 | 0.05451 | -0.032242 |
| Olfr449 | 0.074113 | 0.032185 |
| 2310061I04Rik | 0.090225 | 0.032171 |
| Gtf2a1l | 0.140446 | 0.032143 |
| Eif2ak1 | 0.062638 | 0.031875 |
| Aifm3 | 0.132418 | -0.031847 |
| Fer1l6 | 0.153254 | 0.031635 |
| Srprb | 0.099546 | -0.03155 |
| Strn3 | 0.092536 | -0.031536 |
| Senp2 | 0.081296 | 0.031409 |
| 45545 | 0.075264 | -0.031381 |
| Cdc42se2 | 0.230252 | 0.030421 |

| | | |
|---|---|---|
| Fbxo28 | 0.089133 | 0.030223 |
| Cpz | 0.159778 | -0.02994 |
| Fgf16 | 0.077318 | 0.029842 |
| Gtf2a1 | 0.139775 | 0.029785 |
| BC089597 | 0.100846 | -0.029728 |
| Ypel1 | 0.091127 | 0.0297 |
| Fem1c | 0.143792 | 0.029418 |
| Rhox3f | 0.105774 | 0.029418 |
| Slc25a36 | 0.085614 | 0.029333 |
| Olfr76 | 0.117928 | 0.029262 |
| Gm13212 | 0.051109 | -0.029149 |
| Gm14501 | 0.095157 | -0.029121 |
| Zeb2 | 0.077535 | 0.028866 |
| Sphk2 | 0.098019 | -0.028597 |
| Gm26571 | 0.09082 | 0.028541 |
| 4930415O20Rik | 0.083797 | -0.028343 |
| Irf6 | 0.046845 | -0.028343 |
| Smagp | 0.11191 | 0.028315 |
| Ltc4s | 0.048288 | 0.028258 |
| Porcn | 0.206841 | 0.027947 |
| Smarca5-ps | 0.101754 | -0.027777 |
| Prg3 | 0.088886 | 0.027678 |
| Klk1b27 | 0.077879 | -0.027664 |
| Tmem175 | 0.069881 | -0.027649 |
| Krtap19-3 | 0.117225 | -0.027494 |
| Erich5 | 0.091379 | 0.027423 |
| Alx3 | 0.075238 | -0.02731 |
| Abhd10 | 0.132391 | -0.027126 |
| Usf2 | 0.071007 | 0.027126 |
| Scn2b | 0.065869 | 0.027126 |
| Klk1b22 | 0.051549 | -0.026998 |
| Gm10775 | 0.061186 | -0.02697 |
| Sp9 | 0.087751 | -0.026588 |
| Angptl8 | 0.063445 | -0.026474 |
| Lce1f | 0.070318 | -0.026276 |
| Slc39a5 | 0.084657 | -0.026148 |
| Srcin1 | 0.048197 | 0.026134 |
| Tfe3 | 0.077363 | -0.025964 |
| Cyp4f15 | 0.074244 | -0.025936 |
| Creld1 | 0.086866 | -0.025497 |
| Mphosph9 | 0.074996 | 0.025454 |
| Vmn1r42 | 0.038872 | 0.025369 |
| Snd1 | 0.08159 | 0.024703 |

| | | |
|---|---|---|
| Olfr192 | 0.107922 | 0.024632 |
| Speer4b | 0.0603 | 0.02449 |
| E130112N10Rik | 0.088811 | -0.02432 |
| 9930022D16Rik | 0.071879 | 0.024079 |
| Fpgs | 0.11389 | -0.024008 |
| Gm14180 | 0.064405 | -0.023993 |
| Dhx36 | 0.103209 | 0.023596 |
| Irx2 | 0.053315 | 0.023454 |
| Abcb7 | 0.0509 | -0.02344 |
| Rbm12b1 | 0.046069 | -0.023298 |
| Smim5 | 0.085105 | -0.023113 |
| Usp45 | 0.064871 | -0.023099 |
| Olfr1511 | 0.087917 | 0.023071 |
| Gldc | 0.072914 | 0.023028 |
| 5730522E02Rik | 0.060512 | 0.023 |
| Abce1 | 0.0678 | -0.022886 |
| Pnma2 | 0.058407 | -0.022758 |
| Ccl2 | 0.034253 | 0.022673 |
| Olfr753-ps1 | 0.050193 | -0.022219 |
| Mcts2 | 0.058499 | 0.021295 |
| Ankrd28 | 0.074781 | 0.021195 |
| Klhl40 | 0.041353 | 0.021139 |
| Plpp7 | 0.096631 | 0.021096 |
| Gm21560 | 0.032799 | -0.020669 |
| Odam | 0.073076 | -0.020655 |
| Id4 | 0.116129 | 0.020456 |
| Fubp3 | 0.067723 | -0.020271 |
| Gm9140 | 0.070609 | 0.019787 |
| Nphp1 | 0.085405 | 0.019645 |
| Slc7a6os | 0.061423 | 0.019588 |
| Olfr312 | 0.051638 | -0.019261 |
| Ccdc181 | 0.092731 | -0.019047 |
| Hoxb3 | 0.072525 | 0.018933 |
| Gm9969 | 0.052043 | 0.018307 |
| Lrrd1 | 0.04594 | 0.018235 |
| Elf4 | 0.054128 | 0.018022 |
| Gm20346 | 0.063104 | 0.017965 |
| Trnt1 | 0.033754 | 0.01795 |
| Mfap3 | 0.039115 | 0.01768 |
| Cfl2 | 0.080239 | 0.017665 |
| Gm10251 | 0.036163 | -0.017452 |
| Nkx2-5 | 0.055433 | -0.017281 |
| Olfr458 | 0.065979 | -0.017238 |

| | | |
|---|---|---|
| St3gal4 | 0.062586 | -0.017109 |
| Cntfr | 0.04313 | -0.016525 |
| Efr3a | 0.054797 | -0.016396 |
| Pram1 | 0.015389 | 0.016282 |
| Tbcc | 0.09155 | 0.016254 |
| Zbtb1 | 0.045063 | -0.016026 |
| Ttc12 | 0.026977 | -0.015854 |
| Prap1 | 0.048085 | 0.015583 |
| Dusp2 | 0.031532 | 0.015512 |
| Rragc | 0.023389 | 0.015398 |
| 6430571L13Rik | 0.052449 | -0.015355 |
| Wapl | 0.040419 | 0.015355 |
| Olfr536 | 0.045619 | -0.015341 |
| Thoc7 | 0.057745 | -0.015241 |
| Rps6 | 0.117562 | 0.015126 |
| Cybb | 0.028843 | -0.015084 |
| Epgn | 0.031663 | -0.014855 |
| Trim61 | 0.035206 | 0.014412 |
| Gm1979 | 0.037034 | -0.014327 |
| Foxp4 | 0.074974 | -0.01417 |
| Hyal1 | 0.073159 | 0.014027 |
| Cd226 | 0.009785 | -0.01397 |
| Gm21800 | 0.026755 | -0.013798 |
| H2-M5 | 0.038474 | -0.013641 |
| Aoc2 | 0.032114 | -0.013512 |
| 4930550C14Rik | 0.038505 | -0.013341 |
| Lhfpl1 | 0.047487 | -0.013069 |
| Mybbp1a | 0.063626 | 0.01284 |
| Skap1 | 0.032384 | -0.012726 |
| Tfpt | 0.058441 | 0.011925 |
| Olfr747 | 0.035121 | 0.011753 |
| Morc3 | 0.026291 | -0.011539 |
| Gm13889 | 0.039282 | 0.011438 |
| Cfap206 | 0.035098 | 0.011424 |
| Sox30 | 0.035811 | -0.011152 |
| Hspb2 | 0.017421 | 0.011038 |
| Bves | 0.017295 | 0.01088 |
| Cand1 | 0.031024 | 0.010766 |
| Cxcl11 | 0.020769 | -0.010536 |
| Phb2 | 0.041881 | 0.009978 |
| Uqcrc2 | 0.039018 | -0.00972 |
| Wdr7 | 0.027071 | 0.009333 |
| Reck | 0.00872 | -0.009232 |

| | | |
|---|---|---|
| Jmjd7 | 0.022036 | 0.009204 |
| Panx2 | 0.020894 | -0.008946 |
| Spag6l | 0.038683 | 0.008874 |
| Csmd1 | 0.028441 | 0.008128 |
| Sall4 | 0.020907 | 0.008028 |
| Scfd1 | 0.025784 | 0.00797 |
| 4932416K20Rik | 0.020107 | -0.007827 |
| Faxc | 0.024565 | -0.007511 |
| Maf | 0.014385 | 0.007483 |
| Olfr495 | 0.012284 | -0.00721 |
| Coq2 | 0.018086 | -0.00698 |
| Chst12 | 0.015733 | 0.006478 |
| Rbm12 | 0.026806 | -0.006434 |
| Chchd7 | 0.01928 | -0.006434 |
| Vsir | 0.016812 | -0.006276 |
| Dffa | 0.019659 | 0.006061 |
| Micu2 | 0.011425 | 0.00586 |
| Gm6356 | 0.016326 | -0.005788 |
| Acap3 | 0.018161 | 0.005716 |
| Mesp2 | 0.017012 | -0.005616 |
| 1700028J19Rik | 0.02356 | -0.005601 |
| Serpinb8 | 0.006243 | 0.005299 |
| Plcxd3 | 0.01329 | -0.005184 |
| Map3k4 | 0.014045 | -0.004954 |
| Tbc1d13 | 0.013441 | 0.004796 |
| Mphosph8 | 0.013589 | 0.004566 |
| Hspa12a | 0.011627 | 0.004523 |
| Kcnb1 | 0.016755 | 0.004465 |
| Rpl13a | 0.051852 | 0.004394 |
| Gm9945 | 0.011014 | 0.004106 |
| H2-M3 | 0.010474 | -0.004005 |
| Gadd45a | 0.011271 | -0.003991 |
| Fkbp15 | 0.01142 | -0.003847 |
| Jph4 | 0.009377 | -0.003588 |
| Rnf186 | 0.009035 | 0.003545 |
| Amph | 0.007998 | -0.00353 |
| Socs4 | 0.007284 | -0.003314 |
| Sipa1l3 | 0.009288 | -0.003228 |
| Egr3 | 0.008495 | -0.00294 |
| Tmem41b | 0.014786 | 0.002148 |
| Zbtb9 | 0.003807 | 0.001788 |
| Stam2 | 0.004395 | -0.001615 |
| Krtap4-9 | 0.005314 | 0.001572 |

| | | |
|---|---|---|
| Dvl2 | 0.006092 | -0.001283 |
| Calca | 0.001787 | -0.000808 |
| Sbk2 | 0.002021 | -0.000779 |
| Olfr1219 | 0.00118 | -0.000505 |
| Gm9970 | 0.001183 | -0.000375 |
| Teddm3 | 0.001168 | -0.000361 |
| Micu1 | 0.000266 | -0.000101 |
| Dcaf12 | 0.000233 | -5.77E-05 |
