## Supplementary material for "Lymphatic activation of ACKR3 signaling regulates lymphatic response after ischemic heart injury": Suplementary Methods

### Supplementary Methods

#### *Mice*

Ackr3-TangoGFP transgenic mice were generated by the Animal Models Core Facility at the University of North Carolina at Chapel Hill (UNC-CH). For the generation of Ackr3-Tango, Tango elements were inserted into the *Ackr3* endogenous locus by CRISPR/Cas9-mediated genome editing in C57BL/6J strain. Guide RNAs were cloned into a plasmid donor vector flanking the Tango elements: 1) a codon-optimized tetracycline-controlled transactivator (tTA) with preceding TEV protease cleavage site; 2) a P2A "self-cleaving" peptide sequence; 3) a Beta-arrestin-2- tobacco etch virus (TEV) protease fusion protein; 4) 3X stop codons, FRT site, and loxP-flanked SV40 late polyadenylation sequences. C57BL/6J zygotes were microinjected with 800 nM HiFiCas9 protein, 50 ng/ul guide RNA and 20 ng/ul supercoiled donor vector plasmid. Injected embryos were implanted in recipient pseudopregnant females. The resulting pups were screened by PCR for the presence of the knock-in allele. Founder animals were mated to C57BL/6J-Rosa26-CAG-Flpo transgenic animals to collapse tandem integration events to single-copy knock-ins. F1 animals heterozygous for the correct knock-in event and absence of a vector backbone sequence were used for subsequent breeding to establish the knock-in colony. To genotype the *Ackr3*-Tango knock-in allele, CAT CCC GTT TAC CTG TCA GC forward and CAT AAG AGC CTG TCC TGG TGC reverse primers were used. For H2B-GFP genotyping, GCT CGT TTA GTG AAC CGT CAG forward and TCT TCT GCG CCT TAG TCA CC reverse primers were used. In order to abolish undesired H2B transcription, mice were kept on doxycycline supplemented chow (Envigo, TD.01306). To turn on H2B transcription, mice were kept on normal chow for 7 days before LAD ligation.

*Ackr3*$^{fl/fl}$ and *Ackr3*$^{fl/fl}$ ; Lyve1$^{Cre}$ (Ackr3ΔLyve1) mice were maintained on a C57BL/6 background. For *Ackr3*$^{fl}$ genotyping, TCG GGA GAG GAT TTG GAG TGC TTC forward and TGA AAT CAG CAT GAT ACA GGG TCC reverse primers were used. For genotyping of the constitutive Lyve1 cre allele, TGC CAC CTG AAG TCT CTC CT wild type forward, GAG GAT GGG GAC TGA AAC TG cre forward and TGA GCC ACA GAA GGG TTA GG common reverse primers were used.

*Prox1*$^{GFP}$ mice (generously provided by Young-Kwon Hong) were maintained on 129S/SvEv background by mating heterozygous mice to wild type 129S/SvEv mice. Expression of the Prox1-GFP allele was confirmed by UV illumination of GFP.

For all experiments, 4-8 month old animals were used. Ischemic cardiac injury experiments were conducted on male mice. Animals were housed in SPF housing with ad libitum access to food and water. The mice were provided with environmental enrichment in the form of nesting material and lofts. The mice had a 12h:12h light-dark cycle, with a temperature range between 68-74°F and humidity between 30-70%. All animal studies were approved by the Institutional Animal Care and Use Committee at the University of North Carolina Chapel Hill.

#### *Surgical induction of myocardial infarction*

Mice were anesthetized with intraperitoneal 100mg/kg ketamine and 10mg/kg xylazine, intubated and ventilated (Harvard Apparatus) using 100% O2 at 0.2 L/min and 1% isoflurane. A left lateral thoracotomy was performed between the 3rd and 4th rib.

Following pericardiectomy, the left anterior descending artery (LAD) was ligated at its midpoint with 7-0 suture. Successful ligation was confirmed by tissue blanching and myocardial hypokinesia. Chest wall, muscle and skin were closed with 5-0 suture. Intramuscular cefazolin (50 mg/kg) and subcutaneous buprenorphine (0.1 mg/kg) were administered, and recovery was under a heat lamp. Animals that did not survive permanent LAD ligation surgery or died before recovery from anesthesia were not included in survival data and were excluded from further analysis. Additional subcutaneous buprenorphine (0.1 mg/kg) injection was administered on the first day after surgery. Mice were monitored at least daily until the end of experiment. Ackr3-Tango-GFP mice were euthanized 2-days post LAD ligation, while *Ackr3*$^{fl/fl}$ and *Ackr3*$^{\Delta Lyve1}$ were euthanized 3-, 6-, or 28-days post ligation.

***Histology and immunohistochemistry***

Mice were euthanized with CO2 asphyxiation and trans-cardiac perfusion was performed with 10 mL phosphate-buffered saline (PBS) followed by 10 mL 4% paraformaldehyde (PFA). Dissected organs were fixed in 4% PFA for 24h at 4°C, and then embedded in paraffin for sectioning and staining following standard protocols by the UNC-CH Histology Research Core Facility. Sections were collected 200μm from the beginning of the infarct zone towards the apex of the heart. For immunohistochemistry, paraffin sections were deparaffinized, hydrated, and antigen retrieval using heated sodium citrate buffer (10mM sodium citrate, 0.05% Tween-20 (Fisher Scientific BP337-500), pH 6.0), was performed. Tissues were then permeabilized with 0.2% Triton-X in PBS for 20 minutes and blocked with 5% normal donkey serum (NDS, Jackson ImmunoResearch, 017-000-121) in PBS for 1h at RT. Slides were then stained with primary antibodies including, rabbit anti-LYVE1 (1:200, Fitzgerald, 70R-LR005), goat anti-CD45 (1:50, Novus Biologicals, NB100-77417), and goat anti-F4/80 (1:50, BD Pharminogen, 565409) for 24h at 4°C. To ensure antibody specificity, sections incubated with secondary only antibodies were used as negative controls. Sections were rinsed and incubated with secondary antibodies, made up in 5% NDS in PBS, including, donkey anti-rabbit Cy3 (1:200, Jackson ImmunoResearch, 711-165-152), donkey anti-goat Cy5 (1:200, Jackson ImmunoResearch, 705-175-147), and 1:1000 Bisbenzimide H 33258 (Hoechst) (SigmaAldrich, B1155). Slides were rinsed, mounted with ProLong Gold Antifade Mounting Media (ThermoFisher Scientific, P10144), and coverslips were sealed with clear nail polish. Stained slides were imaged using a Nikon Eclipse Ti2 inverted widefield microscope with automated scanning for large area acquisition equipped with either a Nikon DS-Fi3 CMOS color camera for brightfield imaging or a pco.edge 4.2Q High QE sCMOS fluorescent camera for fluorescent imaging using 4x or 10x air objectives. For quantitative analysis of the images, ImageJ was used.

***Whole mount immunostaining***

Mice were euthanized with CO2 asphyxiation and trans-cardiac perfusion was performed with 10 mL PBS followed by 10 mL 4% PFA. Dissected organs were fixed for 24h in 4% PFA at rocking at 4°C. Hearts were then immersed in 0.1% Triton X-100 in PBS overnight rocking at 4°C. Next, hearts were blocked in 1% BSA, 1% DMSO, 5% NDS (Jackson ImmunoResearch, 017-000-121) in 0.1% Triton X-100 in PBS overnight, rocking at 4°C. Hearts were then immersed in goat anti-LYVE1 IgG (1:200, R&D Systems,

AF2125-SP) antibody dissolved in in blocking solution. To ensure antibody specificity, sections reacting with secondary only antibodies were used as negative controls. Hearts were rinsed 3x for 15min with 1% BSA, 1% DMSO, 0.1% Tween-20 in PBS, then immersed in Alexa Fluor 488-conjugated donkey anti-goat (1:100, Jackson ImmunoResearch, 705-545-003) antibody dissolved in blocking solution. Fluorescent images of the hearts were captured using a Nikon Eclipse Ti2 inverted widefield microscope with automated scanning for large area acquisition equipped with a pco.edge 4.2Q High QE sCMOS fluorescent camera using a 4x air objective. To visualize lymphatic vessels of the whole heart, Extended Depth of Focus images were generated from Z-series of overlapping multipoint acquisition images in NIS Elements software program. For quantitative analysis of the images, ImageJ was used.

#### *Tissue clearing and whole mount immunostaining*

iDisco protocol was performed as described previously[12]. Samples were incubated with rabbit anti-LYVE1 (1:500, Fitzgerald, 70R-LR005) and chicken anti-GFP (1:500, Aves, GFP-1020) primary antibodies for 14 days and incubated with Cy5-conjugated donkey anti-rabbit (1:200, Jackson ImmunoResearch, 711-175-152) and Alexa Fluor 790-conjugated donkey anti-chicken (1:200, Jackson ImmunoResearch, 703-655-155) secondary antibodies for 10 days. For imaging, a LaVision Ultramicroscope II (LaVision) light-sheet system was used. Images were acquired using the Miltenyi Biotec imaging software and analyzed with the Imaris 9 (Oxford Instruments) software program. Surfaces and Spots tools in the Imaris software were used to identify LYVE1+ lymphatic vessels and Tango-GFP+ cells, respectively. Non-vascularized LYVE-1 structures were excluded from the rendering of lymphatic structures. Tango-GFP+ spheres were then clustered based co-localization with the identified lymphatic structures. Relative lymphatic GFP signal was calculated using the following formula: Relative lymphatic GFP signal = $[N(GFP)_{LYVE1+}/ N(GFP)_{LYVE1-}]_{injury\ site} / [N(GFP)_{LYVE1+}/ N(GFP)_{LYVE1+}]_{uninjured\ tissue}$. 3-D images capturing the entire volume of the indicated anatomical locations were quantified. One 3-D image per heart segment was evaluated in each mouse.

#### *Microarray analysis*

For transcriptomic analysis, lymphatic endothelial cells (LECs) and macrophages were collected from $Ackr3^{fl/fl}$ and $Ackr3^{\Delta Lyve1}$ mice at baseline or 7-days post LAD ligation. For this purpose, mice were euthanized with CO2 asphyxiation and trans-cardiac perfusion was performed with 10 mL PBS. The ventricles were then minced and incubated with 2 mg/mL collagenase type 2 (Worthington Biochemical, LS004176) in Hank's balanced salt solution (Gibco, 14025092) at 37°C to make a cell suspension. Cells were then centrifuged and incubated for 1h at 4°C with Dynabead Protein G (Thermo Fisher, 10003D) magnetic beads pre-treated with 1μg/μL anti-CD68 (BioRad, MCA1957) antibodies. RNA was extracted from the captured CD68-positive macrophages using Trizol (Invitrogen, 15596026) following the manufacturer's recommendations. CD68-negative cells were incubated for 1h at 4°C with magnetic beads pre-treated with 1μg/μL anti-LYVE1 (Fritzgerald, 70R-LR005) primary antibodies to collect CD68-negative/LYVE1-positive LECs and CD68/LYVE1 double negative cells, and RNA was extracted from both cell populations using the Trizol protocol. Microarray analysis was performed in the UNC-CH Functional Genomics Core using the mouse Clariom S Pico

Assay kit (Thermo Fisher, 902932) after QC/QA testing. Microarray data was evaluated using the Partek Genomics Suite (Partek) software program. For statistical analysis of the data, one-way ANOVA was performed, and false discovery rate-adjusted p values ≤0.01 were considered significant.

#### *Echocardiography*

Conscious echocardiography was performed using a Vevo 2100 ultra high frequency ultrasound (VisualSonics) on male *Ackr3*$^{\Delta Lyve1}$ and *Ackr3*$^{fl/fl}$ mice before, and 3-, 6-,14-, 21-, and 28-days post LAD ligation. M-mode echocardiographs of left ventricle were analyzed by measuring both diastolic and systolic dimensions of three cardiac cycles from three recordings using Vevo LAB software (VisualSonics). The nine measurements were averaged for each mouse on each experimental day. Stroke volume was calculated using the SV= LV Vol;d – LV Vol;s formula. Cardiac output was calculated using the CO=SV/HR formula.

#### *Cell culture, gene silencing, and hypoxia induction*

Primary human lymphatic endothelial cells (hLECs), isolated from human foreskin (PromoCell, C-12216) were cultured in Endothelial Cell Growth Medium (PromoCell, C-22111) 37°C under 5% CO2 and used within 5 passages. To silence *CALCRL* and *ACKR3* gene expression, hLECs were treated with siRNAs targeting *CALCRL* (Santa Cruz Biotechnology, sc-43705), ACKR3 (Santa Cruz Biotechnology, sc-94573) or scramble control siRNA (Santa Cruz Biotechnology, sc-37007) using Lipofectamine RNAiMAX transfection reagent (Thermo Scientific, 13778030) for 24h following the manufacturers' recommendations. To study hypoxia-induced changes in hLECs, cells were incubated at 1% or 21% O2 in a Tri-gas incubator for the indicated times.

#### *Immunocytochemistry and lymphatic junctional assay*

To assess hLEC junctional arrangement dynamics, confluent monolayers of were fixed in 4% paraformaldehyde for 20 minutes at RT, after siRNA treatment and hypoxia induction, as described above. After fixation, the cells were permeabilized for 2 minutes in 0.2% Tween-20 PBS solution. Samples were then incubated with PBS containing 5% NDS for 1h at RT followed by a 24h incubation with rabbit anti-VE-cadherin (Abcam, ab33168) and anti-VEGFR3 (R&D Systems, AF349) primary antibodies diluted 1:200 in PBS containing 5% NDS at 4°C. After PBS wash, cells were incubated with Cy5-conjugated donkey anti-rabbit (Jackson Immunoresearch, 711-175-152) and Alexa Fluor 488-conjugated donkey anti-goat (Jackson Immunoresearch, 705-545-003) secondary antibodies diluted 1:200 in PBS containing 5% NDS and 1:1000 Hoechst for 2h at RT, then washed with PBS and mounted in ProLong Gold antifade media (P36934, Life Technologies). Cells were imaged using a Zeiss 800 confocal microscope with a 20x air objective. Images were analyzed using ImageJ. Continuous junctions were determined as uninterrupted, convex cell border segments, while discontinuous cellular junctions are characterized by concave patterning. For quantitative analysis, we determined the total length of continuous and discontinuous cellular junction lengths per field of view from at least two images per biological replicate. The average percent ratio of total continuous junctions length among total junctions length (continuous + discontinuous junctions combined) of a multiple field of views from the same biological replicate was used for quantitative assessment of junction dynamics in these experiments.

#### *Adrenomedullin response assay and Western blot*

hLECs were transduced with unconcentrated lentiviral particles containing shRNA targeting RAMP3 (TRCN0000060981) or human beta globin (HBG) (TRCN0000029094) (negative control) (20% v/v and supplemented with 10 µg/mL polybrene) for 24h, after which, the virus-containing media was replaced with fresh LEC growth media. 48h after transduction, hLECs were serum starved overnight by replacing growth media with Opti-MEM reduced serum media (Gibco, 31985070). The next day, time-course experiments were performed by treating cells with 100 nM adrenomedullin from 0 to 60 minutes. To quench signaling at appropriate time-points, the media was quickly replaced with ice-cold 1x PBS, and then lysed in 50 mM Tris-HCl (pH 7.4) supplemented with protease (cOmpleteTM, EDTA-free Protease Inhibitor Cocktail; Roche, 11873580001), phosphatase (PhosSTOP EASYpack; Roche, 4906845001) inhibitors, and Benzonase® nuclease (Millipore, E1014). Protein quantification and normalization of cell lysates was accomplished using a PeirceTM BCA assay kit (ThermoFisher Scientific, 23225), after which, 4x LDS sample buffer (Invitrogen, NP0007) supplemented with 10% DTT (1M stock) was added to each sample. Samples were run on a 4-12% NuPAGE Bis Tris gel (ThermoFisher Scientific, NPO336BOX) using 1x MOPS-SDS Running buffer (Boston BioProducts, BP-178), and transferred to a nitrocellulose membrane using 2x Transfer Buffer (Boston BioProducts, BP-193) supplemented with 20% methanol. Membranes were blocked for 1h in 1x tris-buffered saline (TBS) + 0.1% Tween-20 + 5% BSA prior to incubation with the following primary and secondary antibodies: p-p44/42 MAPK (T202/Y204) (CST, 4370) (1:5,000), p44/42 MAPK (ERK1/2) (CST, 9102) (1:5,000), p-AKT (S473) (CST, 4060), AKT (pan) (CST, 4691) (1:5,000), p-CREB (Ser133) (CST, 9198) (1:5,000), CREB (CST, 9197) (1:5000), GAPDH (Novus Biologicals, NB300-221) (1:10,000), IRDye 800CW Goat anti-Rabbit (Licor, 926-32211) (1:10,000), and IRDye 680LT goat anti-mouse (Licor, 926-68020) (1:10,000). Data was collected on a Licor Odyssey CLx using Image Studio version 5.2 and densitometry performed using ImageJ software. For each independent time-course experiment, two gels were run: one to probe for phosphorylated proteins and the other to probe for total levels. GAPDH levels were determined on each blot as a load control. Relative protein levels were determined by: [(phosphorylation protein level divided by GAPDH) / (Total protein level divided by GAPDH)].

#### *Quantitative real time polymerase chain reaction*

To characterize gene expression levels in cultured hLECs or isolated mouse cells, RNA was extracted using Trizol protocol and cDNA was transcribed using M-MLV reverse transcriptase (Thermo Fisher, 28-025-013) following the manufacturer's protocol. Reverse transcription polymerase chain reaction was performed using the Fast Advanced Master Mix (Thermo Fisher, 4444557), according to the manufacturer's protocol utilizing a Thermo Fisher StepOnePlus real time PCR system. For human LECs, *ACTB*- (Thermo Fisher, Hs01060665_g1), *ACKR3*- (Thermo Fisher, Hs00664172_s1), *ADM*- (Thermo Fisher, Hs00969450_g1), *CALCRL*- (Thermo Fisher, Hs00907738_m1), and *CDH5*- (Thermo Fisher, Hs00901465_m1) specific probes were used. For mouse cells, *Gapdh*-

(Thermo Fisher, Mm99999915_g1), *Cd68*- (Thermo Fisher, Mm03047343_m1), *Lyve1*- (Thermo Fisher, Mm00475056_m1) specific probes were used.

#### *Statistical analyses*

Statistical analyses were performed with the GraphPad Prism 10 software program. Applied statistical tests and groups sizes are indicated for each analysis. P values ≤0.05 were considered significant and are shown in each graph. During analysis of microarray data, false discovery rate-adjusted p values ≤0.01 were considered significant. Data represented as mean ± standard deviation.
